# Systematic Tuning of Rhodamine Spirocyclization for Super-Resolution Microscopy

**DOI:** 10.1101/2021.05.20.444797

**Authors:** Nicolas Lardon, Lu Wang, Aline Tschanz, Philipp Hoess, Mai Tran, Elisa D’Este, Jonas Ries, Kai Johnsson

## Abstract

Rhodamines are the most important class of fluorophores for applications in live-cell fluorescence microscopy. This is mainly because rhodamines exist in a dynamic equilibrium between a fluorescent zwitterion and a non-fluorescent but cell-permeable spirocyclic form. Different imaging applications require different positions of this dynamic equilibrium, which poses a challenge for the design of suitable probes. We describe here how the conversion of the *ortho*-carboxy moiety of a given rhodamine into substituted acyl benzenesulfonamides and alkylamides permits the systematic tuning of the equilibrium of spirocyclization with unprecedented accuracy and over a large range. This allows to transform the same rhodamine into either a highly fluorogenic and cell-permeable probe for live-cell stimulated emission depletion (STED) microscopy, or into a spontaneously blinking dye for single molecule localization microscopy (SMLM). We used this approach to generate differently colored probes optimized for different labeling systems and imaging applications.

## Introduction

Continuous progress in fluorescence microscopy techniques and labeling strategies has led to an increased demand for suitable fluorescent probes.^1-4^ A prominent class of fluorophores are rhodamine derivatives.^5-6^ Their widespread use results from their high photostability and brightness, the broad spectral range they cover, as well as the possibility to tune these properties through synthetic modifications following established procedures.^1, 5, 7-18^ Another crucial feature of rhodamine-based dyes is the capability of spirocyclization.^19^ They exist in a dynamic equilibrium between a non-fluorescent, cell-permeable spirolactone and a fluorescent zwitterion (Figure 1). Binding of rhodamines to their targets often stabilizes the fluorescent zwitterion relative to the non-fluorescent spirolactone. Consequently, rhodamine-based probes can be fluorogenic, which reduces unspecific background signal.^20^ Such cell-permeable and fluorogenic probes are powerful tools for various live-cell microscopy applications.^3^ The spirocyclization equilibrium also plays a central role in the design of rhodamine-based probes for single-molecule localization microscopy (SMLM).^21^ SMLM techniques rely on the precise detection of individual fluorophores to construct super-resolved images.^2^ For this, only a small fraction of the dye population should be in the fluorescent state at the same time requiring the ability to shift the equilibrium strongly towards the spirocyclic form.^22-23^

**Figure 1.**
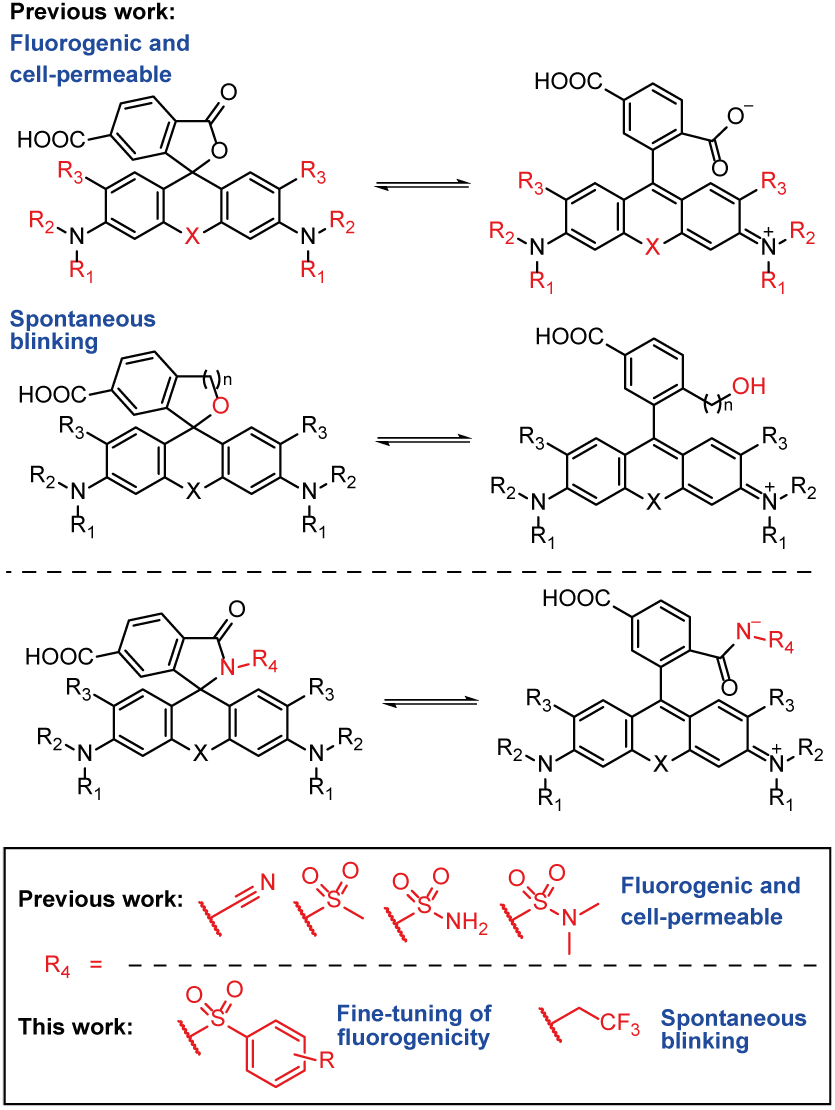
Structures of rhodamine-based fluorophores and their modifications to shift the equilibrium towards the spirocyclic state. Positions colored in red represent structural changes applied to the dye scaffold.

Up to now, strategies to increase cell-permeability and fluorogenicity by shifting the equilibrium towards the spirocyclic state were mainly based on modifications of the xanthene core. Exchanging the bridging oxygen of rhodamines with other functionalities (X) altered the state of the equilibrium and created fluorogenic and cell-permeable probes (Figure 1).^24-26^ However, this approach leads to rather drastic changes in the equilibrium.^12^ Furthermore, it also affects the photophysical properties of the fluorophore including strong shifts in absorbance and emission wavelengths.^27^ Other approaches relied on decreasing the electron density of the xanthene system, such as fluorination and introduction of 3-substituted azetidine moieties.^10-11, 28^ While fluorination of the xanthene core led to a decrease in quantum yield for certain fluorophores, incorporation of electron-deficient azetidines caused a distinct hypsochromic shift in absorbance and emission wavelength.^12, 17^ Xanthenes with reduced electron density are also known to display enhanced reactivity towards intracellular nucleophiles, which can decrease fluorophore brightness.^16, 29-30^ Furthermore, modification of the xanthene core often requires multi-step syntheses and the development of sophisticated methodologies.^8, 12, 31-32^

With respect to probes for SMLM techniques, stable rhodamine spirolactams were previously introduced which could be photoactivated to form a fluorescent state.^22, 33-35^ Another prominent approach is based on the replacement of the *ortho*-carboxy moiety with a more nucleophilic hydroxyl group (Figure 1).^28, 36-39^ The resulting probes can spontaneously switch between their preferred spirocyclic and opened form. Therefore, they exhibit blinking behavior without the need for intense UV irradiation and commonly used additives.^36^ Subsequently, spontaneous blinking also was reported for rhodamine spirolactams^40^ as well as its combination with photoactivation^41^.

We previously developed a general synthetic strategy to transform rhodamine-based dyes into highly fluorogenic and cell-permeable probes without changing their spectroscopic properties.^20^ This was achieved by simple conversion of the *ortho*-carboxy moiety into electron-deficient amides (Figure 1). Here, we show how the equilibrium of spirocyclization can be adjusted much more precisely and over a larger range. Incorporation of benzenesulfonamides with different substituents enabled the systematic fine-tuning of the spirocyclization equilibrium and the fluorogenicity of the resulting probes (Figure 1). Furthermore, incorporation of more electron-rich amides allowed the generation of spontaneously blinking dyes. Thus, the same rhodamine dye can be optimized for either applications in live-cell, no-wash STED imaging or SMLM without changing its spectroscopic properties.

## Results and Discussion

### Fine-Tuning of Rhodamine *500R* Spirocyclization

Our goal was to manipulate the spirocyclization equilibrium of a rhodamine-based scaffold such that it can be transformed into different probes that specifically match the requirements of imaging technique and labeling system. As our synthetic approach modifies the *ortho*-carboxy moiety instead of the xanthene core, we can maintain essential photophysical properties of the original dye. We used rhodamine *500R* (**1**), a green fluorophore with a high quantum yield, but a low tendency to form the spirolactone, as a platform to test this strategy.^11, 17^ We synthesized the basic scaffold of rhodamine *500R* (**2**) by means of Buchwald-Hartwig amination of **3**.^8^ Amidation of **2** followed by deprotection with TFA yielded a library of different rhodamine *500R* spirolactams (Figure 2b and Supplementary Figure 1).^42^ Chloroalkane (CA) and *O*^6^-benzylguanine (BG) ligands were coupled to the 6-carboxy group affording probes for HaloTag and SNAP-tag labeling (Supplementary Figure 1).

**Figure 2.**
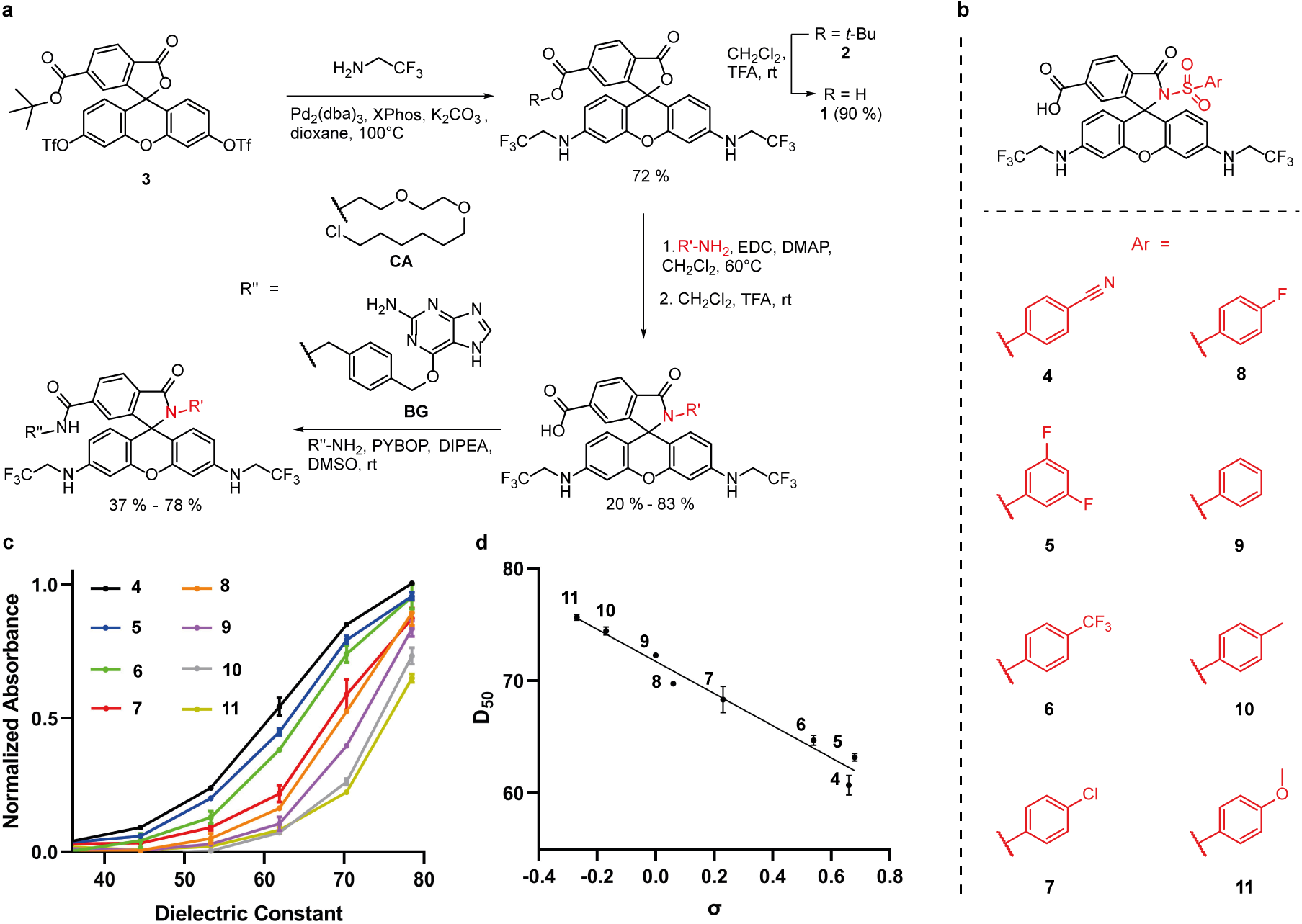
(a) Synthesis of rhodamine *500R* derivatives. (b) Structures of rhodamine *500R* benzenesulfonamide derivatives (**4** - **11**). (c) Normalized maximal absorbance of **4** - **11** (5 µM) in water– dioxane mixtures (v/v: 0/100 - 100/0) at 25°C as a function of the dielectric constant.^43^ The absorbance was normalized to the maximum absorbance of the ortho-carboxy dye **1**. (d) Correlation of D_50_ values versus Hammett constants (σ) of the substituents at the benzenesulfonamide.^44^ Solid line shows linear regression (R^2^ = 0.97). For disubstituted compound **5**, the σ of the substituent was doubled.^44-45^ Error bars show ± s.d. from 3 experiments.

The resulting rhodamine *500R* benzenesulfonamide derivatives (**4 - 11**) did not reveal distinct changes in absorbance and emission wavelengths as well as quantum yield relative to the parental dye, thus confirming that our derivatization strategy does not change the key spectroscopic properties of the underlying rhodamine scaffold (Figure 2b and Supplementary Table 1). Furthermore, we analyzed their spirocyclization behavior by recording the absorbance spectra in dioxane-water mixtures with different dielectric constants (Figure 2c). The D_50_ values derived from these experiments represent the dielectric constant at which half of the fluorophore population is in the opened form.^17, 20, 24^ Thus, they are a measure for the state of the spirocyclization equilibrium. We found that incorporation of various benzenesulfonamides enabled fine-tuning of this equilibrium with outstanding precision (Figures 2b and c). The linear correlation between D_50_ values and Hammett constants (σ) of the aromatic substituents corroborates the rational of our strategy (Figure 2d). Therefore, benzenesulfonamides bearing electron-donating substituents lead to a stronger shift towards the spirocyclic form than electron-deficient benzenesulfonamides.

### Optimizing Rhodamine *500R* for HaloTag Labeling

Self-labeling protein tags are useful tools to attach fluorophores to proteins in living cells.^4^ HaloTag is a widely used protein tagging-system and covalently binds to a synthetic CA ligand which can be conjugated to fluorophores.^46^ To obtain high contrast images without the need for washing-steps, a high fluorogenicity of the corresponding HaloTag probe is required.^3^

We investigated if the fine-tuning of the spirocyclization equilibrium also provides control over the fluorogenicity of the corresponding HaloTag probes (**13** - **20**) (Figure 3a). First, the increase in absorbance and fluorescence upon binding to the HaloTag was tested *in-vitro* (F/F_0_). Due to the weak fluorogenic character of the original rhodamine *500R* HaloTag probe (**12**), labeling of the HaloTag only slightly elevated its absorbance and fluorescence signals (<2-fold). However, we identified various highly fluorogenic probes and the highest turn-on was observed for **19**, exhibiting an increase in absorbance of 54-fold and fluorescence of more than 130-fold (Figures 3a, b and Supplementary Figure 3). The fluorogenicity was further evaluated by live-cell, no-wash microscopy of U-2 OS cells which stably express a SNAP-Halo fusion protein localized to the nucleus and wild-type U-2 OS cells.^20^ We compared the nuclear signal of these U-2 OS FlpIn Halo-SNAP-NLS expressing cells (F_nuc_) with the cytosolic background signal of wild-type U-2 OS cells (F_cyt_) (Figures 3c and d). Consistent with the *in-vitro* test, **19** gave the highest nucleus-to-cytosol signal ratio (F_nuc_/F_cyt_ = 25) and **12** showed no significant fluorogenic behavior (F_nuc_/F_cyt_ = 2.7). Notably, we observed a decrease in nuclear signal intensity with increasing electron density on the benzenesulfonamide. However, since the decrease in background signal intensity was more pronounced, the F_nuc_/F_cyt_ ratio was highest for **19**. We observed that introduction of too nucleophilic amines significantly hampers the brightness of the fluorophore, as in the case of dimethylsulfamide **21**. These results show that the precise control over the spirocyclization equilibrium permits to optimize the fluorogenicity of a fluorophore for HaloTag labeling.

**Figure 3.**
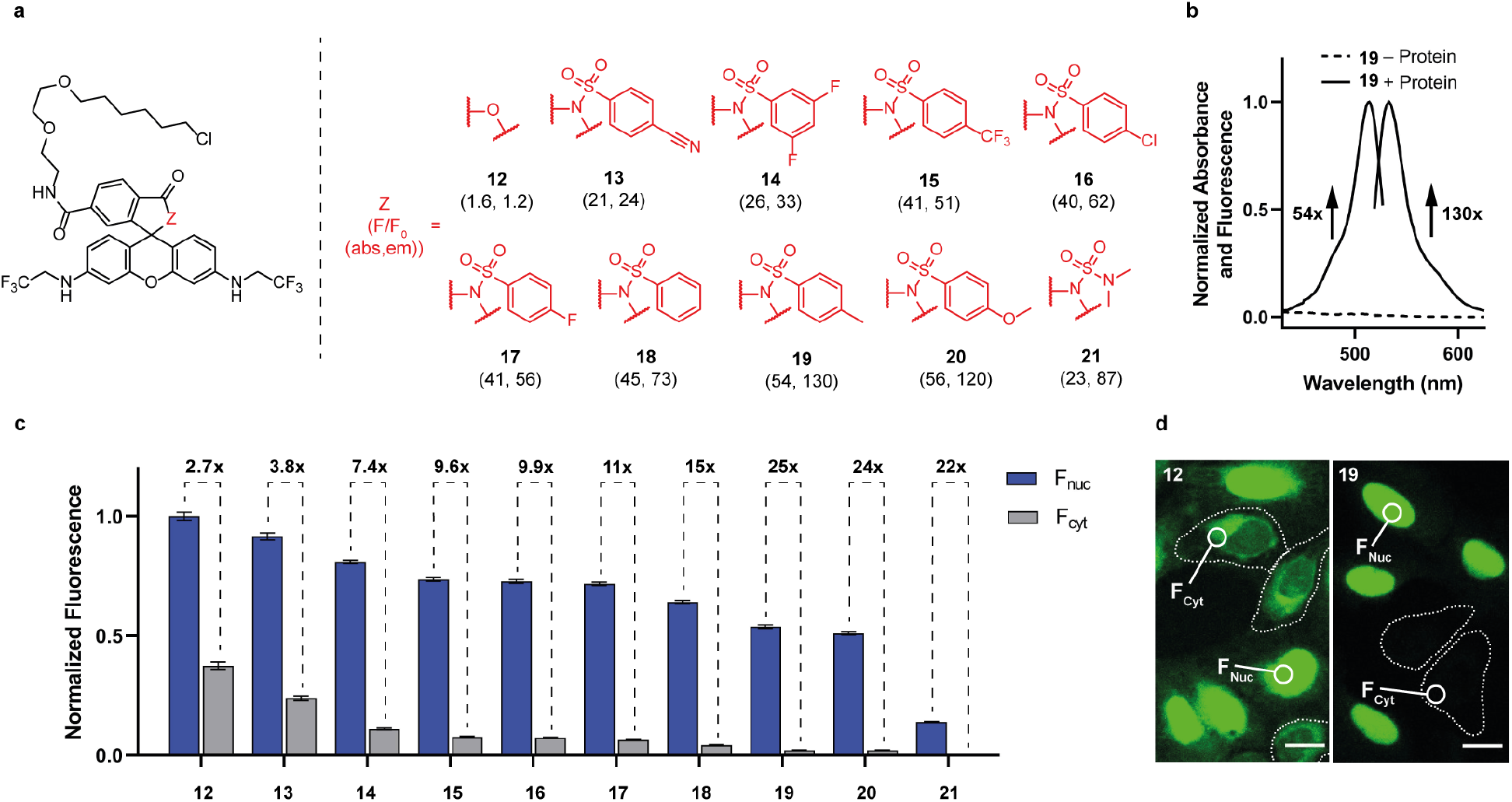
(a) Structures of rhodamine *500R* HaloTag probes (**12** - **21**) and absorbance, fluorescence emission turn-on upon HaloTag labeling (F/F_0_(abs, em)). (b) Normalized absorbance and fluorescence emission spectra of **19** (2.5 µM) in the absence (-Protein) and presence (+Protein) of HaloTag (5 µM) after 2.5 h incubation. (c) Fluorescence ratio (F_nuc_/F_cyt_) of **12** - **21** in live-cell, no-wash confocal microscopy. Bar plot representing the normalized nuclear signal (F_nuc_, U-2 OS FlpIn Halo-SNAP-NLS expressing cells, normalized to the nuclear signal of SiR-BG) and the cytosolic signal (F_cyt_, wild-type U-2 OS cells). Co-cultured U-2 OS FlpIn Halo-SNAP-NLS expressing cells and wild-type U-2 OS cells were prelabeled with SiR-BG (500 nM) overnight and then stained with **12** - **21** (200 nM) for 2.5 h. In total, 180 cells were examined from 3 independent experiments for each probe. Error bars show ± s.e.m. (d) Live-cell, no-wash confocal images of co-cultured U-2 OS FlpIn Halo-SNAP-NLS expressing cells and wild-type U-2 OS cells with **12** and **19**. Wild-type U-2 OS cells are represented with dotted lines. Scale bar, 20 µm.

### Optimizing Rhodamine *500R* for SNAP-tag Labeling

Another prominent example of a self-labeling protein tag is the SNAP-tag which specifically reacts with BG fluorophore derivatives.^47^ Recent studies revealed that SNAP-tag shifts the spirocyclization equilibrium of rhodamines much less to the fluorescent zwitterion than HaloTag, explaining the decreased brightness of certain fluorogenic rhodamines.^48-49^ These findings suggest that the optimal position of the spirocyclization equilibrium will differ for probes for SNAP-tag and HaloTag.

We synthesized and tested different rhodamine *500R* derived SNAP-tag probes (**22, 23** and **24**) for their applicability in live-cell, no-wash microscopy (Figure 4a). Measuring the intracellular labeling kinetics of SNAP-tag localized to the nucleus of U-2 OS cells indicated a low cell-permeability for the original rhodamine *500R* SNAP-tag probe (**22**) (Figure 4b). Furthermore, we observed high background signal intensities for **22** in *in-vitro* turn-on experiments and live-cell, no-wash microscopy (Figures 4c, d and Supplementary Figure 8). The *p*-toluenesulfonamide derivative (**24**) which yielded the highest fluorogenicity as a HaloTag probe (**19**) showed significantly decreased fluorescence signal upon binding to the SNAP-tag (Figure 4b and Supplementary Figure 8). We therefore envisioned to use a probe whose equilibrium is slightly shifted towards the zwitterion compared to **24**. Analyzing the performance of **23** which contains the more electron-deficient 4-fluorobenzenesulfonamide revealed an increased fluorescence signal upon binding to the SNAP-tag *in-vitro* as well as *in-cellulo* relative to **24** (Figure 4b and Supplementary Figure 8). Furthermore, enhanced cell-permeability and a lower cytosolic background signal compared to **22** were detected (Figures 4b, c and d). Our strategy therefore enables us to adjust fluorogenicity and cell-permeability of rhodamine *500R* to transform it into a suitable SNAP-tag probe for live-cell, no-wash microscopy.

**Figure 4.**
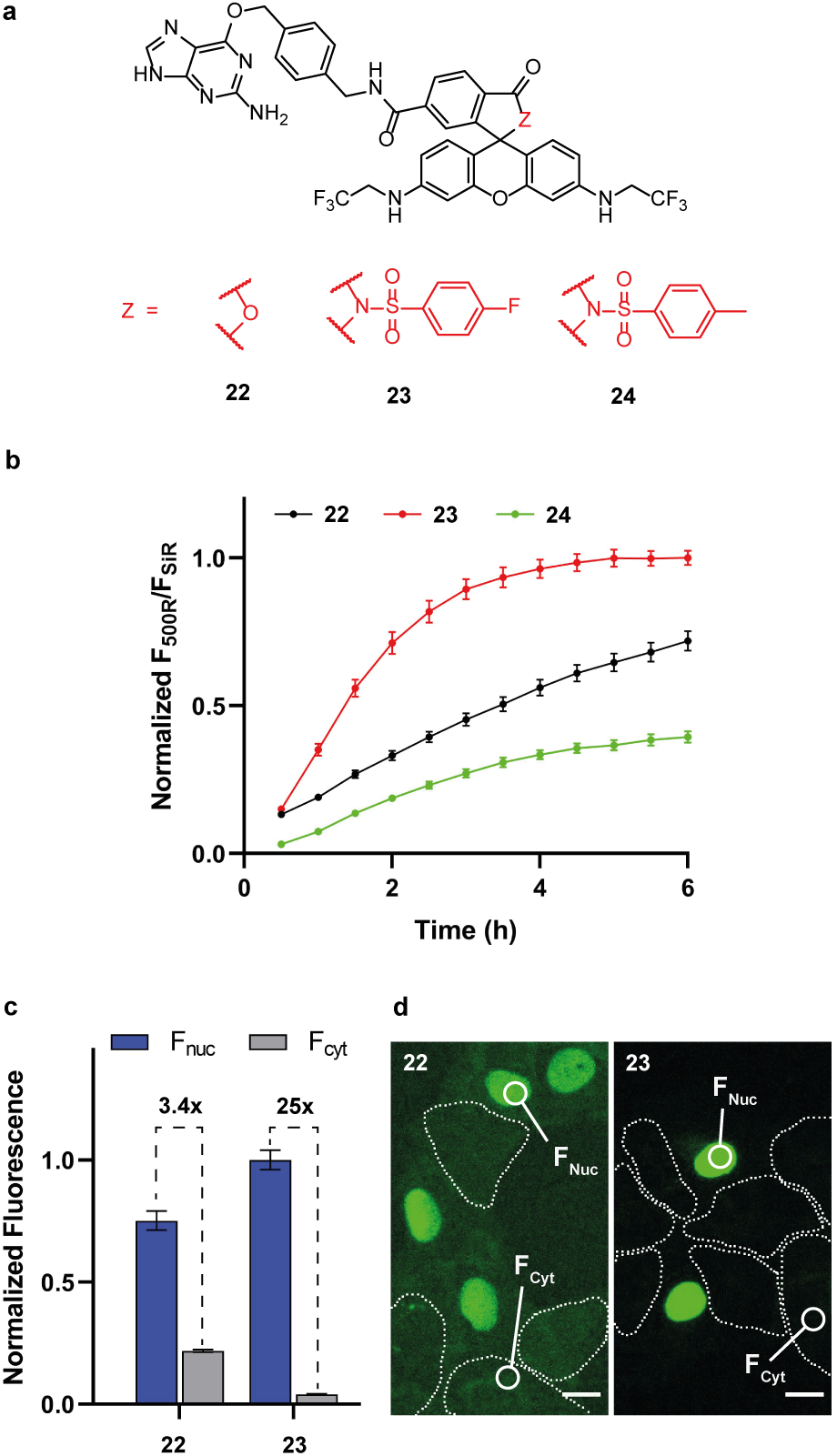
(a) Structures of rhodamine *500R* SNAP-tag probes (**22** - **24**). (b) Normalized ratio of rhodamine *500R* fluorescence to SiR fluorescence at various time points. U-2 OS FlpIn Halo-SNAP-NLS expressing cells were prelabeled with SiR-Halo (200 nM) overnight and treated with **22** - **24** (500 nM). In total, 90 cells were examined from 2 independent experiments for each probe. Error bars show ± s.e.m. (c) Fluorescence ratio (F_nuc_/F_cyt_) of **22** and **23** in live-cell, no-wash confocal microscopy. Bar plot representing the normalized nuclear signal (F_nuc_, U-2 OS FlpIn Halo-SNAP-NLS expressing cells, normalized to the nuclear signal of SiR-Halo) and the cytosolic signal (F_cyt_, wild-type U-2 OS cells). Co-cultured U-2 OS FlpIn Halo-SNAP-NLS expressing cells and wild-type U-2 OS cells were prelabeled with SiR-Halo (200 nM) overnight and then incubated with **22** and **23** (500 nM) for 5 h. In total, 180 cells were examined from 2 independent experiments for each probe. Error bars show ± s.e.m. (d) Live-cell, no-wash confocal images of co-cultured U-2 OS FlpIn Halo-SNAP-NLS expressing cells and wild-type U-2 OS cells with **22** and **23**. Wild-type U-2 OS cells are represented with dotted lines. Scale bar, 20 µm.

### A Spontaneously Blinking Rhodamine *500R* probe for SMLM

SMLM techniques require probes that switch between a fluorescent and a dark state.^3^ In addition to their application as photoswitching probes, recent studies also reported the spontaneous blinking behavior of rhodamine spirolactams.^40-41^ Inspired by these findings, we aimed to investigate if our simple synthetic strategy also enables the transformation of rhodamine *500R* into a spontaneously blinking probe for SMLM. We therefore synthesized HaloTag probes bearing more nucleophilic amides (**25, 26** and **27**) (Figure 5a). Next, we tested their blinking behavior and performance in SMLM by imaging endogenously tagged Nup96-Halo in fixed U-2 OS cells, a protein present in the nuclear pore complex.^50^ Labeling with the trifluoroethyl derivative of rhodamine *500R* (**26**) enabled visualization of the ring shape of the nuclear pore (Figures 5a, d, e and f). The average photon count of each localization was 631 and the localization precision peaked at 8.9 nm (Supplementary Figure 22). Furthermore, fixed U-2 OS cells which stably express HaloTag fused to the microtubule (MT) binding protein Cep41 were imaged with **26**.^51^ The resulting images showed fine structures of tubulin with a full-width-half-maximum (FWHM) of 40.7 nm ± 5.7 nm, which is in agreement with previous reports on MT diameter measurements (Figures 5b, c and Supplementary Figure 23).^51-52^ These observations corroborate the applicability of **26** in SMLM and its spontaneous blinking in standard phosphate-buffered saline (pH 7.4) without the need for low wavelength irradiation and additives (Supplementary Movies 1 and 2).

**Figure 5.**
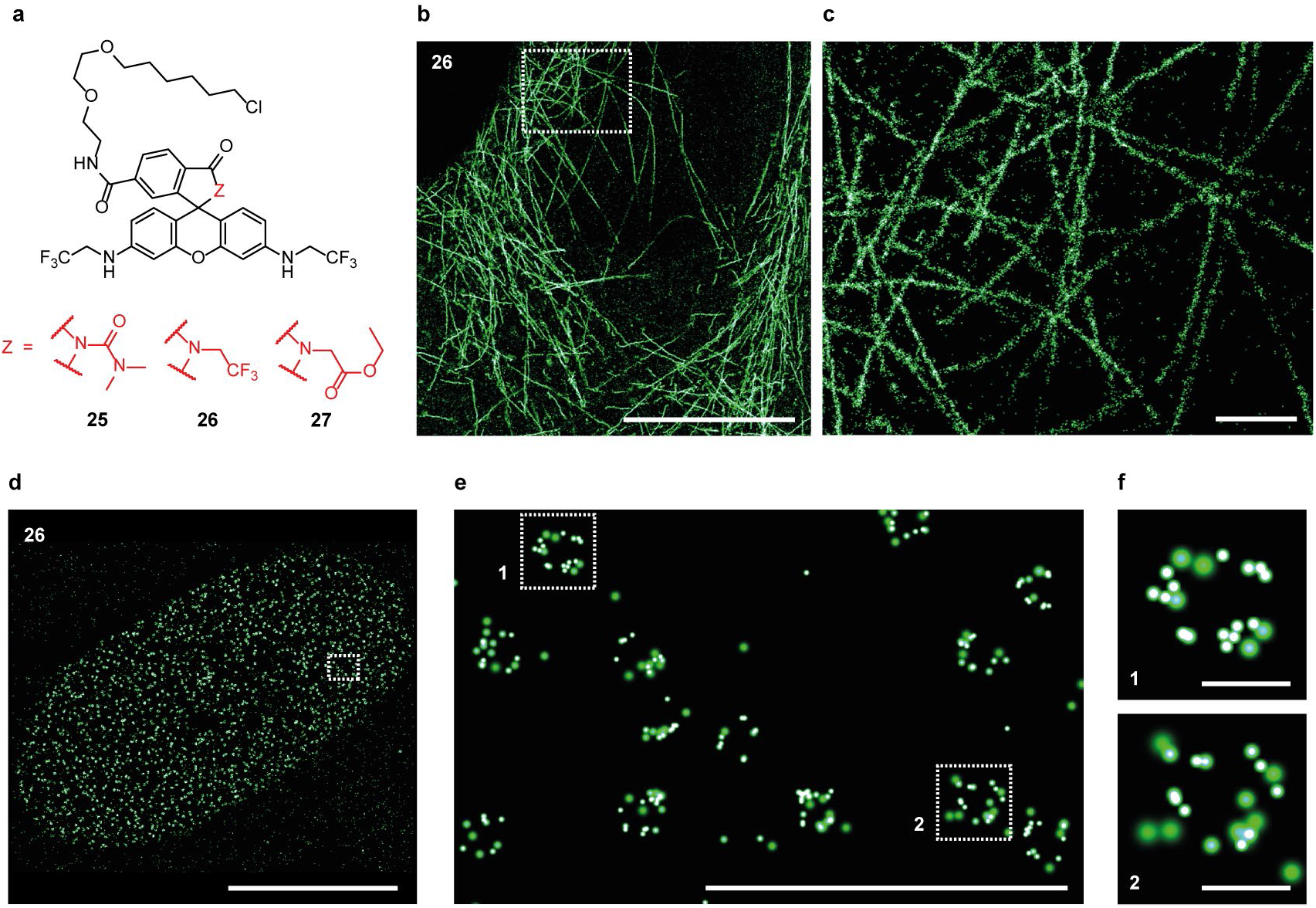
(a) Structures of highly closed rhodamine *500R* HaloTag probes (**25** - **27**). (b) Super-resolution image of fixed U-2 OS cells stably expressing Cep41-Halo labeled with **26** (1 µM) overnight. Scale bar, 10 µm. (c) Super-resolution image of the marked region in (b). Scale bar, 100 nm. (d) Super-resolution image of endogenously tagged Nup96-Halo in fixed U-2 OS cells labeled with **26** (1 µM) overnight. Scale bar, 10 µm. (e) Super-resolution image of the marked region in (d). Scale bar, 1 µm. (f) Individual nuclear pores of the marked regions in (e). Scale bar, 100 nm.

### Extending the Strategy to other Rhodamine-Based Dyes

Since our strategy enabled us to transform rhodamine *500R* into highly fluorogenic probes specifically optimized for SNAP-tag and HaloTag labeling as well as a blinking dye for SMLM, we investigated if our approach could be extended to other rhodamine scaffolds. The far-red silicon-rhodamine (SiR) scaffold has excellent photophysical properties and therefore represents an interesting target for modification.^24^ Due to the high propensity to exist in its spirocyclic state, SiR probes, such as SiR-Halo (**28**), are fluorogenic and cell-permeable (Figure 6a).^24^ Enhancing the fluorogenicity of **28** even further, requires precise fine-tuning of the spirocyclization equilibrium. Due to the higher D_50_ value of SiR (65) compared to rhodamine *500R* (34), we envisioned the introduction of an electron-deficient benzenesulfonamide derivative. We therefore synthesized the corresponding probe **29** (Figure 6a).^20^ **29** exhibited an increase in fluorescence intensity upon binding to HaloTag of more than 1100-fold *in-vitro*, which is almost a hundredfold higher than the turn-on we measured for **28** (Supplementary Figure 16). This simple modification furthermore provided a lower background signal in live-cell imaging, without considerably affecting the brightness of the probe bound to HaloTag (Figures 6b and c). Specifically, we observed an enhanced nucleus-to-cytosol signal ratio in live-cell, no-wash microscopy of U-2 OS FlpIn Halo-SNAP-NLS expressing cells and wild-type U-2 OS cells with **29** (F_nuc_/F_cyt_ = 150) compared to **28** (F_nuc_/F_cyt_ = 18) (Figures 6b and c). Next, we used our strategy to modify carbopyronine (CPY), an orange fluorophore scaffold with high photostability and brightness.^17^ In previous work, we found that replacing the *ortho*-carboxy group of CPY with an acyl dimethylsulfamide delivered a highly fluorogenic HaloTag probe (MaP618-Halo).^20^ However, the corresponding SNAP-tag probe (**31**) only showed negligible increase in fluorescence and absorbance upon binding to the SNAP-tag (Figures 6e and f).^20^ Thus, our goal was to tune the fluorogenicity of CPY and tailor it into a suitable SNAP-tag probe for live-cell, no-wash microscopy. CPY (40) shows a similar D_50_ value compared to rhodamine *500R* indicating a comparable state of the spirocyclization equilibrium. Since incorporation of 4-fluorobenzenesulfonamide transformed rhodamine *500R* into a highly fluorogenic SNAP-tag probe (**23**), we also applied this structural change to CPY to obtain **32** (Figure 6e). We then measured the absorbance and fluorescence increase of **32** upon binding to the SNAP-tag *in-vitro* and compared its fluorogenicity to the original CPY SNAP-tag probe (**30**). Despite a reduced brightness, **32** yielded a significantly increased turn-on in the presence of the SNAP-tag compared to **30** (Figure 6f). Live-cell no-wash microscopy of U-2 OS FlpIn Halo-SNAP-NLS expressing cells and wild-type U-2 OS cells corroborated the enhanced fluorogenicity for **32** (F_nuc_/F_cyt_ = 65) in comparison to **30** (F_nuc_/F_cyt_ = 19) (Supplementary Figure 12). STED microscopy is a powerful method to image cellular structures with high spatial resolution.^53^ The parental rhodamine *500R*, SiR and CPY dyes of **19, 29** and **32** show compatible spectroscopic properties for STED microscopy.^17, 54^ Since our strategy enhances the fluorogenicity without significantly altering these properties, we tested the applicability of **19, 29** and **32** in live-cell, no-wash STED imaging. U-2 OS cells stably expressing Vimentin-Halo were labeled with **19** and **29** as well as Vimentin-SNAP with **32**.^17, 55^ Images were recorded without prior washing-steps and showed bright structures with subdiffraction resolution and exceptionally low background fluorescence (Figures 6d, g and Supplementary Figure 7).

**Figure 6.**
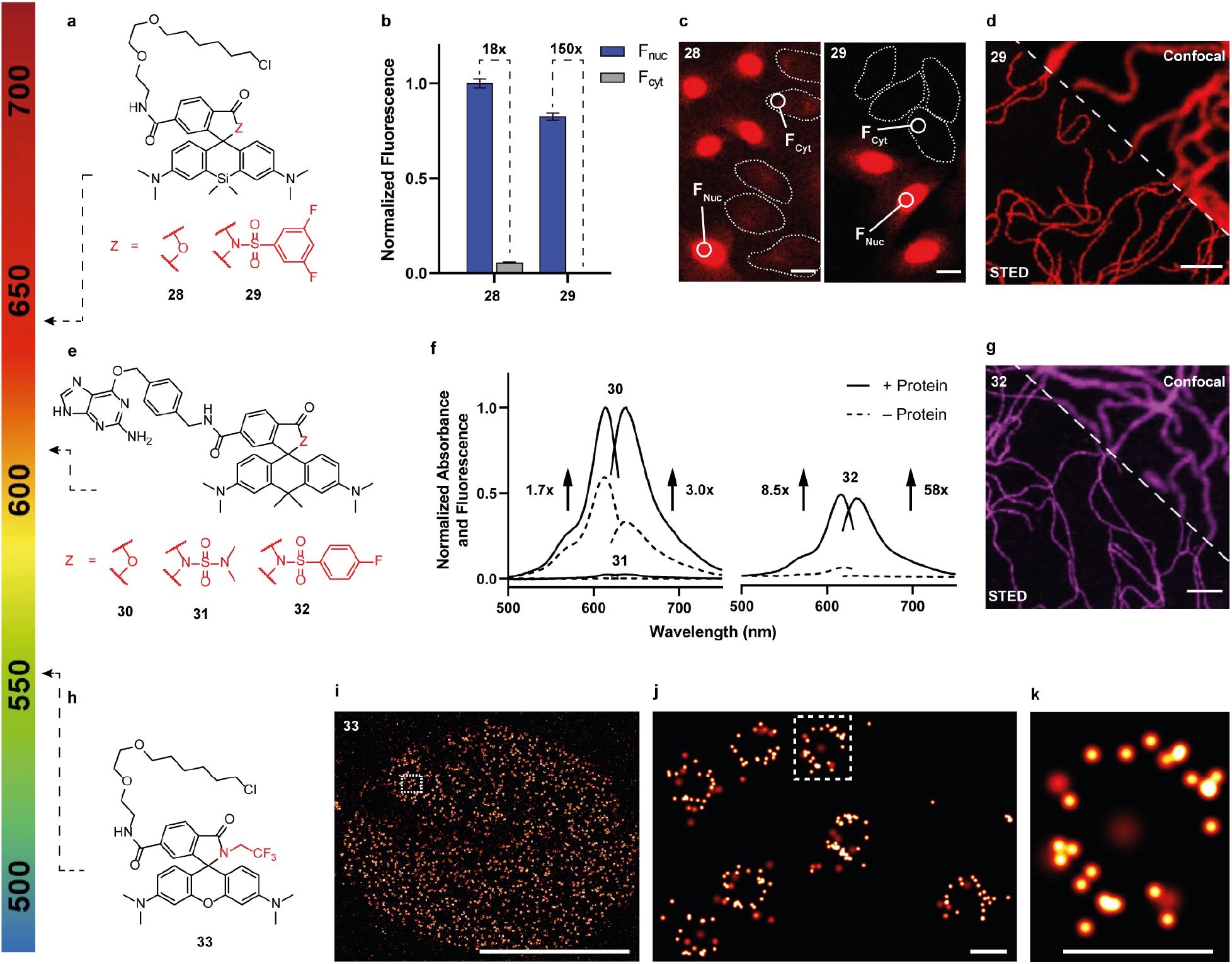
(a) Structures of SiR HaloTag probes (**28** and **29**). (b) Fluorescence ratio (F_nuc_/F_cyt_) of **28** and **29** in live-cell, no-wash confocal microscopy. Bar plot representing the normalized nuclear signal (F_nuc_, U-2 OS FlpIn Halo-SNAP-NLS expressing cells, normalized to the nuclear signal of **23**) and the cytosolic signal (F_cyt_, wild-type U-2 OS cells). Co-cultured U-2 OS FlpIn Halo-SNAP-NLS expressing cells and wild-type U-2 OS cells were prelabeled with **23** (500 nM) overnight and then incubated with **28** and **29** (500 nM) for 2.5 h. In total, 180 cells were examined from 2 independent experiments for each probe. Error bars show ± s.e.m. (c) Live-cell, no-wash confocal images of co-cultured U-2 OS FlpIn Halo-SNAP-NLS expressing cells and wild-type U-2 OS cells with **28** and **29**. Wild-type U-2 OS cells are represented with dotted lines. Scale bar, 20 µm. (d) Live-cell, no-wash confocal and STED images of U-2 OS Vimentin-Halo expressing cells labeled with **29** (500 nM) for 2 h. Image data was smoothed with a 1-pixel low pass Gaussian filter. Scale bar, 1.5 µm. (e) Structures of CPY SNAP-tag probes (**30 - 32**). (f) Normalized absorbance and fluorescence emission spectra of **30 - 32** (2.5 µM) in the absence (-Protein) and presence (+Protein) of SNAP-tag (5 µM) after 2.5 h incubation. (g) Live-cell, no-wash confocal and STED images of U-2 OS Vimentin-SNAP expressing cells labeled with **32** (500 nM) for 4 h. Image data was smoothed with a 1-pixel low pass Gaussian filter. Scale bar, 1.5 µm. (h) Structure of spontaneously blinking TMR HaloTag probe (**33**). (i) Super-resolution image of endogenously tagged Nup96-Halo in fixed U-2 OS cells labeled with **33** (1 µM) overnight. Scale bar, 10 µm. (j) Super-resolution image of the marked region in (i). Scale bar, 100 nm. (k) Individual nuclear pore of the marked region in (j). Scale bar, 100 nm.

Finally, we modified tetramethylrhodamine (TMR), a widely used rhodamine scaffold, by means of our strategy to expand the spectrum of colors for spontaneously blinking probes. We converted the *ortho*-carboxy group into a trifluoroethylamide moiety and introduced the CA ligand to obtain HaloTag probe **33** (Figure 6h). Analogous to **26**, we investigated the blinking behavior of **33** and its performance in SMLM by imaging endogenously tagged Nup96-Halo in fixed U-2 OS cells. **33** exhibited spontaneous blinking without the use of short-wavelength irradiation and any additive (Supplementary Movie 3). Furthermore, it yielded an average photon count of 2146 per localization and the localization precision peaked at 3.6 nm (Supplementary Figure 22). The resulting super-resolution images revealed the ring shape of the nuclear pores (Figures 6i, j and k). Next, we used **33** to image fixed U-2 OS cells which stably express Cep41-Halo. With this probe, we measured a FWHM of 28.5 nm ± 5.8 nm, which is compatible with the MT diameter of 25 nm (Supplementary Figure 23). The difference of measured MT diameters between **33** (28.5 nm ± 5.8 nm) and **26** (40.7 nm ± 5.7 nm) could be attributed to the difference in localization precision (**26** peaked at 9.9 nm, and **33** at 4.8 nm) (Supplementary Figures 22 and 23). However, both results lie within the range of previously reported values.^51-52^ Thus, our synthetic approach enabled us to develop **26** and **33**, two spectrally distinct and spontaneously blinking probes for SMLM.

## Conclusion

We have reported a general strategy that allowed us to adjust the equilibrium of spirocyclization of rhodamines with unprecedented precision and over a large range. We applied this approach to transform rhodamine *500R* into highly fluorogenic probes optimized for HaloTag and SNAP-tag labeling as well as a spontaneously blinking probe for SMLM. Furthermore, we extended our strategy to modify various commonly used rhodamines of different colors. The outstanding fluorogenicity and the blinking behavior of the resulting probes, respectively, make them powerful tools for live-cell, no-wash microscopy and SMLM. We envision that the generality of our strategy will allow the transformation of many other rhodamine-based fluorophores into probes that perfectly match the requirements of different imaging techniques and labeling systems.

## Supporting Information

Supplementary Figures and Tables, Supplementary Information - In-vitro Tests, Cell Culture and Microscopy, Supplementary Information – Synthesis and Characterization, NMR Spectra.

Supplementary Movies 1 - 4: Raw camera frames of **26** and **33**.

## Notes

The authors declare the following competing financial interest(s): K.J. and L.W. are inventors of the patent ‘Cell-permeable fluorogenic fluorophores’ which was filed by the Max Planck Society.

## Acknowledgment

The authors acknowledge funding of the Max Planck Society. Additional support came from the Deutsche Forschungsgemeinschaft (DFG, German Research Foundation, project number 240245660, SFB 1129, project Z3) to K.J. This research was conducted within the Max Planck School Matter to Life supported by the German Federal Ministry of Education and Research (BMBF) in collaboration with the Max Planck Society (K.J. and N.L.). This work was supported by the European Research Council (CoG-724489 to J.R) and the European Molecular Biology Laboratory (P.H., A.T. and J.R.). L.W. and M.T. are recipients of an Alexander von Humboldt Fellowship. We thank Andrea Bergner, Bettina Mathes, Jasmine Hubrich and Ulf Matti for skillful technical assistance, Michelle Frei for helpful discussions and the donation of the Cep41-Halo cells and Stefan Jakobs (Max Planck Institute for Biophysical Chemistry) for the donation of the Vimentin-Halo and Vimentin-SNAP cells.

