## Supporting Information for "Systematic Tuning of Rhodamine Spirocyclization for Super-Resolution Microscopy"

#### Table of Contents

| | | |
|---|---|---|
| 1 | Supplementary Figures and Tables | S-2 |
| 2 | Supplementary Information - *In-Vitro* Tests, Cell Culture and Microscopy | S-26 |
| 3 | Supplementary Information – Synthesis and Characterization | S-34 |
| 4 | References | S-55 |
| 5 | NMR Spectra | S-57 |

### Supplementary Figures and Tables

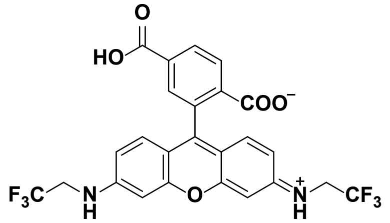

**1**

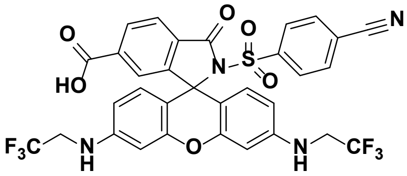

**4**

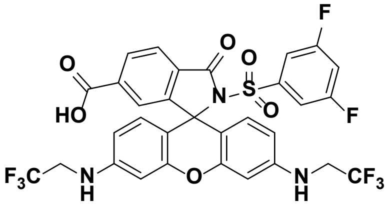

**5**

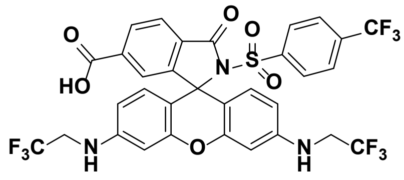

**6**

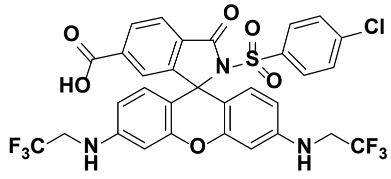

**7**

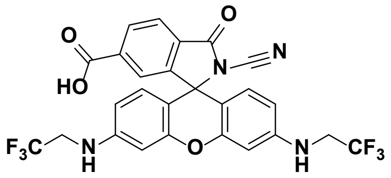

**34**

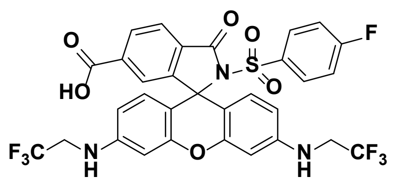

**8**

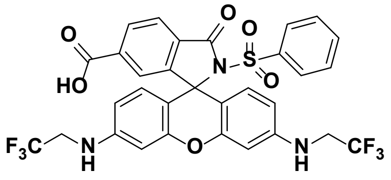

**9**

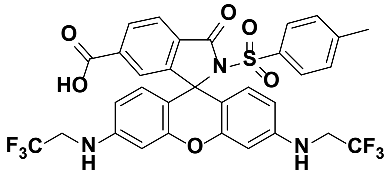

**10**

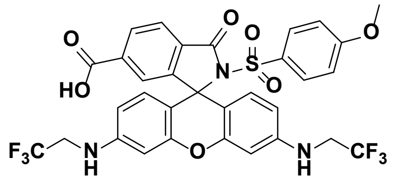

**11**

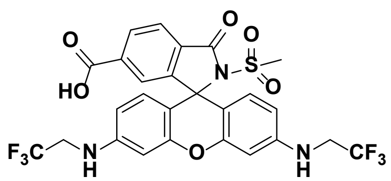

**35**

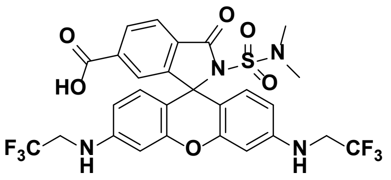

**36**

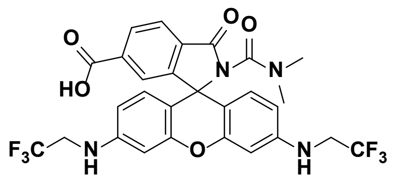

**37**

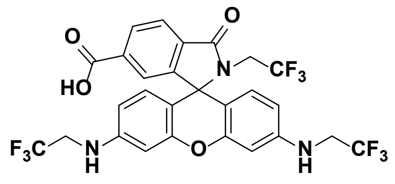

**38**

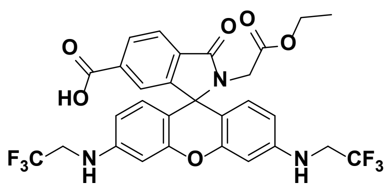

**39**

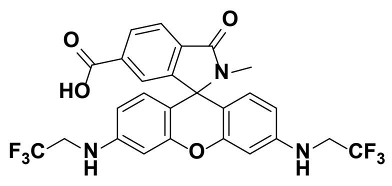

**40**

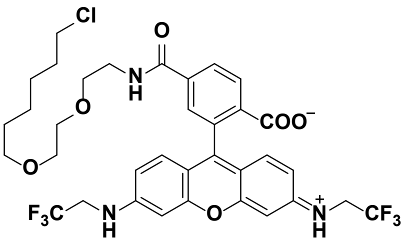

**12**

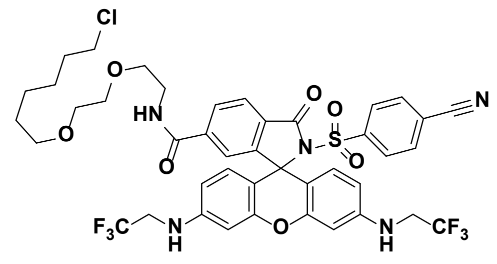

**13**

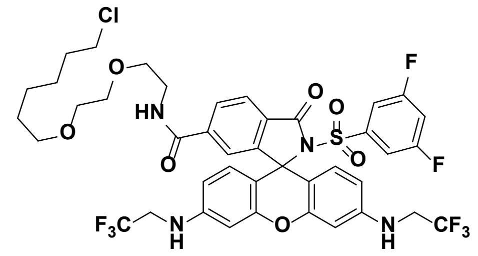

**14**

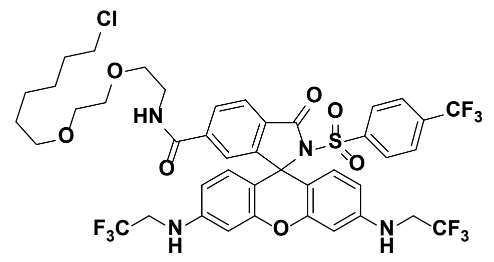

**15**

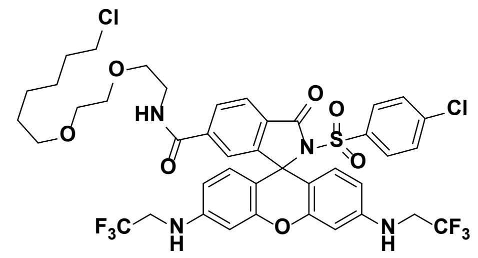

**16**

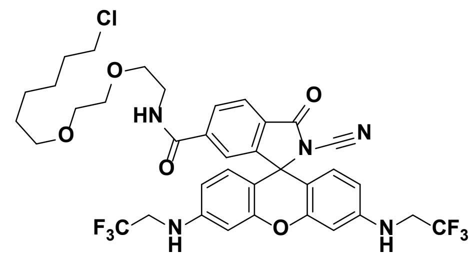

**41**

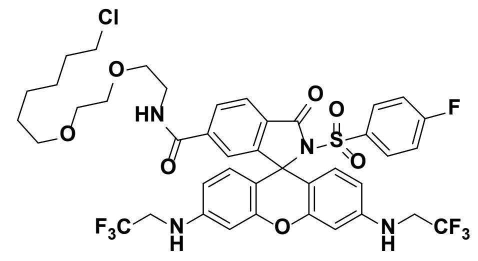

**17**

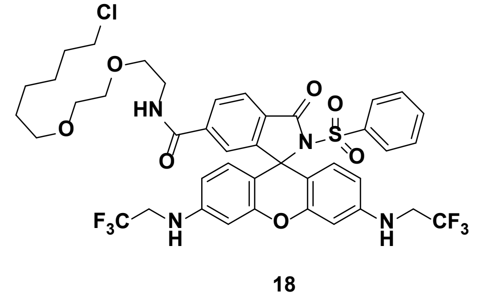

**18**

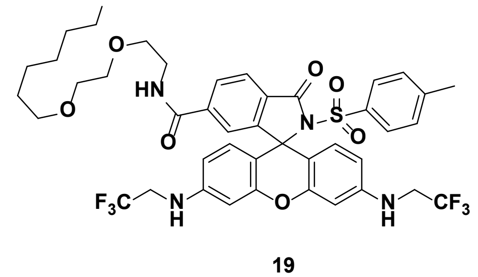

**19**

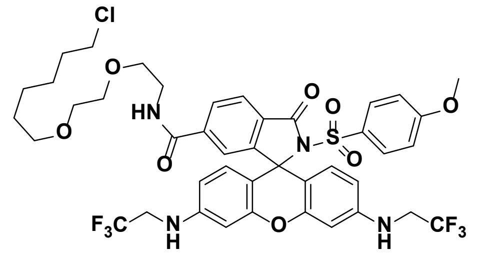

**20**

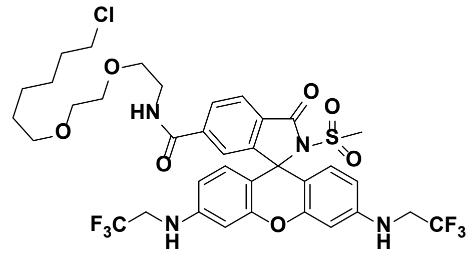

**42**

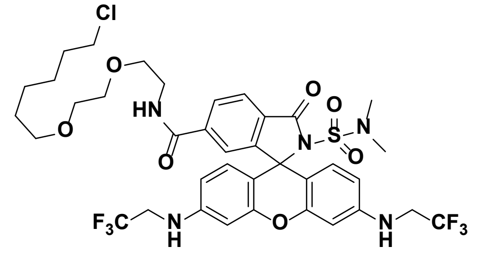

**21**

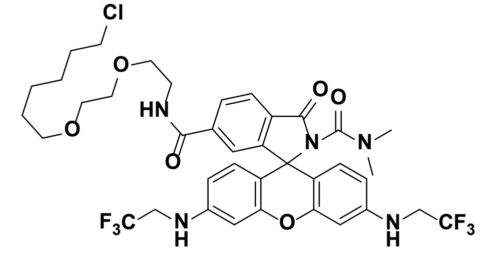

**25**

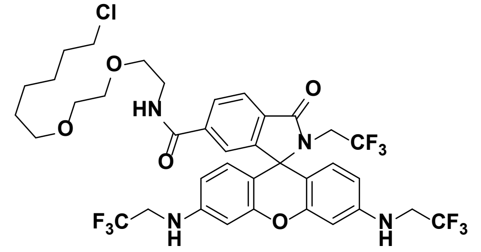

**26**

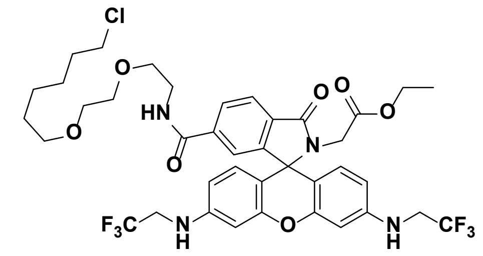

**27**

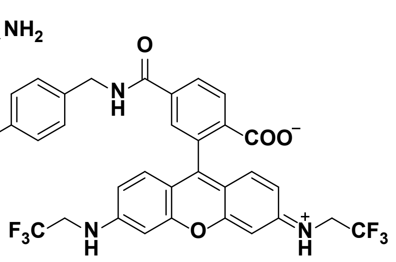

**22**

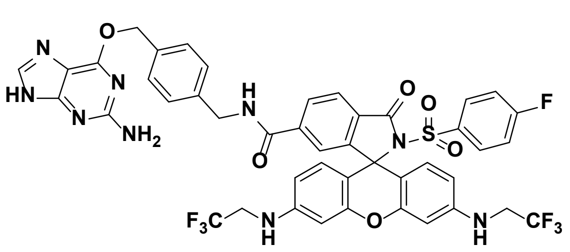

**23**

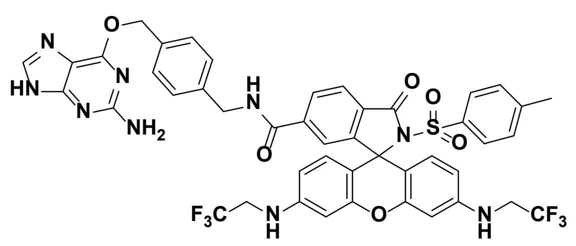

**24**

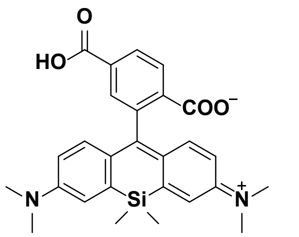

**43**

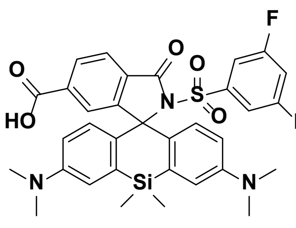

**44**

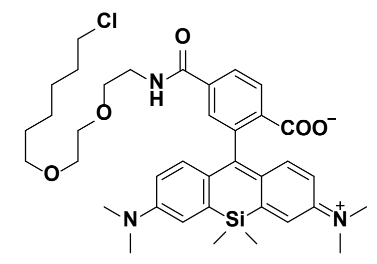

**28**

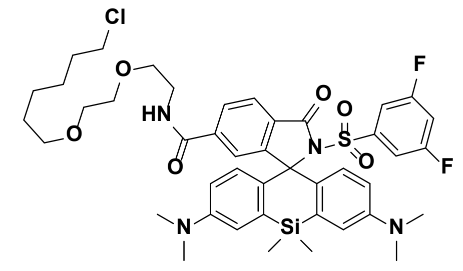

**29**

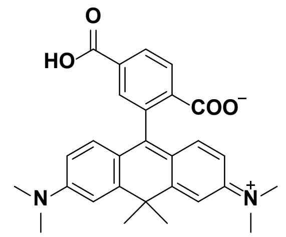

**45**

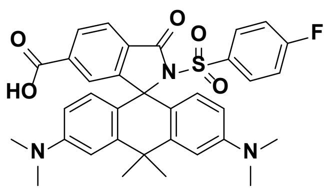

**46**

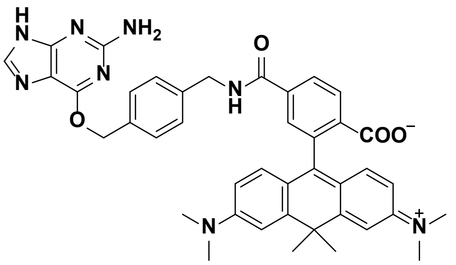

**30**

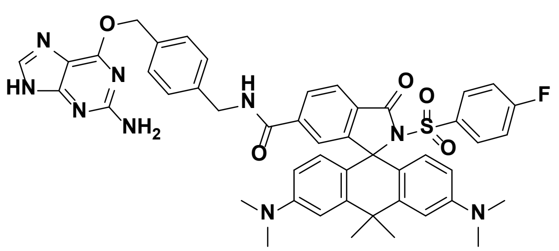

**32**

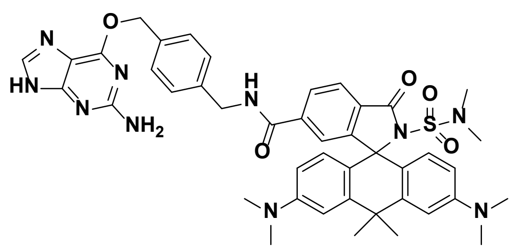

**31**

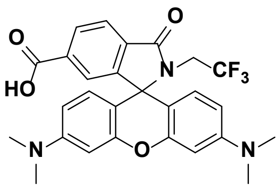

**47**

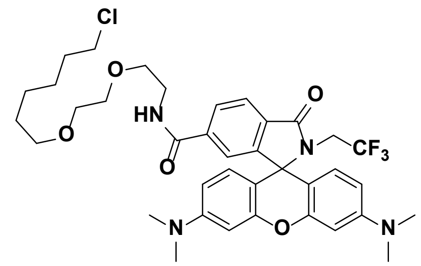

**33**

**Supplementary Figure 1:** Structures of fluorescent molecules.

**a**

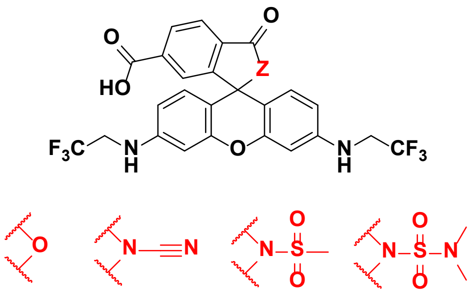

1 34 35 36

**b**

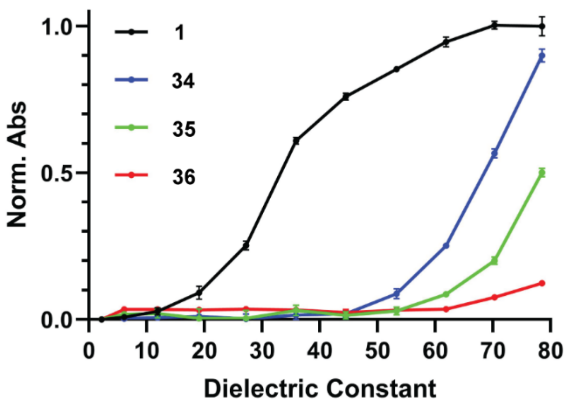


**Supplementary Figure 2:** Comparison of rhodamine *500R* derivatives (**1**, **34** - **36**). **a** Structures of rhodamine *500R* derivatives (**1**, **34** - **36**). **b** Normalized maximal absorbance of **1**, **34** - **36** (5 μM) in water–dioxane mixtures (v/v: 0/100 - 100/0) as a function of the dielectric constant.[1] The absorbance was normalized to the maximum absorbance of **1**. Error bars show ± s.d. from 3 experiments.

.

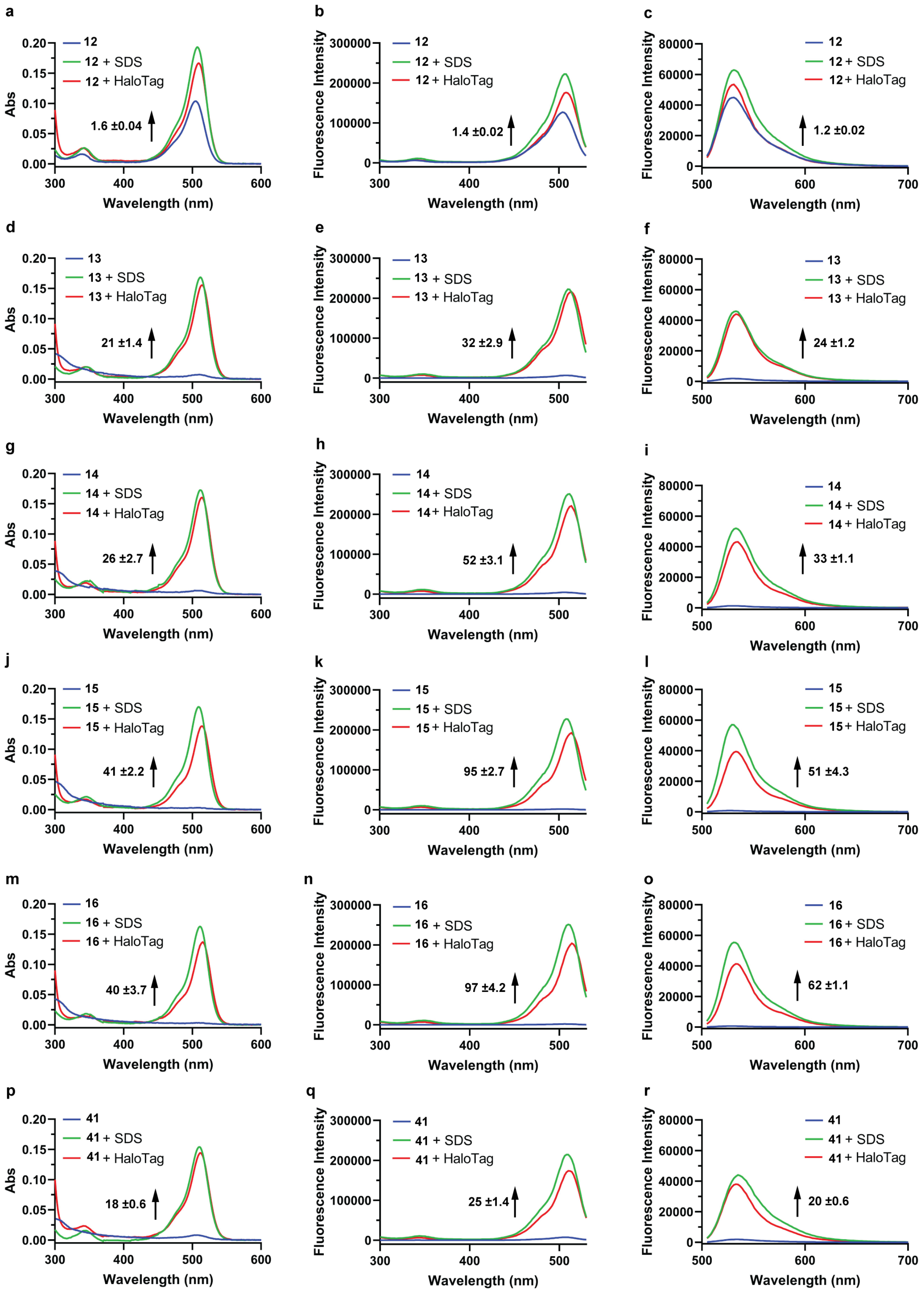
a
12
12 + SDS
12 + HaloTag
1.6 ±0.04
Abs
Wavelength (nm)
b
Fluorescence Intensity
1.4 ±0.02
c
1.2 ±0.02
d
13
13 + SDS
13 + HaloTag
21 ±1.4
e
32 ±2.9
f
24 ±1.2
g
14
14 + SDS
14 + HaloTag
26 ±2.7
h
52 ±3.1
i
33 ±1.1
j
15
15 + SDS
15 + HaloTag
41 ±2.2
k
95 ±2.7
l
51 ±4.3
m
16
16 + SDS
16 + HaloTag
40 ±3.7
n
97 ±4.2
o
62 ±1.1
p
41
41 + SDS
41 + HaloTag
18 ±0.6
q
25 ±1.4
r
20 ±0.6

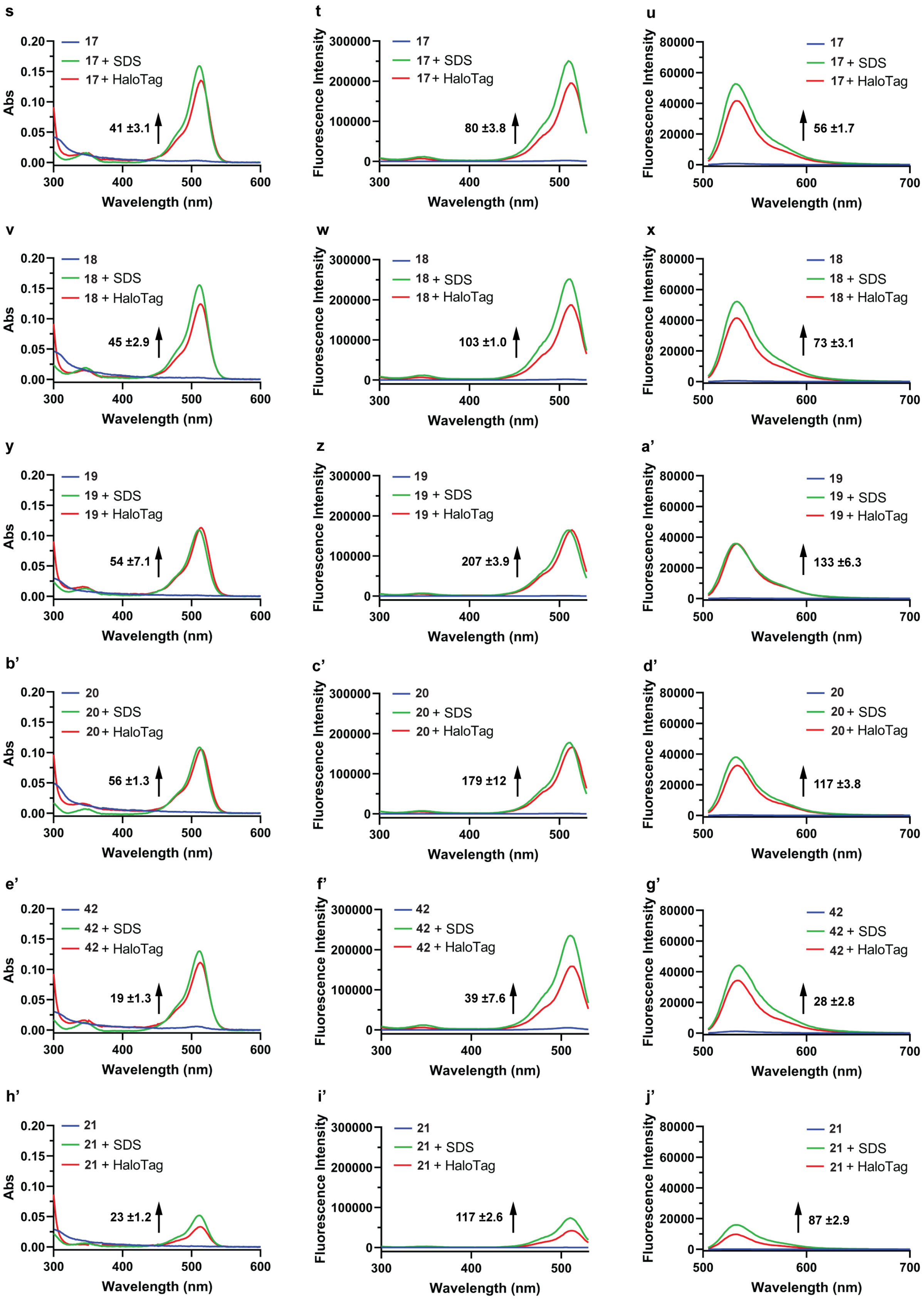
s
17
17 + SDS
17 + HaloTag
41 ±3.1
Abs
Wavelength (nm)
t
17
17 + SDS
17 + HaloTag
80 ±3.8
Fluorescence Intensity
Wavelength (nm)
u
17
17 + SDS
17 + HaloTag
56 ±1.7
Fluorescence Intensity
Wavelength (nm)
v
18
18 + SDS
18 + HaloTag
45 ±2.9
Abs
Wavelength (nm)
w
18
18 + SDS
18 + HaloTag
103 ±1.0
Fluorescence Intensity
Wavelength (nm)
x
18
18 + SDS
18 + HaloTag
73 ±3.1
Fluorescence Intensity
Wavelength (nm)
y
19
19 + SDS
19 + HaloTag
54 ±7.1
Abs
Wavelength (nm)
z
19
19 + SDS
19 + HaloTag
207 ±3.9
Fluorescence Intensity
Wavelength (nm)
a'
19
19 + SDS
19 + HaloTag
133 ±6.3
Fluorescence Intensity
Wavelength (nm)
b'
20
20 + SDS
20 + HaloTag
56 ±1.3
Abs
Wavelength (nm)
c'
20
20 + SDS
20 + HaloTag
179 ±12
Fluorescence Intensity
Wavelength (nm)
d'
20
20 + SDS
20 + HaloTag
117 ±3.8
Fluorescence Intensity
Wavelength (nm)
e'
42
42 + SDS
42 + HaloTag
19 ±1.3
Abs
Wavelength (nm)
f'
42
42 + SDS
42 + HaloTag
39 ±7.6
Fluorescence Intensity
Wavelength (nm)
g'
42
42 + SDS
42 + HaloTag
28 ±2.8
Fluorescence Intensity
Wavelength (nm)
h'
21
21 + SDS
21 + HaloTag
23 ±1.2
Abs
Wavelength (nm)
i'
21
21 + SDS
21 + HaloTag
117 ±2.6
Fluorescence Intensity
Wavelength (nm)
j'
21
21 + SDS
21 + HaloTag
87 ±2.9
Fluorescence Intensity
Wavelength (nm)

**Supplementary Figure 3:** Turn-on of rhodamine *500R* HaloTag probes (**12** – **21**, **41** and **42**) upon labeling of HaloTag and interaction with SDS. Absorption (**a, d, g, j, m, p, s, v, y, b', e', h'**), excitation (**b, e, h, k, n, q, t, w, z, c', f', i'**) and emission (**c, f, i, l, o, r, u, x, a', d', g', j'**) spectra of probes **12** – **21**, **41** and **42** (2.5 μM) measured in HEPES buffer (pH 7.3) in the absence (blue line) and presence of HaloTag (5 μM, red line) or SDS (0.1%, green line) after 2.5 h incubation. The numbers represent the ratio of maximal absorbance and fluorescence intensities (emission: $\lambda_{ex}$: 490 nm, excitation $\lambda_{em}$: 550 nm) in the presence and absence of the HaloTag ($F/F_0$). Representative data for 3 experiments. Errors show ± s.d.

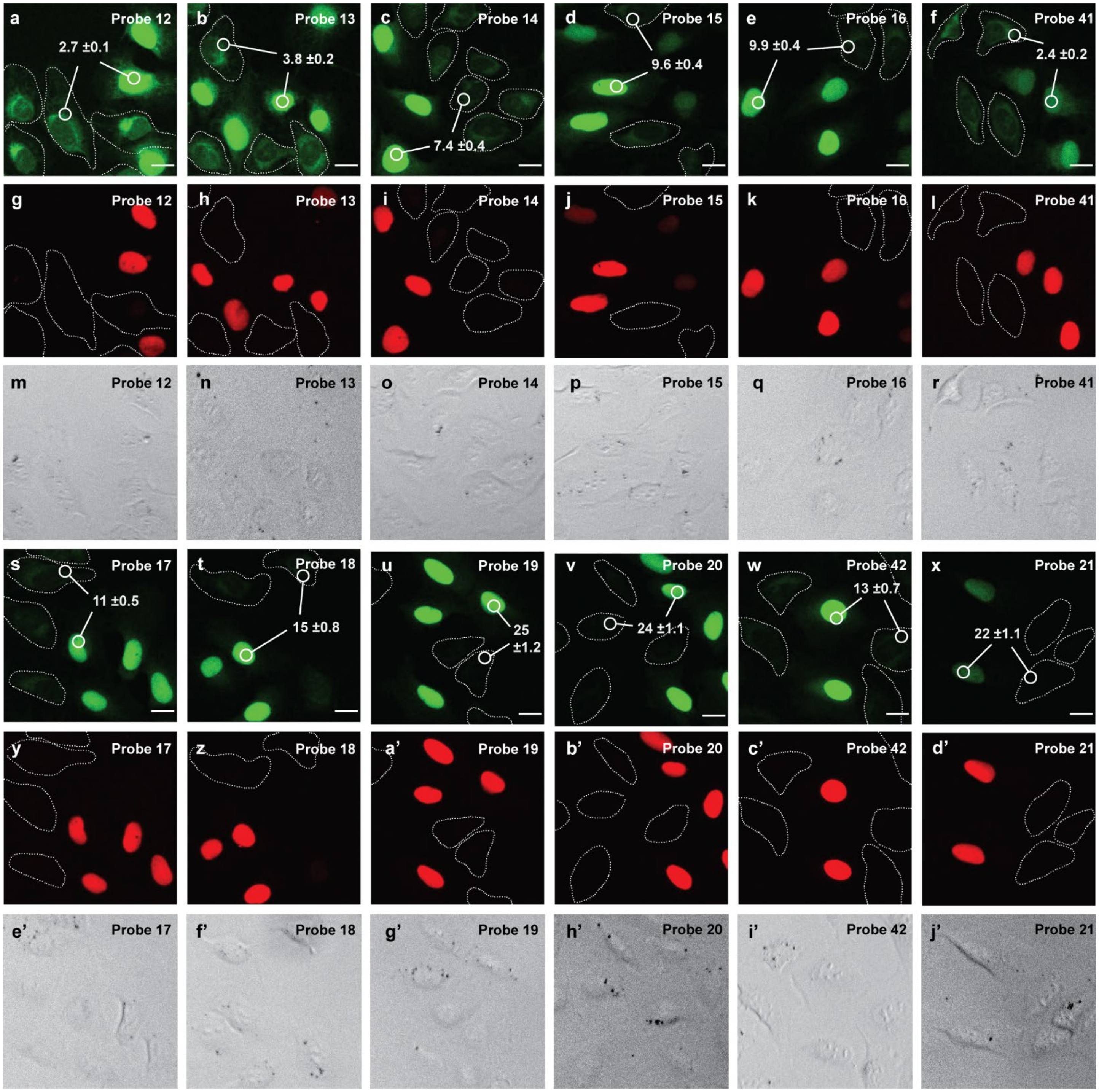


**Supplementary Figure 4:** Evaluation of fluorogenicity of rhodamine *500R* HaloTag probes (**12** – **21**, **41** and **42**) in live-cell, no-wash confocal microscopy. Co-cultured U-2 OS FlpIn Halo-SNAP-NLS expressing cells and wild-type U-2 OS cells were labeled with 500 nM SiR-BG overnight, washed once with imaging medium and then labeled with 200 nM of **12** – **21**, **41** and **42** for 2.5 h. **a – f, s – x** Live-cell, no-wash confocal images (sum projections of z-stacks) showing rhodamine *500R* fluorescence and **g – l, y – d'** SiR fluorescence. Wild-type U-2 OS cells are represented with dotted lines. **m – r, e' – j'** Bright field images. The numbers correspond to the fluorescence ratios $F_{nuc}/F_{cyt}$. $F_{nuc}$: Mean values of rhodamine *500R* fluorescence of regions of interest (ROIs) within the nuclei of U-2 OS FlpIn Halo-SNAP-NLS expressing cells normalized to the SiR fluorescence. $F_{cyt}$: Mean values of rhodamine *500R* fluorescence of ROIs within the cytosol of wild-type U-2 OS cells. In total, 180 cells were examined from 3 independent experiments for each probe. Errors represent ± s.e.m. Scale bar, 20 µm.

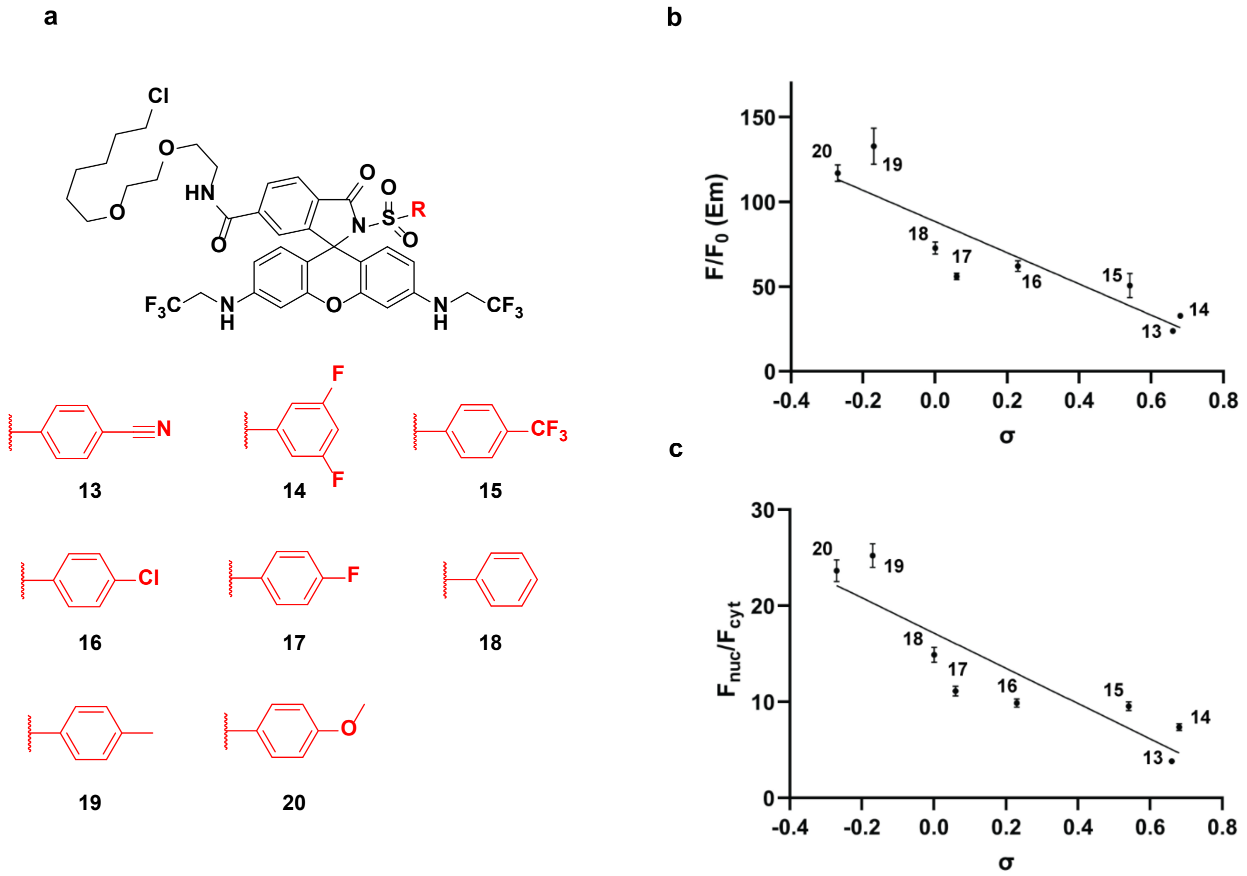


**Supplementary Figure 5:** Effect of structural changes on the aromatic sulfonamide onto the *in-vitro* and *in-cellulo* fluorogenicity. **a** Structures of rhodamine *500R* derivatives (**13** - **20**). **b** Correlation of the ratio of maximal fluorescence intensities in the presence and absence of the HaloTag ($F/F_0$(Em)) versus Hammett constants (σ) of the substituents at the aromatic ring.[2] Error bars show ± s.d. from 3 experiments. Solid line shows linear regression ($R^2$ = 0.80). **c** Correlation of the fluorescence ratios $F_{nuc}/F_{cyt}$ versus σ of the substituents at the aromatic ring. The error bars show ± s.e.m. In total, 180 cells were examined from 3 independent experiments for each probe. Solid line shows linear regression ($R^2$ = 0.80). For disubstituted compound **14**, the σ of the substituent was doubled.

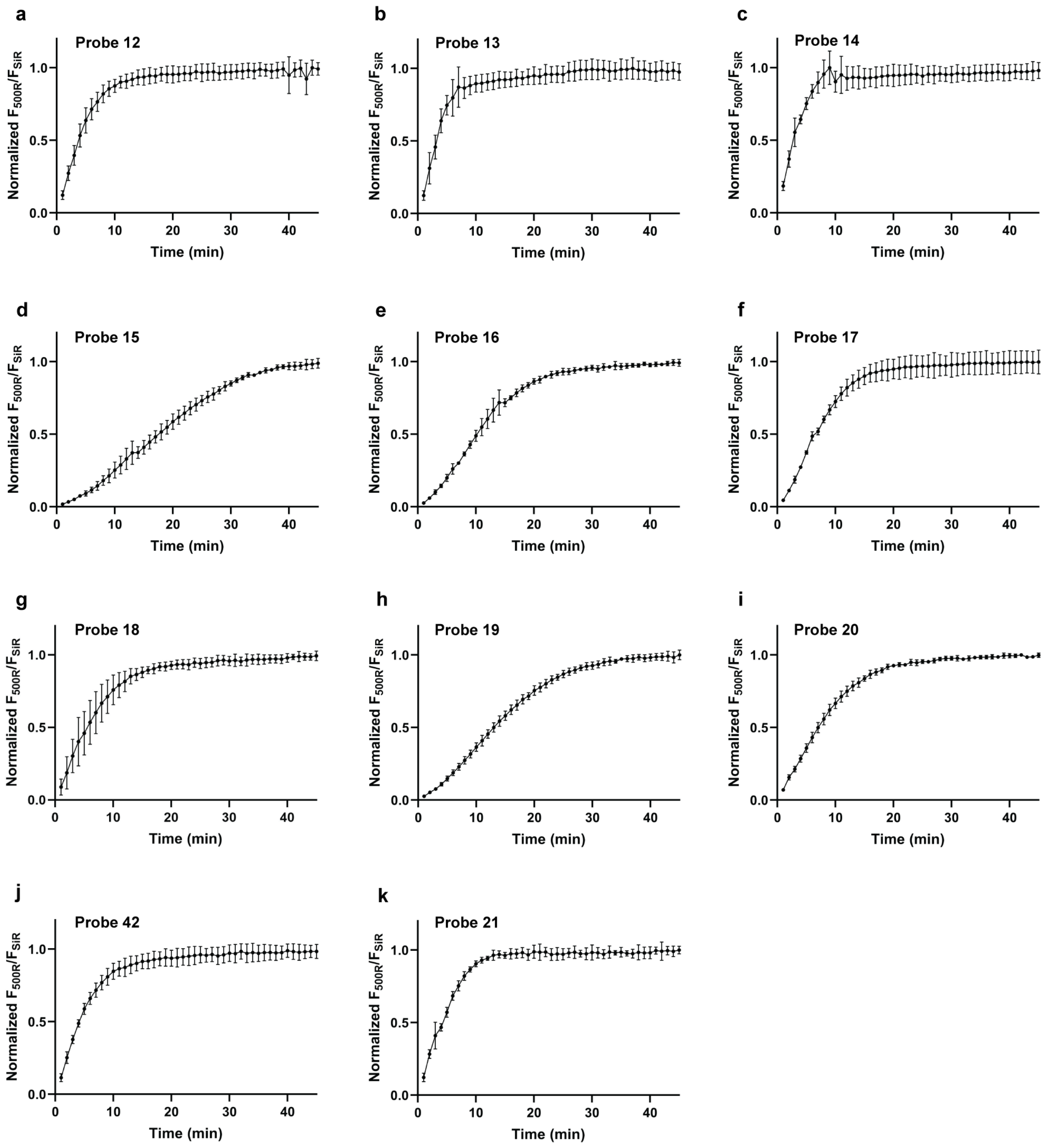


**Supplementary Figure 6:** Intracellular HaloTag labeling with 200 nM of rhodamine *500R* HaloTag probes (**12** – **21** and **42**) as a function of time. U-2 OS FlpIn Halo-SNAP-NLS expressing cells were pre-labeled with 500 nM SiR-BG overnight. Normalized ratio of rhodamine *500R* fluorescence to the fluorescence signal of SiR-BG is plotted at different time points. In total, >51 cells were examined from technical triplicates for each probe. Error bars show ± s.e.m.

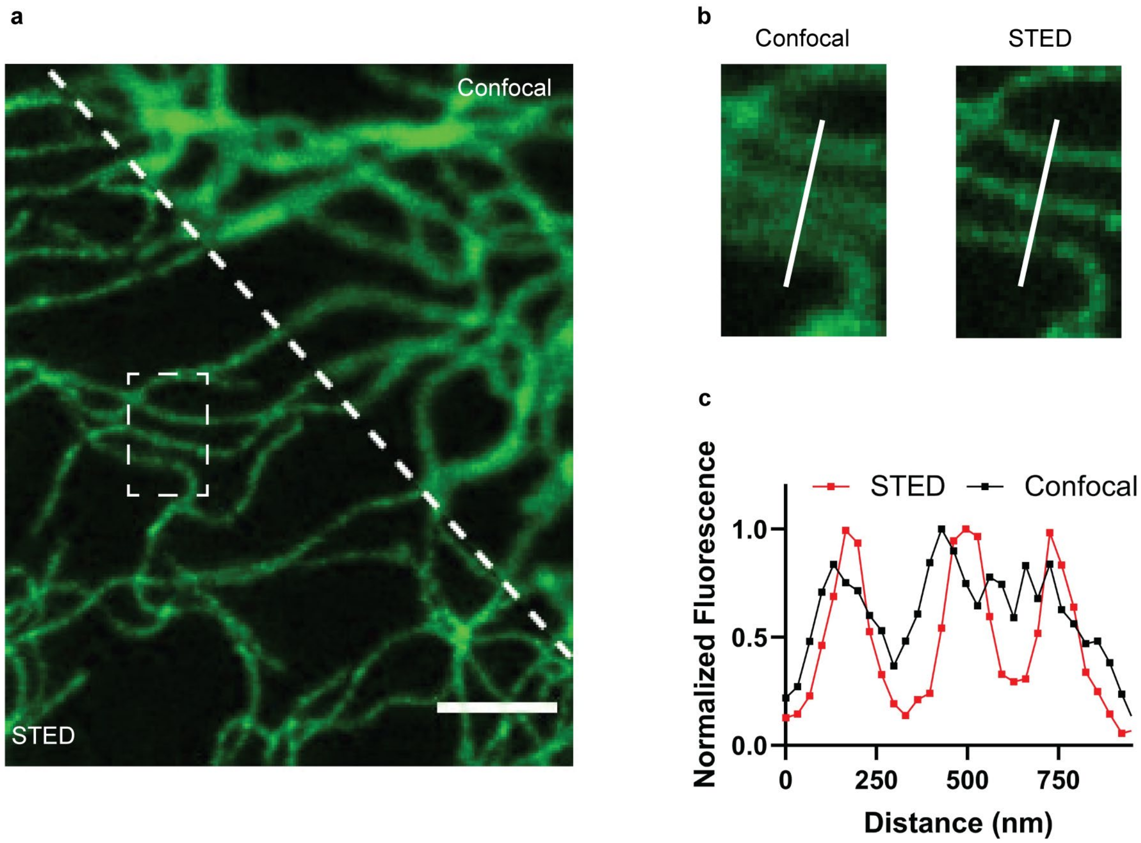


**Supplementary Figure 7:** Comparison of live-cell, no-wash confocal and stimulated emission depletion (STED) microscopy for probe **19**. **a** Confocal and STED image of U-2 OS Vimentin-Halo expressing cells labeled with 200 nM of **19** for 2 h. Scale bar, 1.5 µm. **b** Confocal and STED image from the marked region in **a**. **c** Corresponding line profiles. Image data was smoothed with a 1-pixel low pass Gaussian filter. In total, 32 cells were examined from 2 independent experiments.

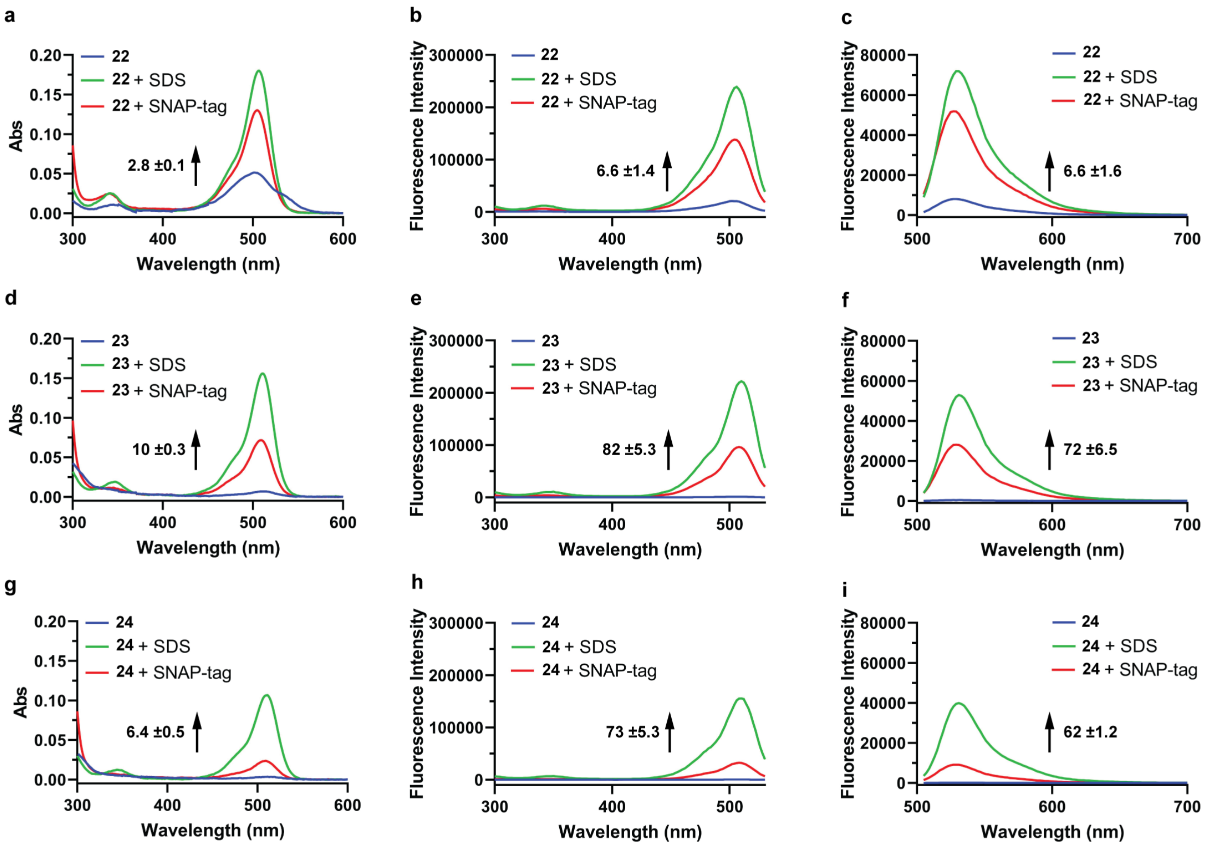


**Supplementary Figure 8:** Turn-on of rhodamine *500R* SNAP-tag probes (**22** - **24**) upon labeling of SNAP-tag and interaction with SDS. Absorption (**a, d, g**), excitation (**b, e, h**) and emission (**c, f, i**) spectra of probes **22** - **24** (2.5 µM) measured in HEPES buffer (pH 7.3) in the absence (blue line) and presence of SNAP-tag (5 µM, red line) or SDS (0.1%, green line) after 2.5 h incubation. The numbers represent the ratio of maximal absorbance and fluorescence intensities (emission: $\lambda_{ex}$: 490 nm, excitation $\lambda_{em}$: 550 nm) in the presence and absence of the SNAP-tag ($F/F_0$). Representative data for 3 experiments. Errors show ± s.d.

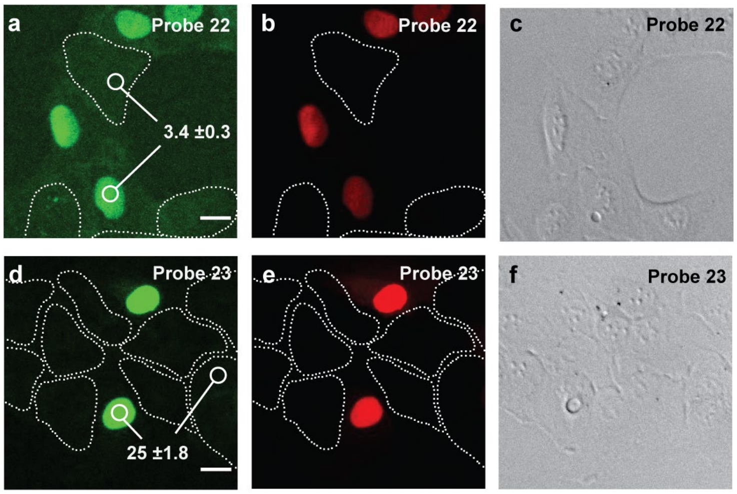


**Supplementary Figure 9:** Evaluation of fluorogenicity of rhodamine *500R* SNAP-tag probes (**22** and **23**) in live-cell, no-wash confocal microscopy. Co-cultured U-2 OS FlpIn Halo-SNAP-NLS expressing cells and wild-type U-2 OS cells were labeled with 200 nM SiR-Halo overnight, washed once with imaging medium and then labeled with 500 nM of **22** and **23** for 5 h. **a, d** Live-cell, no-wash confocal images (sum projections of z-stacks) showing rhodamine *500R* fluorescence and **b, e** SiR fluorescence. Wild-type U-2 OS cells are represented with dotted lines. **c, f** Bright field images. The numbers correspond to the fluorescence ratios $F_{nuc}/F_{cyt}$. $F_{nuc}$: Mean values of rhodamine *500R* fluorescence of ROIs within the nuclei of U-2 OS FlpIn Halo-SNAP-NLS expressing cells normalized to the SiR fluorescence. $F_{cyt}$: Mean values of rhodamine *500R* fluorescence of ROIs within the cytosol of wild-type U-2 OS cells. In total, 180 cells were examined from 2 independent experiments for each probe. Errors represent ± s.e.m. Scale bar, 20 μm.

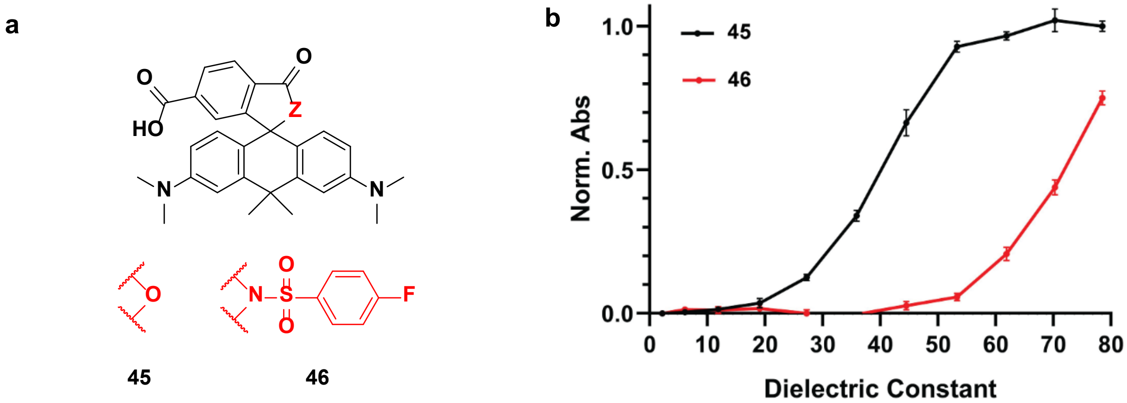


**Supplementary Figure 10:** Comparison of CPY derivatives (**45** and **46**). **a** Structures of CPY derivatives (**45** and **46**). **b** Normalized maximal absorbance of **45** and **46** (5 μM) in water–dioxane mixtures (v/v: 0/100

- 100/0) as a function of the dielectric constant.[1] The absorbance was normalized to the maximum absorbance of **45**. Error bars show ± s.d. from 3 experiments.

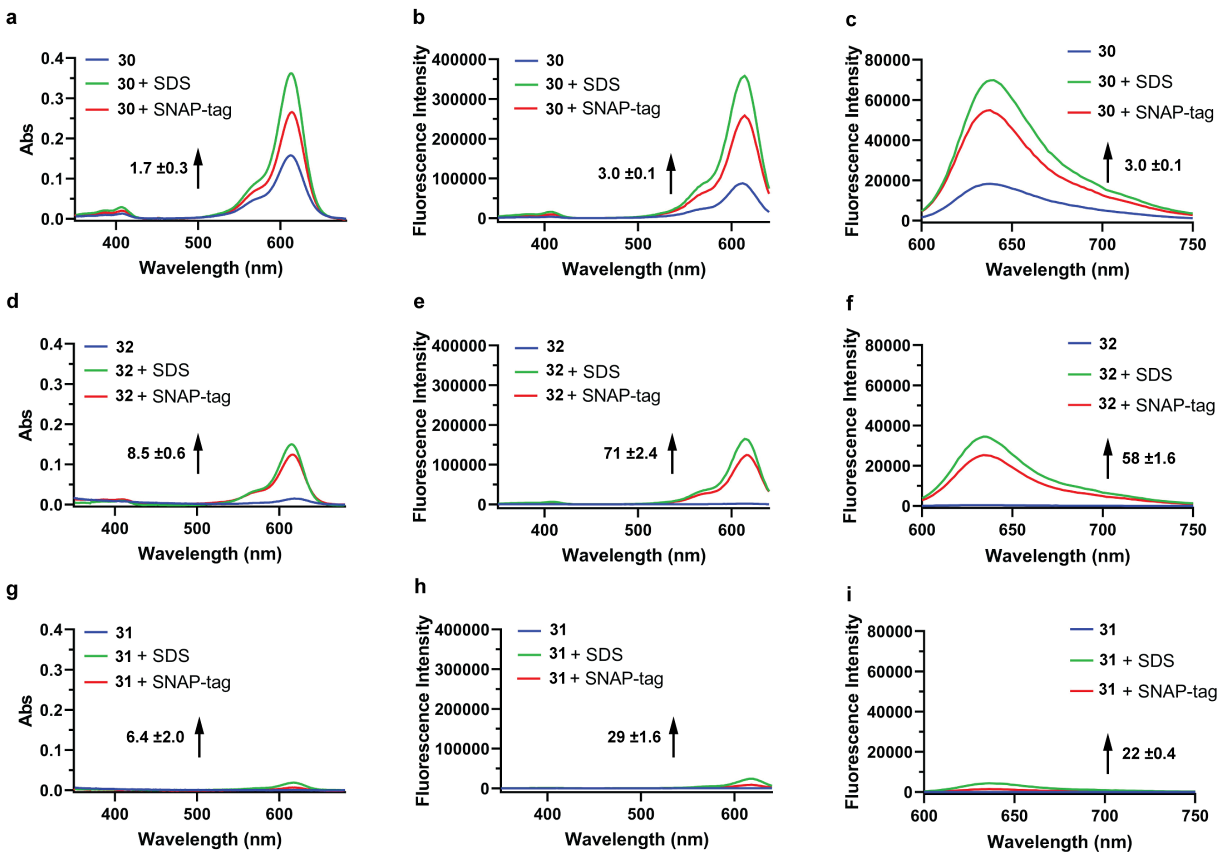


**Supplementary Figure 11:** Turn-on of CPY SNAP-tag probes (**30** - **32**) upon labeling of SNAP-tag and interaction with SDS. Absorption (**a, d, g**), excitation (**b, e, h**) and emission (**c, f, i**) spectra of probes **30** - **32** (2.5 µM) measured in HEPES buffer (pH 7.3) in the absence (blue line) and presence of SNAP-tag (5 µM, red line) or SDS (0.1%, green line) after 2.5 h incubation. The numbers represent the ratio of maximal absorbance and fluorescence intensities (emission: $\lambda_{ex}$: 580 nm, excitation $\lambda_{em}$: 660 nm) in the presence and absence of the SNAP-tag ($F/F_0$). Representative data for 3 experiments. Errors show ± s.d.

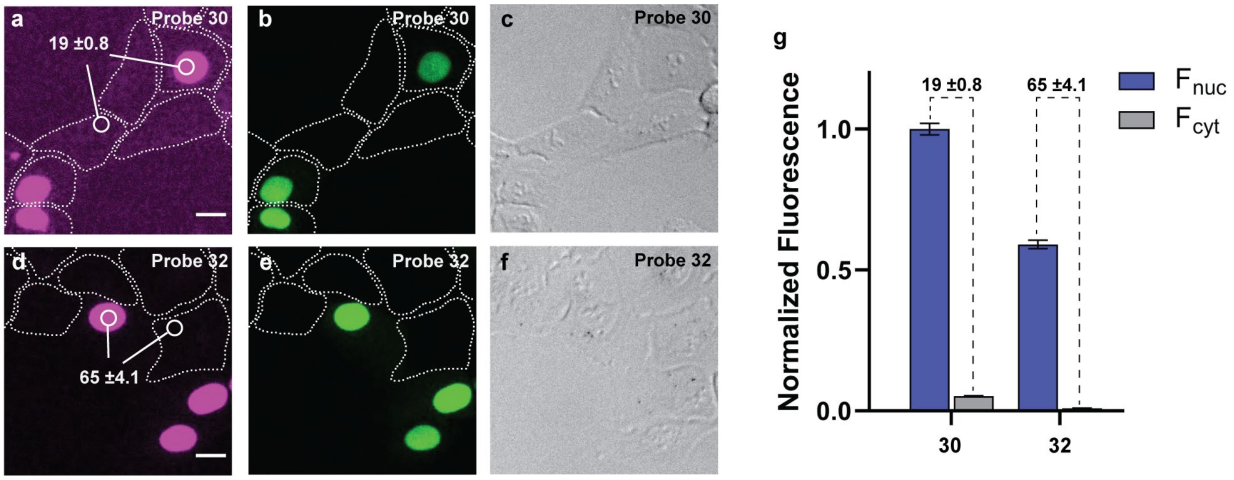


**Supplementary Figure 12:** Evaluation of fluorogenicity of CPY SNAP-tag probes (**30** and **32**) in live-cell, no-wash confocal microscopy. Co-cultured U-2 OS FlpIn Halo-SNAP-NLS expressing cells and wild-type U-2 OS cells were labeled with 200 nM **19** overnight, washed once with imaging medium and then labeled with 500 nM of **30** and **32** for 4 h. **a, d** Live-cell, no-wash confocal images (sum projections of z-stacks) showing CPY fluorescence and **b, e** fluorescence of **19**. Wild-type U-2 OS cells are represented with dotted lines. **c, f** Bright field images. The numbers as well as the bar plot (**g**) represent the fluorescence ratios $F_{nuc}/F_{cyt}$. $F_{nuc}$: Mean values of CPY fluorescence of ROIs within the nuclei of U-2 OS FlpIn Halo-SNAP-NLS expressing cells normalized to the fluorescence of **19**. $F_{cyt}$: Mean values of CPY fluorescence of ROIs within the cytosol of wild-type U-2 OS cells. In total, 180 cells were examined from 2 independent experiments for each probe. Errors represent ± s.e.m. Scale bar, 20 µm.

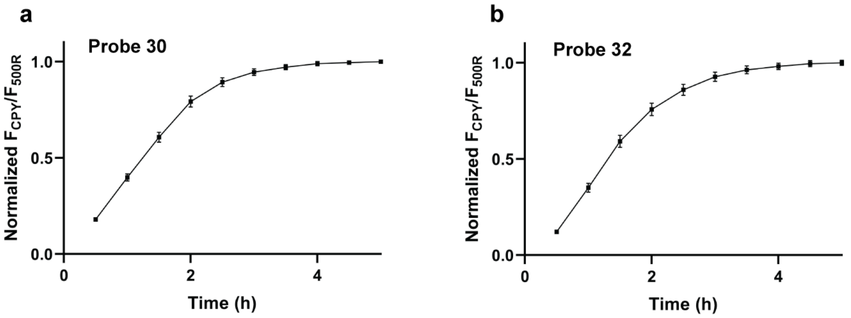


**Supplementary Figure 13:** Intracellular SNAP-tag labeling with 500 nM of CPY SNAP-tag probes (**30** and **32**) as a function of time. U2-O S FlpIn Halo-SNAP-NLS expressing cells were pre-labeled with 200 nM **19** overnight. Normalized ratio of CPY fluorescence to the fluorescence signal of **19** is plotted at different time points. In total, 90 cells were examined from technical triplicates for each probe. Error bars show ± s.e.m.

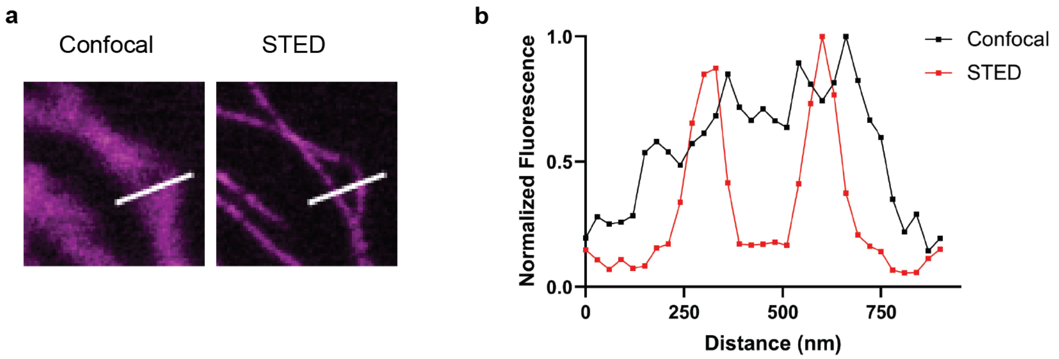


**Supplementary Figure 14:** Comparison of live-cell, no-wash confocal and STED microscopy for probe **32**. **a** Confocal and STED image (from **Figure 6g**) of U-2 OS Vimentin-SNAP expressing cells labeled with 500 nM of **32** for 4 h. **b** Corresponding line profiles. Image data was smoothed with a 1-pixel low pass Gaussian filter. In total, 40 cells were examined from 2 independent experiments.

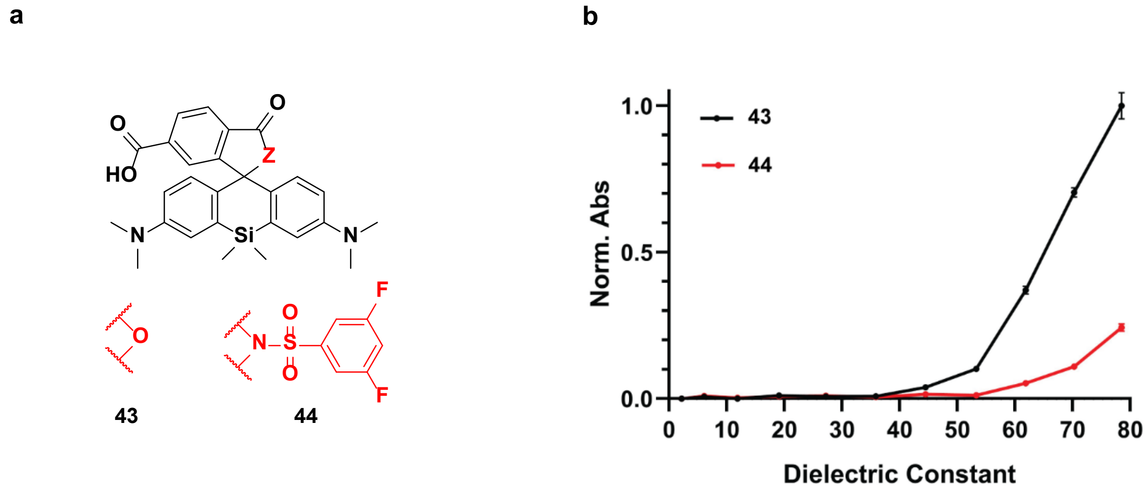


**Supplementary Figure 15:** Comparison of SiR derivatives (**43** and **44**). **a** Structures of SiR derivatives (**43** and **44**). **b** Normalized maximal absorbance of **43** and **44** (5 μM) in water–dioxane mixtures (v/v: 0/100 - 100/0) as a function of the dielectric constant.[1] The absorbance was normalized to the maximum absorbance of **43**. Error bars show ± s.d. from 3 experiments.

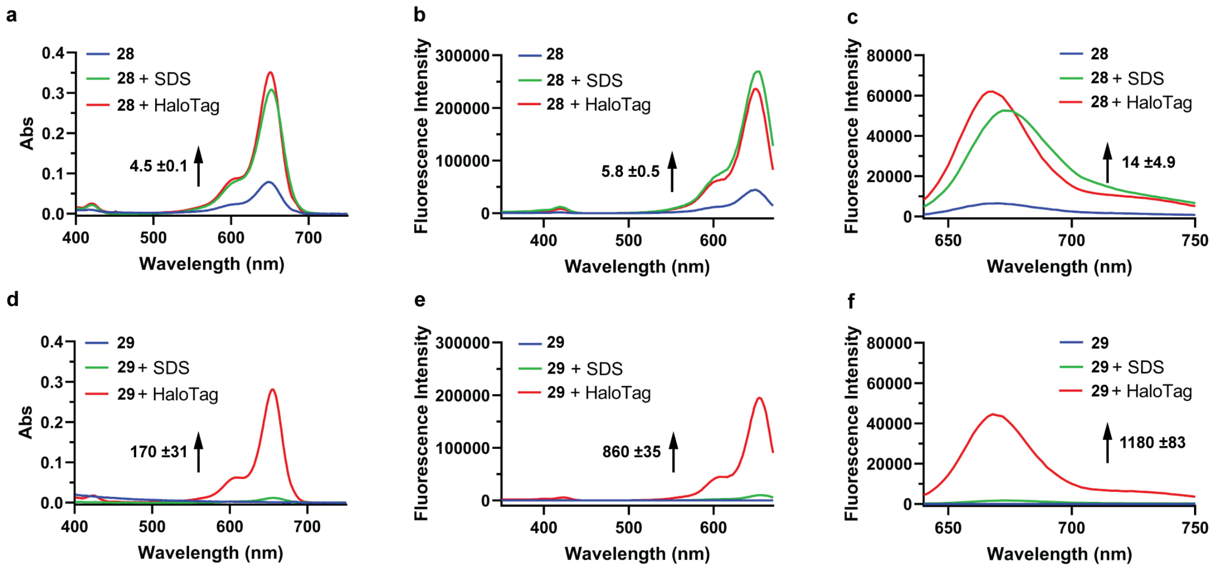


**Supplementary Figure 16:** Turn-on of SiR HaloTag probes (**28** and **29**) upon labeling of HaloTag and interaction with SDS. Absorption (**a** and **d**), excitation (**b** and **e**) and emission (**c** and **f**) spectra of probes **28** and **29** (2.5 µM) measured in HEPES buffer (pH 7.3) in the absence (blue line) and presence of HaloTag (5 µM, red line) or SDS (0.1%, green line) after 2.5 h incubation. The numbers represent the ratio of maximal absorbance and fluorescence intensities (emission: $\lambda_{ex}$: 610 nm, excitation $\lambda_{em}$: 690 nm) in the presence and absence of the HaloTag ($F/F_0$). Representative data for 3 experiments. Errors show ± s.d.

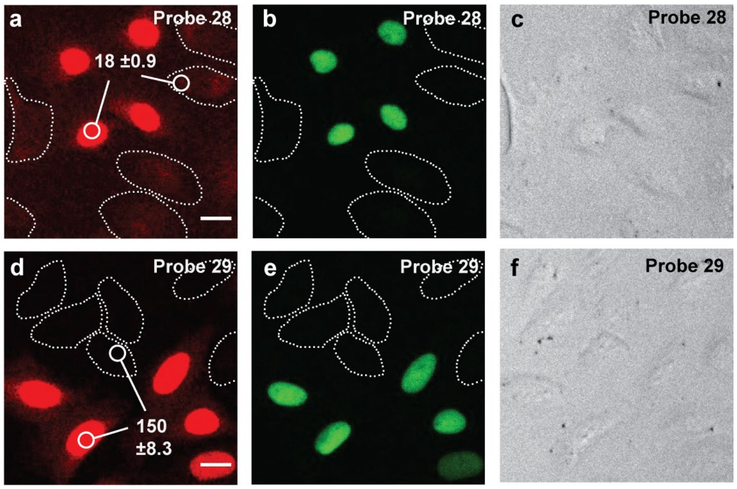


**Supplementary Figure 17:** Evaluation of fluorogenicity of SiR HaloTag probes (**28** and **29**) in live-cell, no-wash confocal microscopy. Co-cultured U-2 OS FlpIn Halo-SNAP-NLS expressing cells and wild-type U-2 OS cells were labeled with 500 nM **23** overnight, washed once with imaging medium and then labeled with 500 nM of **28** and **29** for 2.5 h. **a, d** Live-cell, no-wash confocal images (sum projections of z-stacks)

showing SiR fluorescence and **b, e** fluorescence of **23**. Wild-type U-2 OS cells are represented with dotted lines. **c, f** Bright field images. The numbers correspond to the fluorescence ratios $F_{nuc}/F_{cyt}$. $F_{nuc}$: Mean values of SiR fluorescence of ROIs within the nuclei of U-2 OS FlpIn Halo-SNAP-NLS expressing cells normalized to the fluorescence of **23**. $F_{cyt}$: Mean values of SiR fluorescence of ROIs within the cytosol of wild-type U-2 OS cells. In total, 180 cells were examined from 2 independent experiments for each probe. Errors represent ± s.e.m. Scale bar, 20 µm.

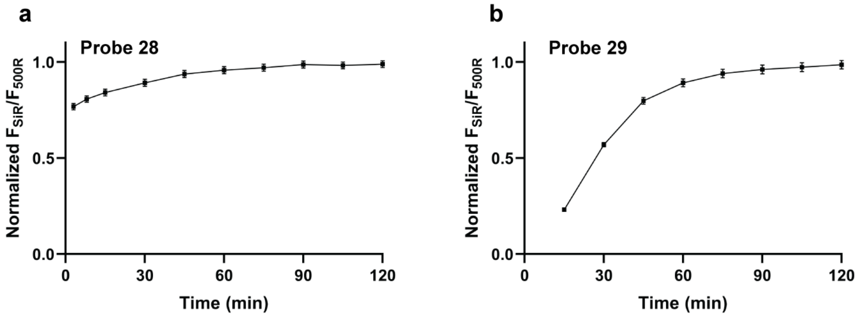


**Supplementary Figure 18:** Intracellular HaloTag labeling with 500 nM of SiR HaloTag probes (**28** and **29**) as a function of time. U-2 OS FlpIn Halo-SNAP-NLS expressing cells were pre-labeled with 500 nM **23** overnight. Normalized ratio of SiR fluorescence to the fluorescence signal of **23** is plotted at different time points. In total, 90 cells were examined from technical triplicates for each probe. Error bars show ± s.e.m.

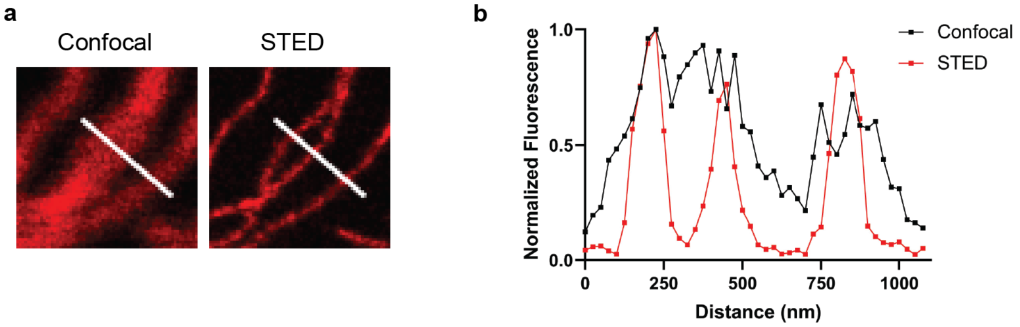


**Supplementary Figure 19:** Comparison of live-cell, no-wash confocal and STED microscopy for probe **29**. **a** Confocal and STED image (from **Figure 6d**) of U-2 OS Vimentin-Halo expressing cells labeled with 500 nM of **29** for 2 h. **b** Corresponding line profiles. Image data was smoothed with a 1-pixel low pass Gaussian filter. In total, 16 cells were examined from 2 independent experiments.

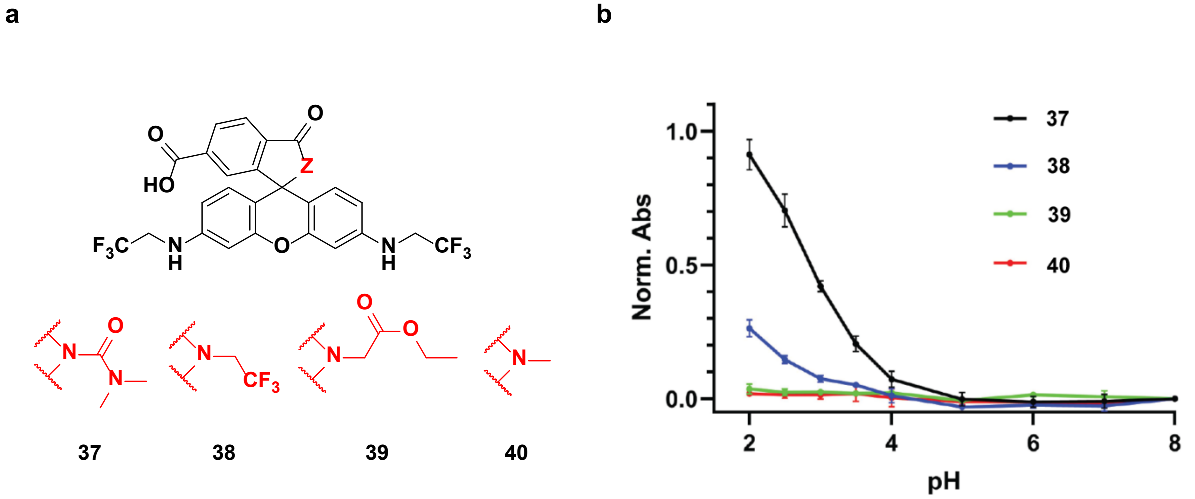


**Supplementary Figure 20:** Comparison of highly closed rhodamine *500R* derivatives (**37** - **40**). **a** Structures of highly closed rhodamine *500R* derivatives (**37** - **40**). **b** Normalized maximal absorbance of **37** - **40** (5 μM) in phosphate-buffered saline (PBS) (10 μM) at different pH. The absorbance was normalized to the maximum absorbance of **1** (5 μM) in PBS (10 μM) at pH = 7.0. Error bars show ± s.d. from 3 experiments.

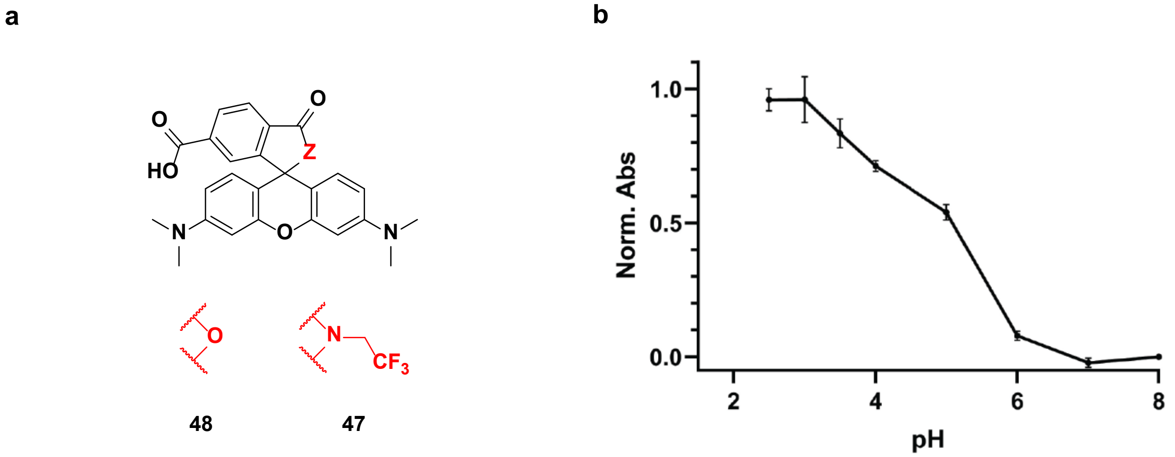


**Supplementary Figure 21:** pH titration of the highly closed TMR derivative (**47**). **a** Structures of TMR derivatives (**47** and **48**). **b** Normalized maximal absorbance of **47** (5 μM) in PBS (10 μM) at different pH. The absorbance was normalized to the maximum absorbance of **48** (5 μM) in PBS (10 μM) at pH = 7.0. Error bars show ± s.d. from 3 experiments.

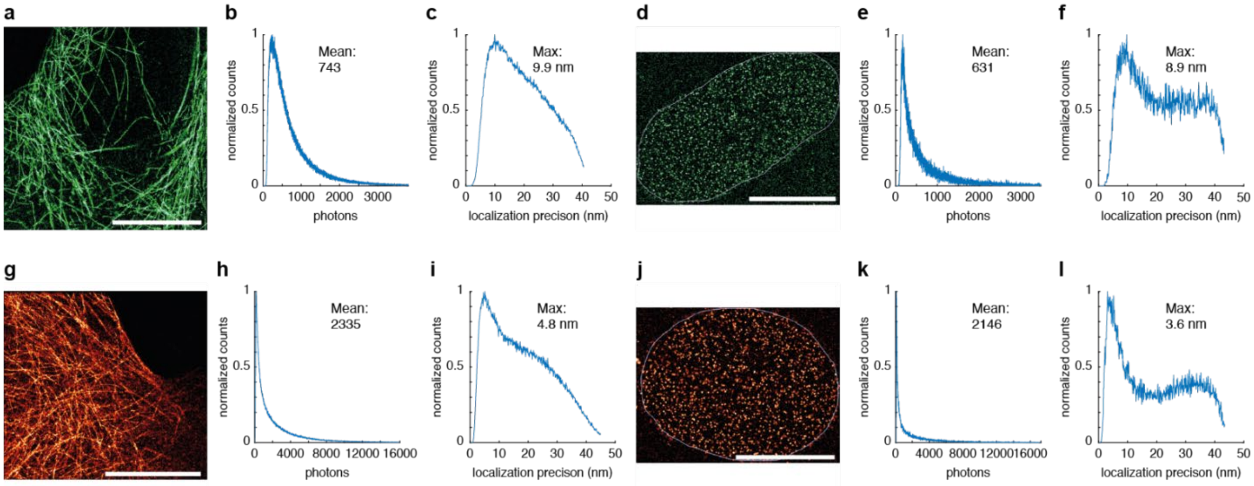


**Supplementary Figure 22:** Localization precision and photon count from single molecule localization microscopy (SMLM) with probe **26** and **33**. **a – c** Analysis of **26** blinking properties in U-2 OS Cep41-Halo cells. **a** Super-resolution image. **b** Photon count (mean: 743). **c** Localization precision (max: 9.9 nm). **d – f** Analysis of **26** blinking properties in U-2 OS Nup96-Halo cells. **d** Super-resolution image. **e** Photon count (mean: 631). **f** Localization precision (max: 8.9 nm). **g – i** Analysis of **33** blinking properties in U-2 OS Cep41-Halo cells. **g** Super-resolution image. **h** Photon count (mean: 2335). **i** Localization precision (max: 4.8 nm). **j – l** Analysis of **33** blinking properties in U-2 OS Cep41-Halo cells. **j** Super-resolution image. **k** Photon count (mean: 2146). **l** Localization precision (max: 3.6 nm). Scale bars, 10 µm. No filters were applied to the localizations for these reconstructions and analyses.

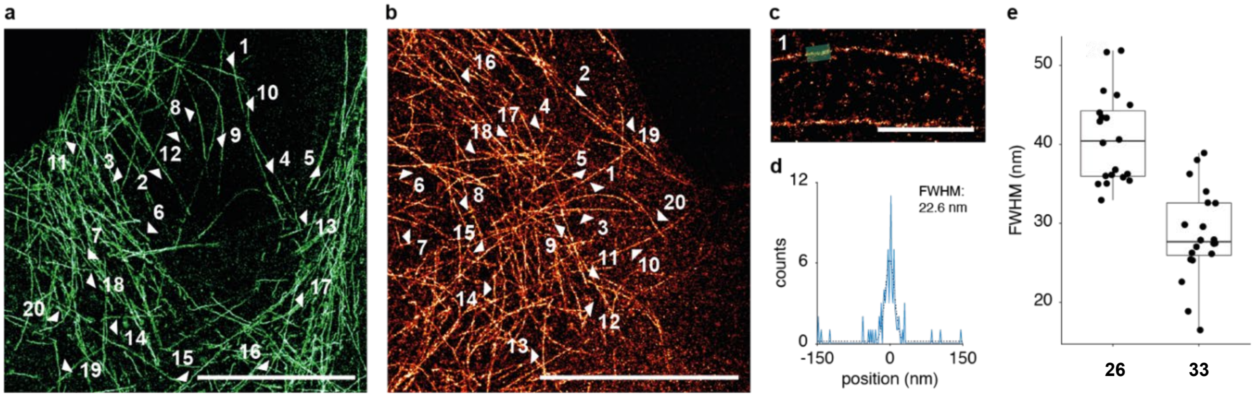


**Supplementary Figure 23:** Microtubule diameter estimation from SMLM with probe **26** and **33**. **a** Super-resolution image of U-2 OS Cep41-Halo cells, labeled with **26**. Arrows indicate the location of diameter estimation. Scale bar, 10 µm. **b** Super-resolution image of U-2 OS Cep41-Halo cells, labeled with **33**. Arrows indicate the location of diameter estimation. Scale bar, 10 µm. **c** Enlarged region of line profile 1 in **b**, highlighted in green. Scale bar, 1 µm. **d** Example histogram (blue) of the line profile marked in **c**. A Gaussian fit (dotted line) was used to estimate the full-width-half-maximum (FWHM) of 22.6 nm. **e** Boxplots depicting the distribution of FWHM for both **26** (mean: 40.7 nm, sd: 5.7 nm) and **33** (mean: 28.5 nm, sd: 5.8 nm). box: 25th–75th percentile, whiskers: 5th–95th percentile, black line: median, individual data points in black, N = 20 tubules per data set.

**Supplementary Movies**

Movie 1: Raw camera frames of **26** in U-2 OS Nup96-Halo cells. Frame rate: 100 fps.

Movie 2: Raw camera frames of **26** in U-2 OS Cep41-Halo cells. Frame rate: 100 fs.

Movie 3: Raw camera frames of **33** in U-2 OS Nup96-Halo cells. Frame rate: 100 fps.

Movie 4: Raw camera frames of **33** in U-2 OS Cep41-Halo cells. Frame rate: 100 fps.

**Supplementary Table 1:** Photophysical properties of rhodamine *500R* derivatives.

| Probe | Z | $\lambda_{abs}$ / $\lambda_{em}$ | ε ($M^{-1}cm^{-1}$) | Φ | $D_{50}$ | $F/F_0$ | $F_{nuc}/F_{cyt}$ |
|---|---|---|---|---|---|---|---|
| **1** | O | $501_a$ / $527_a$ | $82000_a$ | $0.88_a$ | $34_d$ | -- | -- |
| **4** | $NSO_2PhCN$ | $506_a$ / $529_a$ | $79000_a$ | $0.89_a$ | $61_d$ | -- | -- |
| **5** | $NSO_2PhF_2$ | $506_a$ / $529_a$ | $79000_a$ | $0.90_a$ | $63_d$ | -- | -- |
| **6** | $NSO_2PhCF_3$ | $506_a$ / $529_a$ | $74000_a$ | $0.91_a$ | $65_d$ | -- | -- |
| **7** | $NSO_2PhCl$ | $507_a$ / $529_a$ | $73000_a$ | $0.91_a$ | $68_d$ | -- | -- |
| **34** | NCN | $504_a$ / $529_a$ | $71000_a$ | 0.86 | $69_d$ | -- | -- |
| **8** | $NSO_2PhF$ | $505_a$ / $529_a$ | $72000_a$ | $0.91_a$ | $70_d$ | -- | -- |
| **9** | NSO2Ph | $506_a$ / $529_a$ | $67000_a$ | $0.90_a$ | $72_d$ | -- | -- |
| **10** | $NSO_2PhCH_3$ | $506_a$ / $527_a$ | $49000_a$ | $0.89_a$ | $74_d$ | -- | -- |

| | | | | | | | |
|---|---|---|---|---|---|---|---|
| **11** | $NSO_2PhOCH_3$ | $506_a$ / $529_a$ | $47000_a$ | $0.90_a$ | $76_d$ | -- | -- |
| **35** | $NSO_2CH_3$ | $505_a$ / $529_a$ | $39000_a$ | $0.89_a$ | $78_d$ | -- | -- |
| **36** | $NSO_2N(CH_3)_2$ | $505_a$ / $527_a$ | $7800_a$ | -- | $>79_d$ | -- | -- |
| **12** | O | $509_b$ / $531_b$ | $42000_a$ / $67000_b$ / $76000_c$ | $0.85_b$ / $0.95_c$ | -- | $1.2 \pm 0.02_e$ | $2.7 \pm 0.1_f$ |
| **13** | $NSO_2PhCN$ | $514_b$ / $533_b$ | $2900_a$ / $61000_b$ / $66000_c$ | $0.90_b$ / $0.97_c$ | -- | $24 \pm 1.2_e$ | $3.8 \pm 0.2_f$ |
| **14** | $NSO_2PhF_2$ | $514_b$ / $533_b$ | $2400_a$ / $63000_b$ / $67000_c$ | $0.92_b$ / $0.97_c$ | -- | $33 \pm 1.1_e$ | $7.4 \pm 0.4_f$ |
| **15** | $NSO_2PhCF_3$ | $514_b$ / $533_b$ | $1400_a$ / $56000_b$ / $69000_c$ | $0.90_b$ / $0.96_c$ | -- | $51 \pm 4.3_e$ | $9.6 \pm 0.4_f$ |
| **16** | $NSO_2PhCl$ | $515_b$ / $533_b$ | $1300_a$ / $54000_b$ / $63000_c$ | $0.90_b$ / $0.97_c$ | -- | $62 \pm 1.1_e$ | $9.9 \pm 0.4_f$ |
| **41** | NCN | $512_b$ / $533_b$ | $3200_a$ / $59000_b$ / $63000_c$ | $0.88_b$ / $0.94_c$ | -- | $20 \pm 0.6_e$ | $2.4 \pm 0.2_f$ |
| **17** | $NSO_2PhF$ | $514_b$ / $533_b$ | $1300_a$ / $54000_b$ / $64000_c$ | $0.92_b$ / $0.97_c$ | -- | $56 \pm 1.7_e$ | $11 \pm 0.5_f$ |
| **18** | NSO2Ph | $513_b$ / $533_b$ | $1100_a$ / $49000_b$ / $63000_c$ | $0.92_b$ / $0.98_c$ | -- | $73 \pm 3.1_e$ | $15 \pm 0.8_f$ |
| **19** | $NSO_2PhCH_3$ | $514_b$ / $533_b$ | $830_a$ / $45000_b$ / $42000_c$ | $0.90_b$ / $0.97_c$ | -- | $133 \pm 6.3_e$ | $25 \pm 1.2_f$ |
| **20** | $NSO_2PhOCH_3$ | $514_b$ / $533_b$ | $770_a$ / $43000_b$ / $45000_c$ | $0.94_b$ / $0.97_c$ | -- | $117 \pm 3.8_e$ | $24 \pm 1.1_f$ |
| **42** | $NSO_2CH_3$ | $513_b$ / $533_b$ | $2300_a$ / $45000_b$ / $53000_c$ | $0.91_b$ / $0.95_c$ | -- | $28 \pm 2.8_e$ | $13 \pm 0.7_f$ |
| **21** | $NSO_2N(CH_3)_2$ | $513_b$ / $531_b$ | $580_a$ / $13000_b$ / $20000_c$ | $0.91_c$ | -- | $87 \pm 2.9_e$ | $22 \pm 1.1_f$ |
| **22** | O | $505_b$ / $527_b$ | $19000_a$ / $54000_b$ / $73000_c$ | $0.74_b$ / $0.92_c$ | -- | $6.6 \pm 1.6_e$ | $3.4 \pm 0.3_f$ |
| **23** | $NSO_2PhF$ | $509_b$ / $529_b$ | $2800_a$ / $28000_b$ / $64000_c$ | $0.78_b$ / $0.94_c$ | -- | $72 \pm 6.5_e$ | $25 \pm 1.8_f$ |
| **24** | $NSO_2PhCH_3$ | $509_b$ / $527_b$ | $1500_a$ / $9900_b$ / $43000_c$ | $0.95_c$ | -- | $62 \pm 1.2_e$ | -- |

$_a$ HEPES buffer (pH 7.3), $_b$ binding with SNAP- and HaloTag (2 eq.) in HEPES buffer (pH 7.3), $_c$ 0.1% SDS in HEPES buffer (pH 7.3), $_d$ dioxane-$H_2O$ mixture (v/v: 100/0 – 0/100), $_e$ ratio of maximum fluorescence intensities in the presence and absence of SNAP- and HaloTag (2 eq.); errors show ± s.d. from 3 experiments, $_f$ fluorescence ratios; $F_{nuc}$: Mean values of rhodamine *500R* fluorescence of ROIs within the nuclei of U-2 OS FlpIn Halo-SNAP-NLS expressing cells normalized to the SiR fluorescence; $F_{cyt}$: Mean values of rhodamine *500R* fluorescence of ROIs within the cytosol of wild-type U-2 OS cells; in total, 180 cells from 2 - 3 independent experiments were examined for each probe; errors represent ± s.e.m.

**Supplementary Table 2:** Photophysical properties of CPY derivatives.

| **Probe** | **Z** | **$\lambda_{abs}$ / $\lambda_{em}$** | **ε ($M^{-1}cm^{-1}$)** | **Φ** | **$D_{50}$** | **$F/F_0$** | **$F_{nuc}/F_{cyt}$** |
|---|---|---|---|---|---|---|---|
| **45** | O | $609_a$ / $636_a$ | $116000_a$ | $0.51_a$ | $40_d$ | -- | -- |
| **46** | $NSO_2PhF$ | $612_a$ / $634_a$ | $74000_a$ | $0.59_a$ | $72_d$ | -- | -- |
| **30** | O | $614_b$ / $638_b$ | $66000_a$ / $112000_b$ / $149000_c$ | $0.54_b$ / $0.67_c$ | -- | $3.0 \pm 0.1_e$ | $19 \pm 0.8_f$ |
| **32** | $NSO_2PhF$ | $616_b$ / $636_b$ | $5900_a$ / $50000_b$ / $59000_c$ | $0.56_b$ / $0.72_c$ | -- | $58 \pm 1.6_e$ | $65 \pm 4.1_f$ |
| **31** | $NSO_2NMe_2$ | $616_b$ / $636_b$ | $420_a$ / $2700_b$ / $7400_c$ | -- | -- | $22 \pm 0.4_e$ | -- |

$_a$ HEPES buffer (pH 7.3), $_b$ binding with SNAP-tag (2 eq.) in HEPES buffer (pH 7.3), $_c$ 0.1% SDS in HEPES buffer (pH 7.3), $_d$ dioxane-$H_2O$ mixture (v/v: 100/0 – 0/100), $_e$ ratio of maximum fluorescence intensities in the presence and absence of SNAP-tag (2 eq.); errors show ± s.d. from 3 experiments, $_f$ fluorescence ratios; $F_{nuc}$: Mean values of CPY fluorescence of ROIs within the nuclei of U-2 OS FlpIn Halo-SNAP-NLS expressing cells normalized to the fluorescence of **19**; $F_{cyt}$: Mean values of CPY fluorescence of ROIs within the cytosol of wild-type U-2 OS cells; in total, 180 cells from 2 independent experiments were examined for each probe; errors represent ± s.e.m.

**Supplementary Table 3:** Photophysical properties of SiR derivatives.

| **Probe** | **Z** | **$\lambda_{abs}$ / $\lambda_{em}$** | **ε ($M^{-1}cm^{-1}$)** | **Φ** | **$D_{50}$** | **$F/F_0$** | **$F_{nuc}/F_{cyt}$** |
|---|---|---|---|---|---|---|---|

| | | | | | | | |
|---|---|---|---|---|---|---|---|
| **43** | O | 646$_a$ / 668$_a$ | 105000$_a$ | 0.41$_a$ | 65$_d$ | -- | -- |
| **44** | $NSO_2PhF_2$ | 648$_a$ / 666$_a$ | 19000$_a$ | 0.47$_a$ | >79$_d$ | -- | -- |
| **28** | O | 651$_b$ / 668$_b$ | 31000$_a$ / 139000$_b$ / 123000$_c$ | 0.49$_b$ / 0.53$_c$ | -- | 14 ±4.9$_e$ | 18 ±0.9$_f$ |
| **29** | $NSO_2PhF_2$ | 655$_b$ / 668$_b$ | 650$_a$ / 113000$_b$ / 4700$_c$ | 0.57$_b$ | -- | 1180 ±83$_e$ | 150 ±8.3$_f$ |

$_a$ HEPES buffer (pH 7.3), $_b$ binding with HaloTag (2 eq.) in HEPES buffer (pH 7.3), $_c$ 0.1% SDS in HEPES buffer (pH 7.3), $_d$ dioxane-$H_2O$ mixture (v/v: 100/0 – 0/100), $_e$ ratio of maximum fluorescence intensities in the presence and absence of HaloTag (2 eq.); errors show ± s.d. from 3 experiments, $_f$ fluorescence ratios; $F_{nuc}$: Mean values of SiR fluorescence of ROIs within the nuclei of U-2 OS FlpIn Halo-SNAP-NLS expressing cells normalized to the fluorescence of **23**; $F_{cyt}$: Mean values of SiR fluorescence of ROIs within the cytosol of wild-type U-2 OS cells; in total, 180 cells from 2 independent experiments were examined for each probe; errors represent ± s.e.m.

**Supplementary Table 4:** Photophysical properties of highly closed rhodamine *500R* and TMR derivatives.

| **Probe** | **Z** | $\lambda_{abs}$ | $\lambda_{em}$ | **ε ($M^{-1}cm^{-1}$)** | **Φ** |
|---|---|---|---|---|---|
| **37** | $NCON(CH_3)_2$ | 514$_a$ | 539$_a$ | 75000$_a$ | 0.94$_a$ |
| **38** | $NCH_2CF_3$ | 512$_a$ | 535$_a$ | 26000$_a$ | 0.94$_a$ |
| **47** | $NCH_2CF_3$ | 559$_b$ | 582$_b$ | 80000$_b$ | 0.39$_b$ |

$_a$ PBS buffer (pH 2.0), $_b$ PBS buffer (pH 3.0).

### Supplementary Information - *In-Vitro* Tests, Cell Culture and Microscopy

#### Absorbance, Emission, Excitation Spectra and Quantum Yield Measurements

Fluorophores for spectroscopy were prepared as stock solutions (1.5 – 10 mM) in DMSO and diluted such that the DMSO concentration did not exceed 0.5% (v/v). Absorption spectra were recorded on a V-770 spectrophotometer (Jasco) in a Quartz cuvette. Emission and excitation spectra were measured on a Spark® microplate reader (Tecan) using a 96-well plate (Thermo Scientific Nunc with optical bottom). All measurements were performed at 25 °C. Maximum absorption wavelength ($\lambda_{abs}$) and maximum emission wavelength ($\lambda_{em}$) were determined in 50 mM HEPES buffer containing 50 mM NaCl (pH 7.3) or in 10 μM PBS buffer (pH 2.0, 3.0). Absolute fluorescence quantum yields were recorded on a Quantaurus-QY spectrometer (model C11347, Hamamatsu). Measurements were carried out using diluted samples (A ≈ 0.1). The reported values for molar extinction coefficient (ε) and quantum yield (Φ) are averages (N = 3).

For absorbance, emission, excitation spectra and quantum yield measurements upon labeling of HaloTag and SNAP-tag, SNAP-Halo protein was diluted into HEPES buffer (50 mM HEPES, 50 mM NaCl, pH 7.3) to a final concentration of 5 μM. Fluorescent probes were diluted therein to a final concentration of 2.5 μM. The obtained mixtures were incubated until a consistent absorbance signal was observed (2.5 h). As a control experiment, the fluorescent probes (2.5 μM) were incubated together with 0.1% SDS in 50 mM HEPES buffer containing 50 mM NaCl (pH 7.3). The reported values are averages (N = 3).

#### Absorbance Measurements in Water-Dioxane Mixtures

Water-dioxane mixtures containing 0%, 10%, 20%, 30%, 40%, 50%, 60%, 70%, 80%, 90% and 100% anhydrous dioxane (v/v) were prepared. The fluorophores were diluted in these water-dioxane mixtures to 5 μM and the absorbance spectra were measured on a Spark® microplate reader (Tecan) using a 96-well plate (Greiner Bio-One) at 25 °C. The maximal absorbance values of the different fluorophores were normalized to the maximal absorbance of the corresponding *ortho*-carboxylate derivative in 100% water. The normalized absorbance values were plotted against the dielectric constants of the different water-dioxane mixtures.[1] The reported values are averages (N = 3).

#### Absorbance Measurements at various pH

The pH values of PBS (10 μM) solutions were adjusted by addition of NaOH or HCl by means of a pH meter to 2.0, 2.5, 3.0, 3.5, 4.0, 5.0, 6.0, 7.0 and 8.0. The fluorophores were diluted in these PBS mixtures to 5 μM and the absorbance spectra were measured on a Spark® microplate reader (Tecan) using a 96-well plate (Greiner Bio-One) at 25 °C. The maximal absorbance values of the different fluorophores were normalized to the maximal absorbance of the corresponding *ortho*-carboxylate derivative in PBS (10 μM, pH 7.0). The reported values are averages (N = 3).

#### Cloning, Protein Expression and Purification

SNAP-Halo fusion protein was cloned in a pET51b(+) vector (Novagen) for production in *Escherichia coli*, featuring an N-terminal StrepTag-II and an enterokinase cleavage site together with a C-terminal $His_{10}$ tag. Cloning was performed by Gibson assembly[3] and electroporated in E.cloni 10G cells (Lucigen). The protein was expressed in the *E. coli* strain BL21(DE3)-pLysS (Novagen) in lysogeny broth (LB)[4] cultures grown at 37°C to an optical density at 600 nm ($OD_{600nm}$) of 0.8. Protein expression was induced by the addition of 0.5 mM isopropyl-β-D-thiogalactopyranoside (IPTG) and cells were grown at 17°C overnight in the presence of 1 mM $MgCl_2$. Cells were harvested by centrifugation (4,500 g, 10 min, 4ºC) and lysed by sonication. The cell lysate was cleared by centrifugation (75,000g, 4ºC, 10 min) and the protein was purified using HisPur Ni-NTA Superflow Agarose (Thermo Fisher Scientific, Waltham, MA, USA) by batch incubation followed by washing and elution steps on a polypropylene column (Qiagen). The proteins were subsequently purified using a StrepTrap HP column (Cytiva) on an ÄktaPure FPLC followed by a buffer exchange using a HiPrep 26/10 Desalting column (Cytiva) to HEPES 50 mM, NaCl 50 mM pH 7.3 (*i.e.* activity buffer). Proteins were concentrated using Ultra-15 mL centrifugal filter devices (Amicon, Merck KGaA, Darmstadt, Germany) with a molecular weight cut-off (MWCO) smaller than the protein size to a final concentration of 500 µM. Proteins were aliquoted and stored at -80°C after flash freezing in liquid nitrogen. Correct size and purity of proteins were assessed by SDS-PAGE and liquid chromatography-mass spectrometry (LC-MS) analysis.

**Cell Culture, Labeling and Fixation**

U-2 OS cells (wild-type), U-2 OS FlpIn Halo-SNAP-NLS[5], Vimentin-HaloTag[6] and Vimentin-SNAP-tag[7] expressing cells were cultured in Dulbecco's Modified Eagle Medium (DMEM, 4.5 g/L glucose) supplemented with 10% (v/v) fetal bovine serum (FBS), GlutaMAX and sodium pyruvate (all Life Technologies) in a humidified 5% $CO_2$ incubator at 37 °C. Cells were split every 2 – 4 days or at confluency and regularly tested for mycoplasma contamination. Cells were seeded on 10-well glass bottom plates (Greiner Bio-One), 96-well glass bottom plates (Eppendorf) or glass coverslips 1 – 2 days before imaging. Prior to imaging, cells were labeled (see respective experiments for details) in imaging medium (phenol-red free DMEM supplemented with GlutaMAX, sodium pyruvate and 10% (v/v) FBS (all Life Technologies)).

U-2 OS Nup96-Halo cells[8] were cultured in DMEM (catalog no. 11880-02, Gibco) growth medium containing 1x MEM NEAA (catalog no. 11140-035, Gibco), 1x GlutaMAX (catalog no. 35050-038, Gibco) and 10% (v/v) fetal bovine serum (catalog no. 10270-106, Gibco). Stable U-2 OS Flp-In T-Rex Cep41-Halo cells[9] were cultured in DMEM/F-12 (catalog no. 10565, Invitrogen) growth medium containing 10% (v/v) fetal bovine serum (catalog no. 10270-106, Gibco) and 1x ZellShield (Minerva Labs). Both cell lines were cultured in a humidified incubator at 37°C and 5% $CO_2$ and split every 2 – 3 days to ensure confluency at approximately 60 – 70%. 2 days before imaging, cells were seeded at 60 – 70% confluency onto 24 mm round glass coverslips (No. 1.5H, catalog no. 117640, Marienfeld). The coverslips were previously cleaned by overnight incubation in a MeOH:HCl mixture (50:50) while stirring. Then, they were rinsed with water until a neutral pH was reached and placed in a laminar flow hood overnight to dry, followed by a 30 min UV sterilization step. In the case of the U-2 OS Cep41-Halo cells, expression was induced by addition of 1 µg $mL^{-1}$ doxycycline (D9891-1G, Sigma-Aldrich) to the growth medium. After 24 h, cells where incubated overnight at 37°C and 5% $CO_2$ in their respective growth medium containing 1 µM dye (**26** or **33**), followed by a 2 h incubation in growth medium without dye. The samples were then prefixed at room temperature (rt) for 30 s in 2.4% (w/v) formaldehyde (FA) in PBS. Permeabilization was achieved by incubating the sample at rt in 0.4% (v/v) Triton X-100 (Sigma) in PBS for 3 min before completing the fixation process at rt for 30 min in 2.4% (w/v) FA in PBS. The fixation was subsequently quenched by incubating the coverslips in 100 mM $NH_4Cl$ (Sigma) in PBS. Sample preparation was finalized by washing three times in PBS for 5 min each.

#### Confocal Microscopy

Confocal microscopy was performed on a Leica DMi8 microscope (Leica Microsystems) equipped with a Leica TCS SP8 X scanhead, a SuperK white light laser, a HC PL APO CS2 20.0 x/0.75 objective and an incubator with $CO_2$ as well as temperature control (Life Imaging Services, 5%, 37 °C).

#### Time-Dependent Intracellular HaloTag and SNAP-tag Labeling

For rhodamine *500R* derived HaloTag probes (**12** – **21** and **42**), U-2 OS FlpIn Halo-SNAP-NLS expressing cells were pre-labeled with 500 nM of SiR-BG (overnight) and washed with imaging medium. Directly after labeling with 200 nM of **12** – **21** and **42**, images were recorded every minute over 45 min. Microscopy conditions: $\lambda_{ex}$: 510 nm and 645 nm, detection range: 520 – 600 nm and 655 – 720 nm, image size: 580.31 µm x 580.31 µm, pixel size: 567 nm, pixel dwell time: 0.86 µs. Mean values of rhodamine *500R* fluorescence of regions of interest (ROIs) within the nuclei were divided by the mean values of SiR fluorescence at the different time points. >51 cells were examined for each probe from technical triplicates.

For rhodamine *500R* derived SNAP-tag probes (**22** - **24**), U-2 OS FlpIn Halo-SNAP-NLS expressing cells were pre-labeled with 200 nM of SiR-Halo (overnight). Directly after labeling with 500 nM of **22** - **24**, z-stacks (stack size: 20 µm, number of steps: 30) were recorded every 0.5 h over 6 h. Microscopy conditions: $\lambda_{ex}$: 510 nm and 645 nm, detection range: 520 – 600 nm and 655 – 720 nm, image size: 580.31 µm x 580.31 µm, pixel size: 567 nm, pixel dwell time: 0.86 µs. The summed stacks were analyzed. Mean values of rhodamine *500R* fluorescence of ROIs within the nuclei were divided by the mean values of SiR fluorescence at the different time points. In total, 90 cells were examined for each probe in 2 independent experiments.

For CPY derived SNAP-tag probes (**30** and **32**), U-2 OS FlpIn Halo-SNAP-NLS expressing cells were pre-labeled with 200 nM of **19** (overnight). Directly after labeling with 500 nM of **30** and **32**, z-stacks (stack size: 20 µm, number of steps: 30) were recorded every 0.5 h over 5 h. Microscopy conditions: $\lambda_{ex}$: 510 nm and 615 nm, detection range: 520 – 600 nm and 625 – 700 nm, image size: 580.31 µm x 580.31 µm, pixel size: 567 nm, pixel dwell time: 0.86 µs. The summed stacks were analyzed. Mean values of CPY fluorescence of ROIs within the nuclei were divided by the mean fluorescence values of **19** at the different time points. In total, 90 cells were examined for each probe from technical triplicates.

For SiR derived HaloTag probes (**28** and **29**), U-2 OS FlpIn Halo-SNAP-NLS expressing cells were pre-labeled with 500 nM of **23** (overnight) and washed with imaging medium. Directly after labeling with 500 nM of **28** and **29**, z-stacks (stack size: 20 µm, number of steps: 30) were recorded every 15 min over 2 h. For **28**, further z-stacks were measured 3 and 8 min after labeling. Microscopy conditions: $\lambda_{ex}$: 510 nm and 645 nm, detection range: 520 – 600 nm and 655 – 720 nm, image size: 580.31 µm x 580.31 µm, pixel size: 567 nm, pixel dwell time: 0.86 µs. The summed stacks were analyzed. Mean values of SiR fluorescence of ROIs within the nuclei were divided by the mean fluorescence values of **23** at the different time points. In total, 90 cells were examined for each probe from technical triplicates.

**Evaluation of Fluorogenicity - Live-Cell, No-Wash Confocal Microscopy**

For rhodamine *500R* derived HaloTag probes (**12** – **21**, **41** and **42**), co-cultured U-2 OS FlpIn Halo-SNAP-NLS expressing cells and wild-type U-2 OS cells (1:1) were pre-labeled with 500 nM of SiR-BG (overnight) and washed with imaging medium. 2.5 h after labeling with 200 nM **12** – **21**, **41** and **42**, z-stacks (stack size: 30 µm, number of steps: 40) were recorded. Microscopy conditions: $\lambda_{ex}$: 510 nm and 645 nm, detection range: 520 – 600 nm and 655 – 720 nm, image size: 446.02 µm x 446.02 µm, pixel size: 436 nm, pixel dwell time: 0.86 µs. The summed stacks were analyzed to measure the ratios between nuclear signal ($F_{nuc}$) and cytosolic background signal ($F_{cyt}$) of **12** – **21**, **41** and **42**. $F_{nuc}$: Mean values of rhodamine *500R* fluorescence of ROIs within the nuclei of U-2 OS FlpIn Halo-SNAP-NLS expressing cells normalized to the SiR fluorescence. $F_{cyt}$: Mean values of rhodamine *500R* fluorescence of ROIs within the cytosol of wild-type U-2 OS cells. Bright field images were used to locate wild-type U-2 OS cells represented by dotted lines (**Figures 3d** and **Supplementary Figure 6**). In total, 180 cells (90 U-2 OS FlpIn Halo-SNAP-NLS expressing cells and 90 wild-type U-2 OS cells) were examined from 3 independent experiments for each probe.

For rhodamine *500R* derived SNAP-tag probes (**22** and **23**), co-cultured U-2 OS FlpIn Halo-SNAP-NLS expressing cells and wild-type U-2 OS cells (1:1) were pre-labeled with 200 nM of SiR-Halo (overnight) and washed with imaging medium. 5 h after labeling with 500 nM of **22** and **23**, z-stacks (stack size: 30 µm, number of steps: 40) were recorded. Microscopy conditions: $\lambda_{ex}$: 510 nm and 645 nm, detection range: 520 – 600 nm and 655 – 720 nm, image size: 580.31 µm x 580.31 µm, pixel size: 567 nm, pixel dwell time: 0.86 µs. The summed stacks were analyzed as described above. In total, 180 cells (90 U-2 OS FlpIn Halo-SNAP-NLS expressing cells and 90 wild-type U-2 OS cells) were examined from 2 independent experiments for each probe.

For CPY derived SNAP-tag probes (**30** and **32**), co-cultured U-2 OS FlpIn Halo-SNAP-NLS expressing cells and wild-type U-2 OS cells (1:1) were pre-labeled with 200 nM of **19** (overnight) and washed with imaging medium. 4 h after labeling with 500 nM of **30** and **32**, z-stacks (stack size: 30 µm, number of steps: 40) were recorded. Microscopy conditions: $\lambda_{ex}$: 510 nm and 615 nm, detection range: 520 – 600 nm and 625 – 700 nm, image size: 580.31 µm x 580.31 µm, pixel size: 567 nm, pixel dwell time: 0.86 µs. The summed stacks were analyzed as described above. In total, 180 cells (90 U-2 OS FlpIn Halo-SNAP-NLS expressing cells and 90 wild-type U-2 OS cells) were examined from 2 independent experiments for each probe.

For SiR derived HaloTag probes (**28** and **29**), co-cultured U-2 OS FlpIn Halo-SNAP-NLS expressing cells and wild-type U-2 OS cells (1:1) were pre-labeled with 500 nM of **23** (overnight) and washed with imaging medium. 2.5 h after labeling with 500 nM of **28** and **29**, z-stacks (stack size: 30 µm, number of steps: 40) were recorded. Microscopy conditions: $\lambda_{ex}$: 510 nm and 645 nm, detection range: 520 – 600 nm and 655 – 720 nm, image size: 580.31 µm x 580.31 µm, pixel size: 567 nm, pixel dwell time: 0.86 µs. The summed stacks were analyzed as described above. In total, 180 cells (90 U-2 OS FlpIn Halo-SNAP-NLS expressing cells and 90 wild-type U-2 OS cells) were examined from 2 independent experiments for each probe.

**SMLM**

SMLM data were acquired on a custom-built widefield setup described previously.[10-11] Briefly, the free output of a commercial laser box (LightHub, Omicron-Laserage Laserprodukte) equipped with Luxx 488 and Cobolt 561 lasers were coupled into a square multi-mode fiber (catalog no. M103L05, Thorlabs). The fiber was agitated as described previously.[12] The output of the fiber was magnified by an achromatic lens and focused into the sample to homogeneously illuminate an area of about 700 µm$^2$. The laser was guided through a laser cleanup filter (390/482/563/640 HC Quad, AHF) to remove fluorescence generated by the fiber. The emitted fluorescence was collected through a high numerical aperture (NA) oil immersion objective (HCX PL APO 160×/1.43 NA, Leica), filtered by a 525/50 (catalog no. FF03-525/50-25, Semrock) bandpass filter (for imaging of **26**) or by a 600/60 (catalog no. NC458462, Chroma) bandpass filter (for imaging of **33**) on an EMCCD camera (Evolve 512, Photometrics). The z focus was stabilized by an infrared laser that was totally internally reflected off the coverslip onto a quadrant photodiode, which was coupled into closed-loop feedback with the piezo objective positioner (Physik Instrumente). The microscope hardware and data acquisition were handled via Micro-Manager 2.0 using custom-written software.[11, 13] Coverslips containing prepared samples were placed into a custom-built sample holder and 500 µL PBS were added. To image **26**, we used an excitation intensity at 488 nm of 17 kW cm$^{-2}$. For **33** we used an excitation intensity at 561 nm of 23 kW cm$^{-2}$. The exposure time was 10 ms in both cases.

**STED Microscopy**

STED imaging of (**19**) and (**29**) was performed on an Abberior easy3D STED/RESOLFT QUAD scanning expert line microscope (Abberior Instruments GmbH, Göttingen, Germany) built on a motorized inverted microscope IX83 (Olympus, Tokyo, Japan). The microscope is equipped with pulsed STED lasers at 595 nm and 775 nm, 355 nm, 405 nm, 485 nm, 561 nm, and 640 nm excitation lasers, and with 488 nm and 405 nm RESOLFT lines. (**19**) was super-resolved using $\boldsymbol{\lambda_{ex}}$ 485 nm and $\boldsymbol{\lambda_{STED}}$ 595 nm, while (**29**) was super-resolved using $\boldsymbol{\lambda_{ex}}$ 640 nm the $\boldsymbol{\lambda_{STED}}$ 775 nm depletion line. Spectral detection was performed with avalanche photodiodes (APD) in the following spectral windows: 505 – 580 nm (**19**), 650 – 770 nm (**29**). Images were acquired with a 100x/1.40 UPlanSApo Oil immersion objective lens (Olympus). Pixel size was 33 nm (**19**) and 25 nm (**29**). Laser powers and dwell times were optimized for each sample.

STED imaging of (**32**) was performed on an Abberior STED QUAD scanning expert line (Abberior Instruments GmbH, Göttingen, Germany) built on a motorized inverted microscope IX83 (Olympus, Tokyo, Japan). The microscope is equipped with pulsed STED lasers at 655 nm and 775 nm, with 520 nm, 561 nm, 640 nm, and multiphoton (Chameleon Vision II, Coherent, Santa Clara, USA) excitation lasers. Spectral detection was performed with two avalanche photodiodes (APD) in the spectral window 580 – 750 nm. Images were acquired with a 60x/1.42 Oil immersion objective lens (Olympus). Pixel size was 30 nm. Laser powers and dwell times were optimized for each sample.

**Image Processing and Data Analysis**

Images were processed by means of ImageJ/Fiji unless otherwise stated.[14-15] Imaging data in **Figures 6d**, **g** and **Supplementary Figures 9**, **16** and **21** was smoothed with a 1-pixel low pass Gaussian filter in Imspector (Abberior Instruments GmbH, Göttingen Germany). The line profiles represented in **Supplementary Figures 9**, **16** and **21** were measured with 2-pixel wide lines in ImageJ/Fiji before smoothing.

SMLM data was fitted and analyzed as previously described using our custom-written, open-source super-resolution microscopy analysis platform $SMAP_{21}$ in MATLAB (Mathworks, 2020b).[16] The software is available at github.com/jries/SMAP.

The x, y positions were corrected for residual drift by a custom algorithm based on redundant cross-correlation. Localizations persistent over consecutive frames (detected within 35 nm from one another and with a maximum gap of one dark frame) were merged into one localization by calculating the weighted average of x and y positions and the sums of photons per localization and background photons. Localizations were filtered by the localization precision (0–20 nm) to exclude dim localizations, and by the fitted size of the PSF (0–150 nm laterally) to exclude localizations that were detected out-of-focus. Additionally, poorly fitted localizations were excluded if their log-likelihood (LL) was smaller than -1. Super-resolution images were constructed with every localization rendered as a two-dimensional elliptical Gaussian with a width proportional to the localization precision (factor 0.4). The reported average photon counts per localization were calculated based on these merged and filtered localizations.

For the quantification of microtubule width, we constructed a perpendicular line profile from a 250 nm long section of the microtubule. We then fitted a Gaussian distribution (bin width 2 nm). FWHM were then calculated using equation (1) where σ is the standard deviation of the Gaussian fit. Results were plotted in a boxplot (**Supplementary Figure 23**, N = 20 line profiles per dye).

$$(1)\ FWHM = 2\sqrt{2ln\,2}\sigma \approx 2.36\sigma$$

### Supplementary Information – Synthesis and Characterization

#### General Information, Materials and Equipment

All chemical reagents and anhydrous solvents for synthesis were purchased from commercial suppliers (Sigma-Aldrich, TCI, Alfa Aesar) and were used without further purification or distillation. Reactions were monitored by thin-layer chromatography (TLC) performed on TLC-aluminum sheets (Silica gel 60 F254, Merck) or liquid chromatograph-mass spectrometry (LCMS-2020, Shimadzu). The analytical reversed-phase LC-MS was equipped with an SPD-20AV UV-VIS photodiode array detector for product visualization on a C18 1.9 µm, 2.1 x 50 mm column (Supelco) applying 10–95% MeCN/$H_2O$, linear gradient with constant 0.1% v/v formic acid additive in a 6 min run with 1 mL/min flow rate. Concentration under reduced pressure was performed by rotary evaporation at 40°C. Preparative RP-HPLC was performed on an UltiMate 3000 system (Thermo Fisher Scientific) on a C18 5 µm, 10 x 250 mm column (Supelco, flow rate 4 mL/min) or a C18 5 µm, 21.2 x 250 mm column (Supelco, flow rate 8 mL/min), solvent A: 0.1% v/v TFA or formic acid in $H_2O$, solvent B: MeCN. The compounds purified by Prep-HPLC were lyophilized on a lyophilizer (Christ) equipped with a vacuum pump (Vacuubrand). Flash column chromatography was performed with silica gel (230-400 mesh, Silicycle) on an automated purification system (Biotage Isolera One). Yields refer to purified, dried and spectroscopically pure compounds. $^{1}H$ and $^{13}C$ nuclear magnetic resonance (NMR) spectra were recorded on a Bruker DPX 400 at room temperature. Chemical shifts (δ-values) are reported in ppm, spectra were calibrated relative to the residual proton chemical shifts ($CDCl_3$, δ = 7.26; DMSO-$d_6$, δ = 2.50; MeOD-$d_4$, δ = 3.31) and carbon chemical shifts ($CDCl_3$, δ = 77.16; DMSO-$d_6$, δ = 39.52; MeOD-$d_4$, δ = 49.00) of the solvents, multiplicity is reported as follows: s = singlet, d = doublet, t = triplet, q = quartet, m = multiplet or unresolved and coupling constant *J* in Hz.[17] High-resolution mass spectra (HRMS) were measured on a maXis II™ ETD with electron spray ionization (ESI) (Bruker).

### Synthetic Procedures

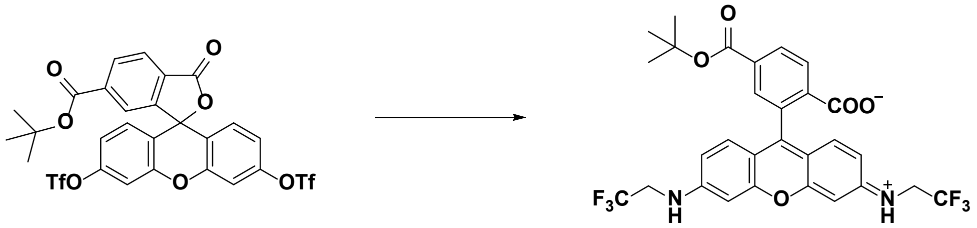

**3** **2**

**(*E*)-4-*tert*-Butoxycarbonyl)-2-(6-((2,2,2-trifluoroethyl)amino)-3-((2,2,2-trifluoro-ethyl)iminio)-*3H*-xanthen-9-yl)benzoate (2): 3** was synthesized according to a procedure reported in.[18] A *Schlenk* flask was dried with a heat gun *in vacuo* prior to dissolution of **3** (50.0 mg, 71.8 µmol, 1.0 eq) in dry 1,4-dioxane (2.5 mL). Potassium carbonate (47.6 mg, 345 µmol, 4.8 eq), 2,2,2-trifluoroethylamine (250 µL, 3.13 mmol, 43.6 eq), tris(dibenzylideneacetone)dipalladium (13.1 mg, 14.4 µmol, 0.20 eq) and 2-(dicyclohexylphosphino)-2',4',6'-triisopropylbiphenyl (10.3 mg, 21.5 µmol, 0.30 eq) were added. The reaction mixture was heated to 100°C and stirred at this temperature for 3 h. Subsequently, the mixture was allowed to cool down to rt and the solvent was evaporated under reduced pressure. The crude product was purified by flash column chromatography ($CH_2Cl_2$/MeOH 10:0 to 10:1) to give **2** (30.7 mg, 51.6 µmol, 72 %) as a red solid.

TLC: $R_f$ = 0.61 ($SiO_2$, $CH_2Cl_2$/MeOH 9:1).

$^1$H NMR (400 MHz, DMSO-$d_6$) δ 8.20 (d, *J* = 8.0, 1H), 8.09 (d, *J* = 8.0 Hz, 1H), 7.58 (s, 1H), 6.79 (t, *J* = 6.8 Hz, 2H), 6.65 (d, *J* = 1.8 Hz, 2H), 6.52 – 6.46 (m, 4H), 4.05 – 3.96 (m, 4H), 1.50 (s, 9H).

$^{13}$C NMR (101 MHz, DMSO-$d_6$) δ 167.9, 163.7, 152.5, 152.2, 150.0, 137.5, 130.7, 130.1, 129.9, 128.7, 127.1, 125.2, 124.3, 125.2, 123.9, 110.2, 106.5, 97.9, 84.6, 82.3, 44.1, 43.8, 43.5, 43.1, 27.6.

HRMS (ESI) Exact mass calculated for $C_{29}H_{24}N_2O_5F_6$ $[M+H]^+$: 595.1662, found: 595.1663.

#### General Procedure A for 9:

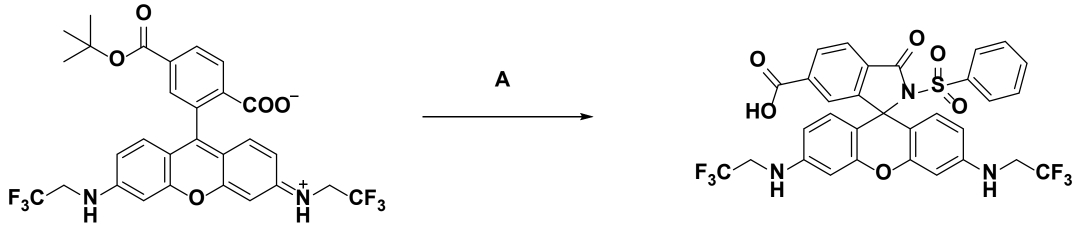

**2** **9**

**3-Oxo-2-(phenylsulfonyl)-3',6'-bis((2,2,2-trifluoroethyl)amino)spiro[isoindoline-1,9'-xanthene]-6-carboxylic acid (9):** A solution of **2** (4.00 mg, 6.73 µmol, 1.0 eq), benzenesulfonamide (5.29 mg, 33.6 µmol, 5 eq.), 1-(3-dimethylaminopropyl)-3-ethylcarbodiimide hydrochloride (5.16 mg, 26.9 µmol, 4 eq.) and 4-dimethylaminopyridine (3.29 mg, 26.9 µmol, 4 eq.) in $CH_2Cl_2$ (0.6 mL) was heated to 60°C and stirred at this temperature for 12 h in a sealed tube. $H_2O$ (1 mL) was added and the aqueous layer was extracted with $CH_2Cl_2$ (3x). The combined organic layers were dried over $MgSO_4$, filtered and concentrated. The residue was dissolved in TFA/$CH_2Cl_2$ (1:4, 0.6 mL) and stirred at rt for 2 h. After the solvent was evaporated, the crude product was dissolved in DMSO (0.8 mL) and purified by preparative HPLC (8 mL/min, 30 % to 90 % MeCN/$H_2O$ (0.1% TFA) in 55 min) to obtain **9** (2.73 mg, 4.03 µmol, 60 %) as a red solid.

$^1$H NMR (400 MHz, MeOD-*d*$_4$) δ 8.22 (d, *J* = 8.1 Hz, 1H), 7.98 (d, *J* = 8.0 Hz, 1H), 7.66 (s, 1H), 7.56 (t, *J* = 7.4 Hz, 1H), 7.46 (d, *J* = 7.9 Hz, 2H), 7.35 (t, *J* = 7.8 Hz, 2H), 6.68 (s, 2H), 6.37 (s, 4H), 3.95 (q, *J* = 9.2 Hz, 4H).

$^{13}$C NMR (101 MHz, DMSO-*d*$_6$) δ 165.9, 164.6, 152.6, 149.4, 138.5, 137.2, 134.1, 130.8, 130.2, 130.0, 128.6, 128.5, 127.7, 127.2, 124.8, 124.4, 124.3, 121.6, 109.5, 107.0, 98.3, 68.9, 44.2, 43.9, 43.6, 43.3.

HRMS (ESI) Exact mass calculated for $C_{31}H_{21}F_6N_3O_6S$ [M+H]$^+$: 678.1128, found: 678.1123.

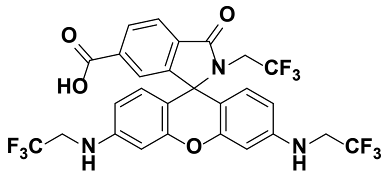


**3-Oxo-2-(2,2,2-trifluoroethyl)-3',6'-bis((2,2,2-trifluoroethyl)-amino)spiro[isoindoline-1,9'-xanthene]-6-carboxylic acid (38):** Following procedure A with 2,2,2-trifluoroethylamine (5.38 µL, 67.3 µmol, 10 eq) at 50°C, **38** (2.08 mg, 3.36 µmol, 50 %) was obtained as a slightly red solid.

$^1$H NMR (400 MHz, DMSO-*d*$_6$) δ 8.10 (dd, *J* = 7.9, 1.1 Hz, 1H), 7.99 (d, *J* = 7.9 Hz, 1H), 7.46 (s, 1H), 6.64 (t, *J* = 6.1 Hz, 2H), 6.58 (d, *J* = 2.1 Hz, 2H), 6.44 (dd, *J* = 8.7, 2.2 Hz, 2H), 6.31 (d, *J* = 8.7 Hz, 2H), 4.00 – 3.92 (m, 4H), 3.75 (q, *J* = 9.6 Hz, 2H).

$^{13}$C NMR (101 MHz, DMSO-*d*$_6$) δ 167.2, 166.2, 153.7, 152.3, 149.2, 135.6, 132.1, 129.9, 129.8, 128.3, 128.0, 127.1, 125.2, 124.4, 124.3, 123.5, 122.4, 121.6, 110.4, 105.3, 98.1, 64.9, 44.3, 43.9, 43.6, 43.3, 41.8, 41.5, 41.1, 40.8.

HRMS (ESI) Exact mass calculated for $C_{27}H_{18}F_9N_3O_4$ [M+H]$^+$: 620.1226, found: 620.1223.

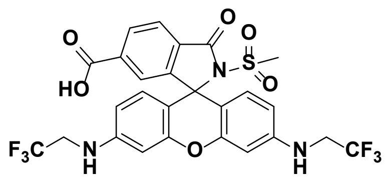

**2-(Methylsulfonyl)-3-oxo-3',6'-bis((2,2,2-trifluoroethyl)amino)spiro[isoindoline-1,9'-xanthene]-6-carboxylic acid (35):** Following procedure A with methanesulfonamide (3.20 mg, 33.7 µmol, 5 eq), **35** (3.56 mg, 5.78 µmol, 86 %) was obtained as a red solid.

$^{1}$H NMR (400 MHz, MeOD-*d₄*) δ 8.25 (d, *J* = 7.8 Hz, 1H), 8.07 (d, *J* = 8.0 Hz, 1H), 7.68 (s, 1H), 6.64 – 6.60 (m, 4H), 6.50 (d, *J* = 7.5 Hz, 2H), 3.90 (q, *J* = 9.1 Hz, 4H), 2.95 (s, 3H).

$^{13}$C NMR (101 MHz, DMSO-$d_6$) δ 165.9, 165.6, 153.3, 152.0, 149.2, 137.1, 130.4, 130.0, 129.9, 128.1, 127.1, 124.6, 124.5, 124.4, 121.6, 109.6, 106.8, 98.2, 68.0, 44.3, 44.0, 43.6, 43.3, 42.0.

HRMS (ESI) Exact mass calculated for $C_{26}H_{19}F_6N_3O_6S$ [M+H]$^+$: 616.0972, found: 616.0969.

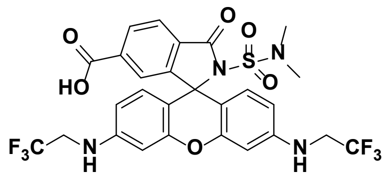

**2-(*N,N*-Dimethylsulfamoyl)-3-oxo-3',6'-bis((2,2,2-trifluoroethyl)-amino)spiro-[isoindoline-1,9'-xanthene]-6-carboxylic acid (36):** Following procedure A with *N,N*-dimethylsulfamide (4.18 mg, 33.7 µmol, 5 eq), **36** (3.07 mg, 4.76 µmol, 71 %) was obtained as a red solid.

$^{1}$H NMR (400 MHz, DMSO-$d_6$) δ 8.12 (d, *J* = 8.0 Hz, 1H), 8.02 (d, *J* = 8.0 Hz, 1H), 7.40 (s, 1H), 6.61 – 6.58 (m, 2H), 6.53 (d, *J* = 1.9 Hz, 2H), 6.48 (d, *J* = 8.7 Hz, 2H), 6.40 (dd, *J* = 8.7, 2.1 Hz, 2H), 3.98 – 3.93 (m, 4H), 2.62 (s, 6H).

$^{13}$C NMR (101 MHz, DMSO-$d_6$) δ 166.5, 166.0, 153.9, 152.9, 149.5, 137.2, 131.3, 130.4, 129.1, 127.6, 125.3, 124.8, 124.6, 109.8, 107.7, 98.5, 68.4, 44.8, 44.4, 44.1, 43.8, 37.9.

HRMS (ESI) Exact mass calculated for $C_{27}H_{22}F_6N_4O_6S$ [M+H]$^+$: 645.1237, found: 645.1239.

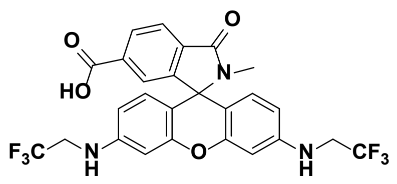

**2-Methyl-3-oxo-3',6'-bis((2,2,2-trifluoroethyl)amino)spiro[isoindoline-1,9'-xanthene]-6-carboxylic acid (40):** Following procedure A with **2** (8.00 mg, 13.5 µmol, 1.0 eq) and methylamine hydrochloride (4.54 mg, 67.3 µmol, 5 eq) **40** (4.03 mg, 7.31 µmol, 54 %) was obtained as a colorless solid.

$^{1}$H NMR (400 MHz, DMSO-$d_6$) δ 8.03 (dd, J = 7.9, 1.3 Hz, 1H), 7.88 (d, J = 7.9 Hz, 1H), 7.42 – 7.42 (m, 1H), 6.64 – 6.60 (m, 4H), 6.45 (dd, J = 8.7, 2.3 Hz, 2H), 6.34 (d, J = 8.6 Hz, 2H), 4.00 – 3.91 (m, 4H), 2.53 (s, 3H).

$^{13}$C NMR (101 MHz, DMSO-$d_6$) δ 166.4, 165.8, 153.5, 152.3, 149.1, 149.0, 134.8, 133.4, 129.9, 129.5, 127.3, 127.2, 124.4, 123.7, 122.9, 121.6, 110.2, 106.1, 106.1, 98.5, 63.7, 44.3, 44.0, 43.7, 43.4, 24.5.

HRMS (ESI) Exact mass calculated for $C_{26}H_{19}F_6N_3O_4$ [M+H]+: 552.1353, found: 552.1350.

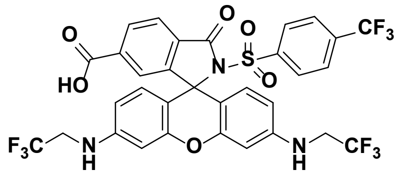


**3-Oxo-3',6'-bis((2,2,2-trifluoroethyl)amino)-2-((4-(trifluoromethyl)phenyl)sulfonyl)-spiro[isoindoline-1,9'-xanthene]-6-carboxylic acid (6):** Following procedure A with 4-(trifluoromethyl)benzenesulfonamide (7.6 mg, 33.7 µmol, 5 eq), **6** (3.51 mg, 4.71 µmol, 70 %) was obtained as a red solid.

$^1$H NMR (400 MHz, MeOD-*d4*) δ 8.22 (dd, *J* = 8.0, 1.1 Hz, 1H), 8.00 (d, *J* = 8.0 Hz, 1H), 7.66 (d, *J* = 8.8 Hz, 3H), 7.61 (d, *J* = 8.5 Hz, 2H), 6.67 (s, 2H), 6.37 – 6.32 (m, 4H), 3.93 (q, *J* = 9.3 Hz, 4H).

$^{13}$C NMR (101 MHz, DMSO-*d6*) δ 165.9, 164.8, 152.6, 152.4, 149.5, 142.1, 137.4, 133.6, 133.3, 130.6, 130.2, 130.0, 128.7, 128.5, 127.2, 125.8, 125.8, 124.8, 124.6, 124.5, 124.4, 121.8, 121.6, 119.1, 109.5, 106.8, 98.2, 69.5, 44.2, 43.9, 43.6, 43.3.

HRMS (ESI) Exact mass calculated for $C_{32}H_{20}F_9N_3O_6S$ [M+H]+: 746.1002, found: 746.0996.

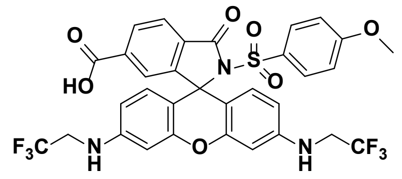


**2-((4-Methoxyphenyl)sulfonyl)-3-oxo-3',6'-bis((2,2,2-trifluoroethyl)amino)spiro-[isoindoline-1,9'-xanthene]-6-carboxylic acid (11):** Following procedure A with 4-(methoxy)benzenesulfonamide (6.3 mg, 33.7 µmol, 5 eq), **11** (3.49 mg, 4.93 µmol, 73 %) was obtained as a red solid.

$^1$H NMR (400 MHz, MeOD-*d4*) δ 8.21 (d, *J* = 7.9 Hz, 1H), 7.97 (d, *J* = 8.0 Hz, 1H), 7.65 (s, 1H), 7.38 (d, *J* = 8.6 Hz, 2H), 6.83 (d, *J* = 8.9 Hz, 2H), 6.66 (s, 2H), 6.37 (s, 4H), 3.96 – 3.91 (m, 4H), 3.83 (s, 3H).

$^{13}$C NMR (101 MHz, DMSO-*d6*) δ 166.0, 164.6, 163.4, 152.6, 152.5, 149.3, 137.0, 130.9, 130.2, 130.1, 130.1, 130.0, 128.5, 127.2, 124.7, 124.4, 124.2, 121.6, 113.6, 109.5, 107.1, 98.2, 68.5, 55.7, 44.2, 43.9, 43.6, 43.3.

HRMS (ESI) Exact mass calculated for $C_{32}H_{23}F_6N_3O_7S$ [M+H]+: 708.1234, found: 708.1231.

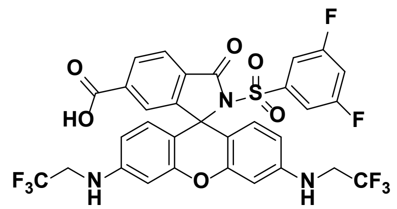


**2-((3,5-Difluorophenyl)sulfonyl)-3-oxo-3',6'-bis((2,2,2-trifluoroethyl)amino)spiro-[isoindoline-1,9'-xanthene]-6-carboxylic acid (5):** Following procedure A with 3,5-difluorobenzenesulfonamide (6.5 mg, 33.7 µmol, 5 eq), **5** (2.89 mg, 4.05 µmol, 60 %) was obtained as a red solid.

$^{1}H$ NMR (400 MHz, MeOD-$d_4$) δ 8.23 (d, *J* = 8.1 Hz, 1H), 8.02 (d, *J* = 8.0 Hz, 1H), 7.66 (s, 1H), 7.22 (tt, *J* = 8.7, 2.2 Hz, 1H), 7.00 – 6.96 (m, 2H), 6.67 (d, *J* = 1.5 Hz, 2H), 6.39 – 6.33 (m, 4H), 3.91 (q, *J* = 9.2 Hz, 4H).

$^{13}C$ NMR (101 MHz, MeOD-$d_4$) δ 166.4, 165.5, 163.5, 163.4, 161.0, 160.9, 153.9, 150.6, 142.1, 142.0, 141.9, 137.2, 131.8, 130.3, 128.7, 126.8, 126.1, 124.2, 124.0, 111.3, 111.2, 111.1, 111.0, 109.9, 109.1, 108.9, 108.6, 108.1, 98.3, 44.9, 44.6, 44.2, 43.9

HRMS (ESI) Exact mass calculated for $C_{31}H_{19}F_8N_3O_6S$ $[M+H]^+$: 714.0940, found: 714.0938.

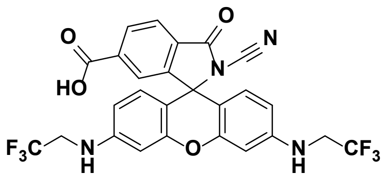


**2-Cyano-3-oxo-3',6'-bis((2,2,2-trifluoroethyl)amino) spiro[iso-indoline-1,9'-xanthene]-6-carboxylic acid (34):** Following procedure A with **2** (2.00 mg, 3.37 µmol, 1.0 eq) and cyanamide (1.42 mg, 33.7 µmol, 10 eq), **34** (0.850 mg, 1.51 µmol, 45 %) was obtained as a red solid.

$^{1}H$ NMR (400 MHz, MeOD-$d_4$) δ 8.28 (d, *J* = 8.1 Hz, 1H), 8.11 (d, *J* = 8.0 Hz, 1H), 7.71 (s, 1H), 6.61 (d, *J* = 2.3 Hz, 2H), 6.58 (d, *J* = 8.7 Hz, 2H), 6.51 (dd, *J* = 8.8, 2.4 Hz, 2H), 3.88 (q, *J* = 9.3 Hz, 4H).

HRMS (ESI) Exact mass calculated for $C_{26}H_{16}F_6N_4O_4$ $[M+H]^+$: 563.1149, found: 563.1148.

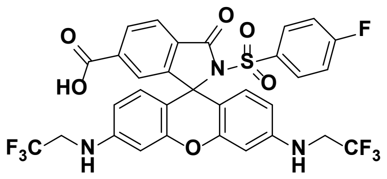


**2-((4-Fluorophenyl)sulfonyl)-3-oxo-3',6'-bis((2,2,2-trifluoroethyl)amino)spiro[iso-indoline-1,9'-xanthene]-6-carboxylic acid (8):** Following procedure A with 4-fluorobenzenesulfonamide (5.9 mg, 33.7 µmol, 5 eq), **8** (3.63 mg, 5.22 µmol, 78 %) was obtained as a red solid.

$^{1}H$ NMR (400 MHz, MeOD-$d_4$) δ 8.22 (dd, *J* = 8.0, 1.3 Hz, 1H), 7.99 (d, *J* = 8.1 Hz, 1H), 7.66 (s, 1H), 7.51 – 7.48 (m, 2H), 7.10 – 7.05 (m, 2H), 6.73 – 6.61 (m, 2H), 6.39 – 6.34 (m, 4H), 3.94 (q, *J* = 9.2 Hz, 4H).

$^{13}C$ NMR (101 MHz, DMSO-$d_6$) δ 166.3, 165.9, 164.7, 163.7, 152.5, 149.5, 137.2, 134.8, 134.8, 130.9, 130.8, 130.7, 130.2, 130.0, 128.6, 127.2, 124.8, 124.4, 124.4, 121.6, 115.9, 115.7, 109.5, 106.9, 98.3, 69.0, 44.3, 43.9, 43.6, 43.3.

HRMS (ESI) Exact mass calculated for $C_{31}H_{20}F_7N_3O_6S$ $[M+H]^+$: 696.1034, found: 696.1032.

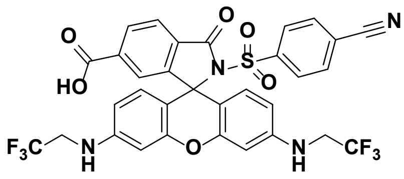

**2-((4-Cyanophenyl)sulfonyl)-3-oxo-3',6'-bis((2,2,2-trifluoroethyl)amino)spiro[iso-indoline-1,9'-xanthene]-6-carboxylic acid (4):** Following procedure A with 4-cyanobenzenesulfonamide (6.15 mg, 33.7 µmol, 5 eq), **4** (3.93 mg, 5.59 µmol, 83 %) was obtained as a red solid.

$^1$H NMR (400 MHz, DMSO-*d$_6$*) δ 8.10 (dd, *J* = 8.1, 1.4 Hz, 1H), 7.98 (d, *J* = 8.0 Hz, 1H), 7.89 – 7.85 (m, 2H), 7.45 – 7.40 (m, 3H), 6.72 (t, *J* = 6.9 Hz, 2H), 6.65 (d, *J* = 2.2 Hz, 2H), 6.34 (dd, *J* = 8.7, 2.3 Hz, 2H), 6.27 (d, *J* = 8.7 Hz, 2H), 4.06 – 3.97 (m, 4H).

$^{13}$C NMR (101 MHz, DMSO-*d$_6$*) δ 165.9, 164.9, 152.5, 152.3, 149.6, 149.5, 142.0, 137.6, 132.7, 130.4, 130.2, 130.0, 128.7, 128.2, 127.2, 124.8, 124.5, 124.4, 121.6, 117.4, 116.3, 109.6, 106.8, 98.2, 69.6, 44.3, 43.9, 43.6, 43.3, 40.2, 39.9, 39.7, 39.5, 39.3, 39.1, 38.9.

HRMS (ESI) Exact mass calculated for $C_{32}H_{20}F_6N_4O_6S$ [M+H]$^+$: 703.1081, found: 703.1077.

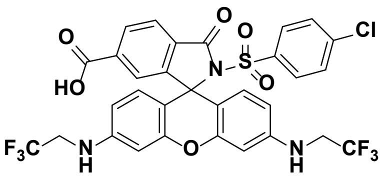

**2-((4-Chlorophenyl)sulfonyl)-3-oxo-3',6'-bis((2,2,2-trifluoro-ethyl)amino)spiro[iso-indoline-1,9'-xanthene]-6-carboxylic acid (7):** Following procedure A with 4-chlorobenzenesulfonamide (6.45 mg, 33.7 µmol, 5 eq), **7** (2.89 mg, 4.06 µmol, 60 %) was obtained as a red solid.

$^1$H NMR (400 MHz, MeOD-*d$_4$*) δ 8.21 (dd, *J* = 8.0, 1.3 Hz, 1H), 7.99 (d, *J* = 8.1 Hz, 1H), 7.64 (s, 1H), 7.41 – 7.34 (m, 4H), 6.66 (d, *J* = 1.7 Hz, 2H), 6.38 – 6.32 (m, 4H), 3.93 (q, *J* = 9.3 Hz, 4H).

$^{13}$C NMR (101 MHz, DMSO-*d$_6$*) δ 165.9, 164.7, 152.5, 152.5, 149.5, 139.2, 137.3, 137.2, 130.6, 130.2, 130.0, 129.6, 128.7, 128.6, 127.2, 124.8, 124.4, 121.6, 109.5, 106.9, 98.2, 69.1, 44.2, 43.9, 43.6, 43.3.

HRMS (ESI) Exact mass calculated for $C_{31}H_{20}ClF_6N_3O_6S$ [M+H]$^+$: 712.0738, found: 712.0734.

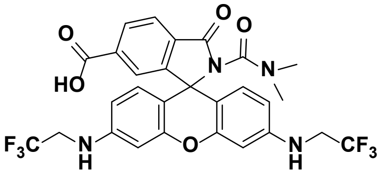

**2-(Dimethylcarbamoyl)-3-oxo-3',6'-bis((2,2,2-trifluoroethyl) amino)spiro[iso-indoline-1,9'-xanthene]-6-carboxylic acid (37):** Following procedure A with 1,1-dimethylurea (5.93 mg, 67.3 µmol, 10 eq), **37** (0.820 mg, 1.35 µmol, 20 %) was obtained as a colorless solid.

$^1$H NMR (400 MHz, MeOD-*d$_4$*) δ 8.24 (d, *J* = 8.0 Hz, 1H), 8.03 (d, *J* = 8.0 Hz, 1H), 7.68 (s, 1H), 6.51 (d, *J* = 2.1 Hz, 2H), 6.47 (d, *J* = 8.6 Hz, 2H), 6.39 (dd, *J* = 8.7, 2.3 Hz, 2H), 3.87 – 3.80 (m, 4H), 2.80 (s, 6H).

HRMS (ESI) Exact mass calculated for $C_{28}H_{22}F_6N_4O_5$ [M+H]$^+$: 609.1567, found: 609.1572.

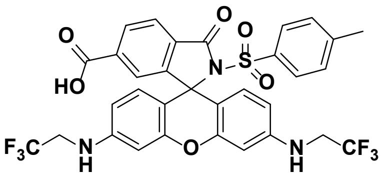

**3-Oxo-2-tosyl-3',6'-bis((2,2,2-trifluoroethyl)amino) spiro[isoindoline-1,9'-xanthene]-6-carboxylic acid (10):** Following procedure A with *p*-toluenesulfonamide (5.75 mg, 33.7 µmol, 5 eq), **10** (2.55 mg, 3.69 µmol, 55 %) was obtained as a red solid.

$^1$H NMR (400 MHz, MeOD-*d₄*) δ 8.20 (dd, *J* = 8.1, 1.1 Hz, 1H), 7.97 (d, *J* = 8.0 Hz, 1H), 7.63 (s, 1H), 7.32 (d, *J* = 8.3 Hz, 2H), 7.15 (d, *J* = 8.1 Hz, 2H), 6.65 (s, 2H), 6.38 – 6.32 (m, 4H), 3.92 (q, *J* = 9.2 Hz, 4H), 2.36 (s, 3H).

$^{13}$C NMR (101 MHz, DMSO-$d_6$) δ 166.0, 164.6, 152.6, 152.5, 149.4, 144.8, 137.1, 135.7, 130.8, 130.1, 130.0, 128.9, 128.6, 127.9, 127.2, 124.8, 124.4, 124.2, 121.6, 109.4, 107.1, 98.3, 68.7, 44.3, 43.9, 43.6, 43.3, 21.1.

HRMS (ESI) Exact mass calculated for $C_{32}H_{23}F_6N_3O_6S$, $[M+H]^+$: 692.1285, found: 692.1284.

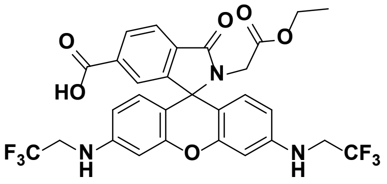

**2-(2-Ethoxy-2-oxoethyl)-3-oxo-3',6'-bis((2,2,2-trifluoroethyl) amino)spiro[iso-indoline-1,9'-xanthene]-6-carboxylic acid (39):** Following procedure A with glycine ethyl ester hydrochloride (4.70 mg, 33.7 µmol, 5 eq), **39** (1.04 mg, 1.67 µmol, 25 %) was obtained as a colorless solid.

$^1$H NMR (400 MHz, MeOD-$d_4$) δ 8.21 (d, *J* = 7.9 Hz, 1H), 7.99 (d, *J* = 8.0 Hz, 1H), 7.66 (s, 1H), 6.54 (d, *J* = 1.6 Hz, 2H), 6.43 – 6.37 (m, 4H), 3.89 – 3.80 (m, 6H), 1.08 (t, *J* = 7.1 Hz, 3H).

HRMS (ESI) Exact mass calculated for $C_{29}H_{23}F_6N_3O_6$ $[M+H]^+$: 624.1564, found: 624.1562.

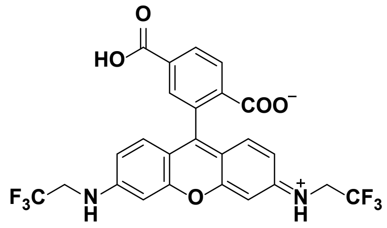

**3-Oxo-3',6'-bis((2,2,2-trifluoroethyl)amino)-3H-spiro[isobenzofuran-1,9'-xanthene]-6-carboxylic acid (1): 2** (4.00 mg, 6.73 µmol, 1.0 eq) was dissolved in TFA/$CH_2Cl_2$ (1:4, 0.6 mL) and stirred at rt for 2 h. After the solvent was evaporated, the crude product was dissolved in DMSO (0.8 mL) and purified by preparative HPLC (8 mL/min, 30 % to 90 % MeCN/$H_2O$ (0.1 % TFA) in 55 min) to obtain **1** (3.26 mg, 6.06 µmol, 90 %) as a red solid.

$^1$H NMR (400 MHz, DMSO-$d_6$) δ 8.21 (d, *J* = 8.0 Hz, 1H), 8.09 (d, *J* = 7.9 Hz, 1H), 7.63 (s, 1H), 6.80 – 6.75 (m, 2H), 6.65 (s, 2H), 6.52 – 6.47 (m, 4H), 4.05 – 3.96 (m, 4H).

$^{13}C$ NMR (101 MHz, DMSO-*d6*) δ 168.5, 166.5, 153.1, 152.6, 150.4, 137.6, 131.2, 130.4, 129.1, 127.6, 125.6, 124.8, 122.0, 110.6, 107.1, 98.4, 84.9, 44.6, 44.3, 43.9, 43.6.

HRMS (ESI) Exact mass calculated for $C_{25}H_{16}F_6N_2O_5$ $[M+H]^+$: 539.1036, found: 539.1036.

**General Procedure B for 15:**

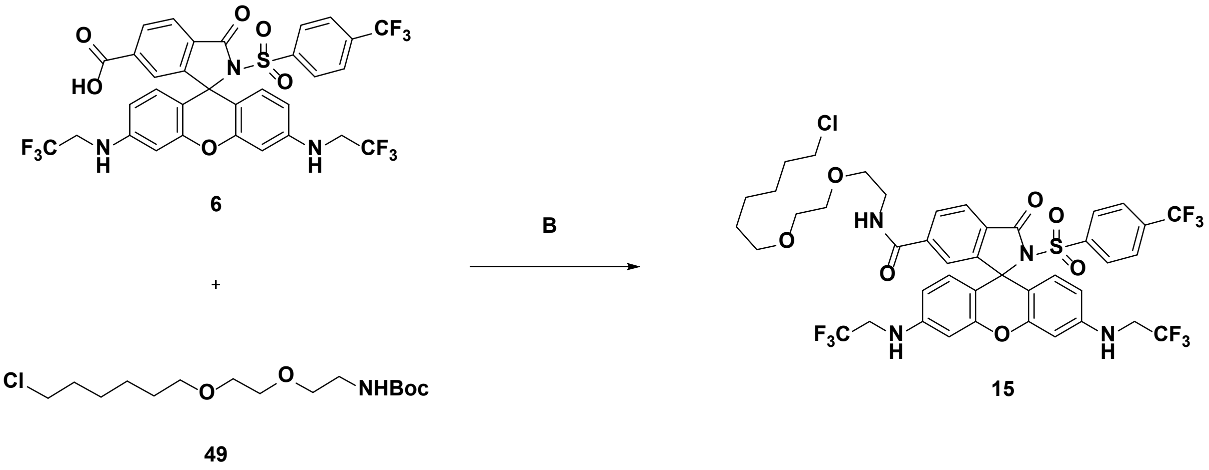


***N*-(2-(2-((6-Chlorohexyl)oxy)ethoxy)ethyl)-3-oxo-3',6'-bis((2,2,2-trifluoroethyl)amino)-2-((4-(trifluoromethyl)phenyl)sulfonyl)spiro[isoindoline-1,9'-xanthene]-6-carboxamide (15):** A solution of Halo-NHBoc (**49**, 1.98 mg, 6.12 µmol, 1.3 eq) in TFA (0.3 mL) was stirred at rt for 5 min. The solvent was evaporated, the residue dissolved in DMSO (0.3 mL) and transferred to a solution of **6** (3.51 mg, 4.71 µmol, 1.0 eq), PyBOP (3.67 mg, 7.06 µmol, 1.5 eq), and DIPEA (7.78 µL, 47.1 µmol, 10 eq) in DMSO (0.3 mL). The reaction mixture was stirred at rt for 40 min followed by purification using preparative HPLC (8 mL/min, 30 % to 90 % MeCN/$H_2O$ (0.1 % TFA) in 55 min) to obtain **15** (2.90 mg, 3.05 µmol, 65 %) as a red solid.

$^1H$ NMR (400 MHz, DMSO-*d6*) δ 8.71 (t, *J* = 5.5 Hz, 1H), 8.07 (d, *J* = 8.1 Hz, 1H), 7.95 (d, *J* = 8.1 Hz, 1H), 7.76 (d, *J* = 8.4 Hz, 2H), 7.55 (s, 1H), 7.48 (d, *J* = 8.3 Hz, 2H), 6.71 (t, *J* = 6.8 Hz, 2H), 6.65 (d, *J* = 1.6 Hz, 2H), 6.33 (dd, *J* = 8.7, 1.8 Hz, 2H), 6.26 (d, *J* = 8.6 Hz, 2H), 4.06 – 3.97 (m, 4H), 3.57 (t, *J* = 6.6 Hz, 2H), 3.45 – 3.38 (m, 6H), 3.34 – 3.26 (m, 4H), 1.64 (p, J = 6.7 Hz, 2H), 1.43 – 1.36 (m, 2H), 1.35 – 1.28 (m, 2H), 1.25 – 1.20 (m, 2H).

HRMS (ESI) Exact mass calculated for $C_{42}H_{40}ClF_9N_4O_7S$ $[M+H]^+$: 951.2235, found: 951.2233.

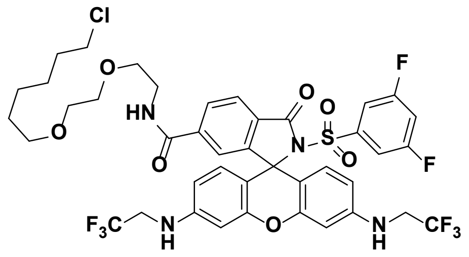

***N*-(2-(2-((6-Chlorohexyl)oxy)ethoxy)ethyl)-2-((3,5-difluorophenyl)sulfonyl)-3-oxo-3',6'-bis((2,2,2-trifluoroethyl)amino)spiro [isoindoline-1,9'-xanthene]-6-carboxamide (14):** Following procedure B with **5** (2.89 mg, 4.05 µmol, 1.0 eq), **14** (1.96 mg, 2.13 µmol, 53 %) was obtained as a red solid.

$^1$H NMR (400 MHz, DMSO-$d_6$) δ 8.72 (t, *J* = 5.3 Hz, 1H), 8.08 (d, *J* = 8.1 Hz, 1H), 7.97 (d, *J* = 8.0 Hz, 1H), 7.64 (t, *J* = 8.8 Hz, 1H), 7.56 (s, 1H), 6.83 (d, *J* = 4.3 Hz, 2H), 6.72 (t, *J* = 6.0 Hz, 2H), 6.64 (s, 2H), 6.34 (d, *J* = 8.8 Hz, 2H), 6.27 (d, *J* = 8.6 Hz, 2H), 4.04 – 3.95 (m, 4H), 3.58 (t, *J* = 6.6 Hz, 2H), 3.45 – 3.39 (m, 6H), 3.33 – 3.27 (m, 4H), 1.64 (p, *J* = 6.8 Hz, 2H), 1.43 – 1.36 (m, 2H), 1.35 – 1.28 (m, 2H), 1.25 – 1.20 (m, 2H).

HRMS (ESI) Exact mass calculated for $C_{41}H_{39}ClF_8N_4O_7S$ $[M+H]^+$: 919.2173, found: 919.2175.

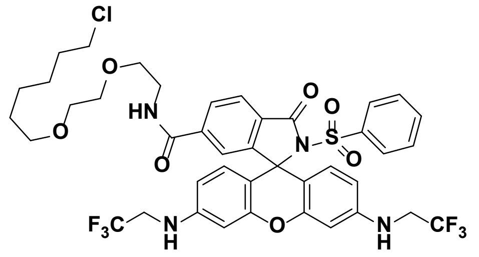

***N*-(2-(2-((6-Chlorohexyl)oxy)ethoxy)ethyl)-3-oxo-2-(phenylsulfonyl)-3',6'-bis((2,2,2-trifluoroethyl) amino)spiro[isoindoline-1,9'-xanthene]-6-carboxamide (18):** Following procedure B with **9** (2.73 mg, 4.03 µmol, 1.0 eq), **18** (1.70 mg, 1.92 µmol, 48 %) was obtained as a red solid.

$^1$H NMR (400 MHz, DMSO-$d_6$) δ 8.70 (t, *J* = 5.5 Hz, 1H), 8.05 (d, *J* = 8.1 Hz, 1H), 7.92 (d, *J* = 8.1 Hz, 1H), 7.64 (t, *J* = 7.4 Hz, 1H), 7.54 (s, 1H), 7.39 (t, *J* = 7.8 Hz, 2H), 7.29 (d, *J* = 7.8 Hz, 2H), 6.68 (t, *J* = 6.7 Hz, 2H), 6.64 (d, *J* = 1.7 Hz, 2H), 6.35 (dd, *J* = 9.0, 1.6 Hz, 2H), 6.23 (d, *J* = 8.6 Hz, 2H), 4.05 – 3.97 (m, 4H), 3.57 (t, *J* = 6.6 Hz, 2H), 3.45 – 3.39 (m, 6H), 3.31 – 3.27 (m, 4H), 1.64 (p, *J* = 6.8 Hz, 2H), 1.43 – 1.36 (m, 2H), 1.35 – 1.28 (m, 2H), 1.25 – 1.20 (m, 2H).

HRMS (ESI) Exact mass calculated for $C_{41}H_{41}ClF_6N_4O_7S$ $[M+H]^+$: 883.2361, found: 883.2362.

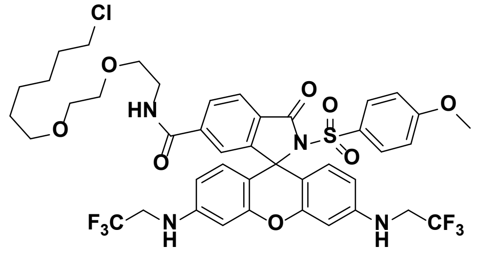

***N*-(2-(2-((6-Chlorohexyl)oxy)ethoxy)ethyl)-2-((4-methoxyphenyl)sulfonyl)-3-oxo-3',6'-bis((2,2,2-trifluoroethyl)amino)spiro [isoindoline-1,9'-xanthene]-6-carboxamide (20):** Following procedure B with **11** (3.49 mg, 4.93 µmol, 1.0 eq), **20** (3.03 mg, 3.32 µmol, 67 %) was obtained as a red solid.

$^1$H NMR (400 MHz, DMSO-$d_6$) δ 8.70 (t, *J* = 5.3 Hz, 1H), 8.05 (d, *J* = 8.0 Hz, 1H), 7.91 (d, *J* = 8.0 Hz, 1H), 7.53 (s, 1H), 7.22 (d, *J* = 8.8 Hz, 2H), 6.87 (d, *J* = 8.9 Hz, 2H), 6.67 (t, *J* = 6.2 Hz, 2H), 6.63 (s, 2H), 6.35

(d, $J$ = 8.7 Hz, 2H), 6.22 (d, $J$ = 8.6 Hz, 2H), 4.05 – 3.97 (m, 4H), 3.80 (s, 3H), 3.57 (t, $J$ = 6.6 Hz, 2H), 3.45 – 3.39 (m, 6H), 3.32 – 3.27 (m, 4H), 1.65 (p, $J$ = 6.8 Hz, 2H), 1.43 – 1.36 (m, 2H), 1.34 – 1.28 (m, 2H), 1.26 – 1.20 (m, 2H).

HRMS (ESI) Exact mass calculated for $C_{42}H_{43}ClF_6N_4O_8S$ $[M+H]^+$: 913.2467, found: 913.2462.

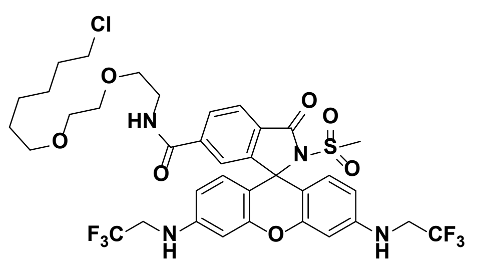


**_N_-(2-(2-((6-Chlorohexyl)oxy)ethoxy)ethyl)-2-(methylsulfonyl)-3-oxo-3',6'-bis((2,2,2-trifluoroethyl)amino)spiro[isoindoline-1,9'-xanthene]-6-carboxamide (42):** Following procedure B with **35** (3.56 mg, 5.78 µmol, 1.0 eq), **42** (3.32 mg, 4.04 µmol, 70 %) was obtained as a red solid.

$^1$H NMR (400 MHz, DMSO-$d_6$) δ 8.72 (t, $J$ = 5.5 Hz, 1H), 8.08 (d, $J$ = 8.1 Hz, 1H), 8.02 (d, $J$ = 8.0 Hz, 1H), 7.50 (s, 1H), 6.62 (t, $J$ = 6.6 Hz, 2H), 6.53 – 6.51 (m, 4H), 6.42 (dd, J = 8.7, 1.9 Hz, 2H), 4.01 – 3.91 (m, 4H), 3.59 (t, $J$ = 6.6 Hz, 2H), 3.46 – 3.39 (m, 6H), 3.34 – 3.27 (m, 4H), 2.97 (s, 3H), 1.66 (p, $J$ = 6.8 Hz, 2H), 1.44 – 1.37 (m, 2H), 1.35 – 1.29 (m, 2H), 1.27 – 1.20 (m, 2H).

HRMS (ESI) Exact mass calculated for $C_{36}H_{39}ClF_6N_4O_7S$ $[M+H]^+$: 821.2205, found: 821.2202.

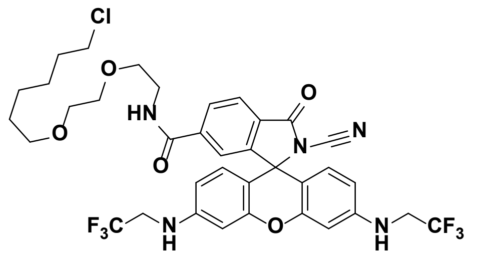


**_N_-(2-(2-((6-Chlorohexyl)oxy)ethoxy)ethyl)-2-cyano-3-oxo-3',6'-bis((2,2,2-trifluoroethyl)amino)spiro[isoindoline-1,9'-xanthene]-6-carboxamide (41):** Following procedure B with **34** (0.930 mg, 1.65 µmol, 1.0 eq), **41** (1.03 mg, 1.34 µmol, 81 %) was obtained as a red solid.

$^1$H NMR (400 MHz, DMSO-$d_6$) δ 8.77 (t, $J$ = 5.5 Hz, 1H), 8.14 (d, $J$ = 8.6 Hz, 1H), 8.09 (d, $J$ = 7.9 Hz, 1H), 7.63 (s, 1H), 6.81 (t, $J$ = 6.7 Hz, 2H), 6.63 – 6.60 (m, 4H), 6.52 (d, $J$ = 8.6 Hz, 2H), 4.06 – 3.96 (m, 4H), 3.59 (t, $J$ = 6.6 Hz, 2H), 3.47 – 3.39 (m, 6H), 3.32 – 3.27 (m, 4H) 1.65 (p, $J$ = 6.8 Hz, 2H), 1.44 – 1.37 (m, 2H), 1.34 – 1.29 (m, 2H), 1.26 – 1.19 (m, 2H).

HRMS (ESI) Exact mass calculated for $C_{36}H_{36}ClF_6N_5O_5$ $[M+H]^+$: 768.2382, found: 768.2382.

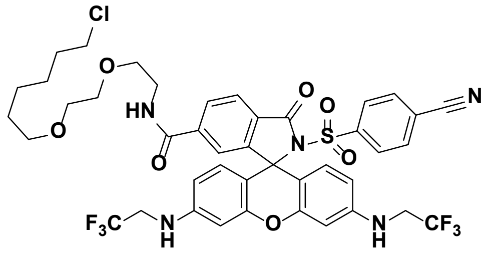


***N*-(2-(2-((6-Chlorohexyl)oxy)ethoxy)ethyl)-2-((4-cyanophenyl)sulfonyl)-3-oxo-3',6'-bis((2,2,2-trifluoroethyl)-amino)spiro [isoindoline-1,9'-xanthene]-6-carboxamide (13):** Following procedure B with **4** (3.93 mg, 5.59 µmol, 1.0 eq), **13** (1.90 mg, 2.09 µmol, 37 %) was obtained as a red solid.

$^1$H NMR (400 MHz, DMSO-*d$_6$*) δ 8.71 (t, *J* = 5.4 Hz, 1H), 8.07 (d, *J* = 8.1 Hz, 1H), 7.95 (d, *J* = 8.0 Hz, 1H), 7.86 (d, *J* = 8.3 Hz, 2H), 7.55 (s, 1H), 7.41 (d, *J* = 8.3 Hz, 2H), 6.71 (t, *J* = 6.7 Hz, 2H), 6.64 (s, 2H), 6.33 (d, *J* = 8.7 Hz, 2H), 6.24 (d, *J* = 8.6 Hz, 2H), 4.06 – 3.97 (m, 4H), 3.57 (t, *J* = 6.6 Hz, 2H), 3.45 – 3.38 (m, 6H), 3.30 – 3.26 (m, 4H), 1.64 (p, *J* = 6.8 Hz, 2H), 1.43 – 1.36 (m, 2H), 1.33 – 1.28 (m, 2H), 1.25 – 1.20 (m, 2H).

HRMS (ESI) Exact mass calculated for $C_{42}H_{40}ClF_6N_5O_7S$ [M+H]$^+$: 908.2314, found: 908.2314.

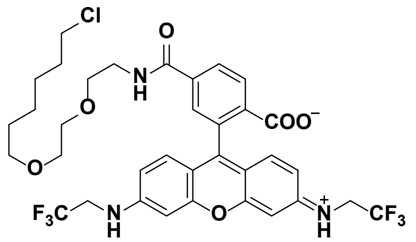


**(*E*)-4-((2-(2-((6-Chlorohexyl)oxy)ethoxy) ethyl)carbamoyl)-2-(6-((2,2,2-trifluoroethyl)-amino)-3-((2,2,2-trifluoroethyl)iminio)-*3H*-xanthen-9-yl)benzoate (12):** Following procedure B with **1** (3.26 mg, 6.06 µmol, 1.0 eq), **12** (3.51 mg, 4.72 µmol, 78 %) was obtained as a red solid.

$^1$H NMR (400 MHz, MeOD-*d$_4$*) δ 8.33 (d, *J* = 8.2 Hz, 1H), 8.20 (dd, *J* = 8.2, 1.6 Hz, 1H), 7.79 (d, *J* = 1.4 Hz, 1H), 7.14 (d, *J* = 9.2 Hz, 2H), 7.08 (d, *J* = 2.0 Hz, 2H), 6.95 (dd, *J* = 9.2, 2.2 Hz, 2H), 4.19 (q, *J* = 9.0 Hz, 4H), 3.66 – 3.54 (m, 8H), 3.52 (t, *J* = 6.6 Hz, 2H), 3.42 (t, *J* = 6.5 Hz, 2H), 1.71 (p, *J* = 6.7 Hz, 2H), 1.49 (p, *J* = 6.7 Hz, 2H), 1.43 – 1.36 (m, 2H), 1.34 – 1.28 (m, 2H).

HRMS (ESI) Exact mass calculated for $C_{35}H_{36}ClF_6N_3O_6$ [M+H]$^+$: 744.2270, found: 744.2268.

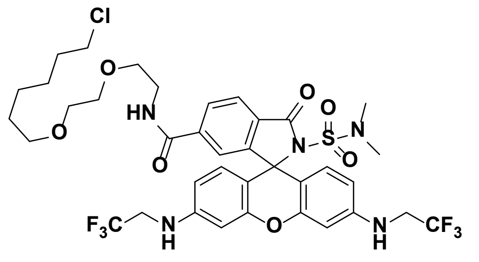


***N*-(2-(2-((6-Chlorohexyl)oxy)ethoxy)ethyl)-2-(*N,N*-dimethylsulfamoyl)-3-oxo-3',6'-bis((2,2,2-trifluoroethyl)amino)spiro [isoindoline-1,9'-xanthene]-6-carboxamide (21):** Following procedure B with **36** (3.07 mg, 4.76 µmol, 1.0 eq), **21** (2.11 mg, 2.48 µmol, 52 %) was obtained as a red solid.

$^1$H NMR (400 MHz, DMSO-*d$_6$*) δ 8.70 (t, *J* = 5.4 Hz, 1H), 8.08 (d, *J* = 8.1 Hz, 1H), 7.98 (d, *J* = 8.0 Hz, 1H), 7.50 (s, 1H), 6.58 (t, *J* = 6.4 Hz, 2H), 6.52 (s, 2H), 6.45 (d, *J* = 8.6 Hz, 2H), 6.39 (d, *J* = 8.6 Hz, 2H), 4.01 –

3.90 (m, 4H), 3.59 (t, *J* = 6.6 Hz, 2H), 3.46 – 3.39 (m, 6H), 3.32 – 3.28 (m, 4H), 2.61 (s, 6H), 1.66 (p, *J* = 6.7 Hz, 2H), 1.44 – 1.37 (m, 2H), 1.35 – 1.29 (m, 2H), 1.27 – 1.21 (m, 2H).

HRMS (ESI) Exact mass calculated for $C_{37}H_{42}ClF_6N_5O_7S$ [M+H]$^+$: 850.2470, found: 850.2474.

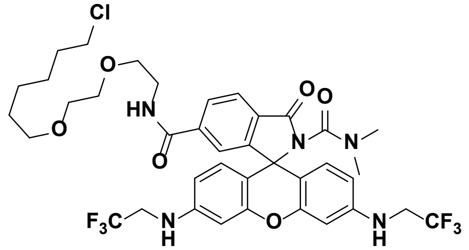


***N*[6]-(2-(2-((6-Chlorohexyl)oxy)ethoxy)ethyl)-*N*[2],*N*[2]-dimethyl-3-oxo-3',6'-bis((2,2,2-trifluoroethyl)amino)spiro[isoindoline-1,9'-xanthene]-2,6-dicarboxamide (25):** Following procedure B with **37** (0.820 mg, 1.35 µmol, 1.0 eq), **25** (0.590 mg, 0.725 µmol, 54 %) was obtained as a colorless solid.

$^1$H NMR (400 MHz, DMSO-$d_6$) δ 8.68 (t, *J* = 5.5 Hz, 1H), 8.06 (d, *J* = 8.0 Hz, 1H), 7.97 (d, *J* = 8.0 Hz, 1H), 7.52 (s, 1H), 6.53 (t, *J* = 6.9 Hz, 2H), 6.49 (d, *J* = 1.8 Hz, 2H), 6.41 (d, *J* = 8.7 Hz, 2H), 6.37 (dd, *J* = 8.7, 2.1 Hz, 2H), 3.98 – 3.89 (m, 4H), 3.58 (t, *J* = 6.6 Hz, 2H), 3.46 – 3.39 (m, 6H), 3.31 – 3.28 (m, 4H), 2.71 (s, 6H), 1.65 (p, *J* = 6.8 Hz, 2H), 1.44 – 1.37 (m, 2H), 1.35 – 1.29 (m, 2H), 1.24 – 1.21 (m, 2H).

HRMS (ESI) Exact mass calculated for $C_{38}H_{42}ClF_6N_5O_6$ [M+H]$^+$: 814.2801, found: 814.2801.

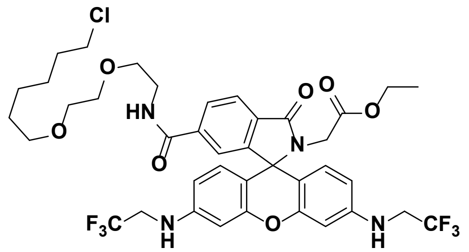


**Ethyl 2-(6-((2-(2-((6-chlorohexyl)oxy)ethoxy) ethyl)-carbamoyl)-3-oxo-3',6'-bis((2,2,2-trifluoroethyl)amino)spiro-[isoindoline-1,9'-xanthen]-2-yl)acetate (27):** Following procedure B with **39** (1.04 mg, 1.67 µmol, 1.0 eq), **27** (1.08 mg, 1.30 µmol, 78 %) was obtained as a colorless solid.

$^1$H NMR (400 MHz, DMSO-$d_6$) δ 8.65 (t, *J* = 5.5 Hz, 1H), 8.03 (d, *J* = 8.0 Hz, 1H), 7.91 (d, *J* = 7.9 Hz, 1H), 7.53 (s, 1H), 6.60 (t, *J* = 6.8 Hz, 2H), 6.55 (d, *J* = 1.7 Hz, 2H), 6.41 (dd, *J* = 8.7, 1.8 Hz, 2H), 6.33 (d, *J* = 8.7 Hz, 2H), 3.99 – 3.90 (m, 4H), 3.76 (q, *J* = 7.1 Hz, 2H), 3.68 (s, 2H), 3.58 (t, *J* = 6.6 Hz, 2H), 3.47 – 3.39 (m, 6H), 3.31 – 3.28 (m, 4H), 1.66 (p, *J* = 6.8 Hz, 2H), 1.44 – 1.37 (m, 2H), 1.34 – 1.29 (m, 2H), 1.26 – 1.21 (m, 2H), 0.99 (t, *J* = 7.1 Hz, 3H).

HRMS (ESI) Exact mass calculated for $C_{39}H_{43}ClF_6N_4O_7$ [M+H]$^+$: 829.2797, found: 829.2799.

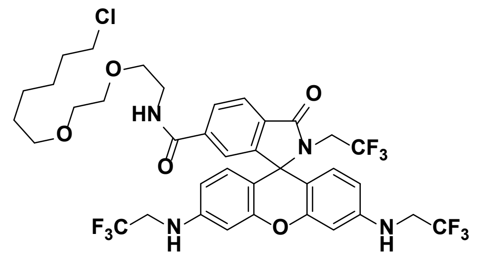

***N*-(2-(2-((6-Chlorohexyl)oxy)ethoxy)ethyl)-3-oxo-2-(2,2,2-trifluoroethyl)-3',6'-bis((2,2,2-trifluoroethyl)amino)-spiro[isoindoline-1,9'-xanthene]-6-carboxamide (26):** Following procedure B with **38** (2.06 mg, 3.33 µmol, 1.0 eq), **26** (1.78 mg, 2.16 µmol, 65 %) was obtained as a colorless solid.

$^1$H NMR (400 MHz, DMSO-*d$_6$*) δ 8.67 (t, *J* = 5.5 Hz, 1H), 8.05 (d, *J* = 8.0 Hz, 1H), 7.94 (d, *J* = 8.0 Hz, 1H), 7.54 (s, 1H), 6.62 (t, *J* = 6.7 Hz, 2H), 6.57 (d, *J* = 2.1 Hz, 2H), 6.43 (dd, *J* = 8.7, 2.2 Hz, 2H), 6.28 (d, *J* = 8.7 Hz, 2H), 4.00 – 3.91 (m, 4H), 3.72 (q, *J* = 9.6 Hz, 2H), 3.58 (t, *J* = 6.6 Hz, 2H), 3.46 – 3.40 (m, 6H), 3.33 – 3.28 (m, 4H), 1.66 (p, *J* = 6.7 Hz, 2H), 1.44 – 1.36 (m, 2H), 1.34 – 1.29 (m, 2H), 1.26 – 1.21 (m, 2H).

HRMS (ESI) Exact mass calculated for $C_{37}H_{38}ClF_9N_4O_5$ $[M+H]^+$: 825.2460, found: 825.2463.

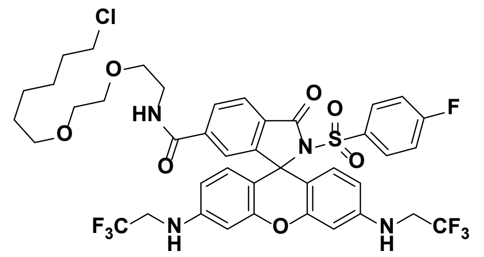

***N*-(2-(2-((6-Chlorohexyl)oxy)ethoxy)ethyl)-2-((4-fluorophenyl)sulfonyl)-3-oxo-3',6'-bis((2,2,2-trifluoroethyl)-amino)spiro[isoindoline-1,9'-xanthene]-6-carboxamide (17):** Following procedure B with **8** (2.80 mg, 4.03 µmol, 1.0 eq), **17** (2.27 mg, 2.52 µmol, 63 %) was obtained as a red solid.

$^1$H NMR (400 MHz, DMSO-*d$_6$*) δ 8.71 (t, *J* = 5.5 Hz, 1H), 8.06 (dd, *J* = 8.0, 1.0 Hz, 1H), 7.94 (d, *J* = 8.1 Hz, 1H), 7.54 (s, 1H), 7.36 – 7.32 (m, 2H), 7.22 (t, *J* = 8.8 Hz, 2H), 6.70 – 6.67 (m, 2H), 6.64 (d, *J* = 2.1 Hz, 2H), 6.35 (dd, *J* = 8.7, 2.2 Hz, 2H), 6.23 (d, *J* = 8.7 Hz, 2H), 4.05 – 3.97 (m, 4H), 3.57 (t, J = 6.5 Hz, 2H), 3.45 – 3.38 (m, 6H), 3.33 – 3.26 (m, 4H), 1.64 (p, *J* = 6.7 Hz, 2H), 1.43 – 1.36 (m, 2H), 1.35 – 1.28 (m, 2H), 1.25 – 1.17 (m, 2H).

HRMS (ESI) Exact mass calculated for $C_{41}H_{40}ClF_7N_4O_7S$ $[M+H]^+$: 901.2267, found: 901.2269.

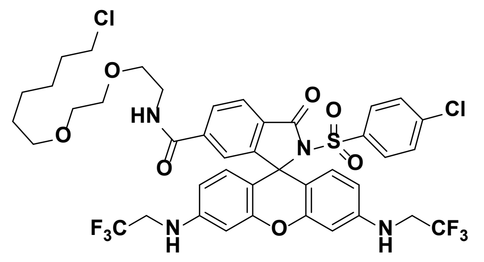

***N*-(2-(2-((6-Chlorohexyl)oxy)ethoxy)ethyl)-2-((4-chloro-phenyl)sulfonyl)-3-oxo-3',6'-bis((2,2,2-trifluoro-ethyl)amino)spiro[isoindoline-1,9'-xanthene]-6-carboxamide (16):** Following procedure B with **7** (2.89 mg, 4.06 µmol, 1.0 eq), **16** (2.53 mg, 2.76 µmol, 68 %) was obtained as a red solid.

$^1$H NMR (400 MHz, DMSO-*d$_6$*) δ 8.71 (t, *J* = 5.5 Hz, 1H), 8.06 (dd, *J* = 8.1, 1.0 Hz, 1H), 7.94 (d, *J* = 8.1 Hz, 1H), 7.54 (s, 1H), 7.46 (d, *J* = 8.7 Hz, 2H), 7.26 (d, *J* = 8.7 Hz, 2H), 6.70 (t, *J* = 6.6 Hz, 2H), 6.64 (d, *J* = 2.1 Hz, 2H), 6.35 (dd, *J* = 8.7, 2.2 Hz, 2H), 6.25 (d, *J* = 8.7 Hz, 2H), 4.06 – 3.97 (m, 4H), 3.57 (t, *J* = 6.6 Hz,

2H), 3.45 – 3.39 (m, 6H), 3.33 – 3.26 (m, 4H), 1.64 (p, *J* = 6.7 Hz, 2H), 1.43 – 1.36 (m, 2H), 1.35 – 1.28 (m, 2H), 1.25 – 1.18 (m, 2H).

HRMS (ESI) Exact mass calculated for $C_{41}H_{40}Cl_2F_6N_4O_7S$ $[M+H]^+$: 917.1972, found: 917.1966.

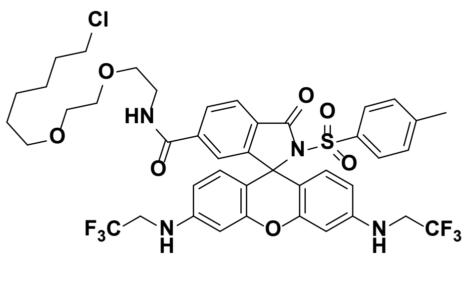

***N*-(2-(2-((6-Chlorohexyl)oxy)ethoxy)ethyl)-3-oxo-2-tosyl-3',6'-bis((2,2,2-trifluoroethyl) amino)spiro[isoindoline-1,9'-xanthene]-6-carboxamide (19):** Following procedure B with **10** (2.55 mg, 3.69 µmol, 1.0 eq), **19** (2.26 mg, 2.52 µmol, 68 %) was obtained as a red solid.

$^1$H NMR (400 MHz, DMSO-*d6*) δ 8.70 (t, *J* = 5.6 Hz, 1H), 8.05 (dd, *J* = 8.1, 1.2 Hz, 1H), 7.91 (d, *J* = 8.1 Hz, 1H), 7.53 (s, 1H), 7.20 – 7.16 (m, 4H), 6.67 (t, *J* = 6.7 Hz, 2H), 6.63 (d, *J* = 2.2 Hz, 2H), 6.35 (dd, *J* = 8.7, 2.3 Hz, 2H), 6.23 (d, *J* = 8.7 Hz, 2H), 4.05 – 3.96 (m, 4H), 3.57 (t, *J* = 6.6 Hz, 2H), 3.46 – 3.39 (m, 6H), 3.33 – 3.27 (m, 4H), 2.33 (s, 3H), 1.65 (p, J = 6.7 Hz, 2H), 1.43 – 1.36 (m, 2H), 1.35 – 1.28 (m, 2H), 1.25 – 1.16 (m, 2H).

HRMS (ESI) Exact mass calculated for $C_{42}H_{43}ClF_6N_4O_7S$ $[M+H]^+$: 897.2518, found: 897.2518.

**General Procedure C for 23:**

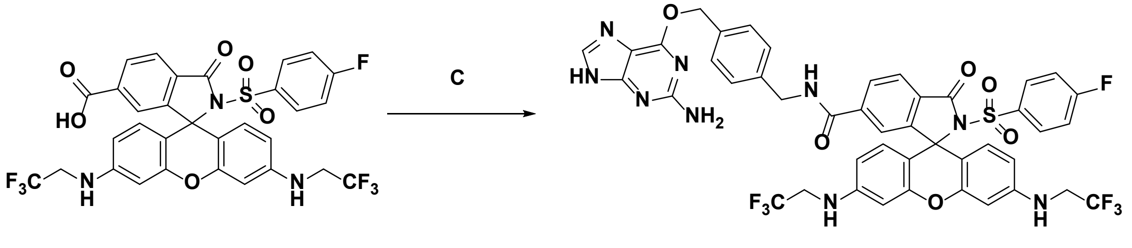

8 23

***N*-(4-(((2-Amino-*9H*-purin-6-yl)oxy)methyl)benzyl)-2-((4-fluorophenyl)sulfonyl)-3-oxo-3',6'-bis((2,2,2-trifluoroethyl)amino)spiro[isoindoline-1,9'-xanthene]-6-carboxamide (23):** 6-(4-Aminomethylbenzyloxy)-*9H*-purin-2-amine (1.63 mg, 6.04 µmol, 1.5 eq) was added to a solution of **8** (2.80 mg, 4.03 µmol, 1 eq), PyBOP (2.72 mg, 5.23 µmol, 1.3 eq) and DIPEA (6.65 µL, 40.3 µmol, 10 eq) in DMSO (600 µL). The reaction mixture was stirred at rt for 30 min followed by purification using preparative HPLC (8 mL/min, 35 % to 90 % MeCN/$H_2O$ (0.1 % TFA) in 55 min) to obtain **23** (3.01 mg, 3.18 µmol, 79%) as a red solid.

$^1$H NMR (400 MHz, MeOD-$d_4$) δ 8.31 (s, 1H), 8.04 (dd, *J* = 8.1, 1.4 Hz, 1H), 7.97 (d, *J* = 8.1 Hz, 1H), 7.54 (s, 1H), 7.48 – 7.43 (m, 4H), 7.30 (d, *J* = 8.2 Hz, 2H), 7.10 – 7.04 (m, 2H), 6.61 (d, *J* = 2.1 Hz, 2H), 6.33 – 6.26 (m, 4H), 5.58 (s, 2H), 4.46 – 4.45 (m, 2H), 3.89 (q, *J* = 9.4 Hz, 4H).

HRMS (ESI) Exact mass calculated for $C_{44}H_{32}F_7N_9O_6S$ $[M+2H]^{2+}$: 474.6115, found: 474.6113.

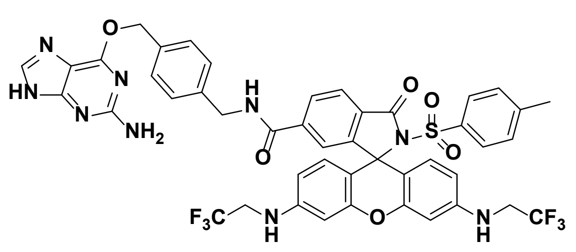

***N*-(4-(((2-Amino-*9H*-purin-6-yl)oxy)methyl) benzyl)-3-oxo-2-tosyl-3',6'-bis((2,2,2-trifluoro-ethyl)amino)spiro[isoindoline-1,9'-xanthene]-6-carboxamide (24):** Following procedure C with **10** (3.10 mg, 4.48 µmol, 1 eq), **24** (2.61 mg, 2.77 µmol, 62 %) was obtained as a red solid.

$^1$H NMR (400 MHz, DMSO-$d_6$) δ 9.22 (t, *J* = 6.0 Hz, 1H), 8.37 (s, 1H), 8.09 (dd, *J* = 8.1, 1.5 Hz, 1H), 7.93 (d, *J* = 8.0 Hz, 1H), 7.55 (s, 1H), 7.44 (d, *J* = 8.1 Hz, 2H), 7.27 (d, *J* = 8.1 Hz, 2H), 7.22 – 7.13 (m, 4H), 6.67 (t, *J* = 7.0 Hz, 2H), 6.63 (d, *J* = 2.3 Hz, 2H), 6.35 (dd, *J* = 8.7, 2.4 Hz, 2H), 6.24 (d, *J* = 8.6 Hz, 2H), 5.47 (s, 2H), 4.37 (d, *J* = 5.8 Hz, 2H), 4.05 – 3.94 (m, 4H), 2.33 (s, 3H).

HRMS (ESI) Exact mass calculated for $C_{45}H_{35}N_9O_6F_6S$ $[M+2H]^{2+}$: 472.6240, found: 472.6236.

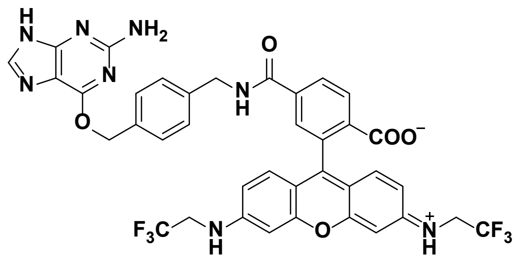

**(*E*)-4-((4-(((2-amino-*9H*-purin-6-yl)oxy)methyl) benzyl)carbamoyl)-2-(6-((2,2,2-trifluoroethyl) amino)-3-((2,2,2-trifluoroethyl)iminio)-*3H*-xanthen-9-yl)benzoate (22):** Following procedure C with **1** (3.30 mg, 6.13 µmol, 1 eq), **22** (4.20 mg, 5.31 µmol, 87 %) was obtained as a red solid.

$^1$H NMR (400 MHz, MeOD-$d_4$) δ 8.40 (d, *J* = 8.2 Hz, 1H), 8.23 (dd, *J* = 8.3, 1.8 Hz, 1H), 7.96 (d, *J* = 6.2 Hz, 1H), 7.84 (t, *J* = 1.9 Hz, 1H), 7.50 – 7.48 (m, 2H), 7.38 (d, *J* = 7.8 Hz, 2H), 7.18 (dd, *J* = 9.2, 1.3 Hz, 2H), 7.14 (d, *J* = 2.2 Hz, 2H), 7.01 – 6.98 (m, 2H), 5.55 (d, *J* = 2.5 Hz, 2H), 4.60 – 4.58 (m, 2H), 4.24 (q, *J* = 9.0 Hz, 4H).

HRMS (ESI) Exact mass calculated for $C_{38}H_{28}F_6N_8O_5$ $[M+2H]^{2+}$: 396.1116, found: 396.1113.

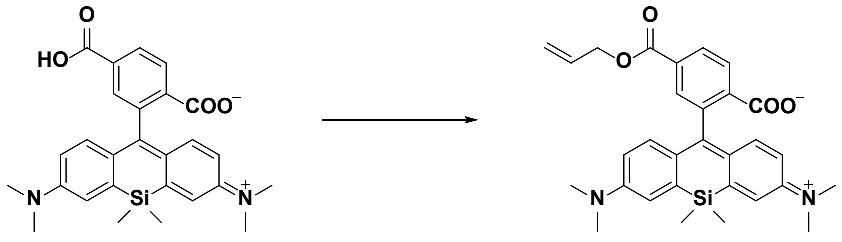

43 50

**4-((Allyloxy)carbonyl)-2-(7-(dimethylamino)-3-(dimethyliminio)-5,5-dimethyl-3,5-dihydrodibenzo[b,e]silin-10-yl)benzoate (50):** Potassium carbonate (40.8 mg, 295 µmol, 2.0 eq) and triethylamine (41.0 µL, 295 µmol, 2.0 eq) were added to a solution of **43** (70.0 mg, 147 µmol, 1.0 eq.) in dry DMF (3 mL). The reaction mixture was cooled to 0 °C and a solution of allyl bromide (19.3 µL, 221 µmol, 1.5 eq.) in dry DMF (0.5 mL) was added dropwise. Afterward, the mixture was allowed to warm to rt and stirred for 2 h, before it was quenched with water and extracted with $CH_2Cl_2$ (3 x). The combined organic layers were dried over $MgSO_4$, filtered and concentrated. The residue was purified by flash column chromatography ($CH_2Cl_2$/MeOH 20:1) to give **50** (65.2 mg, 127 µmol, 86 %) as a yellow solid.

TLC: $R_f$ = 0.51 ($SiO_2$, $CH_2Cl_2$/MeOH 20:1).

$^1$H NMR (400 MHz, $CDCl_3$) δ 8.22 (dd, *J* = 8.0, 1.3 Hz, 1H), 8.02 (d, *J* = 8.0 Hz, 1H), 7.97 (s, 1H), 6.99 (d, *J* = 2.6 Hz, 2H), 6.82 (d, *J* = 8.9 Hz, 2H), 6.58 (dd, *J* = 8.9, 2.8 Hz, 2H), 6.04 – 5.94 (m, 1H), 5.40 – 5.35 (m, 1H), 5.30 – 5.27 (m, 1H), 4.79 (dt, *J* = 5.9, 1.2 Hz, 2H), 2.97 (s, 12H), 0.69 (s, 3H), 0.62 (s, 3H).

$^{13}$C NMR (101 MHz, $CDCl_3$) δ 170.0, 165.2, 155.2, 149.4, 136.7, 135.2, 131.8, 131.3, 130.2, 130.0, 128.0, 125.8, 125.7, 119.2, 116.8, 113.6, 92.1, 66.4, 40.4, 0.4, -1.1.

HRMS (ESI) Exact mass calculated for $C_{30}H_{32}N_2O_4Si$ $[M+H]^+$: 513.2204, found: 513.2197.

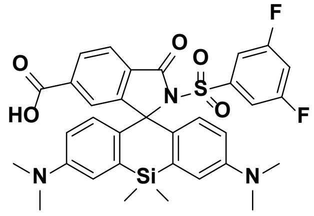

**2'-((3,5-Difluorophenyl)sulfonyl)-3,7-bis(dimethylamino)-5,5-dimethyl-3'-oxo-*5H*-spiro[dibenzo[b,e]siline-10,1'-isoindoline]-6'-carboxylic acid (44):** A *Schlenk* flask was dried with a heat gun *in vacuo* prior to dissolution of **50** (12.0 mg, 23.4 µmol, 1 eq) in dry $CH_2Cl_2$ (2 mL). Phosphorus oxychloride (32.7 µL, 351 µmol, 15 eq) was added, the mixture was heated to 50°C and stirred at this temperature for 2 h. Subsequently, 3,5-difluorobenzenesulfonamide (45.2 mg, 234 µmol, 10 eq) and DIPEA (116 µL, 702 µmol, 30 eq) dissolved in dry MeCN (2 mL) were added. After further dry MeCN (2 mL) was added, the mixture was stirred at 70°C for 10 min. The solvent was evaporated, $H_2O$ (1 mL) was added and the aqueous layer was extracted with $CH_2Cl_2$ (3x). The combined organic layers were dried over $MgSO_4$, filtered and concentrated. The residue was dissolved in MeOH/$CH_2Cl_2$ (5:1,

1.8 mL) in a *Schlenk* flask which had been dried with a heat gun *in vacuo* before. 1,3-Dimethylbarbituric acid (11.0 mg, 70.2 µmol, 3 eq) and tetrakis(triphenylphosphine)palladium (13.5 mg, 11.7 µmol, 0.5 eq) were added and the reaction mixture was stirred at rt for 30 min. After the solvent was evaporated, the crude product was dissolved in DMSO (1.4 mL) and purified by preparative HPLC (8 mL/min, 50 % to 80 % MeCN/$H_2O$ (0.1% formic acid) in 55 min) to obtain **44** (9.50 mg, 14.7 µmol, 63 %) as a green solid.

$^1$H NMR (400 MHz, DMSO-*d*$_6$) δ 8.01 (s, 2H), 7.66 (tt, *J* = 9.0, 2.4 Hz, 1H), 7.16 (s, 1H), 6.96 (d, *J* = 2.9 Hz, 2H), 6.81 – 6.72 (m, 2H), 6.50 (dd, *J* = 9.1, 2.9 Hz, 2H), 6.32 (d, *J* = 9.0 Hz, 2H), 2.93 (s, 12H), 0.59 (s, 3H), 0.55 (s, 3H).

$^{13}$C NMR (101 MHz, DMSO-*d*$_6$) δ 165.9, 165.8, 162.7, 162.6, 160.2, 160.1, 155.6, 148.6, 140.9, 137.4, 135.4, 129.6, 129.5, 129.3, 129.0, 124.7, 124.0, 115.1, 114.4, 111.5, 111.2, 110.3, 110.0, 109.8, 75.6, 0.1, -1.0.

HRMS (ESI) Exact mass calculated for $C_{33}H_{31}F_2N_3O_5SSi$ [M+H]$^+$: 648.1795, found: 648.1790.

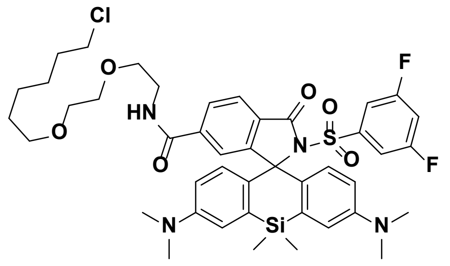

***N*-(2-(2-((6-Chlorohexyl)oxy)ethoxy)ethyl)-2'-((3,5-difluorophenyl)sulfonyl)-3,7-bis(dimethylamino)-5,5-dimethyl-3'-oxo-*5H*-spiro[dibenzo[b,e]siline-10,1'-isoindoline]-6'-carboxamide (29):** Following procedure B with **44** (2.30 mg, 3.55 µmol, 1 eq), **29** (2.37 mg, 2.78 µmol, 78 %) was obtained as a green solid by preparative HPLC (8 mL/min, 55 % to 90 % MeCN/$H_2O$ (0.1% TFA) in 55 min).

$^1$H NMR (400 MHz, DMSO-*d*$_6$) δ 8.70 (t, *J* = 5.6 Hz, 1H), 8.02 – 7.92 (m, 2H), 7.64 (tt, *J* = 9.1, 2.4 Hz, 1H), 7.15 (s, 1H), 6.97 (d, *J* = 2.9 Hz, 2H), 6.79 – 6.69 (m, 2H), 6.50 (dd, *J* = 9.1, 2.9 Hz, 2H), 6.30 (d, *J* = 9.0 Hz, 2H), 3.57 (t, J = 6.6 Hz, 2H), 3.44 – 3.37 (m, 6H), 3.30 – 3.25 (m, 4H), 2.93 (s, 12H), 1.68 – 1.61 (m, 2H), 1.42 – 1.27 (m, 4H), 1.24 – 1.17 (m, 2H), 0.59 (s, 3H), 0.57 (s, 3H).

HRMS (ESI) Exact mass calculated for $C_{43}H_{51}ClF_2N_4O_6SSi$ [M+H]$^+$: 853.3028, found: 853.3030.

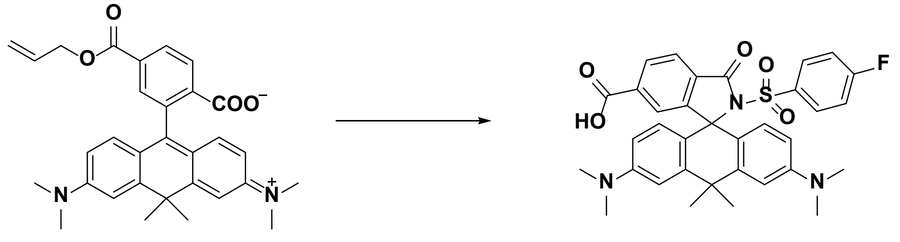

51 46

**3,6-Bis(dimethylamino)-2'-((4-fluorophenyl)sulfonyl)-10,10-dimethyl-3'-oxo-*10H*-spiro[anthracene-9,1'-isoindoline]-6'-carboxylic acid (46): 51** was synthesized according to a procedure reported in.[5] A *Schlenk* flask was dried with a heat gun *in vacuo* prior to dissolution of **51** (11.0 mg, 22.2 µmol, 1 eq) in dry $CH_2Cl_2$ (2 mL). Phosphorus oxychloride (31.0 µL, 332 µmol, 15 eq) was added, the mixture was heated to 50°C and stirred at this temperature for 2 h. Subsequently, 4-fluorobenzenesulfonamide (38.8 mg, 222 µmol, 10 eq) and DIPEA (110 µL, 665 µmol, 30 eq) dissolved in dry MeCN (2 mL) were added. After further dry MeCN (2 mL) was added, the mixture was stirred at 70°C for 10 min. The solvent was evaporated, $H_2O$ (1 mL) was added and the aqueous layer was extracted with $CH_2Cl_2$ (3x). The combined organic layers were dried over $MgSO_4$, filtered and concentrated. The residue was dissolved in MeOH/$CH_2Cl_2$ (5:1, 1.8 mL) in a *Schlenk* flask which had been dried with a heat gun *in vacuo* before. 1,3-Dimethylbarbituric acid (10.4 mg, 66.5 µmol, 3 eq) and tetrakis-(triphenylphosphine)palladium (7.69 mg, 6.65 µmol, 0.3 eq) were added and the reaction mixture was stirred at rt for 30 min. After the solvent was evaporated, the crude product was dissolved in DMSO (1.4 mL) and purified by preparative HPLC (8 mL/min, 40 % to 80 % MeCN/$H_2O$ (0.1 % TFA) in 55 min) to obtain **46** (10.6 mg, 17.3 µmol, 78%) as a blue solid.

$^1$H NMR (400 MHz, DMSO-*d$_6$*) δ 8.02 – 7.96 (m, 2H), 7.43 – 7.40 (m, 2H), 7.31 (t, *J* = 8.8 Hz, 2H), 7.12 (s, 1H), 7.00 (s, 2H), 6.51 (d, *J* = 8.4 Hz, 2H), 6.33 (d, *J* = 8.8 Hz, 2H), 2.98 (s, 12H), 1.89 (s, 3H), 1.82 (s, 3H).

$^{13}$C NMR (101 MHz, DMSO-*d$_6$*) δ 166.4, 165.9, 165.8, 163.9, 158.5, 158.1, 155.2, 149.5, 145.4, 137.1, 134.8, 131.3, 131.2, 129.5, 129.5, 128.0, 124.6, 124.1, 116.0, 115.8, 112.5, 110.5, 71.8, 40.5, 37.7, 36.1, 32.5.

HRMS (ESI) Exact mass calculated for $C_{34}H_{32}FN_3O_5S$ $[M+H]^+$: 614.2119, found: 614.2120.

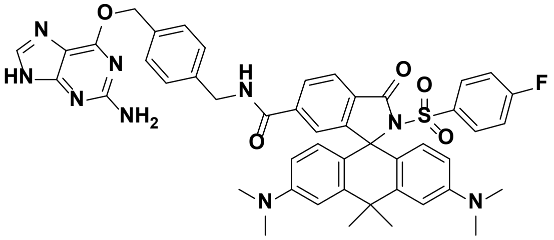


***N*-(4-(((2-Amino-*9H*-purin-6-yl)oxy)methyl) benzyl)-3,6-bis(dimethylamino)-2'-((4-fluorophenyl)sulfonyl)-10,10-dimethyl-3'-oxo-10H-spiro[anthracene-9,1'-isoindoline]-6'-carboxamide (32):** Following procedure C with **46** (4.00 mg, 6.52 µmol, 1 eq), **32** (3.66 mg, 4.23 µmol, 65 %) was obtained as a blue solid.

$^1$H NMR (400 MHz, DMSO-*d$_6$*) δ 9.23 (t, *J* = 5.9 Hz, 1H), 8.31 (s, 1H), 7.99 (dd, *J* = 8.1, 1.3 Hz, 1H), 7.94 (d, *J* = 8.1 Hz, 1H), 7.43 – 7.38 (m, 4H), 7.30 – 7.23 (m, 4H), 7.12 (s, 1H), 6.92 (d, *J* = 2.4 Hz, 2H), 6.45 (dd, *J* = 8.9, 2.5 Hz, 2H), 6.28 (d, *J* = 8.8 Hz, 2H), 5.46 (s, 2H), 4.35 (d, *J* = 5.7 Hz, 2H), 2.95 (s, 12H), 1.89 (s, 3H), 1.82 (s, 3H).

HRMS (ESI) Exact mass calculated for $C_{47}H_{44}FN_9O_5S$ $[M+2H]^{2+}$: 433.6658, found: 433.6660.

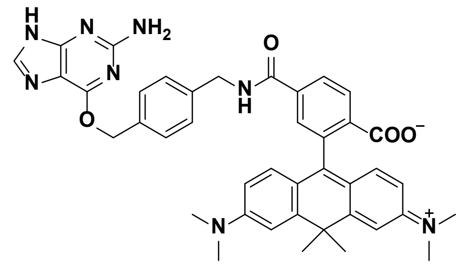

**4-((4-(((2-Amino-*9H*-purin-6-yl)oxy)methyl)benzyl)-carbamoyl)-2-(6-(dimethylamino)-3-(dimethyl-iminio)-10,10-dimethyl-3,10-dihydroanthracen-9-yl)benzoate (30):** Following procedure C with **45** (3.00 mg, 6.57 µmol, 1 eq), **30** (3.95 mg, 5.57 µmol, 85 %) was obtained as a dark blue solid.

$^1$H NMR (400 MHz, MeOD-*d$_4$*) δ 9.29 (t, *J* = 5.9 Hz, 1H), 8.34 (d, *J* = 8.2 Hz, 1H), 8.16 (dd, *J* = 8.2, 1.7 Hz, 1H), 8.06 (s, 1H), 7.77 (d, *J* = 1.6 Hz, 1H), 7.48 (d, *J* = 8.1 Hz, 2H), 7.38 (d, *J* = 8.1 Hz, 2H), 7.22 (d, *J* = 2.4 Hz, 2H), 6.99 (d, *J* = 9.4 Hz, 2H), 6.78 (dd, *J* = 9.4, 2.5 Hz, 2H), 5.57 (s, 2H), 4.59 – 4.57 (m, 2H), 3.31 (s, 12H), 1.86 (s, 3H), 1.75 (s, 3H).

HRMS (ESI) Exact mass calculated for $C_{41}H_{40}N_8O_4$ $[M+2H]^{2+}$: 355.1659, found: 355.1659.

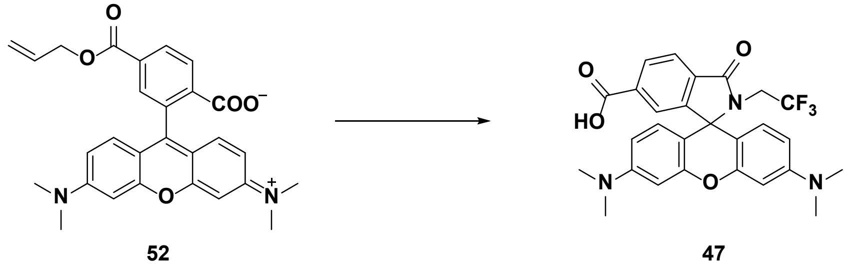

**3',6'-Bis(dimethylamino)-3-oxo-2-(2,2,2-trifluoroethyl)spiro[isoindoline-1,9'-xanthene]-6-carboxylic acid (47): 52** was synthesized according to a procedure reported in.[5] A solution of **52** (25.0 mg, 53.1 µmol, 1 eq.), 2,2,2-trifluoroethylamine (84.9 µL, 1.06 mmol, 20 eq.), 1-(3-dimethylaminopropyl)-3-ethylcarbodiimide hydrochloride (40.7 mg, 213 µmol, 4 eq.) and 4-dimethylaminopyridine (26.0 mg, 213 µmol, 4 eq.) in $CH_2Cl_2$ (1 mL) was heated to 50°C and stirred at this temperature for 12 h in a sealed tube. $H_2O$ (2 mL) was added and the aqueous layer was extracted with $CH_2Cl_2$ (3x). The combined organic layers were dried over $MgSO_4$, filtered and concentrated. The residue was dissolved in MeOH/$CH_2Cl_2$ (5:1, 3 mL) in a *Schlenk* flask which had been dried with a heat gun *in vacuo* before. 1,3-Dimethylbarbituric acid (16.6 mg, 106 µmol, 2 eq.) and tetrakis(triphenylphosphine)palladium (12.3 mg, 10.6 µmol, 0.1 eq.) were added and the reaction mixture was stirred at rt for 30 min. After the solvent was evaporated, the crude product was dissolved in DMSO (3.2 mL) and purified by preparative HPLC (8 mL/min, 45% to 80% MeCN/$H_2O$ (0.1% formic acid) in 55 min) to obtain **47** (16.7 mg, 32.6 µmol, 61 %) as a slightly magenta solid.

$^1$H NMR (400 MHz, DMSO-*d$_6$*) δ 8.09 (dd, *J* = 7.9, 1.2 Hz, 1H), 7.98 (d, *J* = 7.9 Hz, 1H), 7.42 (s, 1H), 6.45 – 6.43 (m, 4H), 6.37 – 6.35 (m, 2H), 3.75 (q, *J* = 9.7 Hz, 2H), 2.92 (s, 12H).

$^{13}C$ NMR (101 MHz, DMSO-$d_6$) δ 167.4, 166.3, 153.9, 152.4, 151.3, 136.0, 131.8, 129.7, 128.2, 128.0, 125.2, 124.4, 123.4, 122.4, 109.2, 104.2, 98.2, 64.9, 41.8, 41.5, 41.1, 40.7.

HRMS (ESI) Exact mass calculated for $C_{27}H_{24}F_3N_3O_4$ $[M+H]^+$: 512.1792, found: 512.1793.

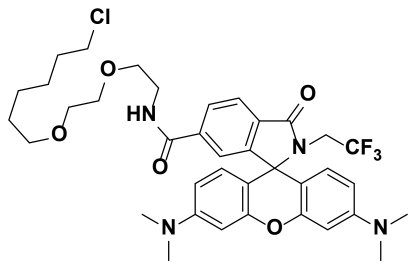


***N*-(2-(2-((6-Chlorohexyl)oxy)ethoxy)ethyl)-3',6'-bis(dimethylamino)-3-oxo-2-(2,2,2-trifluoroethyl)spiro-[isoindoline-1,9'-xanthene]-6-carboxamide (33):** Following procedure B with **47** (4.00 mg, 7.82 µmol, 1 eq.), **33** (3.10 mg, 4.32 µmol, 55 %) was obtained as a colorless solid by preparative HPLC (8 mL/min, 60 % to 95 % MeCN/$H_2O$ (0.1 % formic acid) in 55 min).

$^1H$ NMR (400 MHz, DMSO-$d_6$) δ 8.67 (t, *J* = 5.6 Hz, 1H), 8.04 (dd, *J* = 8.0, 1.3 Hz, 1H), 7.95 (d, *J* = 8.0 Hz, 1H), 7.49 (s, 1H), 6.45 – 6.42 (m, 4H), 6.36 – 6.33 (m, 2H), 3.72 (q, *J* = 9.7 Hz, 2H), 3.58 (t, *J* = 6.6 Hz, 2H), 3.47 – 3.43 (m, 4H), 3.41 – 3.39 (m, 2H), 3.31 – 3.27 (m, 4H), 2.92 (s, 12H), 1.66 (p, *J* = 6.7 Hz, 2H), 1.40 (p, *J* = 6.8 Hz, 2H), 1.36 – 1.19 (m, 4H).

HRMS (ESI) Exact mass calculated for $C_{37}H_{44}N_4O_5ClF_3$ $[M+H]^+$: 717.3025, found: 717.3019.

1H (DMSO); 298.0 K; 400.15 MHZ

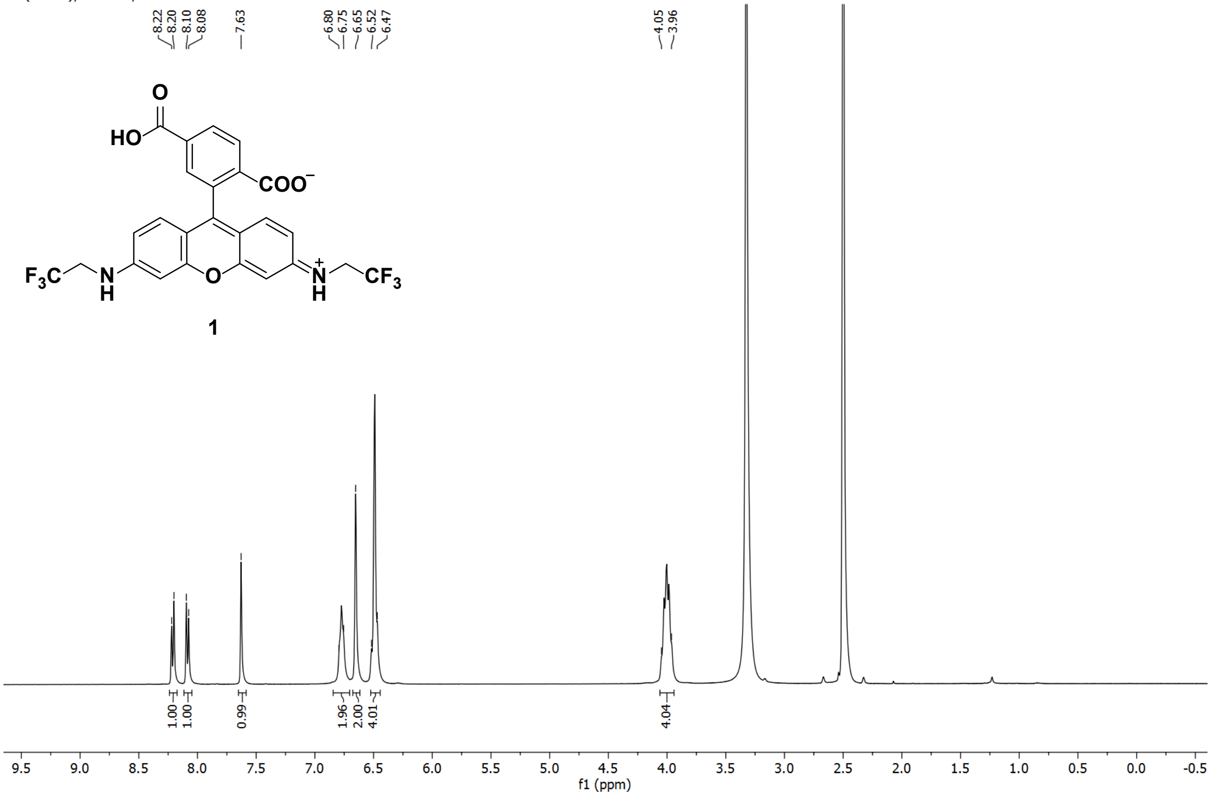


13C (DMSO); 298.0 K; 100.63 MHZ

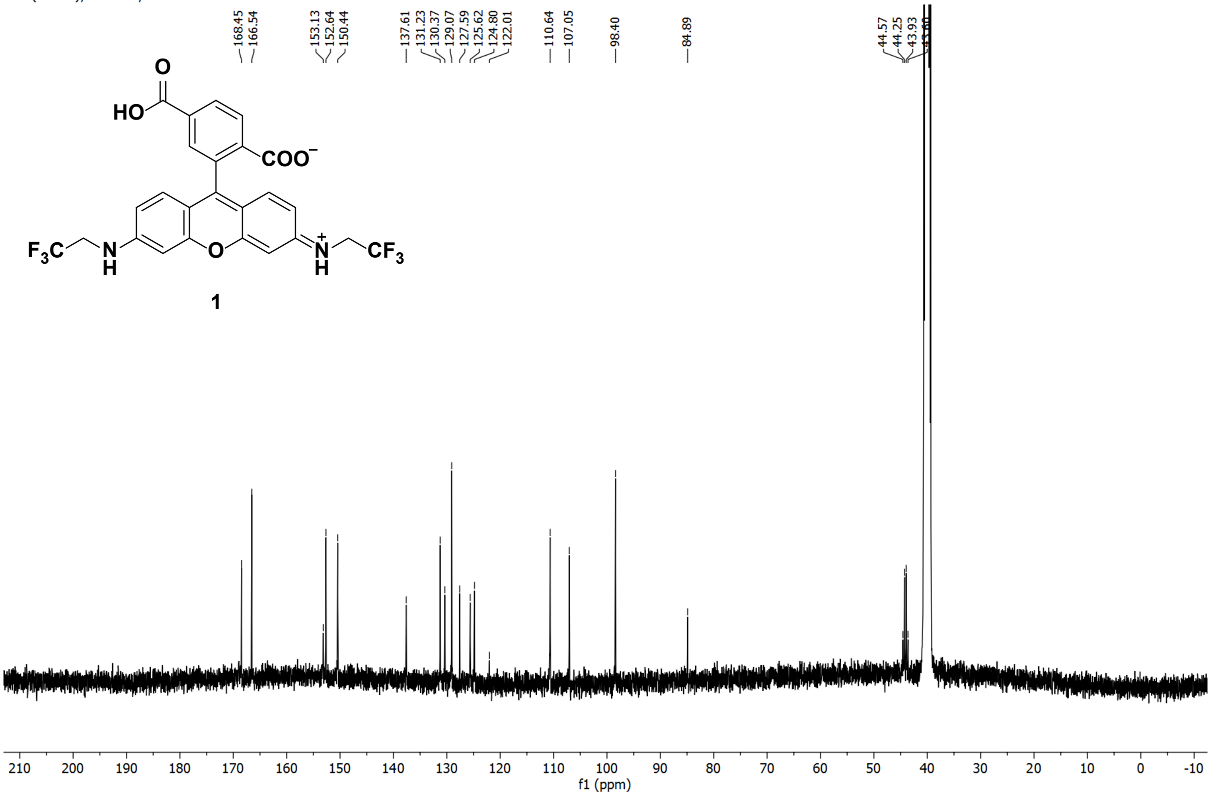

1H (DMSO); 298.0 K; 400.15 MHZ

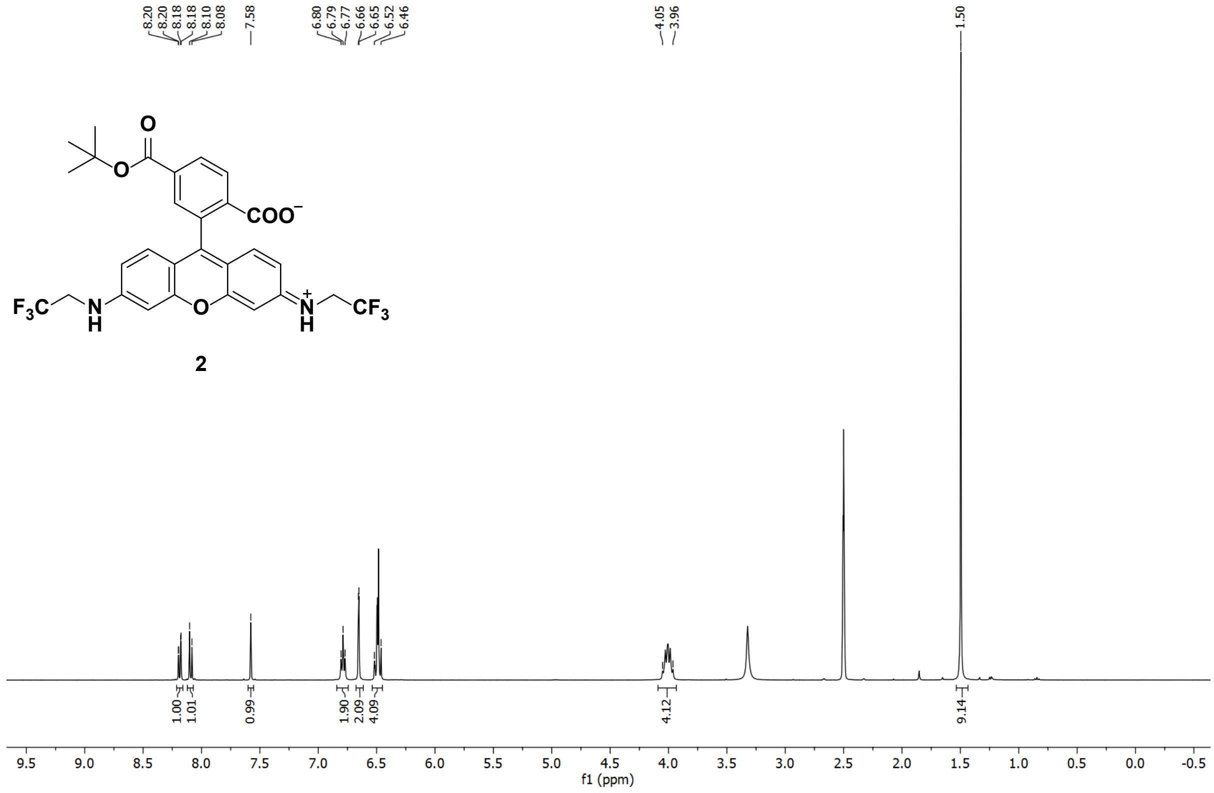


13C (DMSO); 298.0 K; 100.63 MHZ

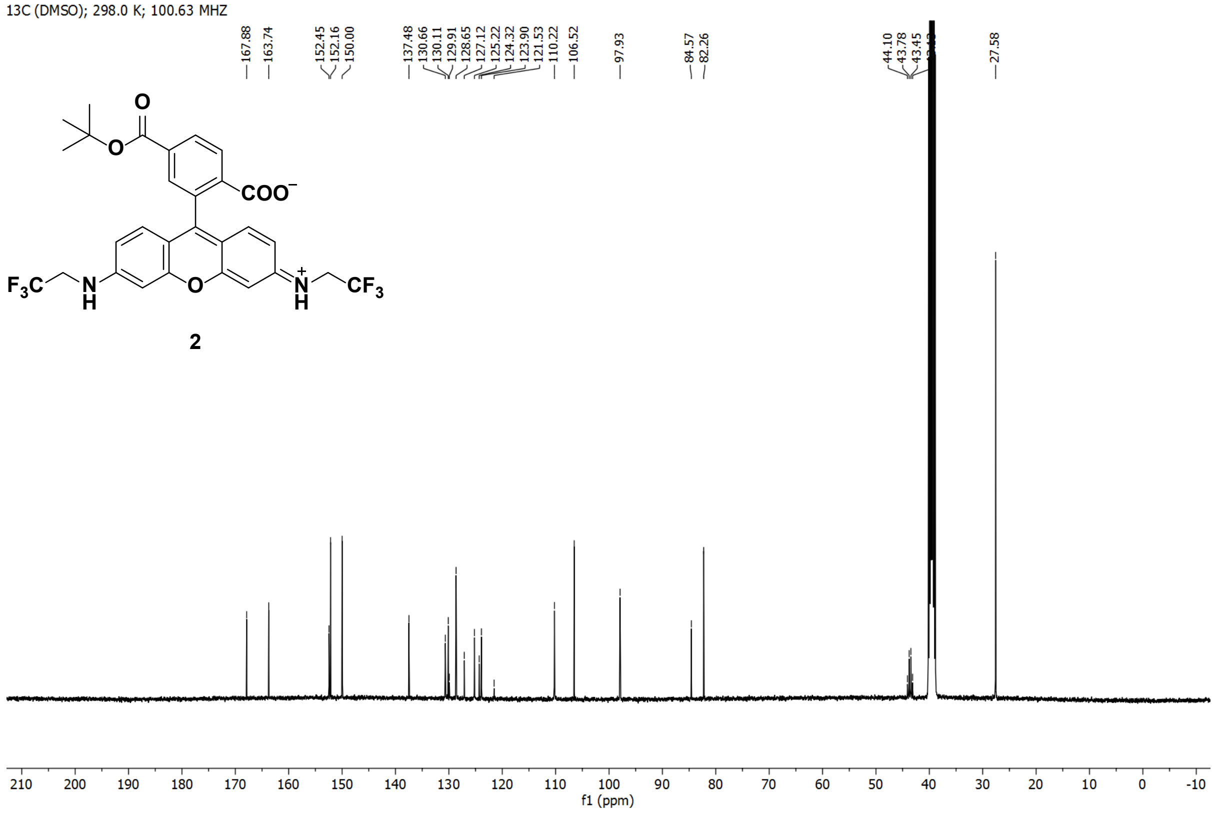

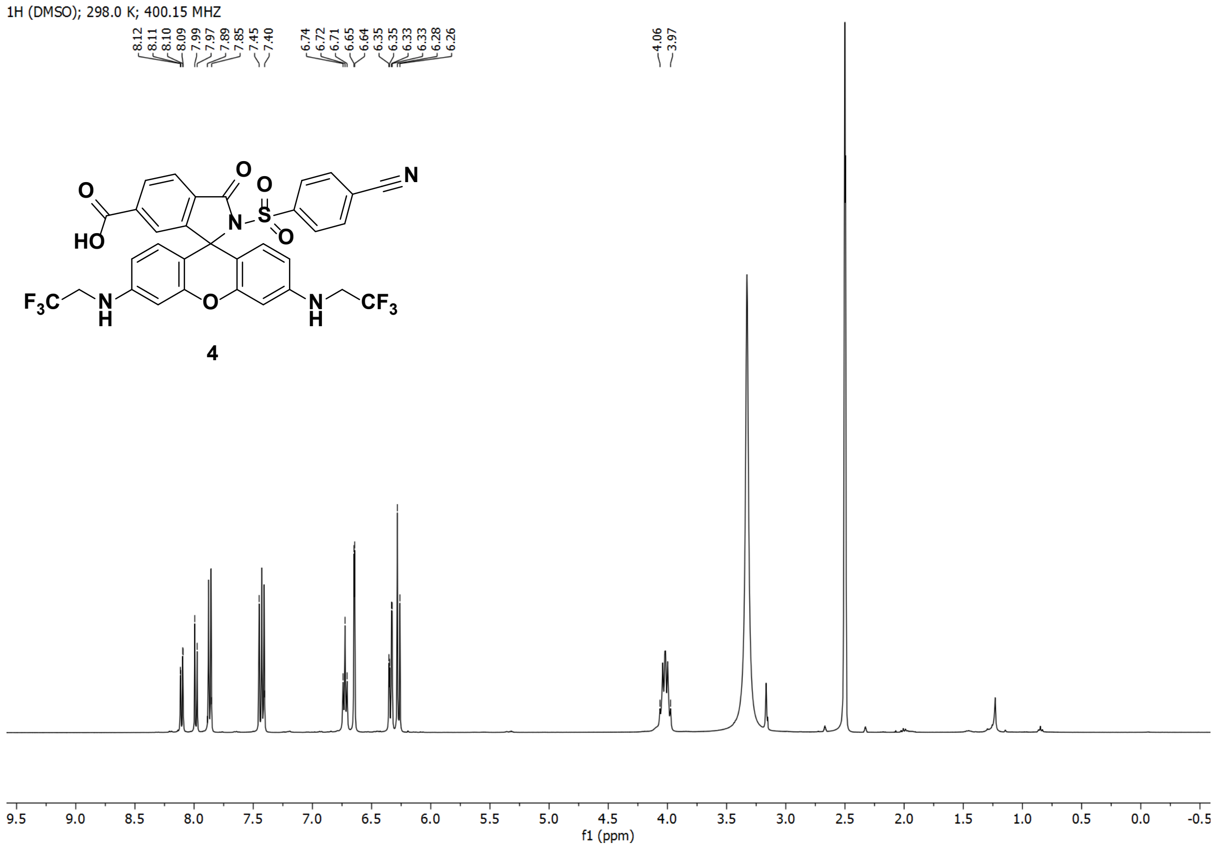
1H (DMSO); 298.0 K; 400.15 MHZ
8.12 8.11 8.10 8.09 7.99 7.97 7.89 7.85 7.45 7.40
6.74 6.72 6.71 6.65 6.64 6.35 6.35 6.33 6.33 6.28 6.26
4.06 3.97
4
f1 (ppm)

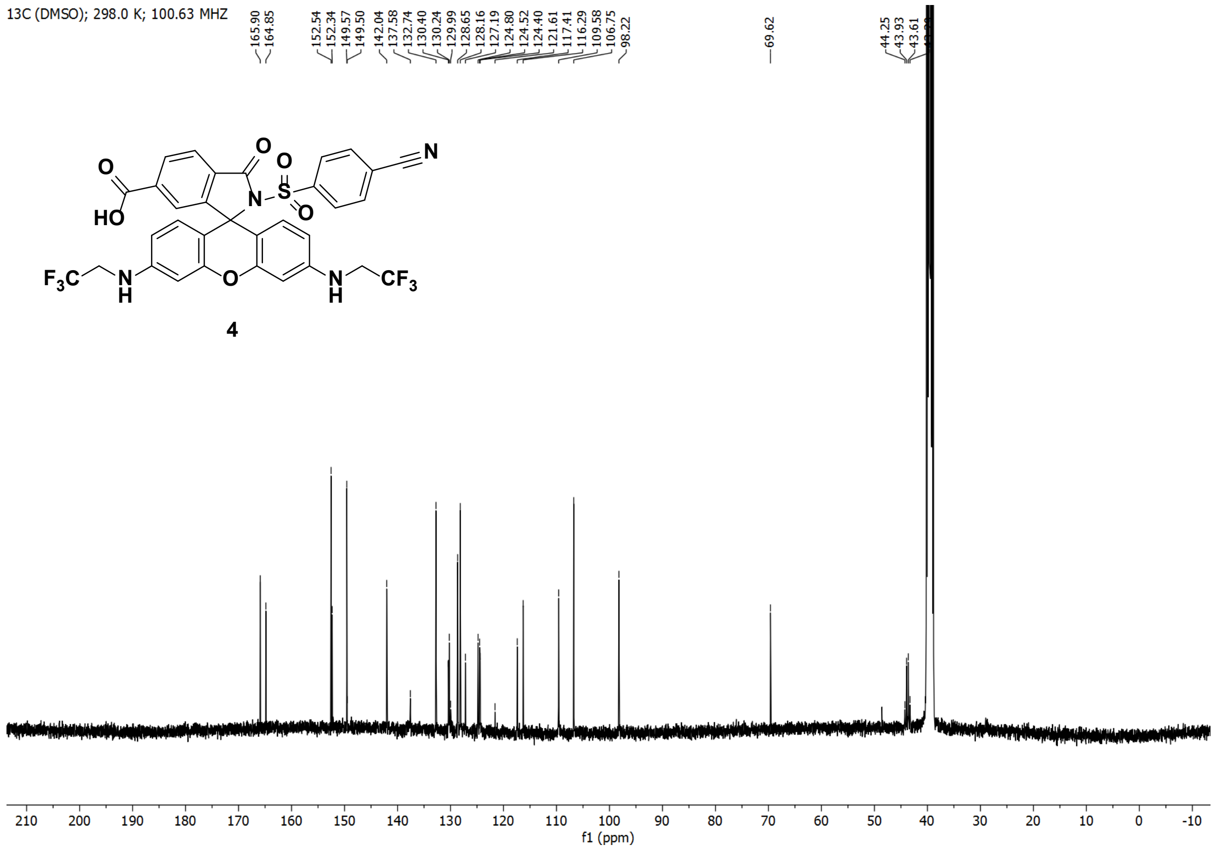
13C (DMSO); 298.0 K; 100.63 MHZ
165.90 164.85
152.54 152.34 149.57 149.50 142.04 137.58 132.74 130.40 130.24 129.99 128.65 128.16 127.19 124.80 124.52 124.40 121.61 117.41 116.29 109.58 106.75 98.22
69.62
44.25 43.93 43.61
4
f1 (ppm)

1H (MeOD); 298.0 K; 400.15 MHZ

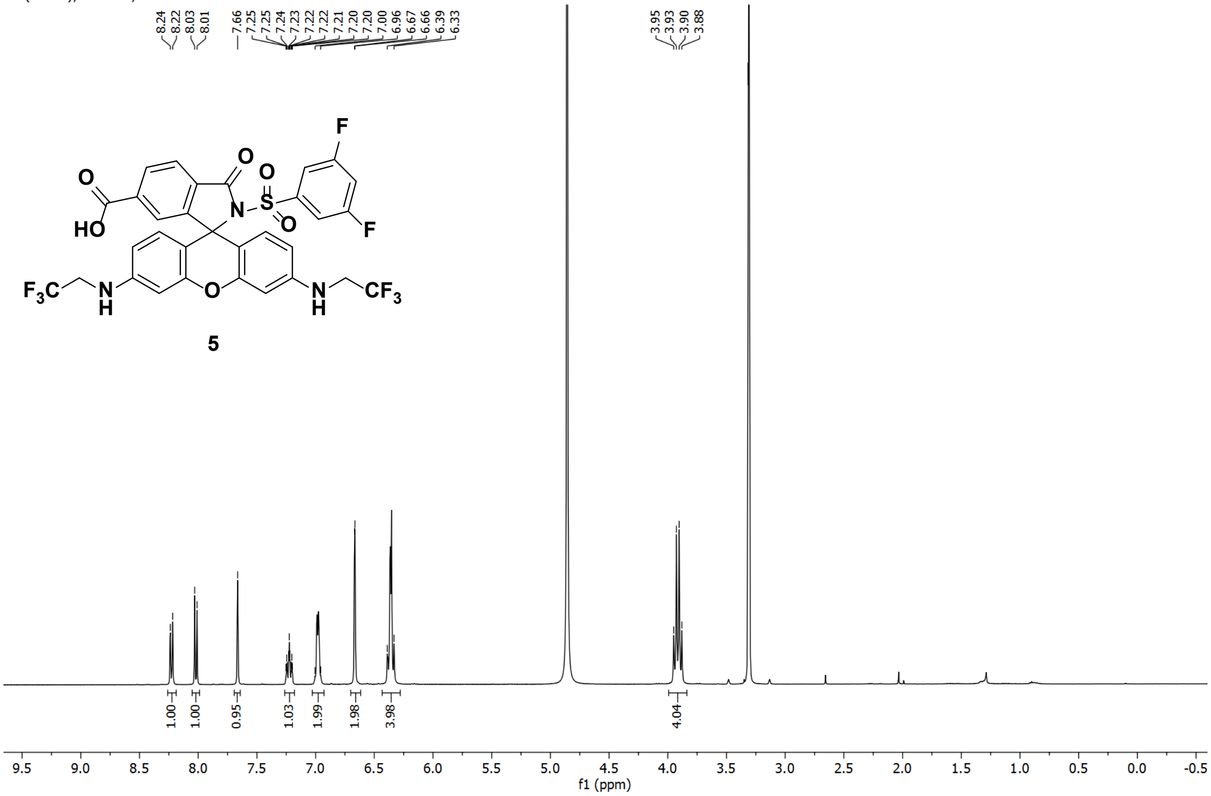


13C (MeOD); 298.0 K; 100.63 MHZ

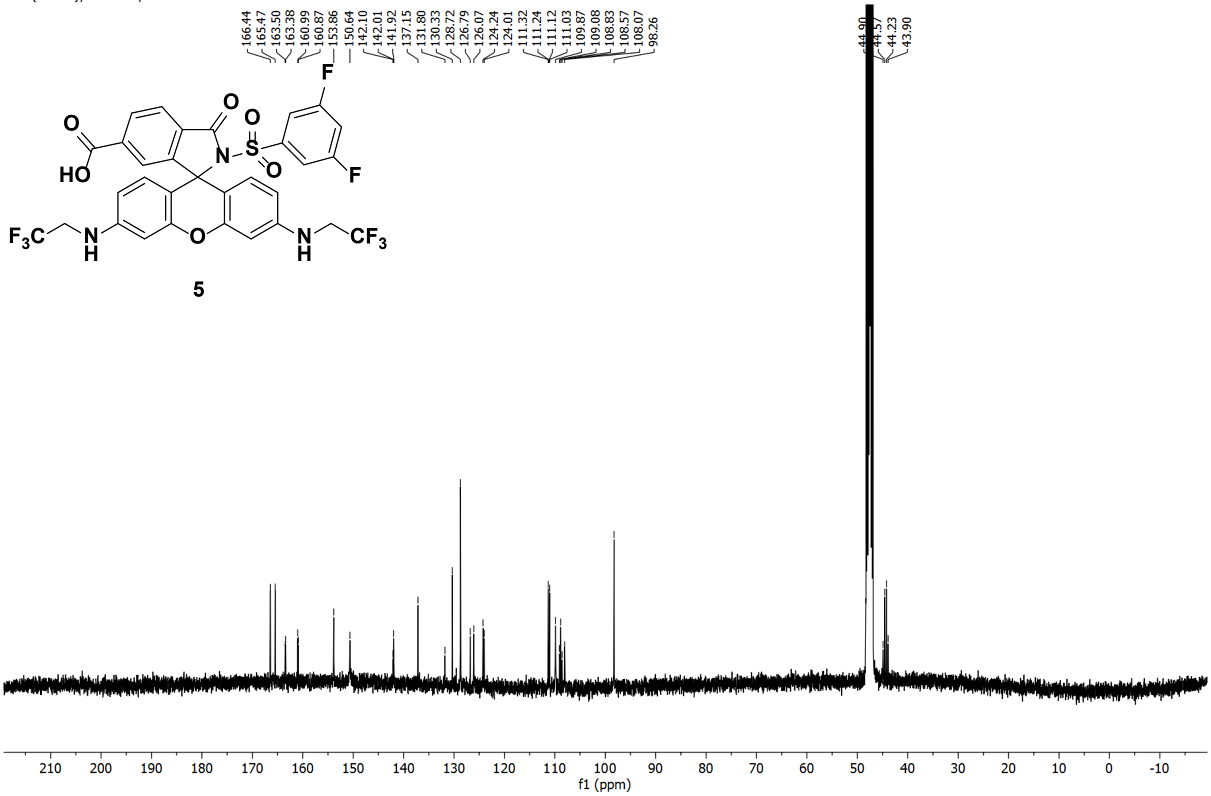

1H (MeOD); 298.0 K; 400.15 MHZ

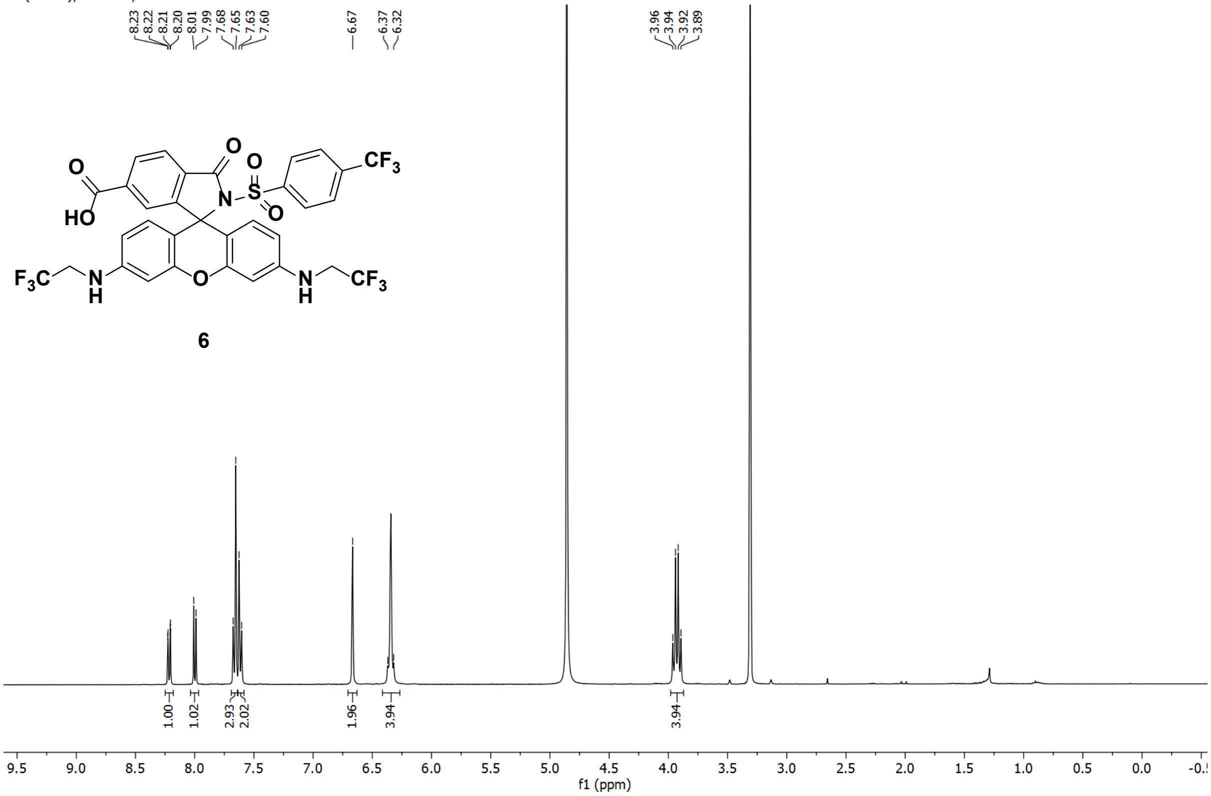


13C (DMSO); 298.0 K; 100.63 MHZ

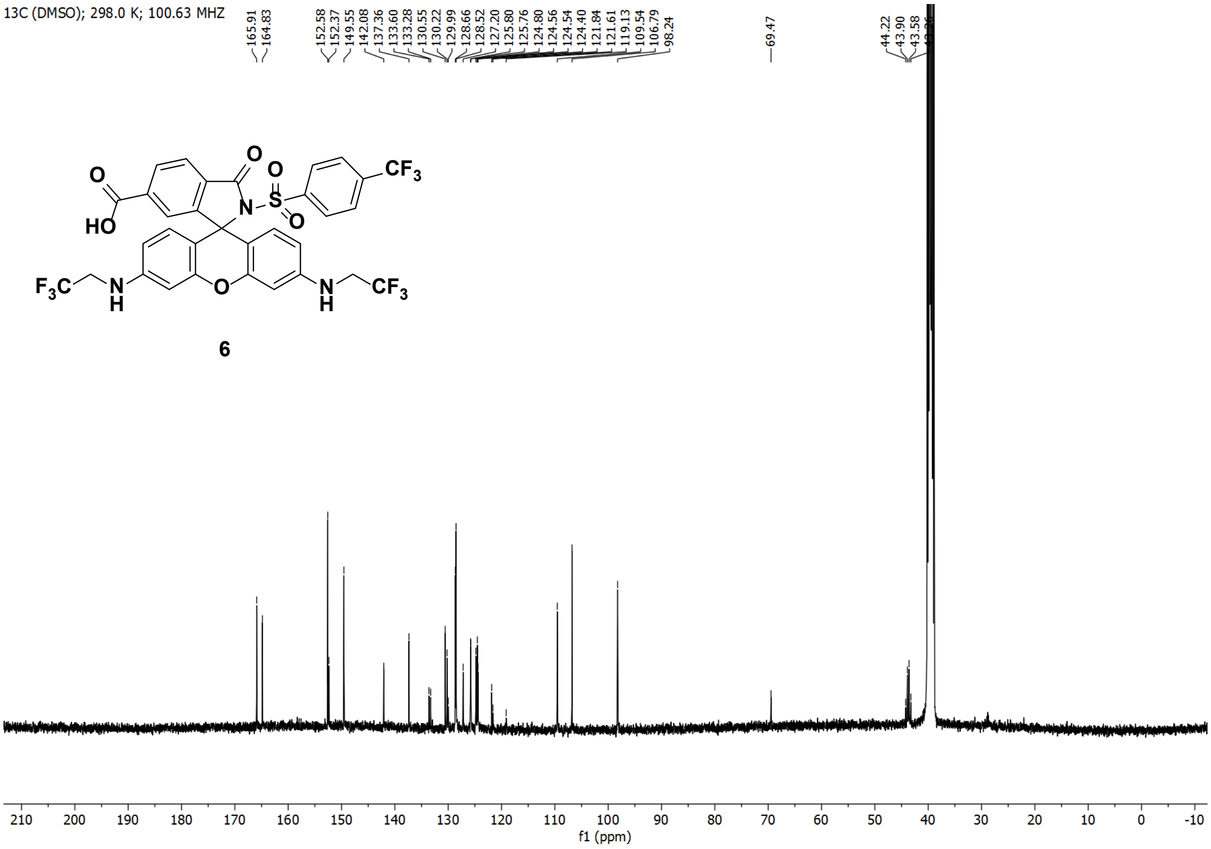

1H (MeOD); 298.0 K; 400.15 MHZ

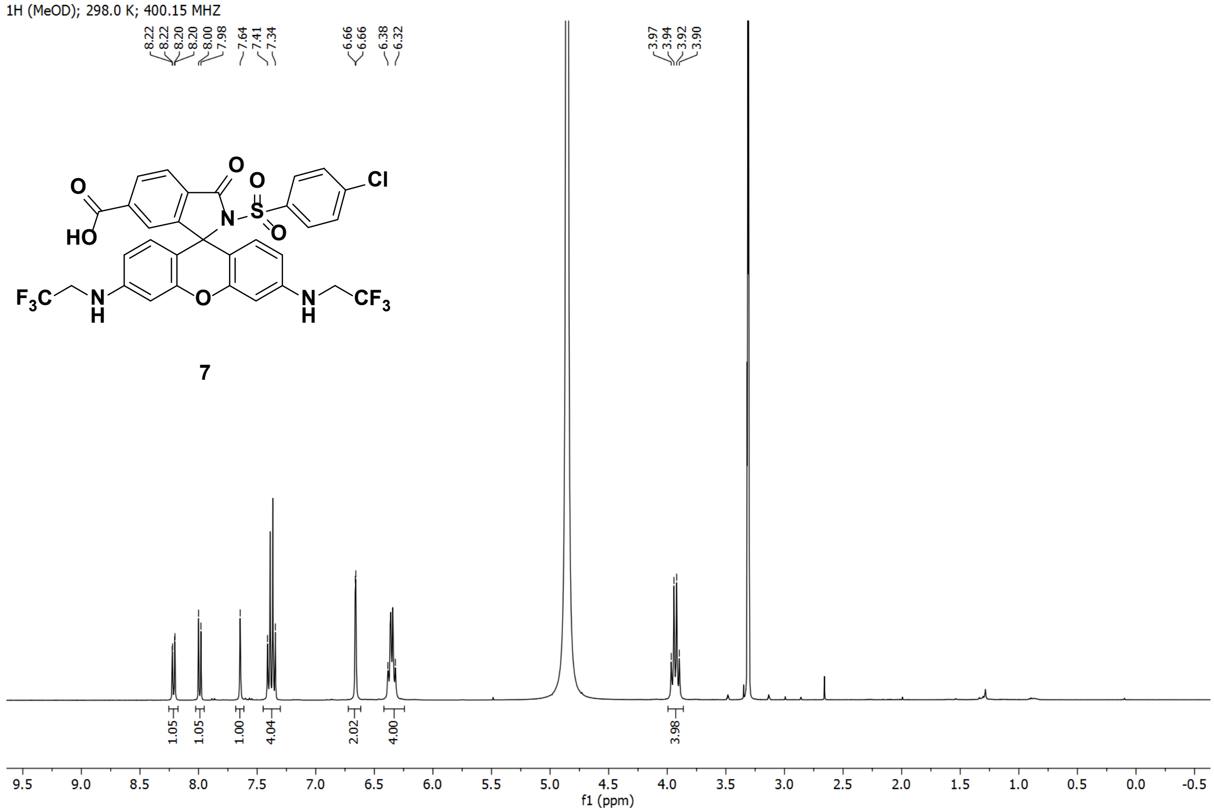


13C (DMSO); 298.0 K; 100.63 MHZ

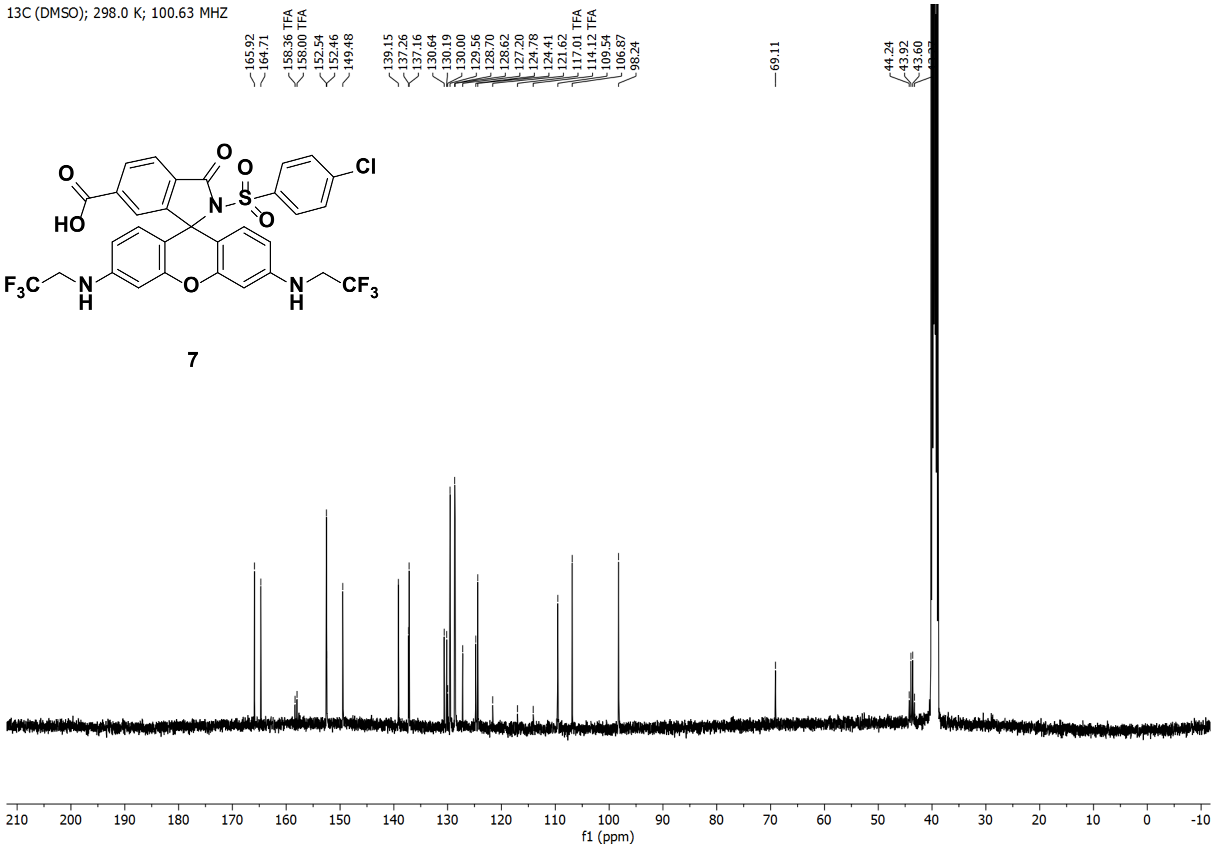

1H (MeOD); 298.0 K; 400.15 MHZ

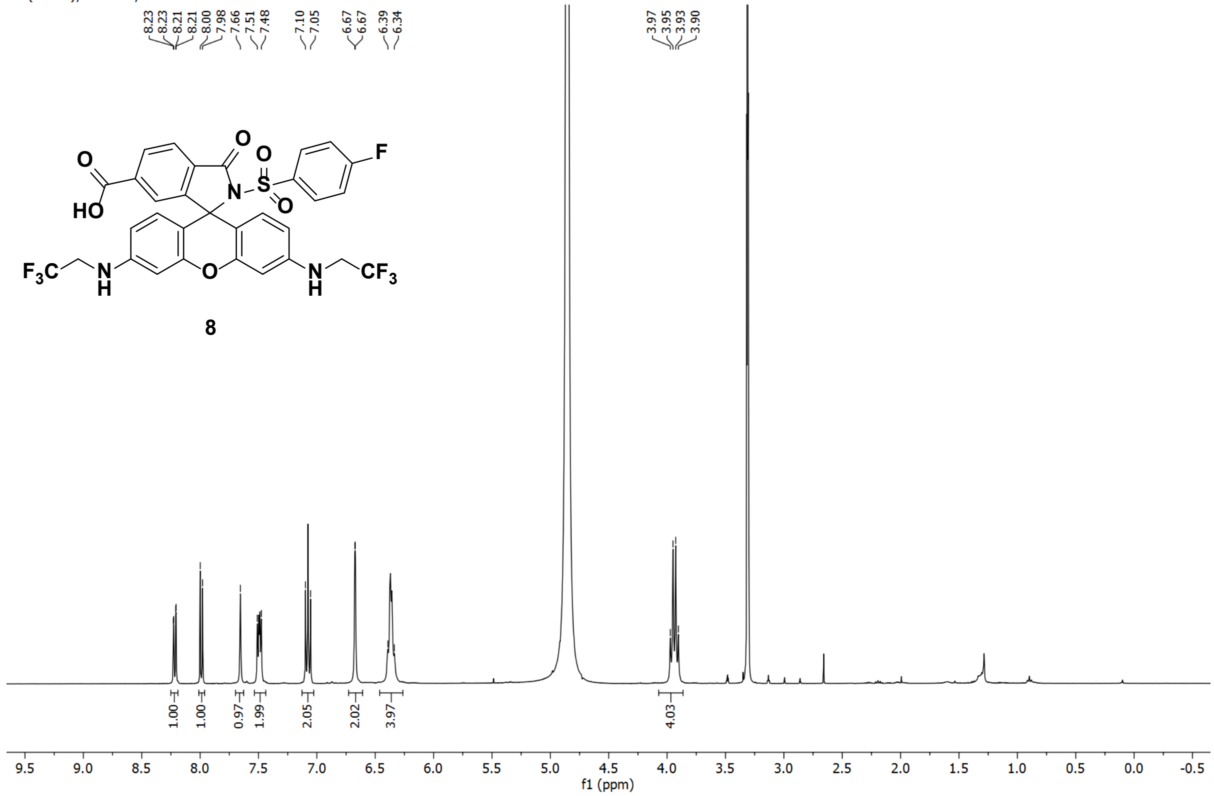


13C (DMSO); 298.0 K; 100.63 MHZ

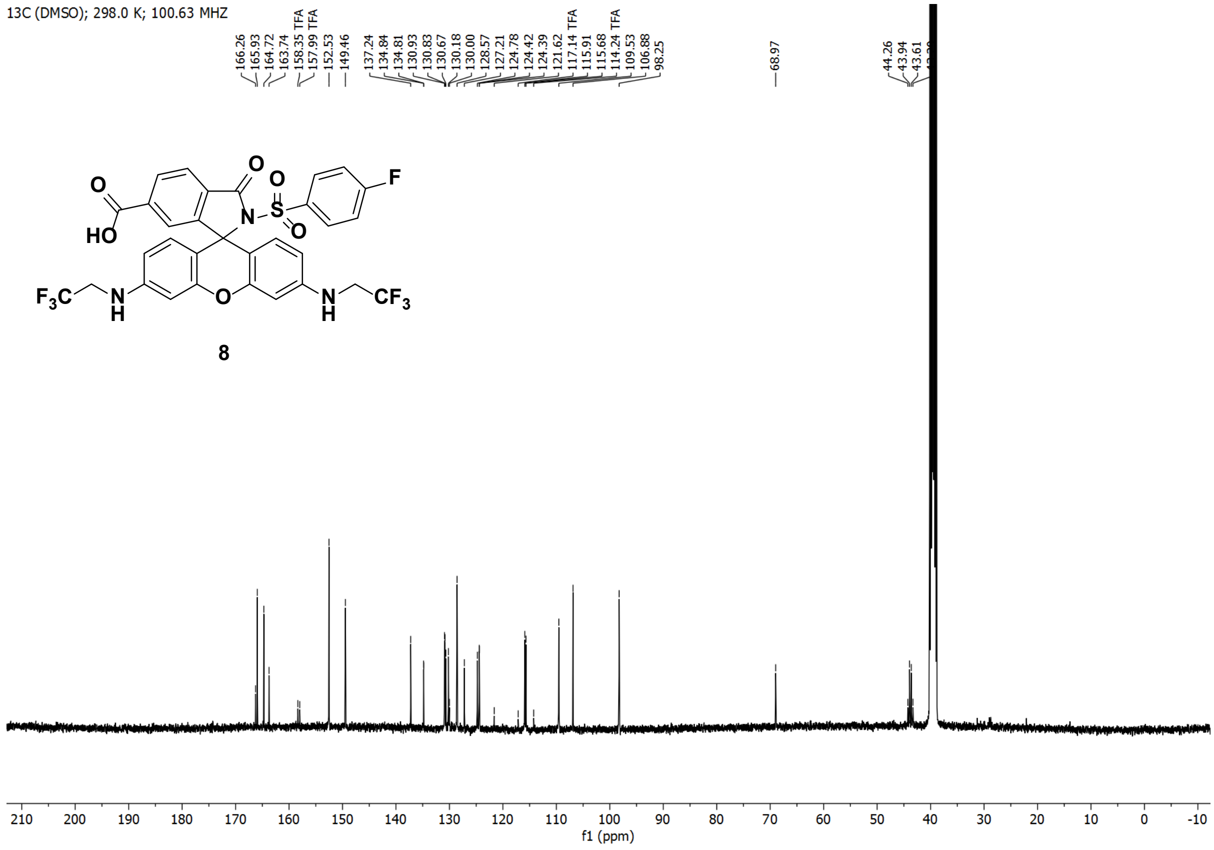

1H (MeOD); 298.0 K; 400.15 MHZ

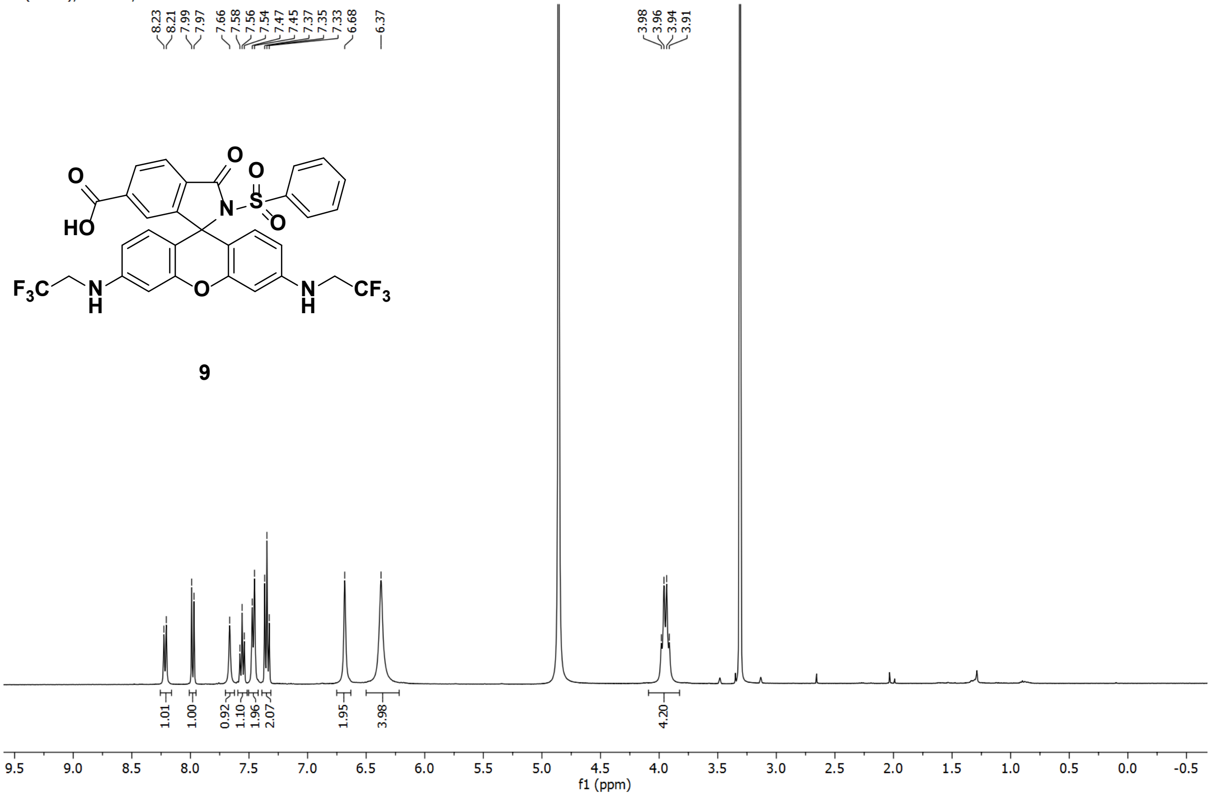


**9**

13C (DMSO); 298.0 K; 100.63 MHZ

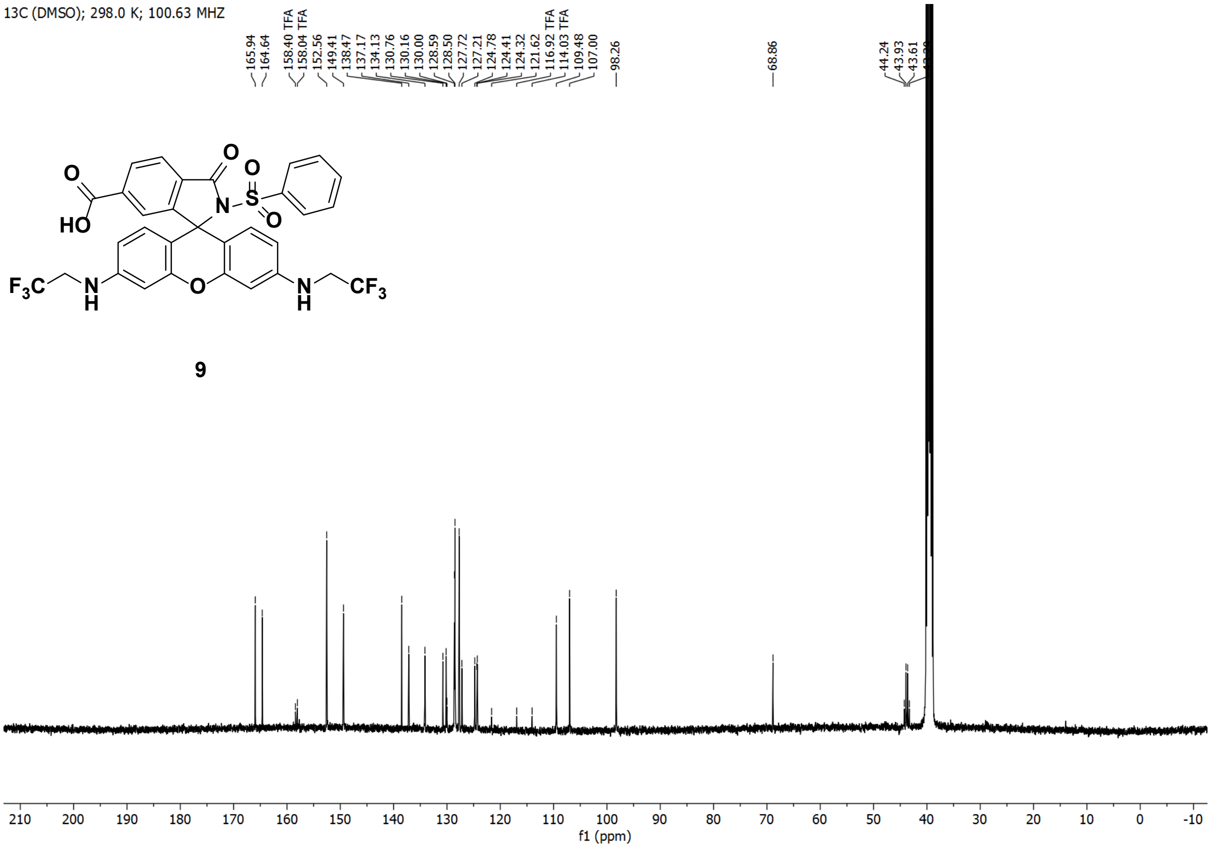


**9**

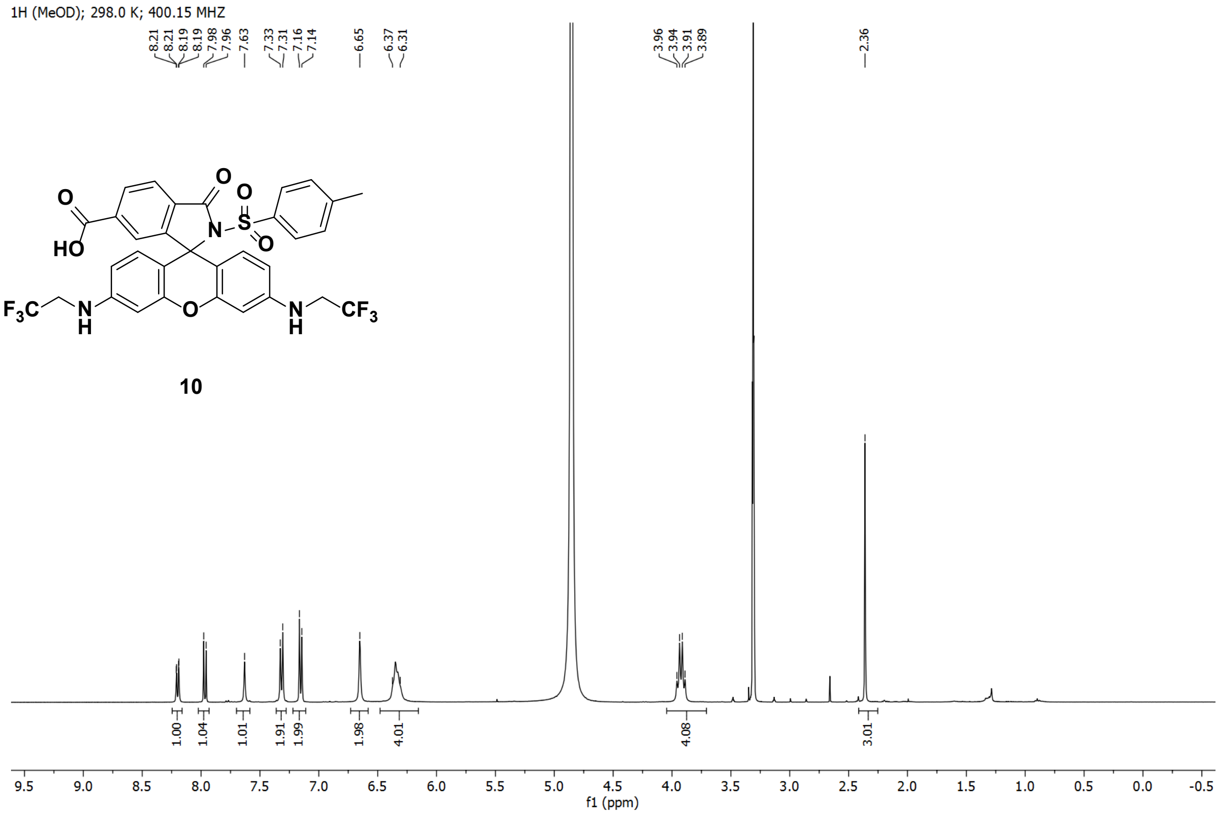

1H (MeOD); 298.0 K; 400.15 MHZ
8.21
8.21
8.19
8.19
7.98
7.96
7.63
7.33
7.31
7.16
7.14
6.65
6.37
6.31
3.96
3.94
3.91
3.89
2.36
10
1.00
1.04
1.01
1.91
1.99
1.98
4.01
4.08
3.01
f1 (ppm)


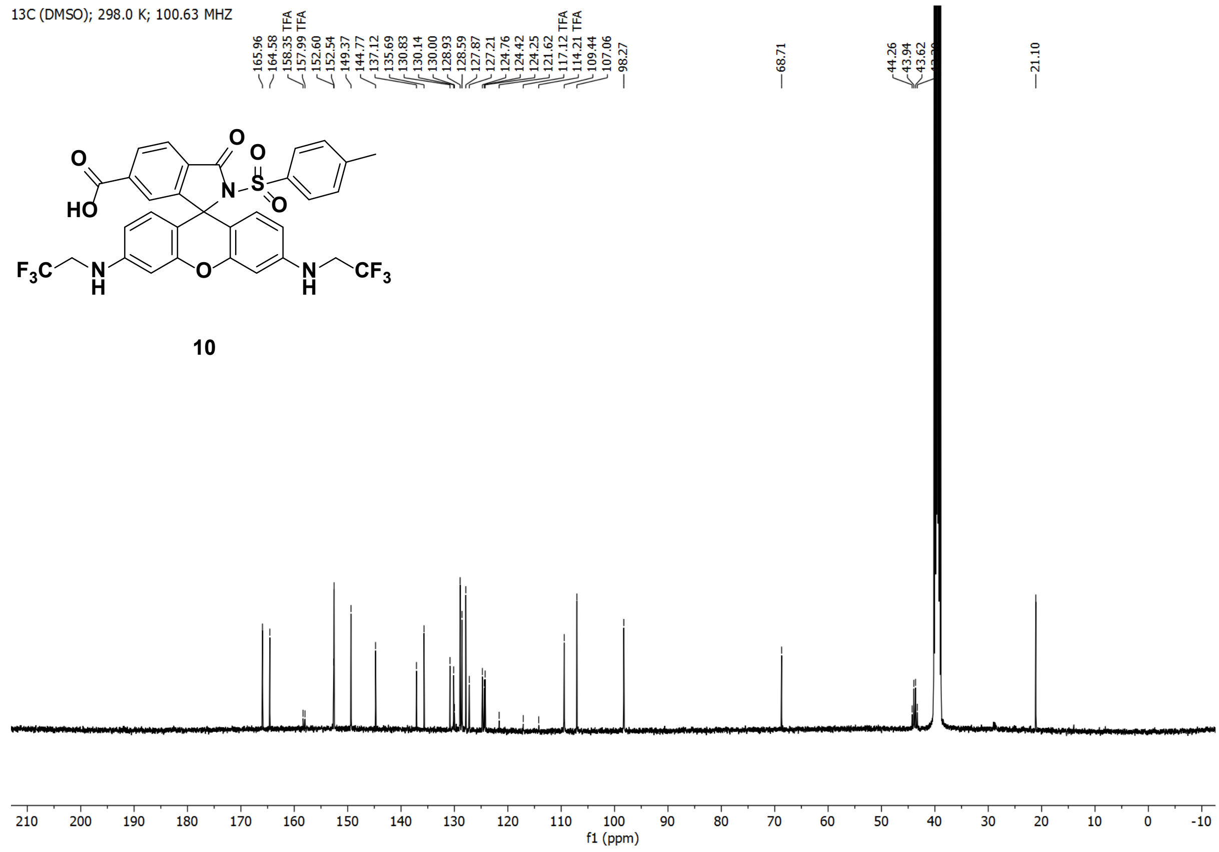

13C (DMSO); 298.0 K; 100.63 MHZ
165.96
164.58
158.35 TFA
157.99 TFA
152.60
152.54
149.37
144.77
137.12
135.69
130.83
130.14
130.00
128.93
128.59
127.87
127.21
124.76
124.42
124.25
121.62
117.12 TFA
114.21 TFA
109.44
107.06
98.27
68.71
44.26
43.94
43.62
21.10
10
f1 (ppm)

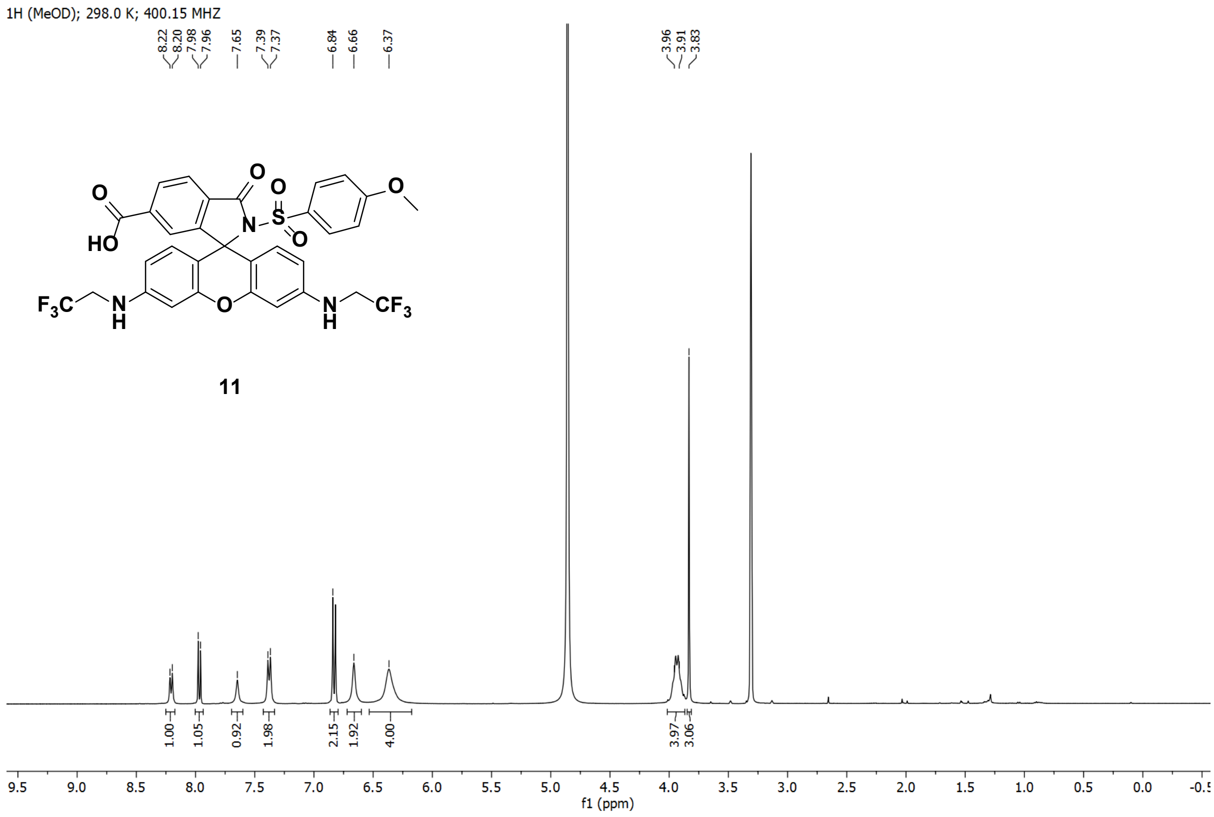
1H (MeOD); 298.0 K; 400.15 MHZ
8.22 8.20 7.98 7.96 7.65 7.39 7.37 6.84 6.66 6.37 3.96 3.91 3.83
1.00 1.05 0.92 1.98 2.15 1.92 4.00 3.97 3.06
HO O O O S O N O F3C N H O N H CF3 O
11
f1 (ppm)

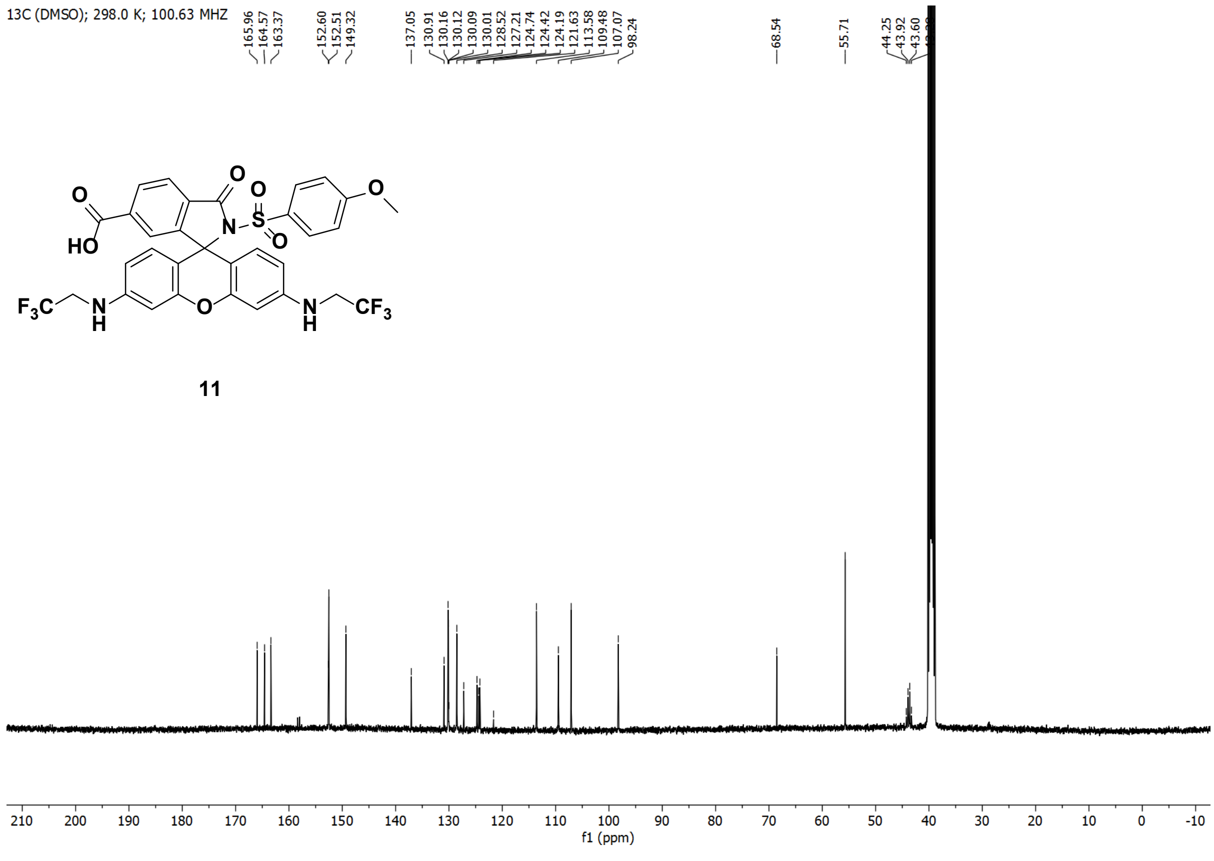
13C (DMSO); 298.0 K; 100.63 MHZ
165.96 164.57 163.37 152.60 152.51 149.32 137.05 130.91 130.16 130.12 130.09 130.01 128.52 127.21 124.74 124.42 124.19 121.63 113.58 109.48 107.07 98.24 68.54 55.71 44.25 43.92 43.60
HO O O O S O N O F3C N H O N H CF3 O
11
f1 (ppm)

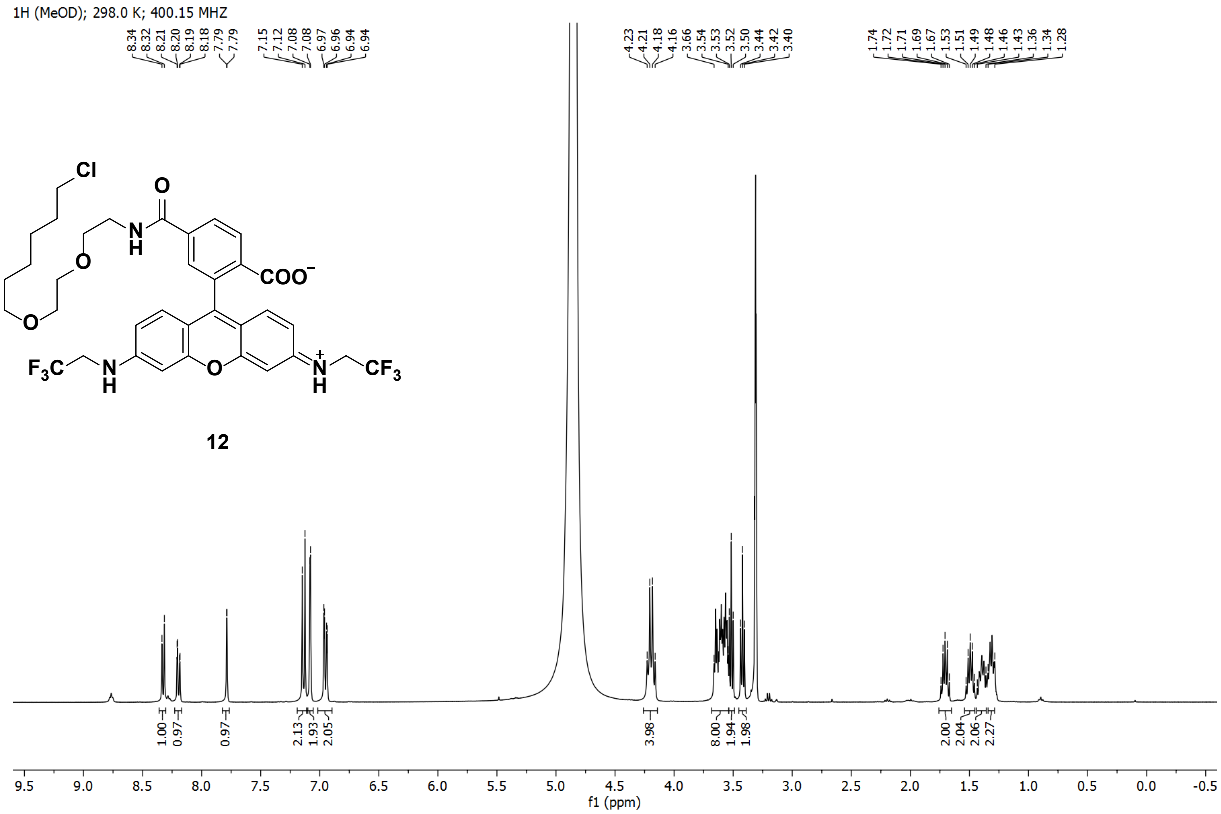
1H (MeOD); 298.0 K; 400.15 MHZ
8.34 8.32 8.21 8.20 8.19 8.18 7.79 7.79 7.15 7.12 7.08 7.08 6.97 6.96 6.94 6.94
4.23 4.21 4.18 4.16 3.66 3.54 3.53 3.52 3.50 3.44 3.42 3.40
1.74 1.72 1.71 1.69 1.67 1.53 1.51 1.49 1.48 1.46 1.43 1.36 1.34 1.28
1.00 0.97 0.97 2.13 1.93 2.05 3.98 8.00 1.94 1.98 2.00 2.04 2.06 2.27
f1 (ppm)
12

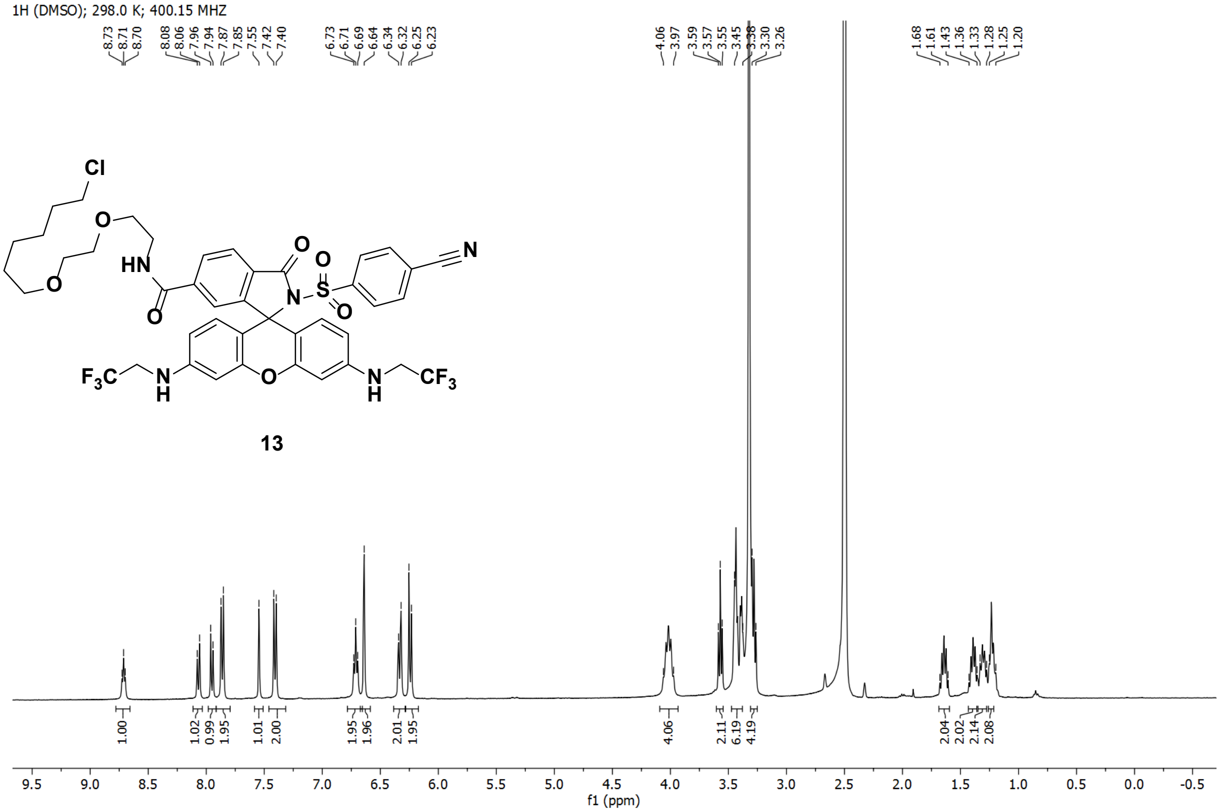
1H (DMSO); 298.0 K; 400.15 MHZ
8.73 8.71 8.70 8.08 8.06 7.96 7.94 7.87 7.85 7.55 7.42 7.40 6.73 6.71 6.69 6.64 6.34 6.32 6.25 6.23
4.06 3.97 3.59 3.57 3.55 3.45 3.38 3.30 3.26
1.68 1.61 1.43 1.36 1.33 1.28 1.25 1.20
1.00 1.02 0.99 1.95 1.01 2.00 1.95 1.96 2.01 1.95 4.06 2.11 6.19 4.19 2.04 2.02 2.14 2.08
f1 (ppm)
13

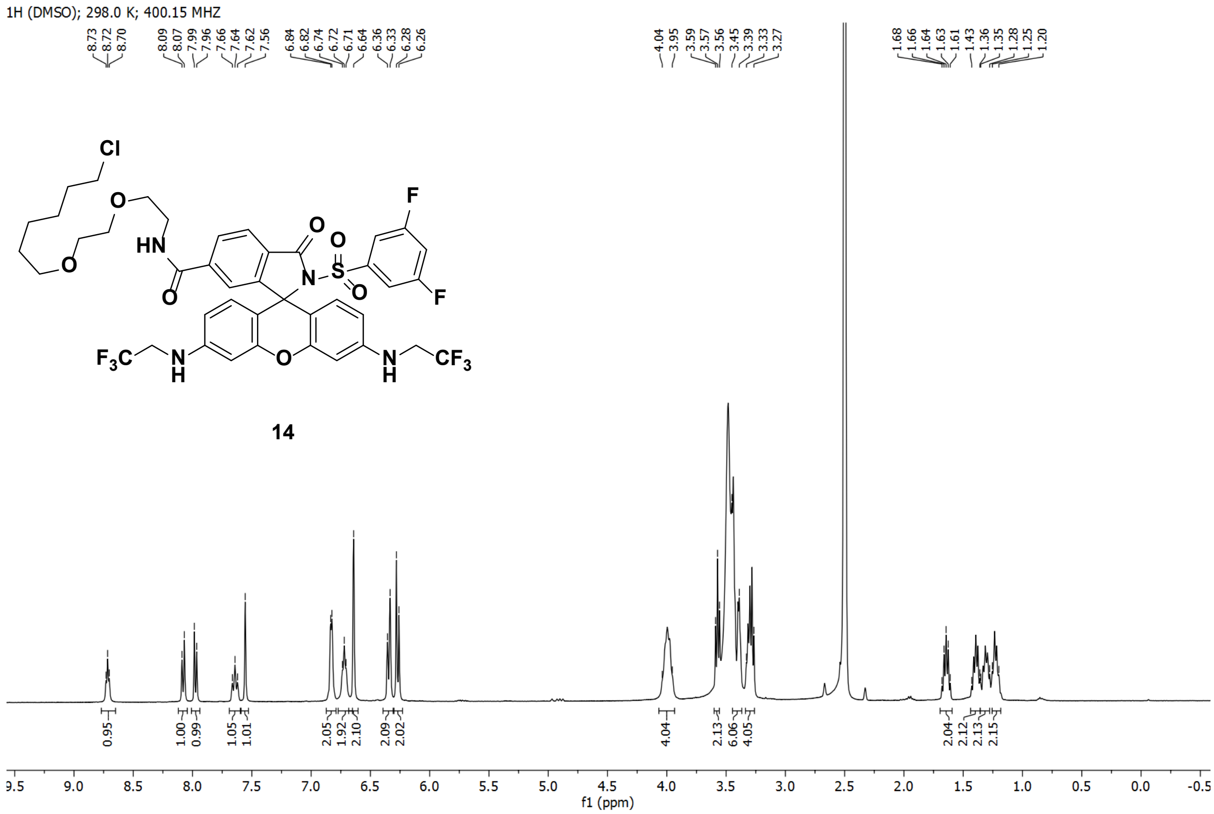
1H (DMSO); 298.0 K; 400.15 MHZ
14
f1 (ppm)

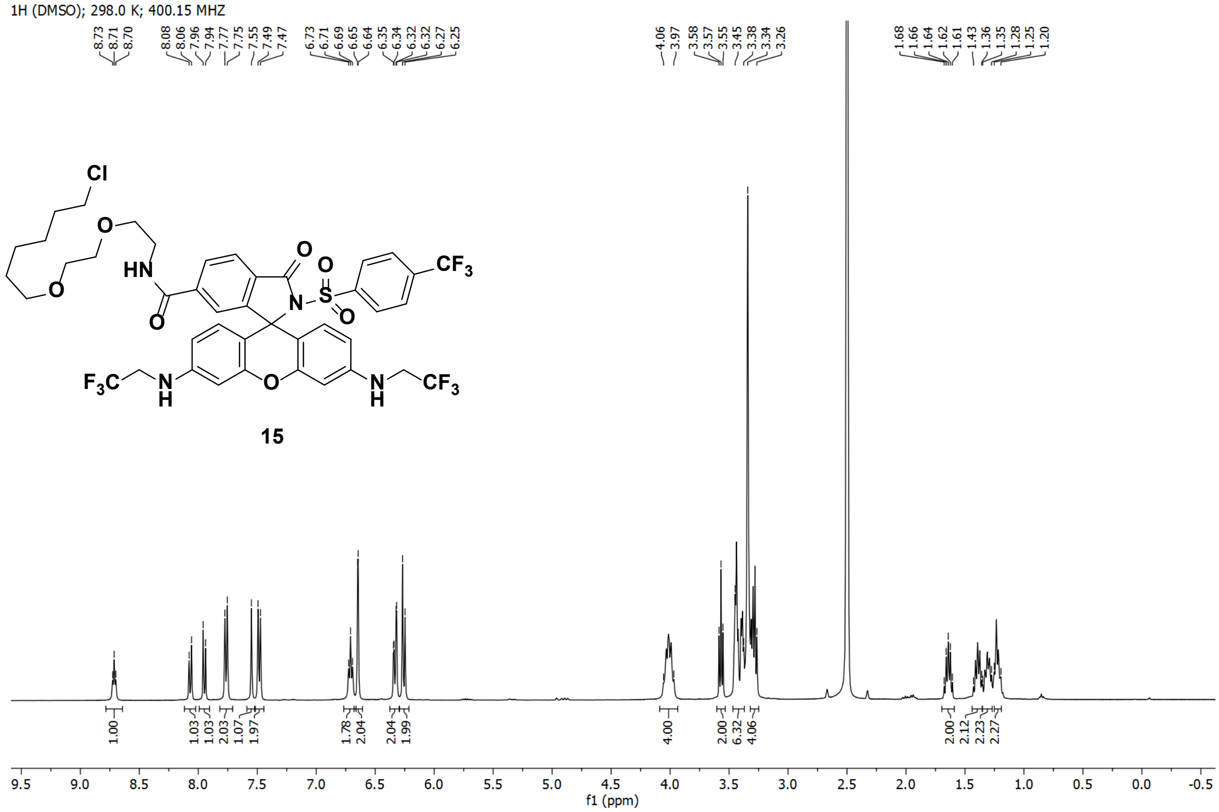
1H (DMSO); 298.0 K; 400.15 MHZ
15
f1 (ppm)

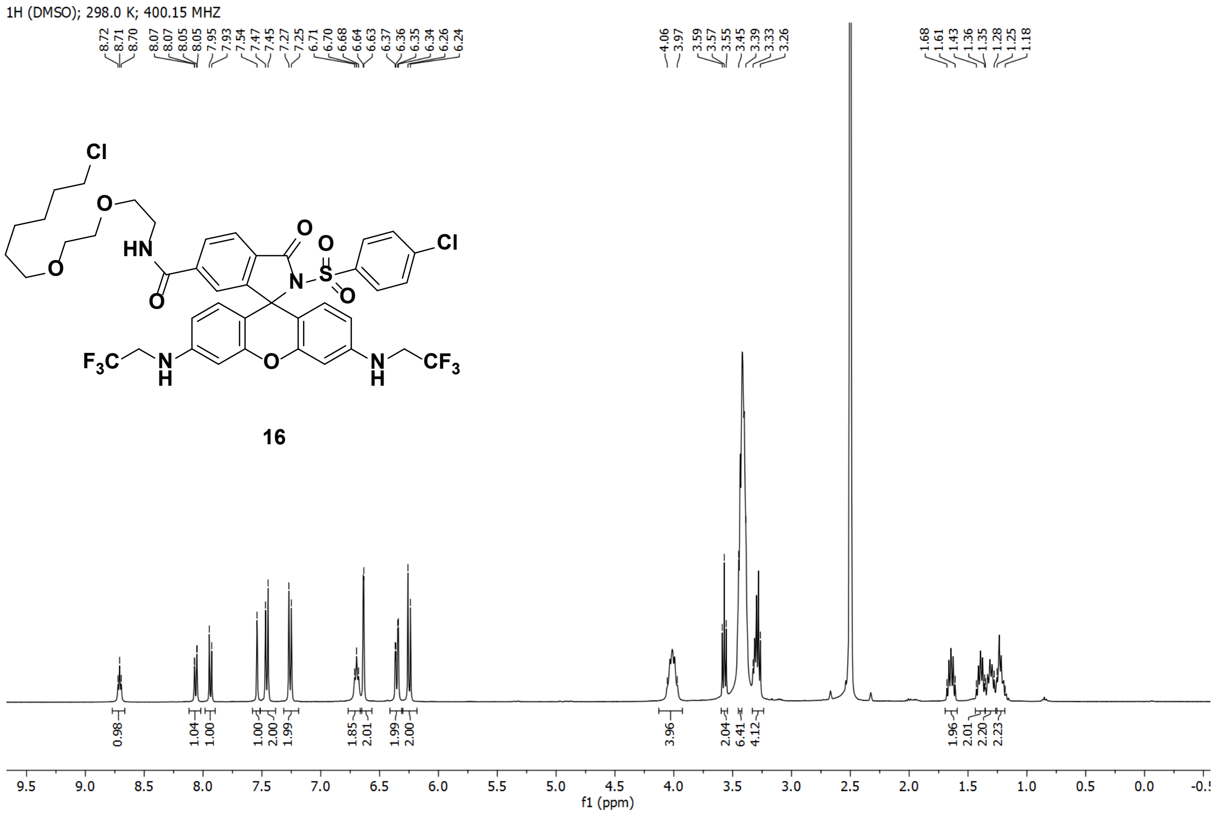
1H (DMSO); 298.0 K; 400.15 MHZ
16
f1 (ppm)

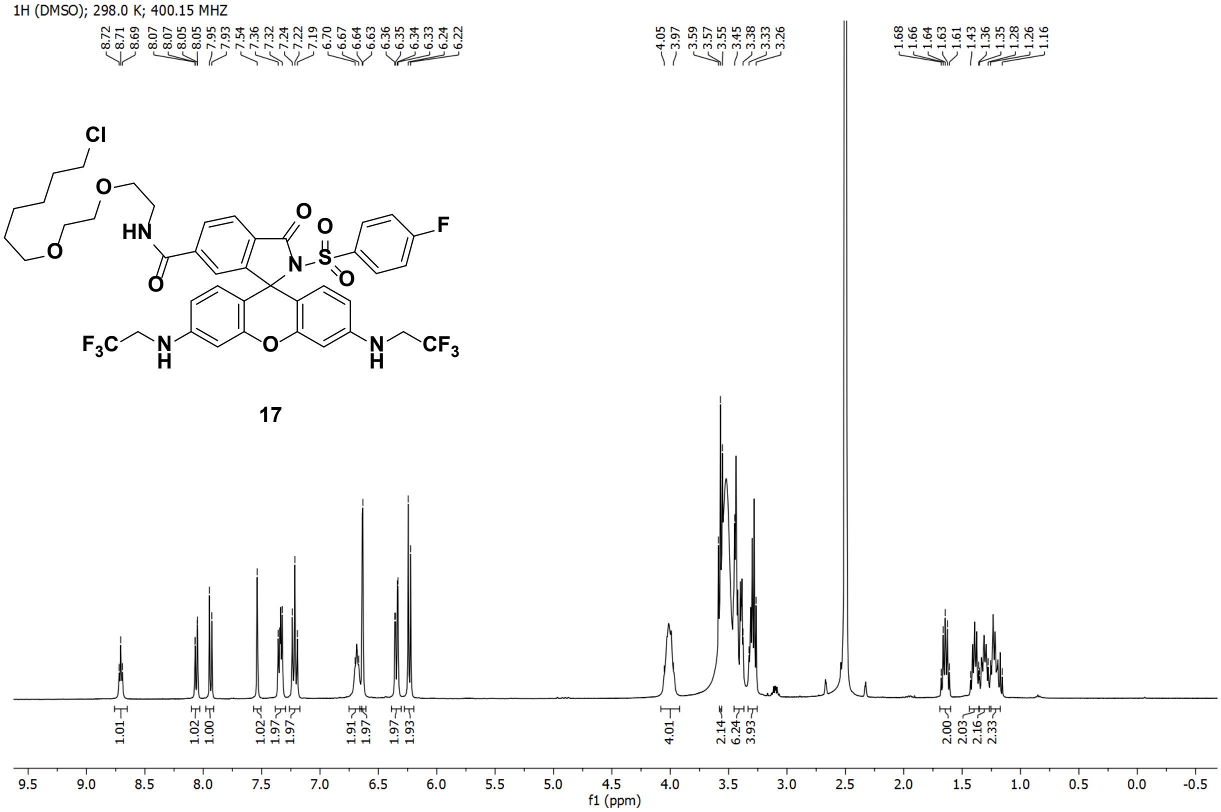
1H (DMSO); 298.0 K; 400.15 MHZ
17
f1 (ppm)

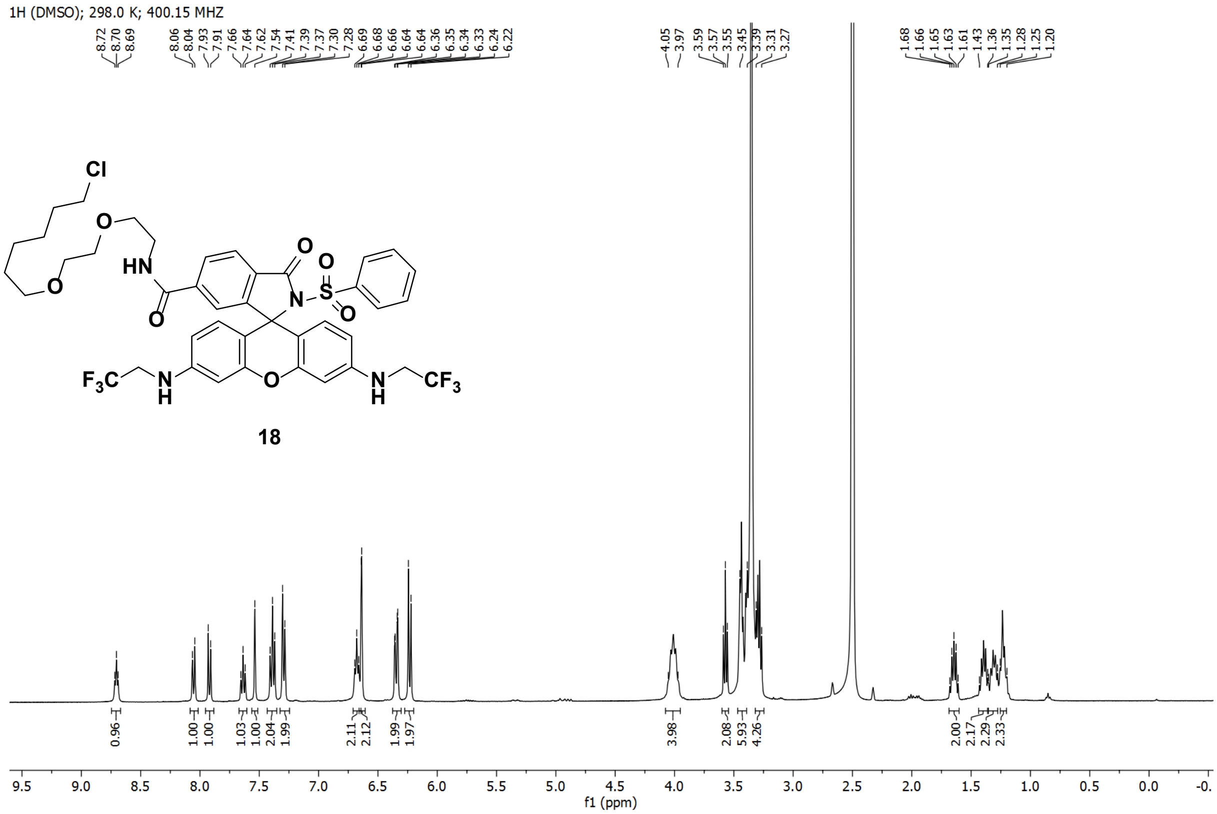
1H (DMSO); 298.0 K; 400.15 MHZ
8.72 8.70 8.69 8.06 8.04 7.93 7.91 7.66 7.64 7.62 7.54 7.41 7.39 7.37 7.30 7.28 6.69 6.68 6.66 6.64 6.64 6.36 6.35 6.34 6.33 6.24 6.22
4.05 3.97 3.59 3.57 3.55 3.45 3.39 3.31 3.27
1.68 1.66 1.65 1.63 1.61 1.43 1.36 1.35 1.28 1.25 1.20
Cl
O
O
HN
O
O
S
O
N
O
$F_3C$
N
H
O
N
H
$CF_3$
18
0.96 1.00 1.00 1.03 1.00 2.04 1.99 2.11 2.12 1.99 1.97 3.98 2.08 5.93 4.26 2.00 2.17 2.29 2.33
9.5 9.0 8.5 8.0 7.5 7.0 6.5 6.0 5.5 5.0 4.5 4.0 3.5 3.0 2.5 2.0 1.5 1.0 0.5 0.0
f1 (ppm)

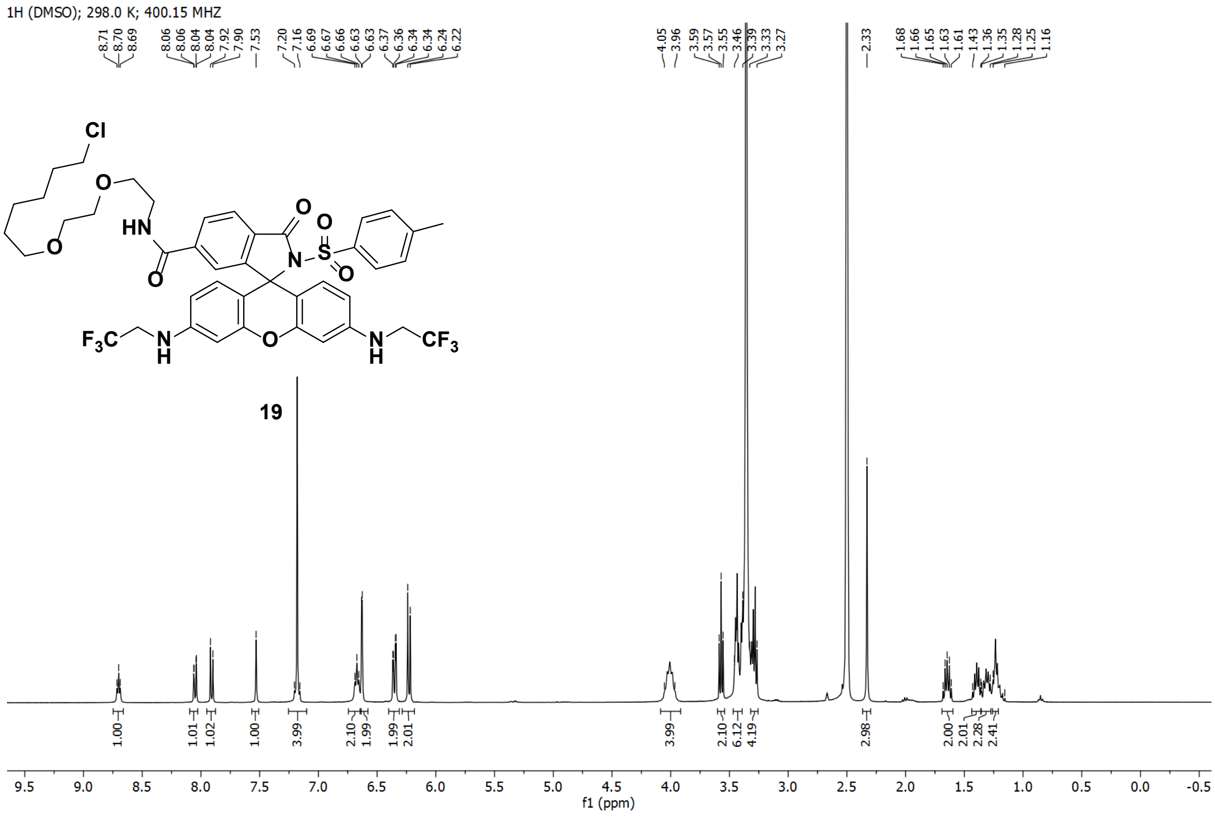
1H (DMSO); 298.0 K; 400.15 MHZ
8.71 8.70 8.69 8.06 8.06 8.04 8.04 7.92 7.90 7.53 7.20 7.16 6.69 6.67 6.66 6.63 6.63 6.37 6.36 6.34 6.34 6.24 6.22
4.05 3.96 3.59 3.57 3.55 3.46 3.39 3.33 3.27 2.33
1.68 1.66 1.65 1.63 1.61 1.43 1.36 1.35 1.28 1.25 1.16
Cl
O
O
HN
O
O
S
O
N
O
$F_3C$
N
H
O
N
H
$CF_3$
19
1.00 1.01 1.02 1.00 3.99 2.10 1.99 1.99 2.01 3.99 2.10 6.12 4.19 2.98 2.00 2.01 2.28 2.41
9.5 9.0 8.5 8.0 7.5 7.0 6.5 6.0 5.5 5.0 4.5 4.0 3.5 3.0 2.5 2.0 1.5 1.0 0.5 0.0 -0.5
f1 (ppm)

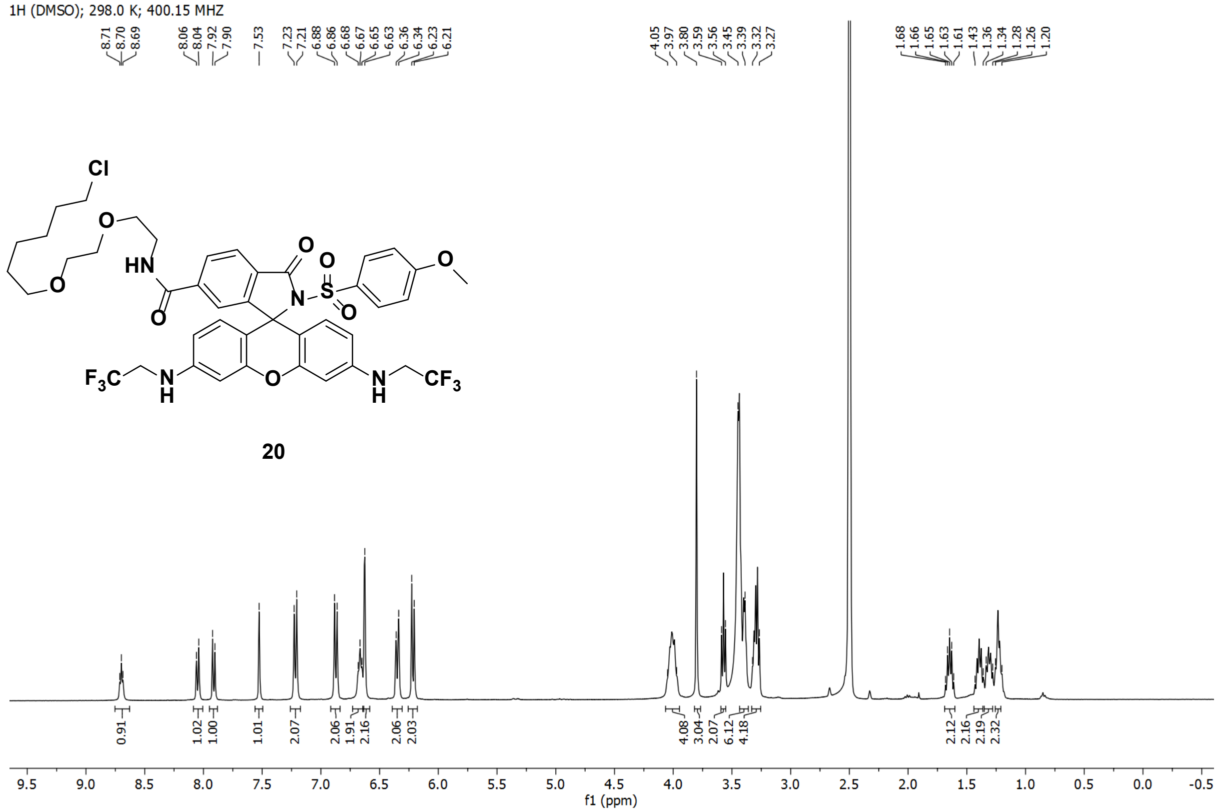
1H (DMSO); 298.0 K; 400.15 MHZ
20
f1 (ppm)

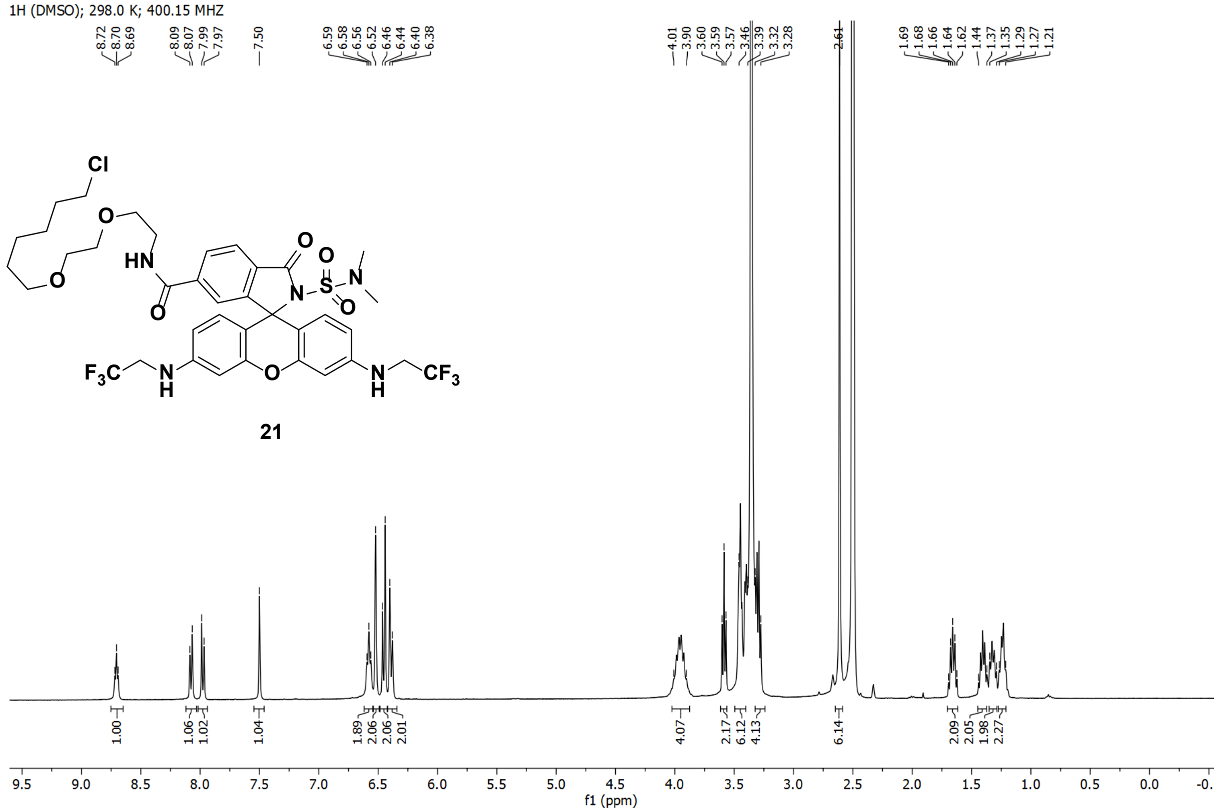
1H (DMSO); 298.0 K; 400.15 MHZ
21
f1 (ppm)

1H (MeOD); 298.0 K; 400.15 MHZ

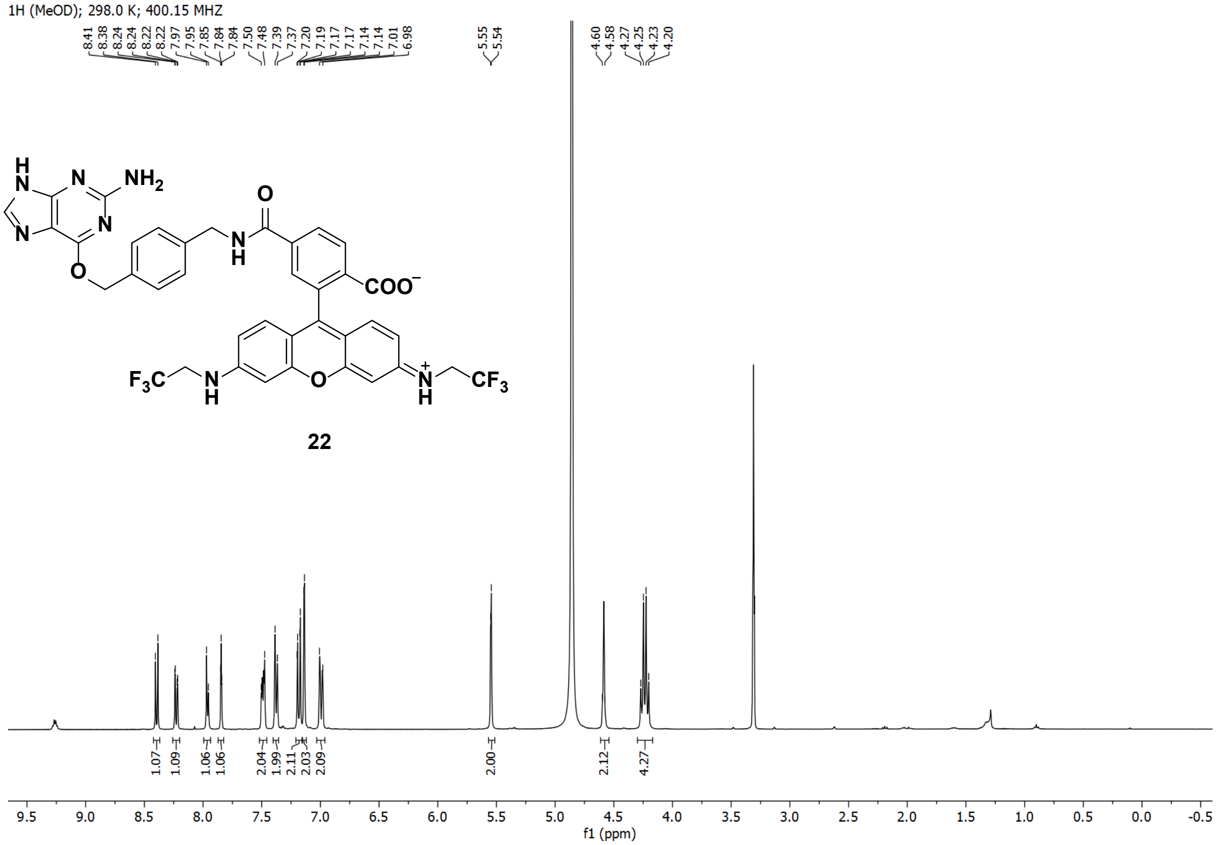

1H (MeOD); 298.0 K; 400.15 MHZ

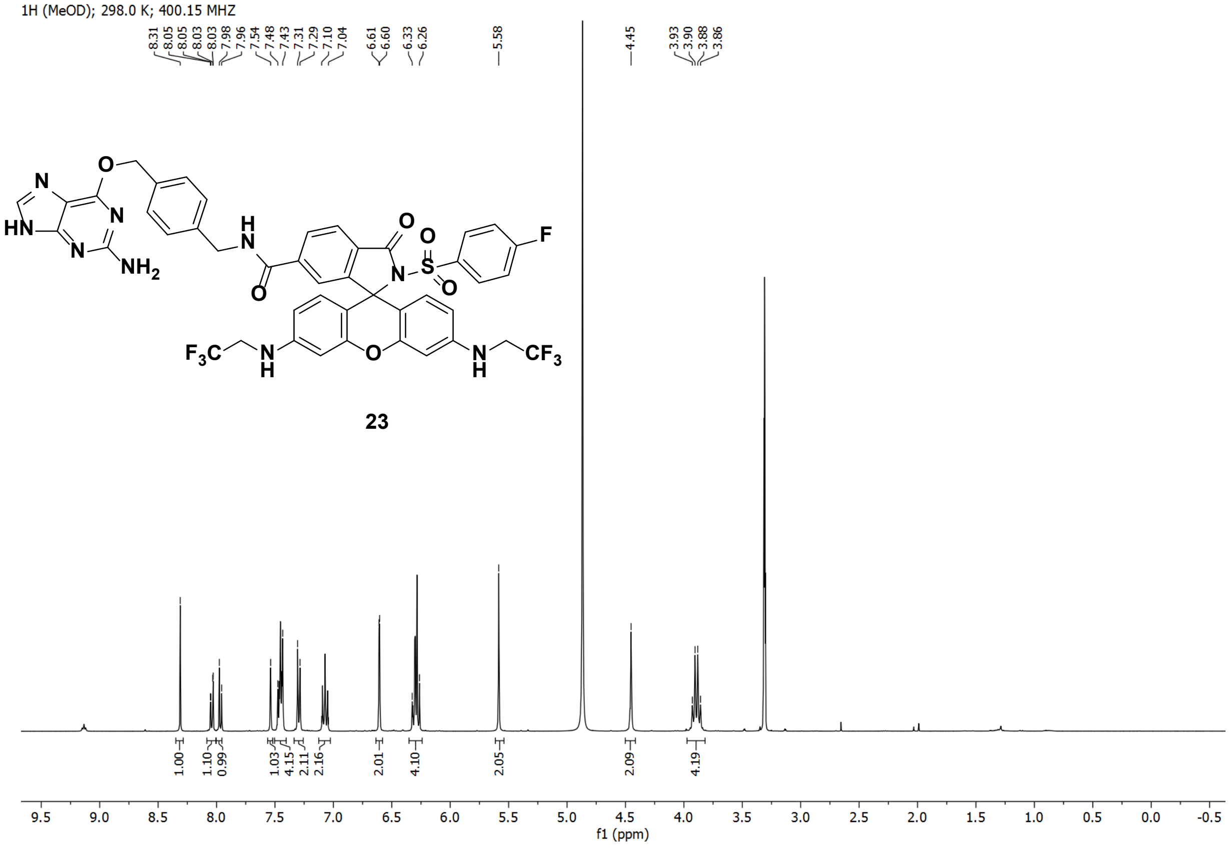

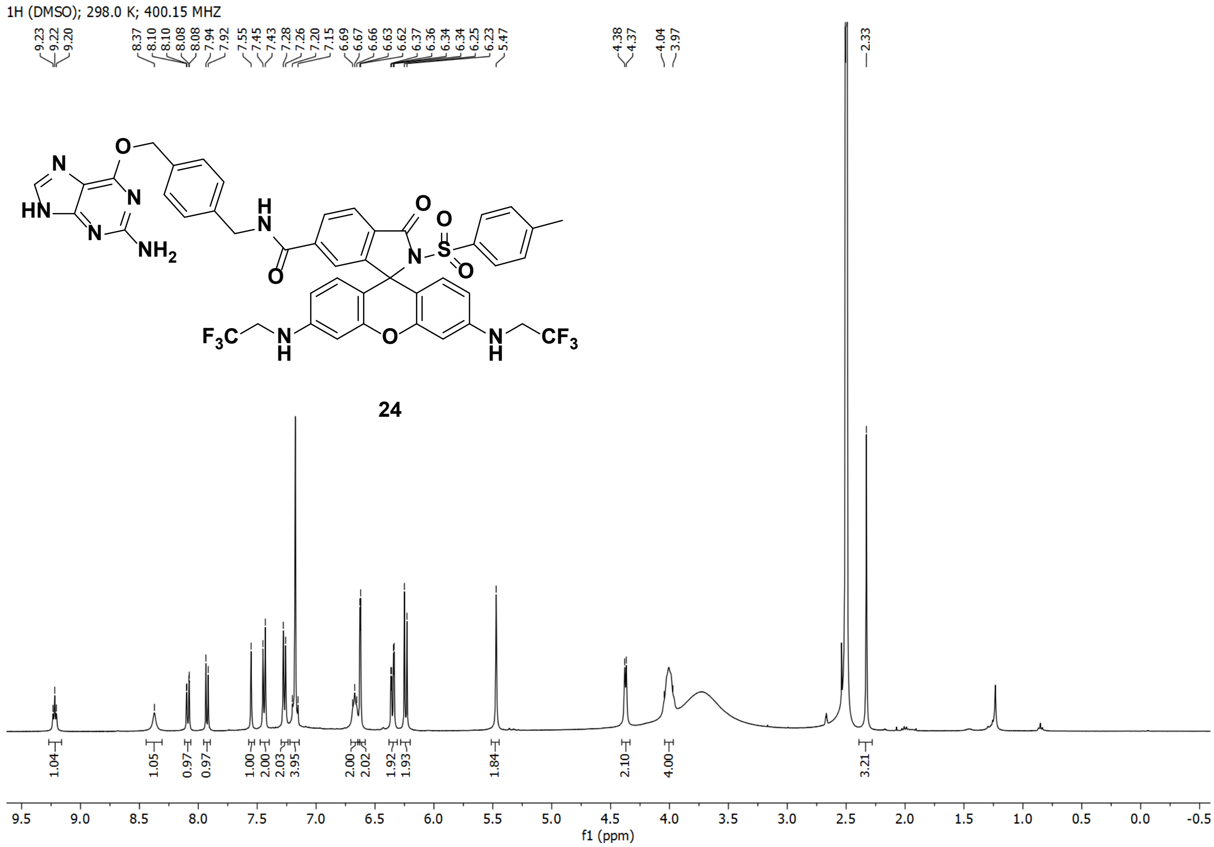
1H (DMSO); 298.0 K; 400.15 MHZ
24
f1 (ppm)

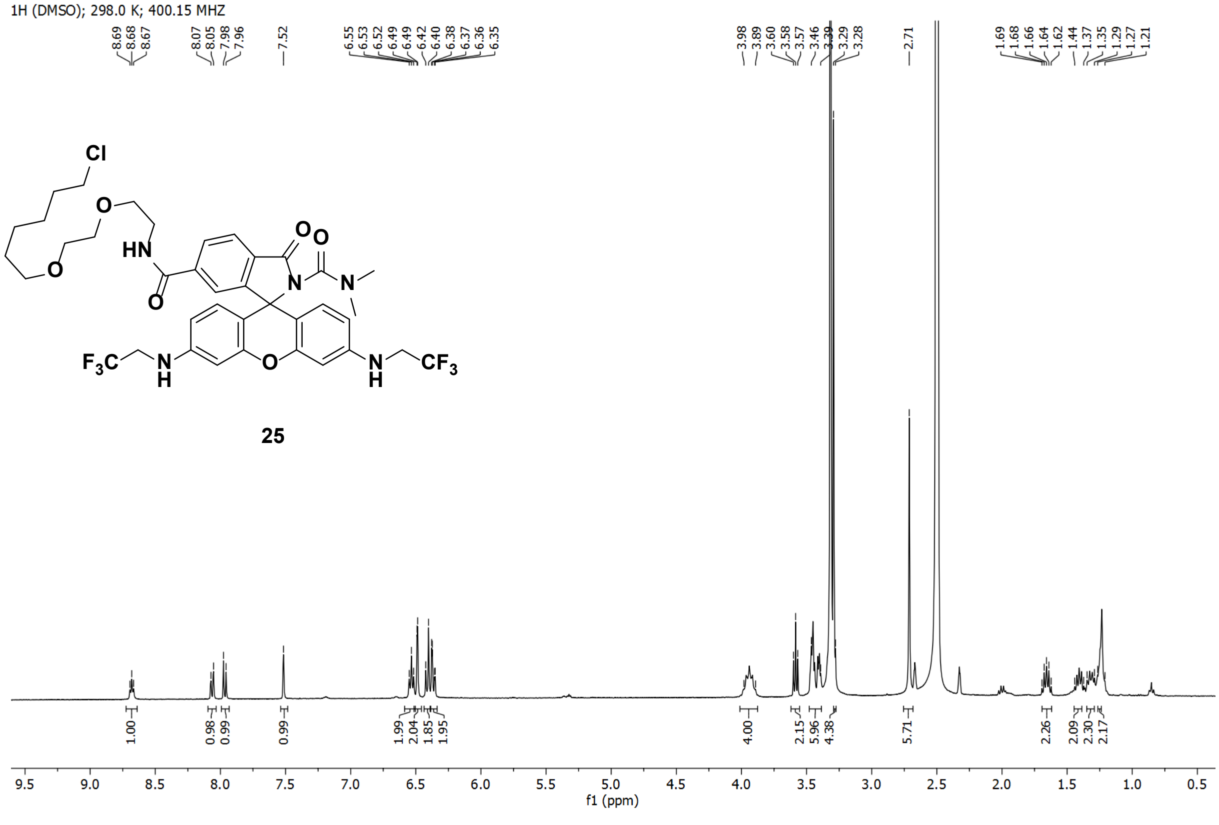
1H (DMSO); 298.0 K; 400.15 MHZ
25
f1 (ppm)

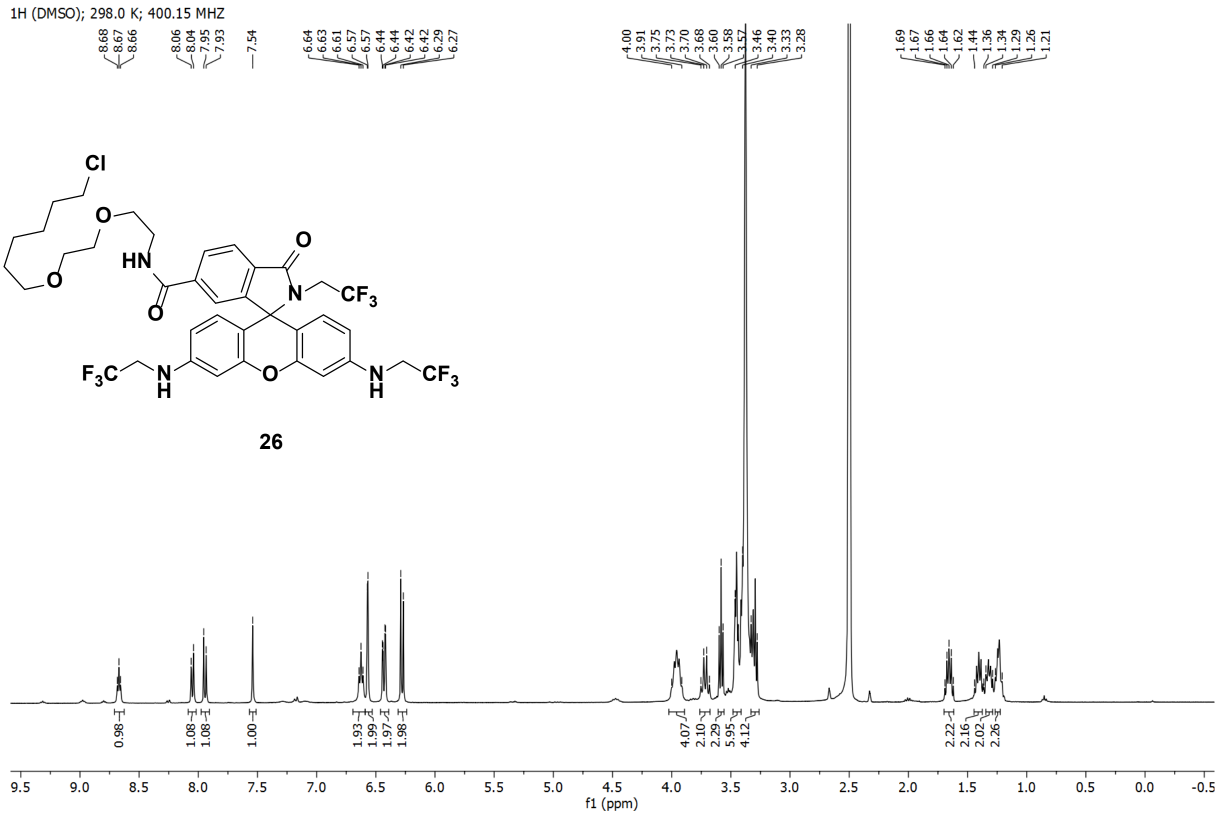
1H (DMSO); 298.0 K; 400.15 MHZ
26
f1 (ppm)

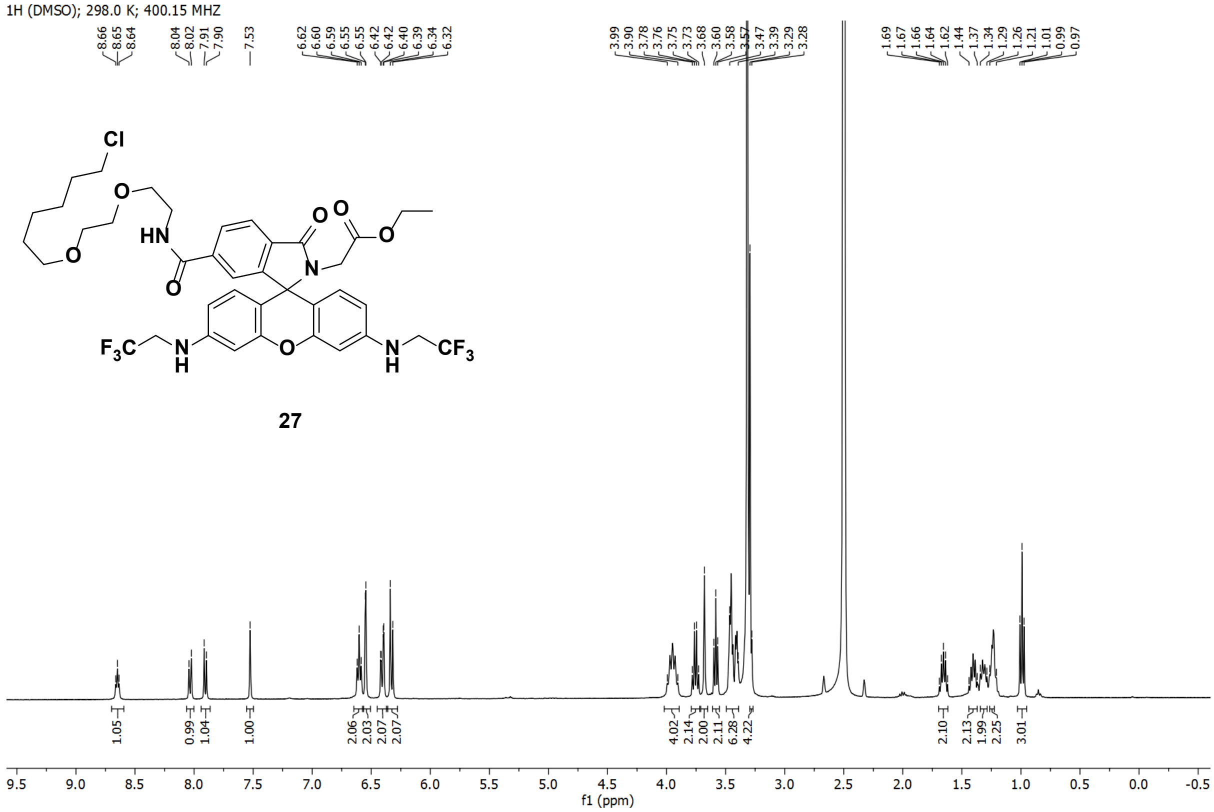
1H (DMSO); 298.0 K; 400.15 MHZ
27
f1 (ppm)

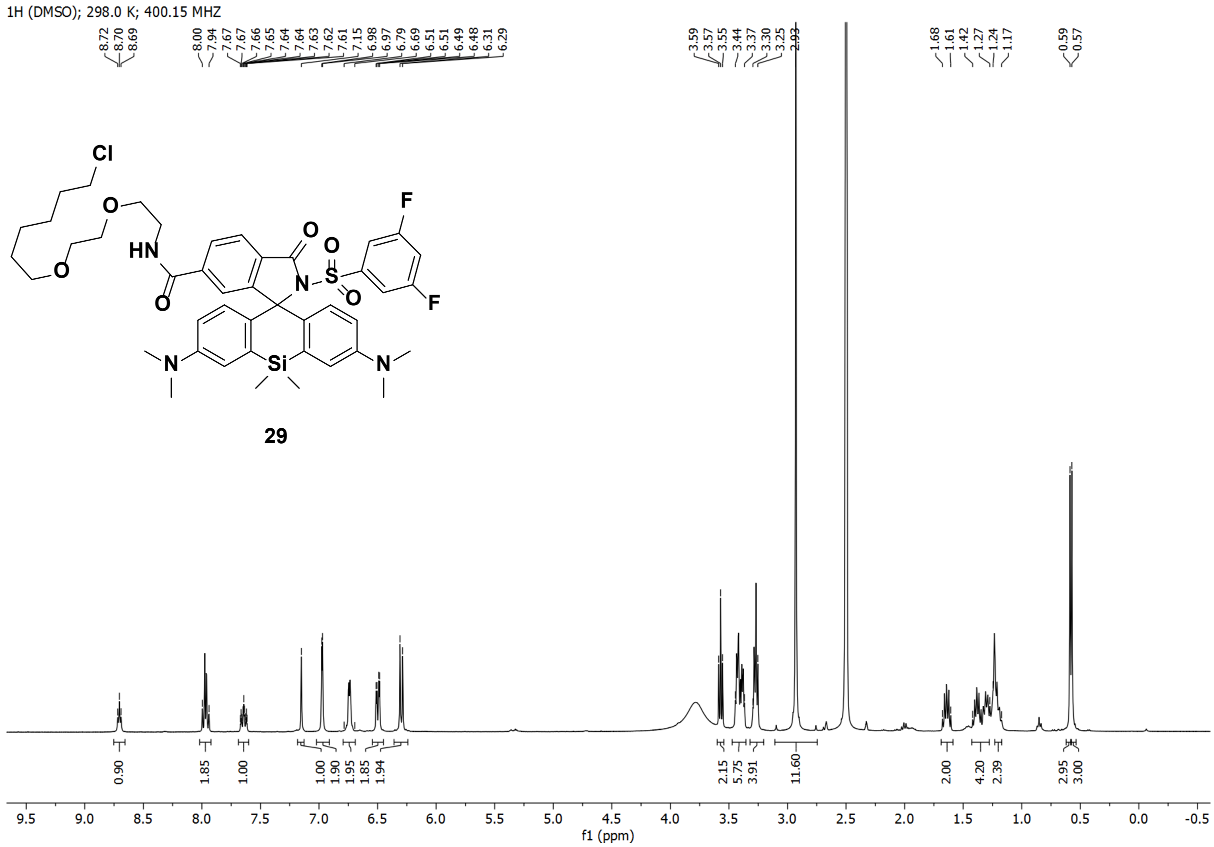
1H (DMSO); 298.0 K; 400.15 MHZ
29
f1 (ppm)

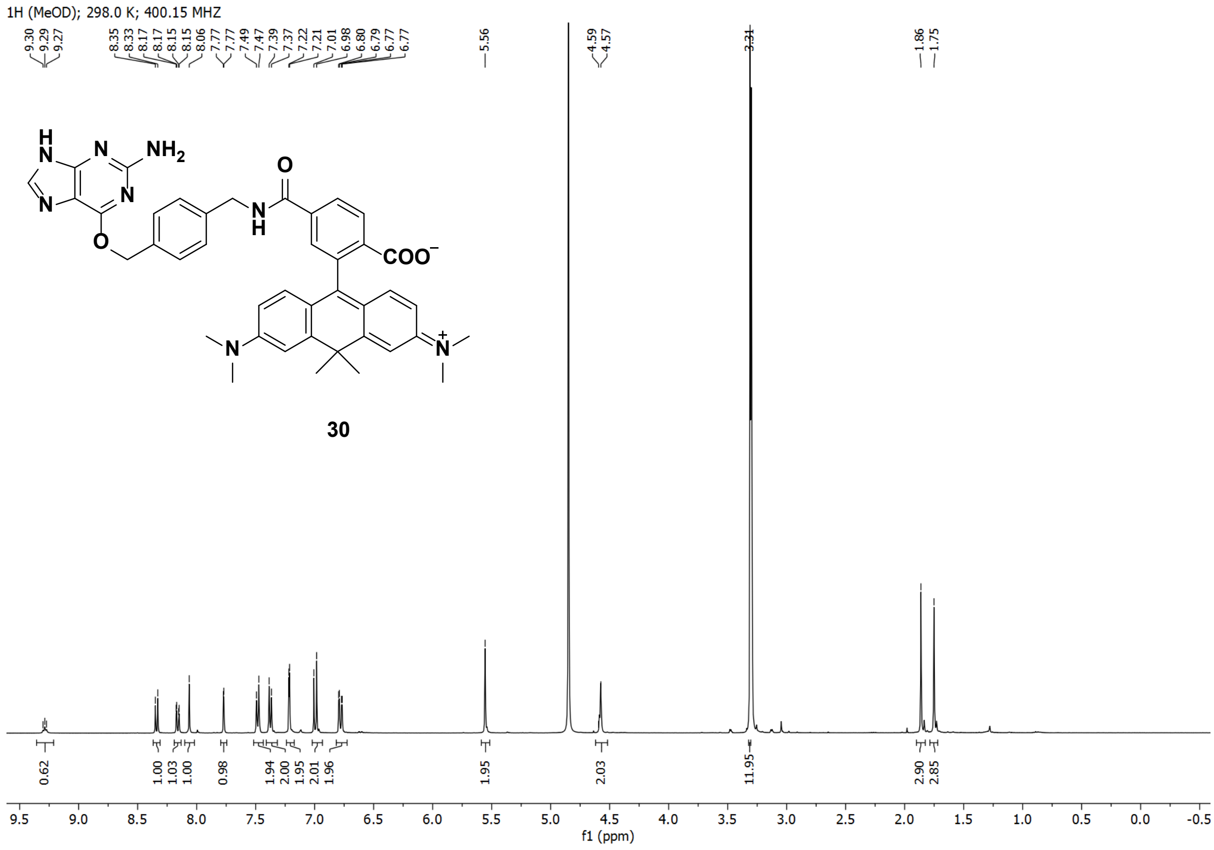
1H (MeOD); 298.0 K; 400.15 MHZ
30
f1 (ppm)

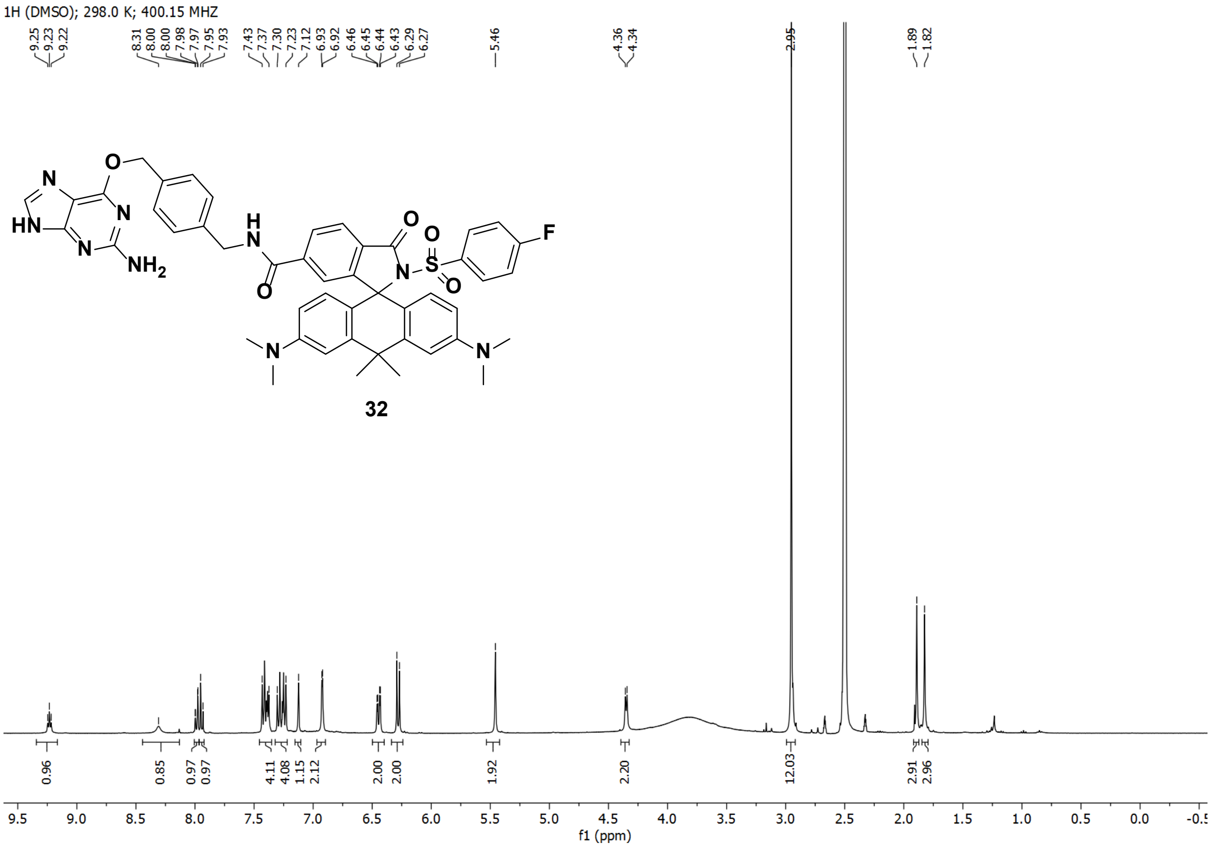
1H (DMSO); 298.0 K; 400.15 MHZ
9.25 9.23 9.22 8.31 8.00 8.00 7.98 7.97 7.95 7.93 7.43 7.37 7.30 7.23 7.12 6.93 6.92 6.46 6.45 6.44 6.43 6.29 6.27 5.46 4.36 4.34 2.95 1.89 1.82
0.96 0.85 0.97 0.97 4.11 4.08 1.15 2.12 2.00 2.00 1.92 2.20 12.03 2.91 2.96
f1 (ppm)

32

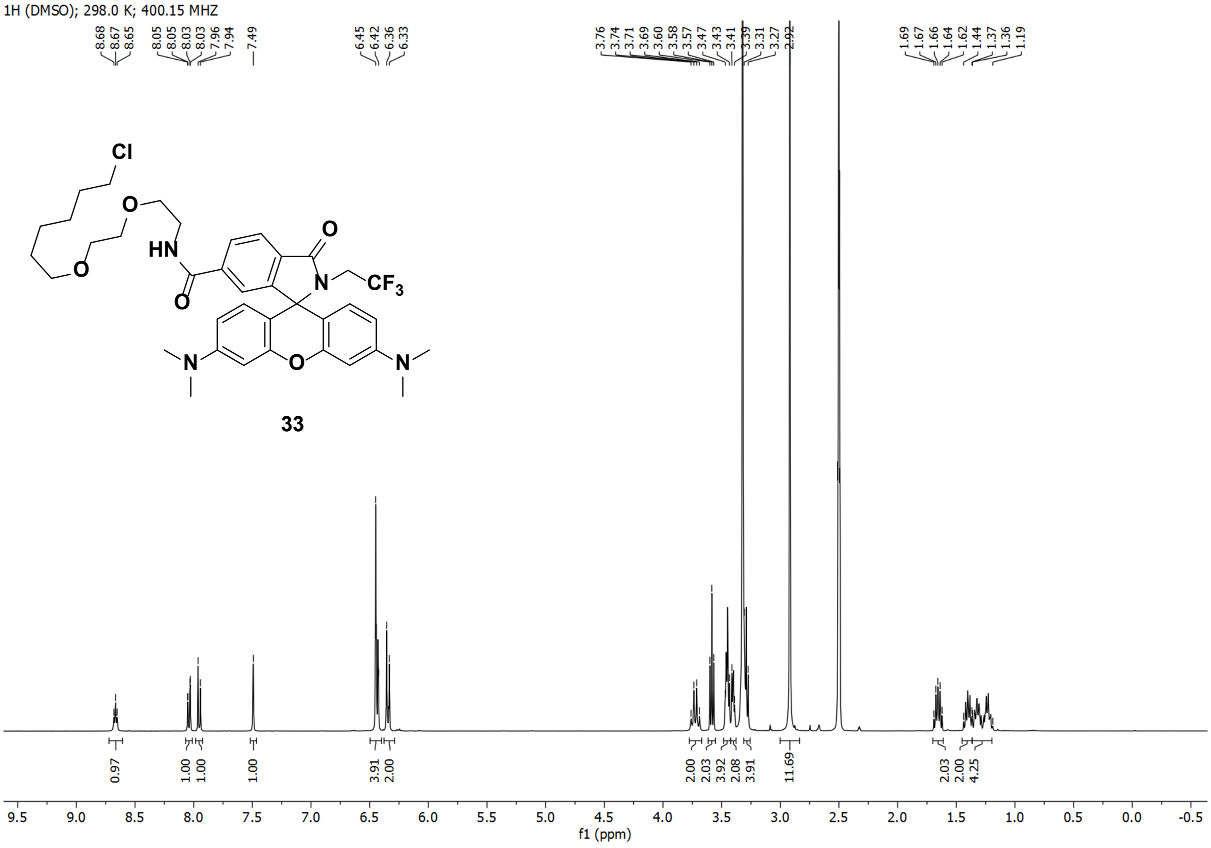
1H (DMSO); 298.0 K; 400.15 MHZ
8.68 8.67 8.65 8.05 8.05 8.03 8.03 7.96 7.94 7.49 6.45 6.42 6.36 6.33 3.76 3.74 3.71 3.69 3.60 3.58 3.57 3.47 3.43 3.41 3.39 3.31 3.27 2.92 1.69 1.67 1.66 1.64 1.62 1.44 1.37 1.36 1.19
0.97 1.00 1.00 1.00 3.91 2.00 2.00 2.03 3.92 2.08 3.91 11.69 2.03 2.00 4.25
f1 (ppm)

33

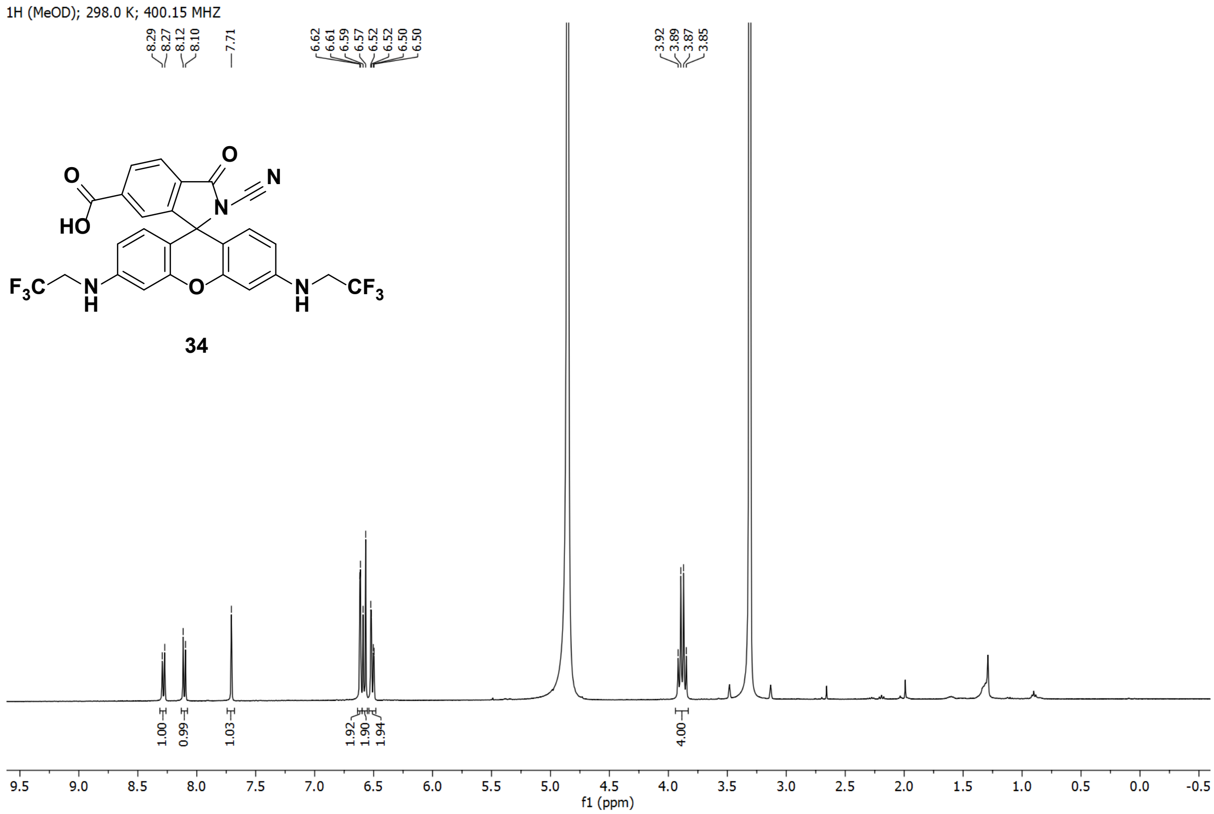

1H (MeOD); 298.0 K; 400.15 MHZ
8.29
8.27
8.12
8.10
7.71
6.62
6.61
6.59
6.57
6.52
6.52
6.50
6.50
3.92
3.89
3.87
3.85
1.00
0.99
1.03
1.92
1.90
1.94
4.00
34
f1 (ppm)


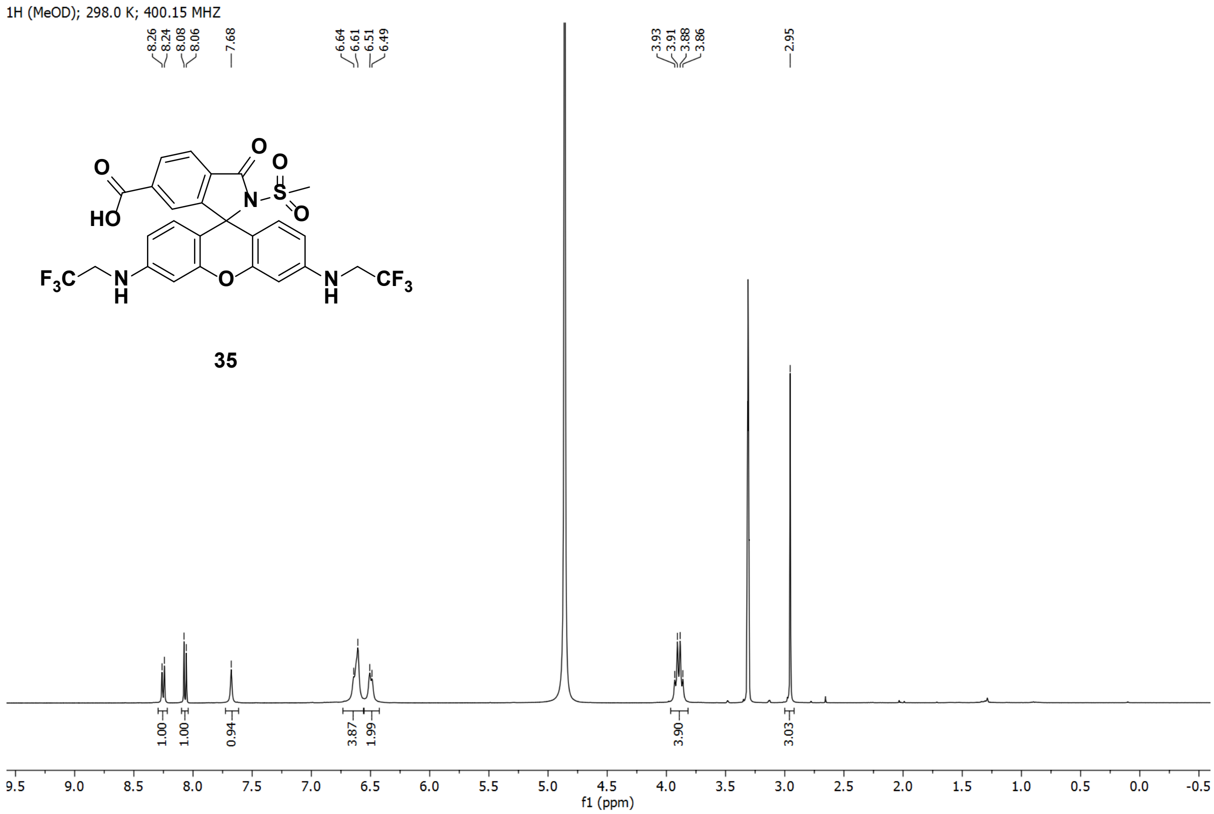

1H (MeOD); 298.0 K; 400.15 MHZ
8.26
8.24
8.08
8.06
7.68
6.64
6.61
6.51
6.49
3.93
3.91
3.88
3.86
2.95
1.00
1.00
0.94
3.87
1.99
3.90
3.03
35
f1 (ppm)

13C (DMSO); 298.0 K; 100.63 MHZ

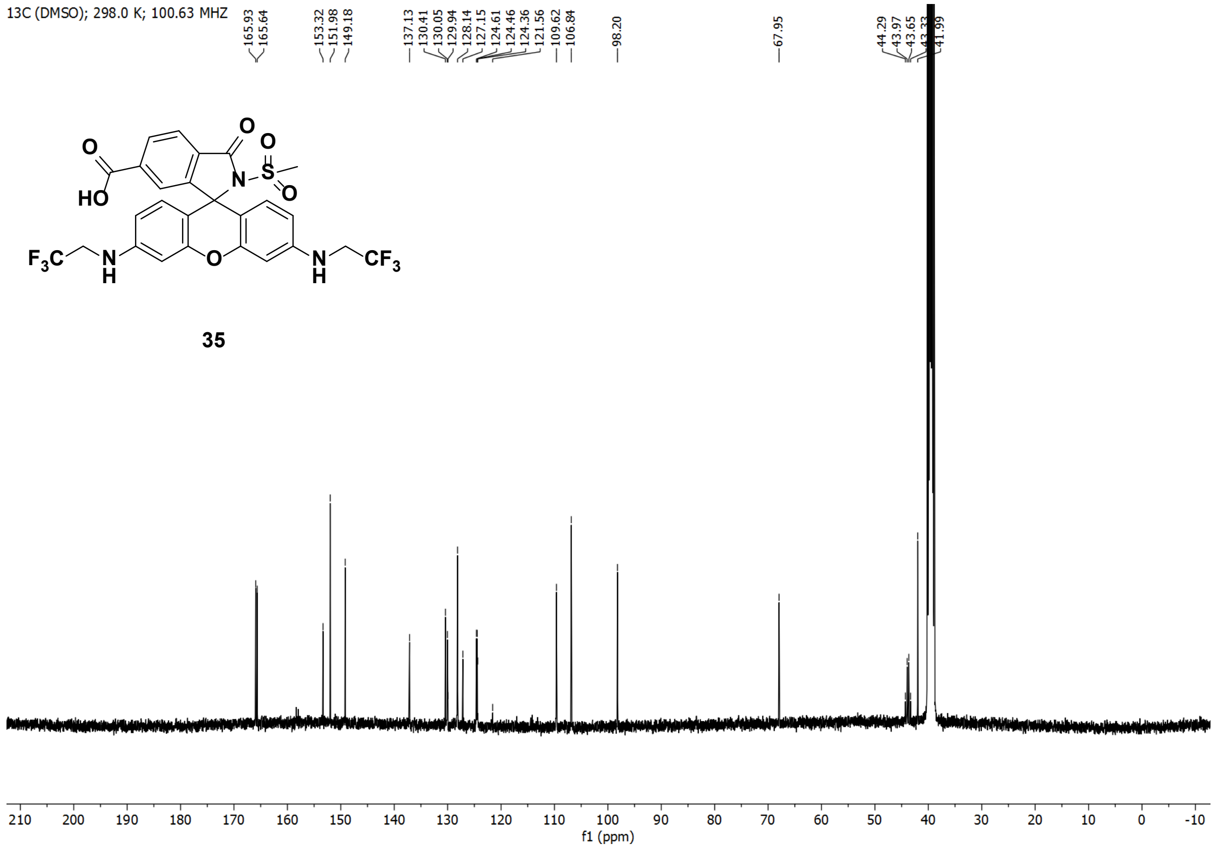


35

1H (DMSO); 298.0 K; 400.15 MHZ

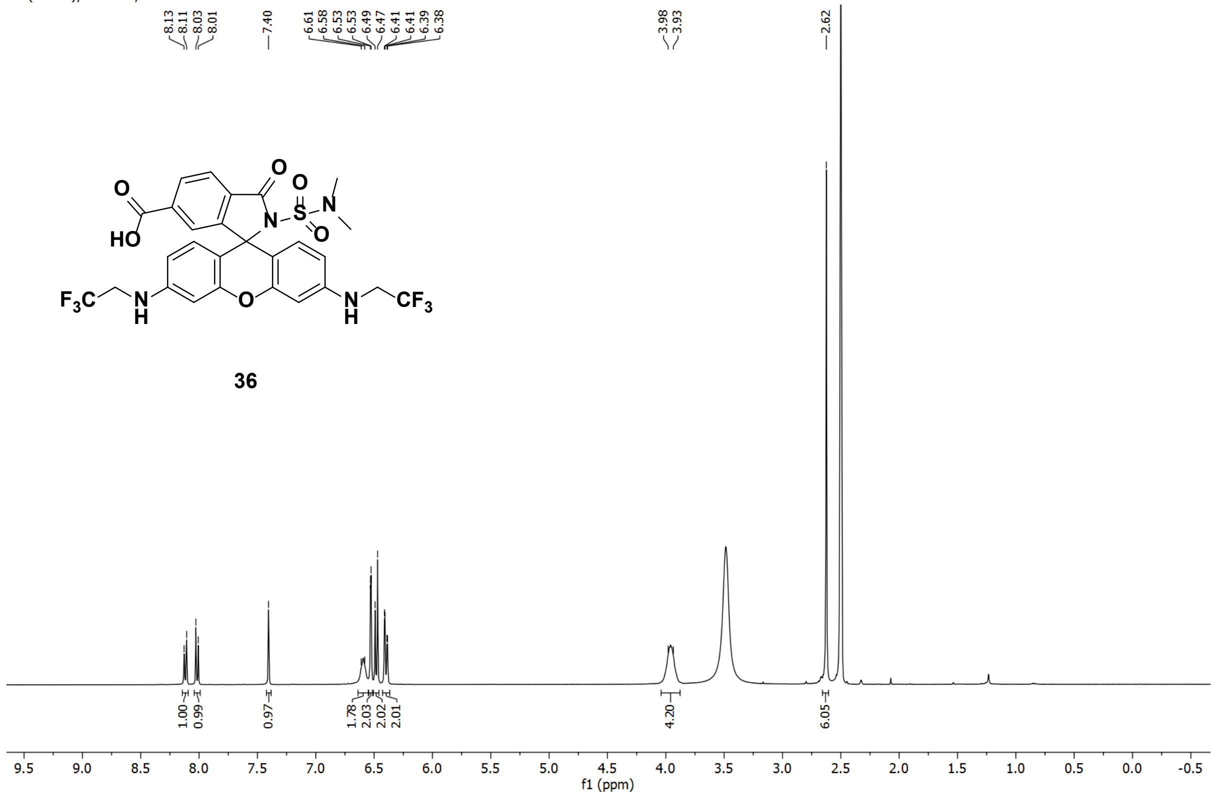


36

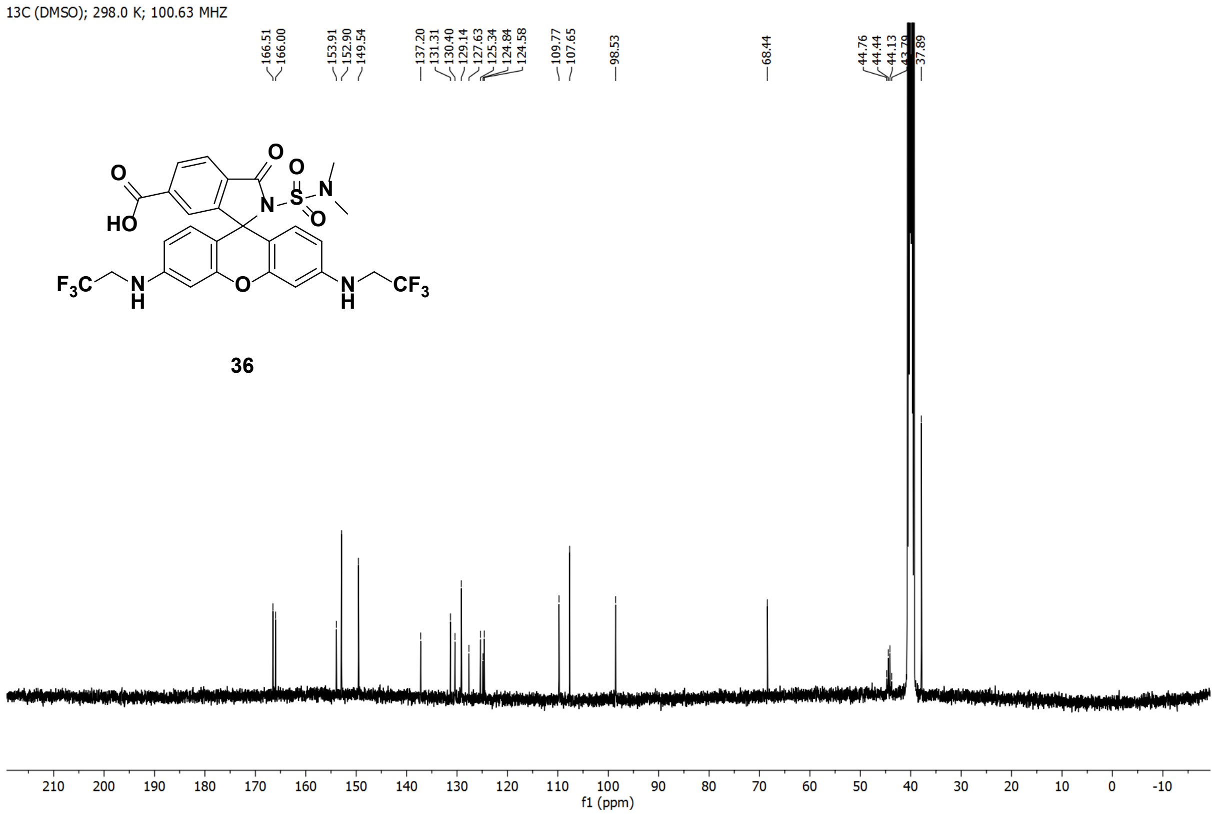
13C (DMSO); 298.0 K; 100.63 MHZ
166.51
166.00
153.91
152.90
149.54
137.20
131.31
130.40
129.14
127.63
125.34
124.84
124.58
109.77
107.65
98.53
68.44
44.76
44.44
44.13
37.89
O
HO
N
S
O
O
N
F3C
N
H
O
N
H
CF3
36
210 200 190 180 170 160 150 140 130 120 110 100 90 80 70 60 50 40 30 20 10 0 -10
f1 (ppm)

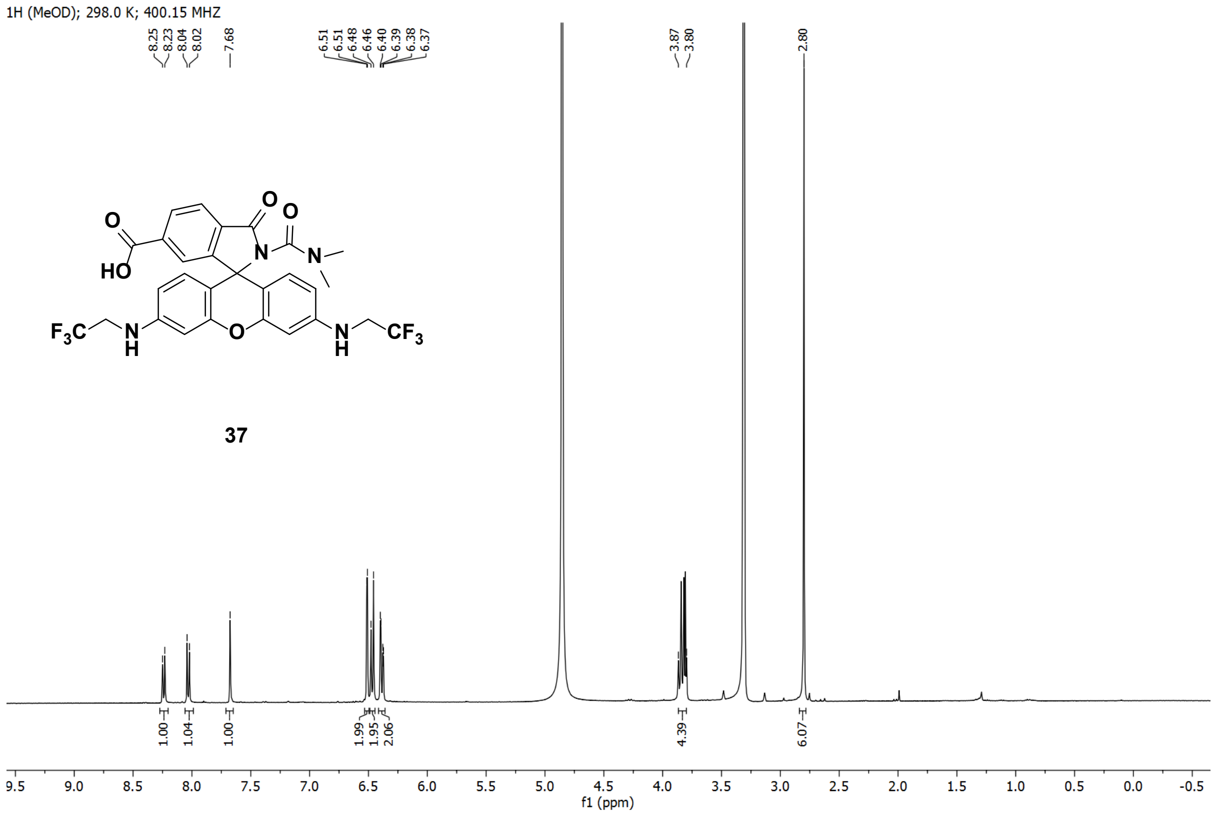
1H (MeOD); 298.0 K; 400.15 MHZ
8.25
8.23
8.04
8.02
7.68
6.51
6.51
6.48
6.46
6.40
6.39
6.38
6.37
3.87
3.80
2.80
O
O
O
HO
N
N
F3C
N
H
O
N
H
CF3
37
1.00
1.04
1.00
1.99
1.95
2.06
4.39
6.07
9.5 9.0 8.5 8.0 7.5 7.0 6.5 6.0 5.5 5.0 4.5 4.0 3.5 3.0 2.5 2.0 1.5 1.0 0.5 0.0 -0.5
f1 (ppm)

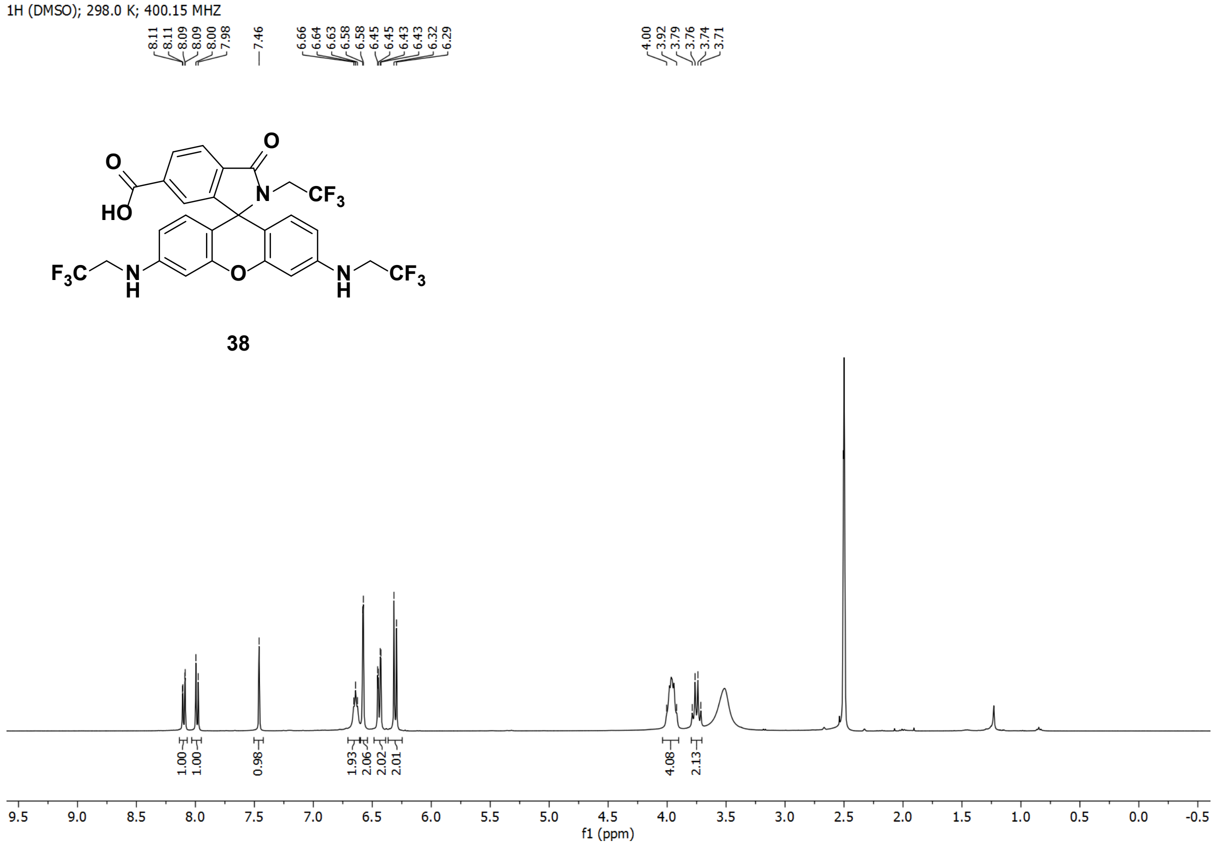
1H (DMSO); 298.0 K; 400.15 MHZ
8.11 8.11 8.09 8.09 8.00 7.98
7.46
6.66 6.64 6.63 6.58 6.58 6.45 6.45 6.43 6.43 6.32 6.29
4.00 3.92 3.79 3.76 3.74 3.71
38
1.00 1.00 0.98 1.93 2.06 2.02 2.01 4.08 2.13
f1 (ppm)

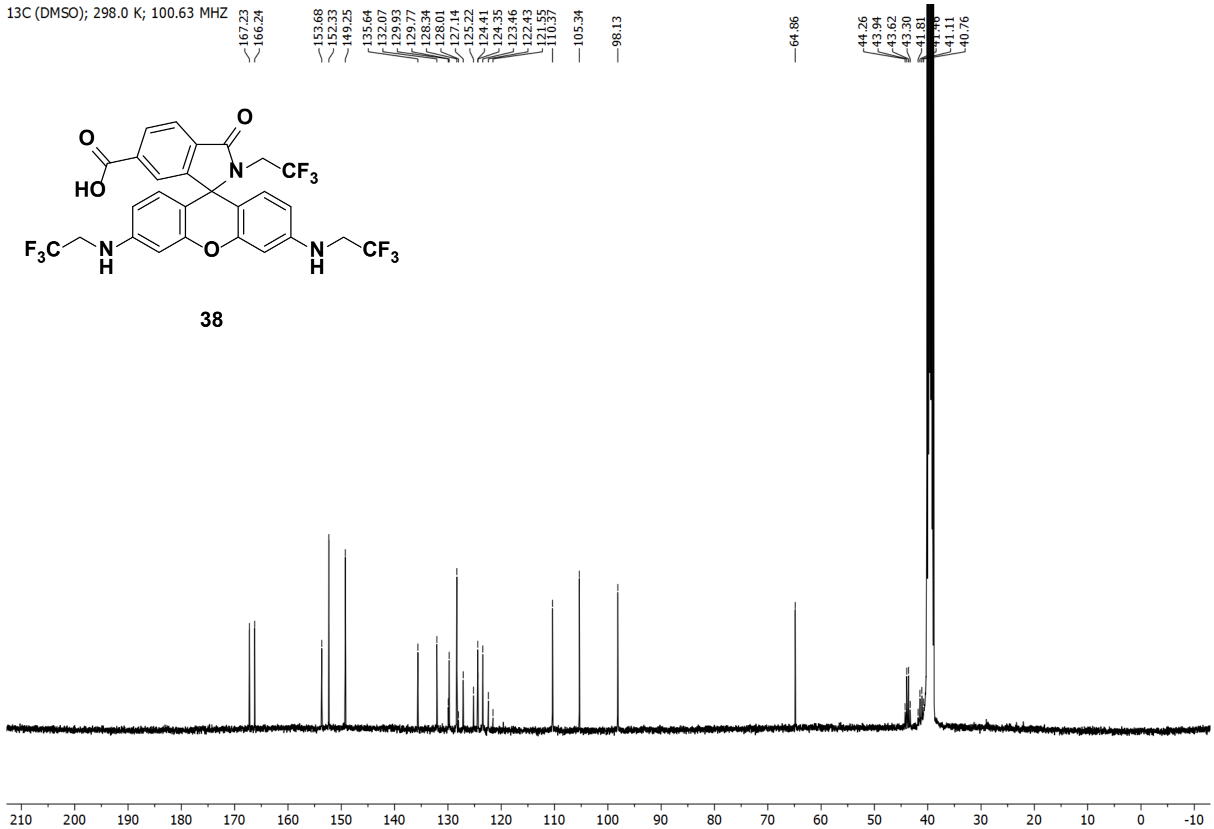
13C (DMSO); 298.0 K; 100.63 MHZ
167.23 166.24
153.68 152.33 149.25
135.64 132.07 129.93 129.77 128.34 128.01 127.14 125.22 124.41 124.35 123.46 122.43 121.55 110.37
105.34
98.13
64.86
44.26 43.94 43.62 43.30 41.81 41.46 41.11 40.76
38
f1 (ppm)

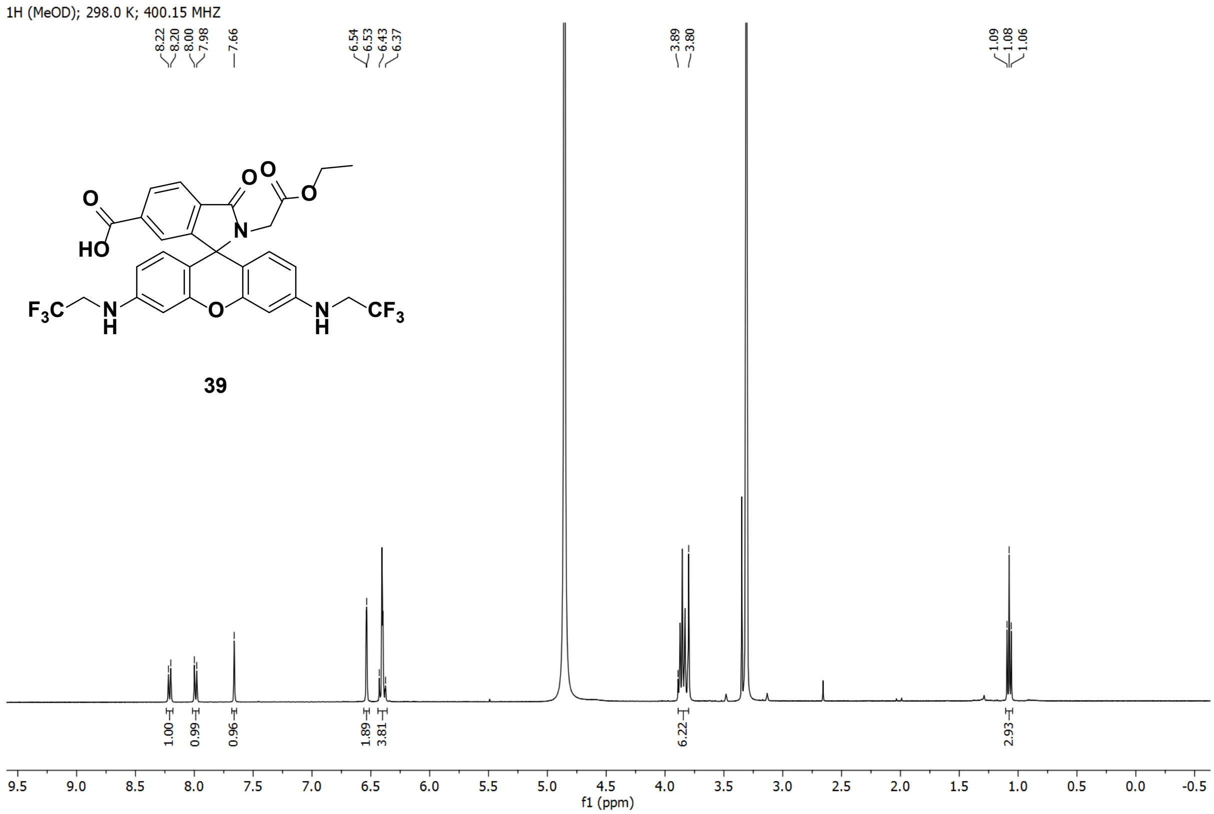
1H (MeOD); 298.0 K; 400.15 MHZ
8.22
8.20
8.00
7.98
7.66
6.54
6.53
6.43
6.37
3.89
3.80
1.09
1.08
1.06
39
1.00
0.99
0.96
1.89
3.81
6.22
2.93
f1 (ppm)

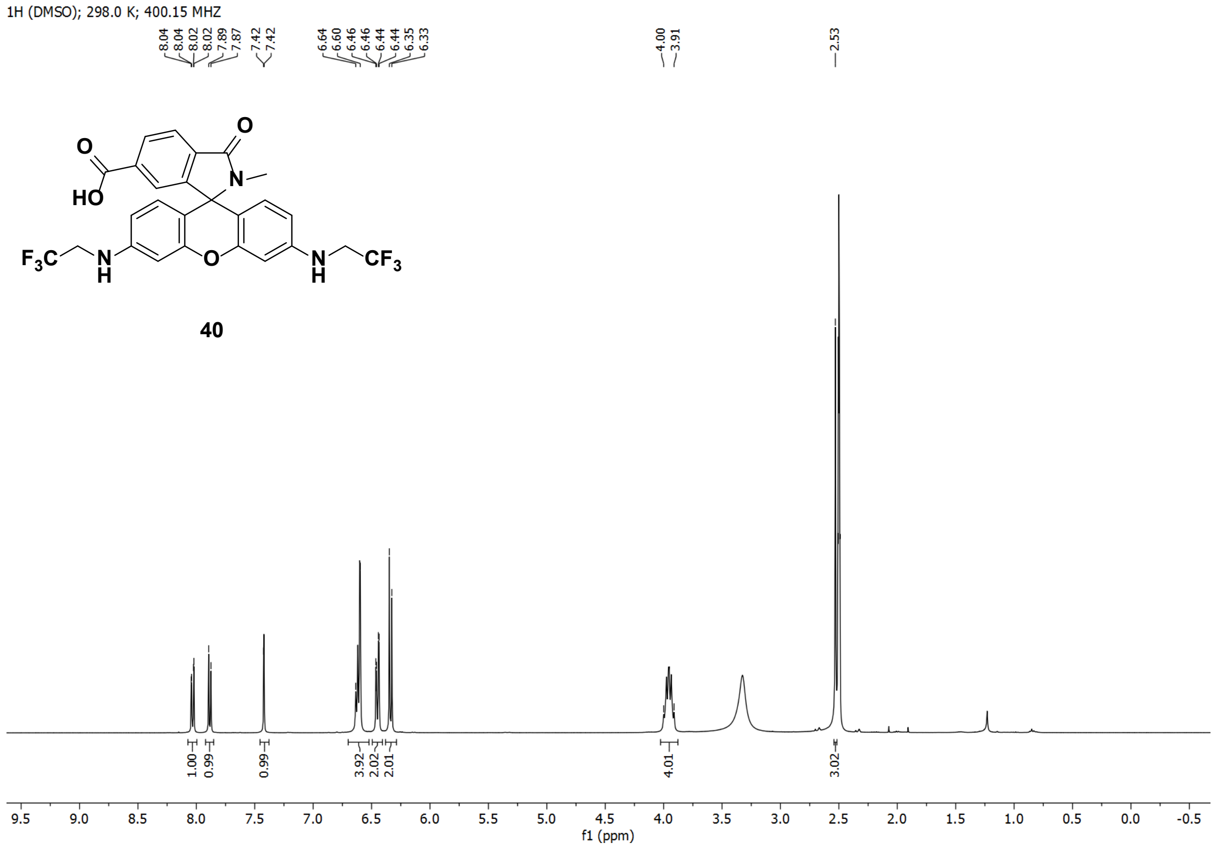
1H (DMSO); 298.0 K; 400.15 MHZ
8.04
8.04
8.02
8.02
7.89
7.87
7.42
7.42
6.64
6.60
6.46
6.46
6.44
6.44
6.35
6.33
4.00
3.91
2.53
40
1.00
0.99
0.99
3.92
2.02
2.01
4.01
3.02
f1 (ppm)

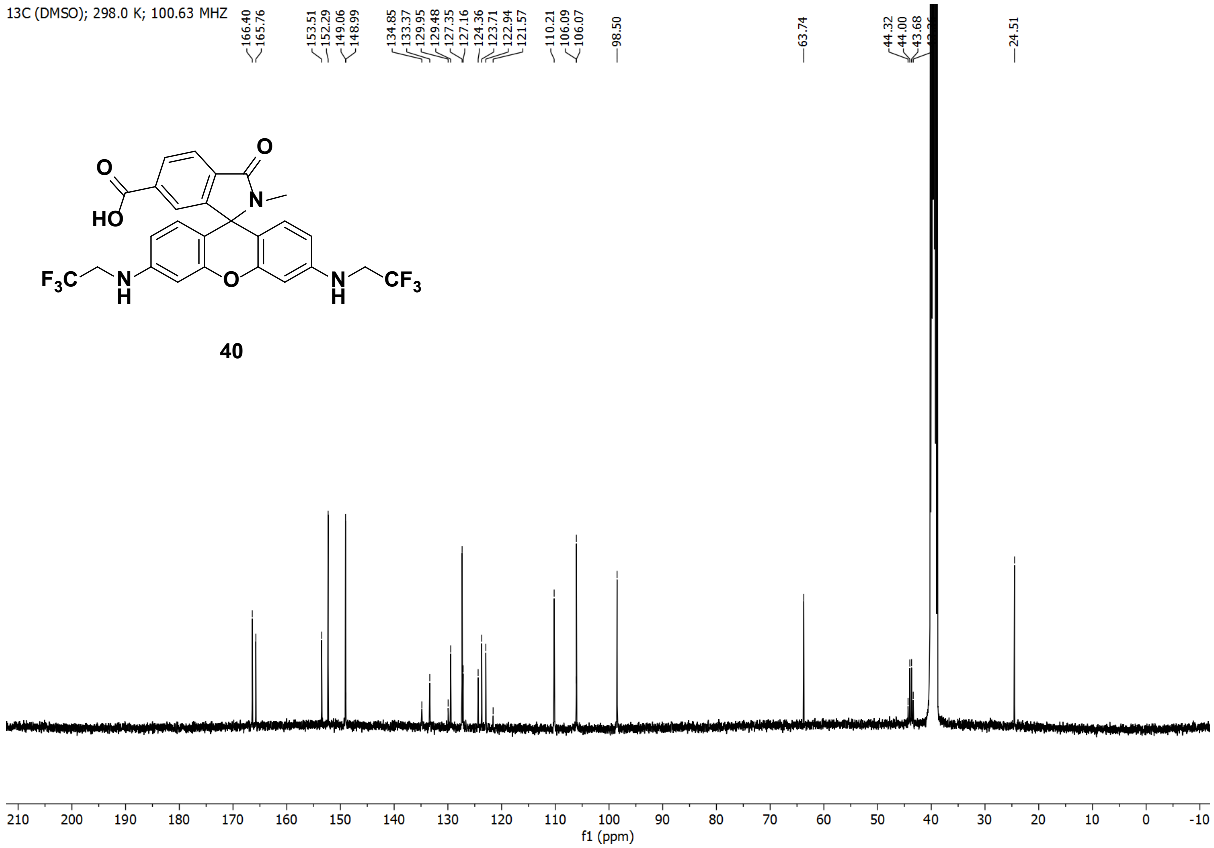
13C (DMSO); 298.0 K; 100.63 MHZ
166.40
165.76
153.51
152.29
149.06
148.99
134.85
133.37
129.95
129.48
127.35
127.16
124.36
123.71
122.94
121.57
110.21
106.09
106.07
98.50
63.74
44.32
44.00
43.68
24.51
O
HO
N
F3C
N
H
O
N
H
CF3
40
f1 (ppm)

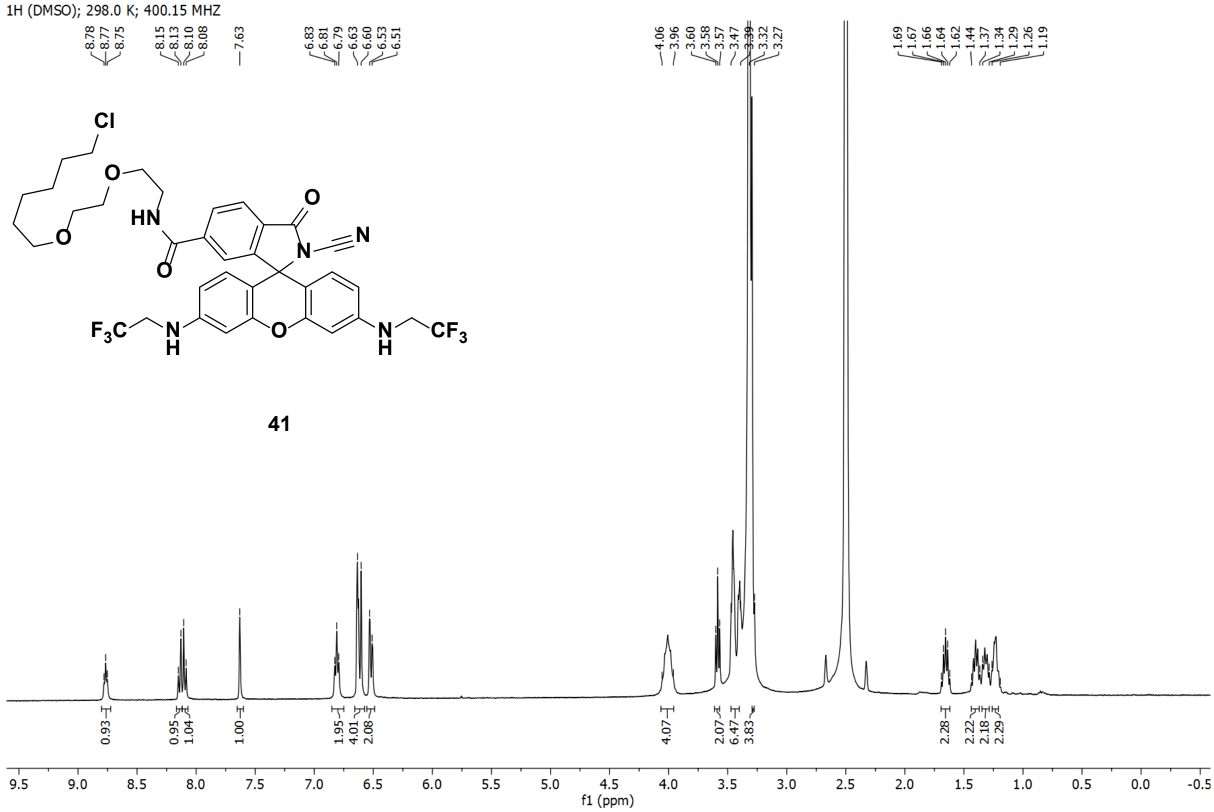
1H (DMSO); 298.0 K; 400.15 MHZ
8.78
8.77
8.75
8.15
8.13
8.10
8.08
7.63
6.83
6.81
6.79
6.63
6.60
6.53
6.51
4.06
3.96
3.60
3.58
3.57
3.47
3.39
3.32
3.27
1.69
1.67
1.66
1.64
1.62
1.44
1.37
1.34
1.29
1.26
1.19
Cl
O
O
HN
O
O
N
N
F3C
N
H
O
N
H
CF3
41
0.93
0.95
1.04
1.00
1.95
4.01
2.08
4.07
2.07
6.47
3.83
2.28
2.22
2.18
2.29
f1 (ppm)

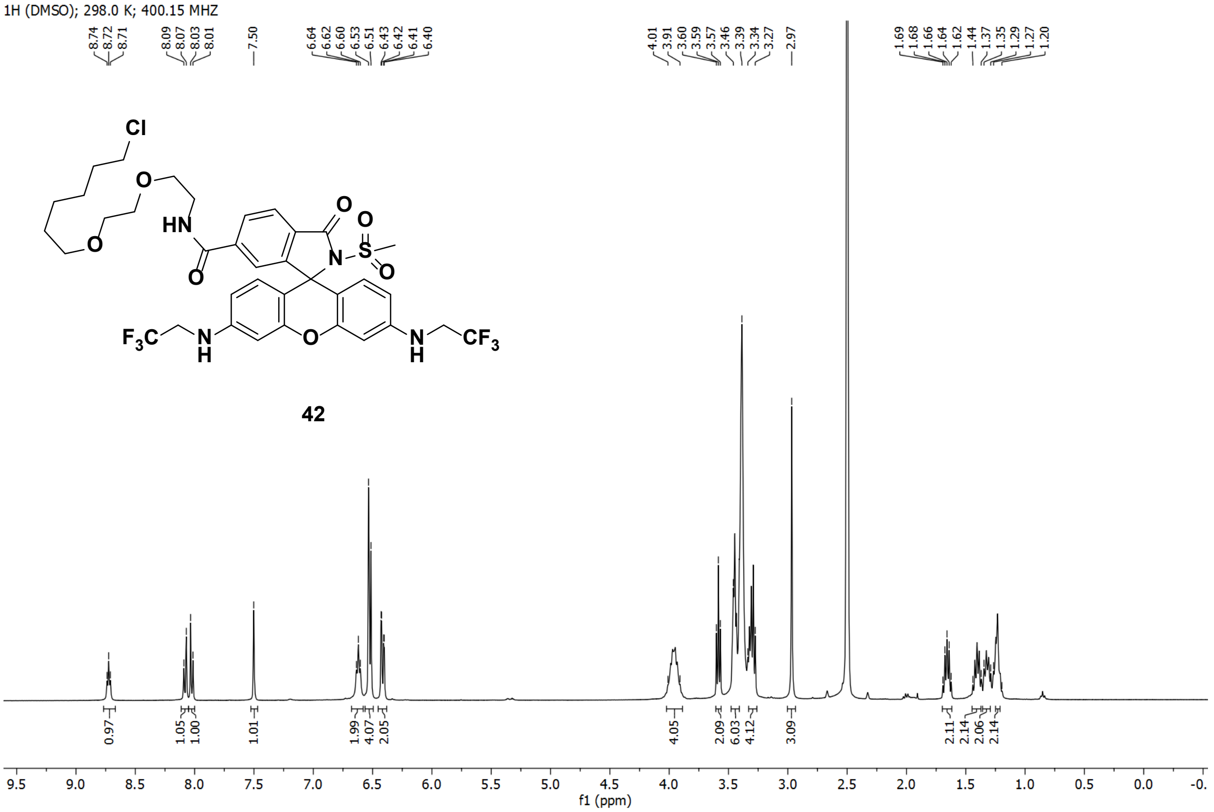
1H (DMSO); 298.0 K; 400.15 MHZ
8.74 8.72 8.71 8.09 8.07 8.03 8.01 7.50 6.64 6.62 6.60 6.53 6.51 6.43 6.42 6.41 6.40
4.01 3.91 3.60 3.59 3.57 3.46 3.39 3.34 3.27 2.97
1.69 1.68 1.66 1.64 1.62 1.44 1.37 1.35 1.29 1.27 1.20
Cl
O
HN
O
O
O
N
S
O
O
F3C
N
H
O
N
H
CF3
42
0.97 1.05 1.00 1.01 1.99 4.07 2.05 4.05 2.09 6.03 4.12 3.09 2.11 2.14 2.06 2.14
9.5 9.0 8.5 8.0 7.5 7.0 6.5 6.0 5.5 5.0 4.5 4.0 3.5 3.0 2.5 2.0 1.5 1.0 0.5 0.0
f1 (ppm)

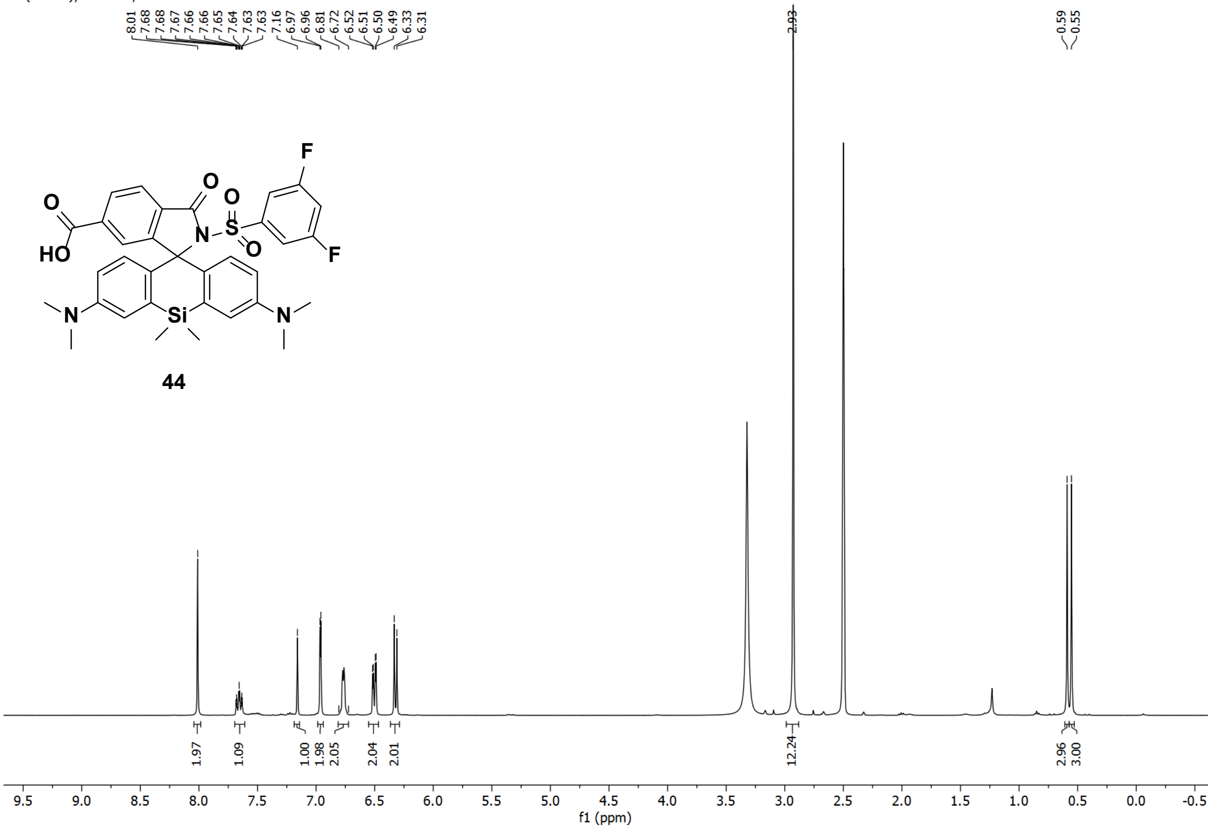
1H (DMSO); 298.0 K; 400.15 MHZ
8.01 7.68 7.68 7.67 7.66 7.65 7.64 7.63 7.63 7.16 6.97 6.96 6.81 6.72 6.52 6.51 6.50 6.49 6.33 6.31
2.93
0.59 0.55
F
O
O
O
HO
N
S
O
F
N
Si
N
44
1.97 1.09 1.00 1.98 2.05 2.04 2.01 12.24 2.96 3.00
9.5 9.0 8.5 8.0 7.5 7.0 6.5 6.0 5.5 5.0 4.5 4.0 3.5 3.0 2.5 2.0 1.5 1.0 0.5 0.0 -0.5
f1 (ppm)

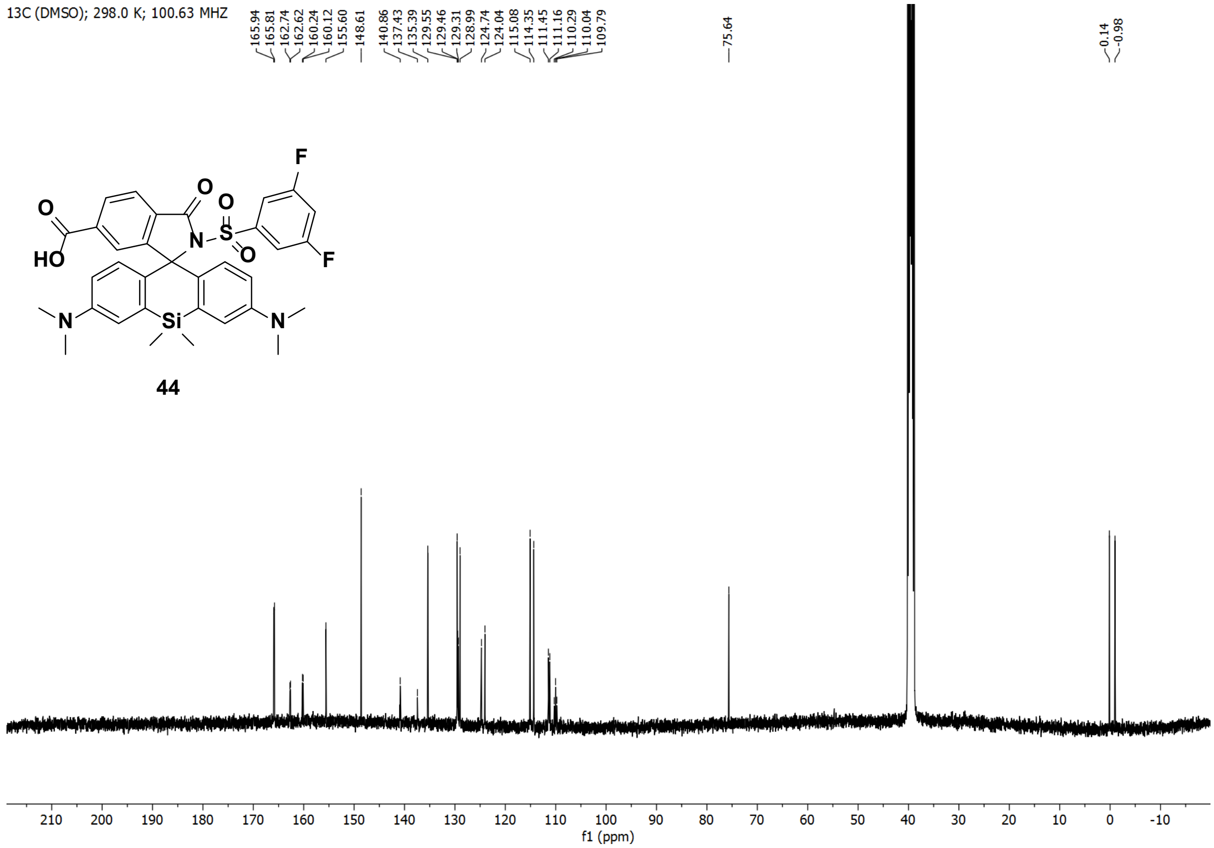
13C (DMSO); 298.0 K; 100.63 MHZ
165.94
165.81
162.74
162.62
160.24
160.12
155.60
148.61
140.86
137.43
135.39
129.55
129.46
129.31
128.99
124.74
124.04
115.08
114.35
111.45
111.16
110.29
110.04
109.79
75.64
0.14
-0.98
44
f1 (ppm)

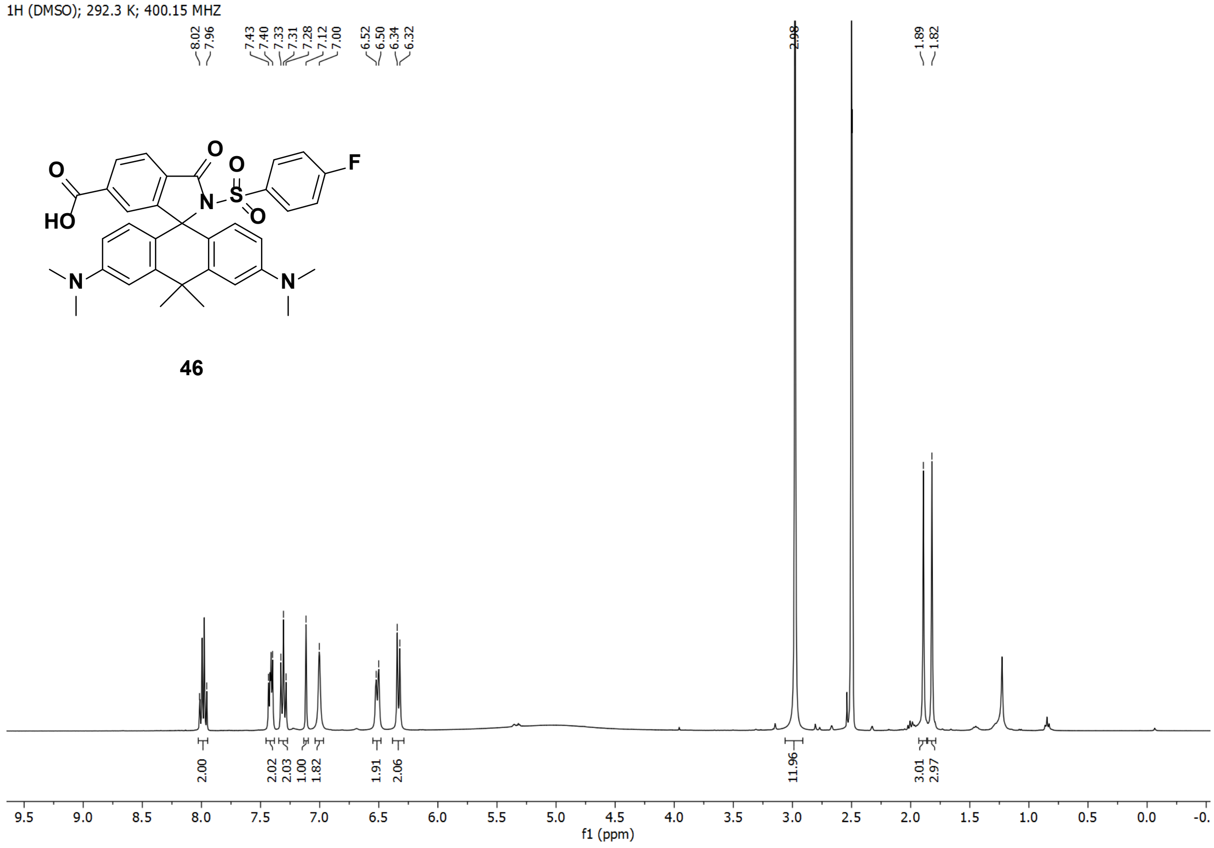
1H (DMSO); 292.3 K; 400.15 MHZ
8.02
7.96
7.43
7.40
7.33
7.31
7.28
7.12
7.00
6.52
6.50
6.34
6.32
2.98
1.89
1.82
46
2.00
2.02
2.03
1.00
1.82
1.91
2.06
11.96
3.01
2.97
f1 (ppm)

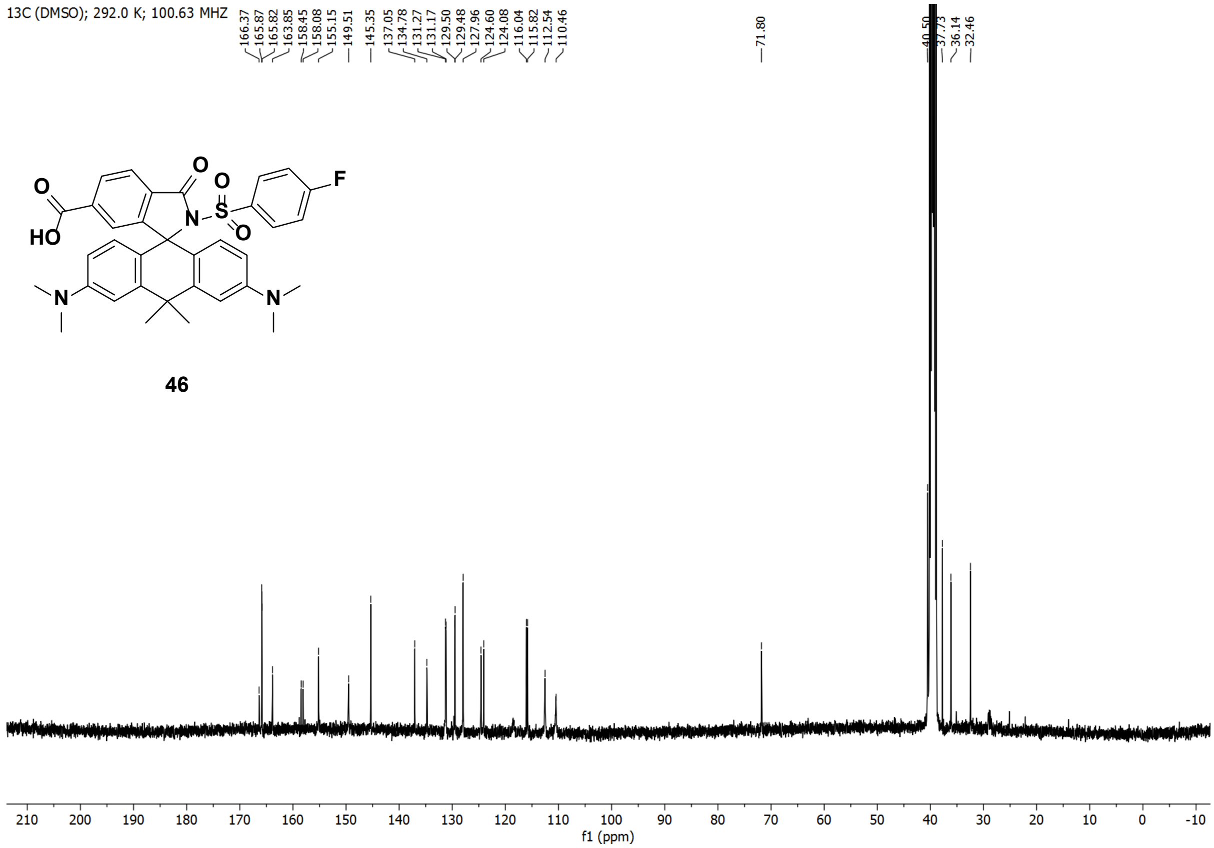

13C (DMSO); 292.0 K; 100.63 MHZ
166.37
165.87
165.82
163.85
158.45
158.08
155.15
149.51
145.35
137.05
134.78
131.27
131.17
129.50
129.48
127.96
124.60
124.08
116.04
115.82
112.54
110.46
71.80
40.50
37.73
36.14
32.46
f1 (ppm)


**46**

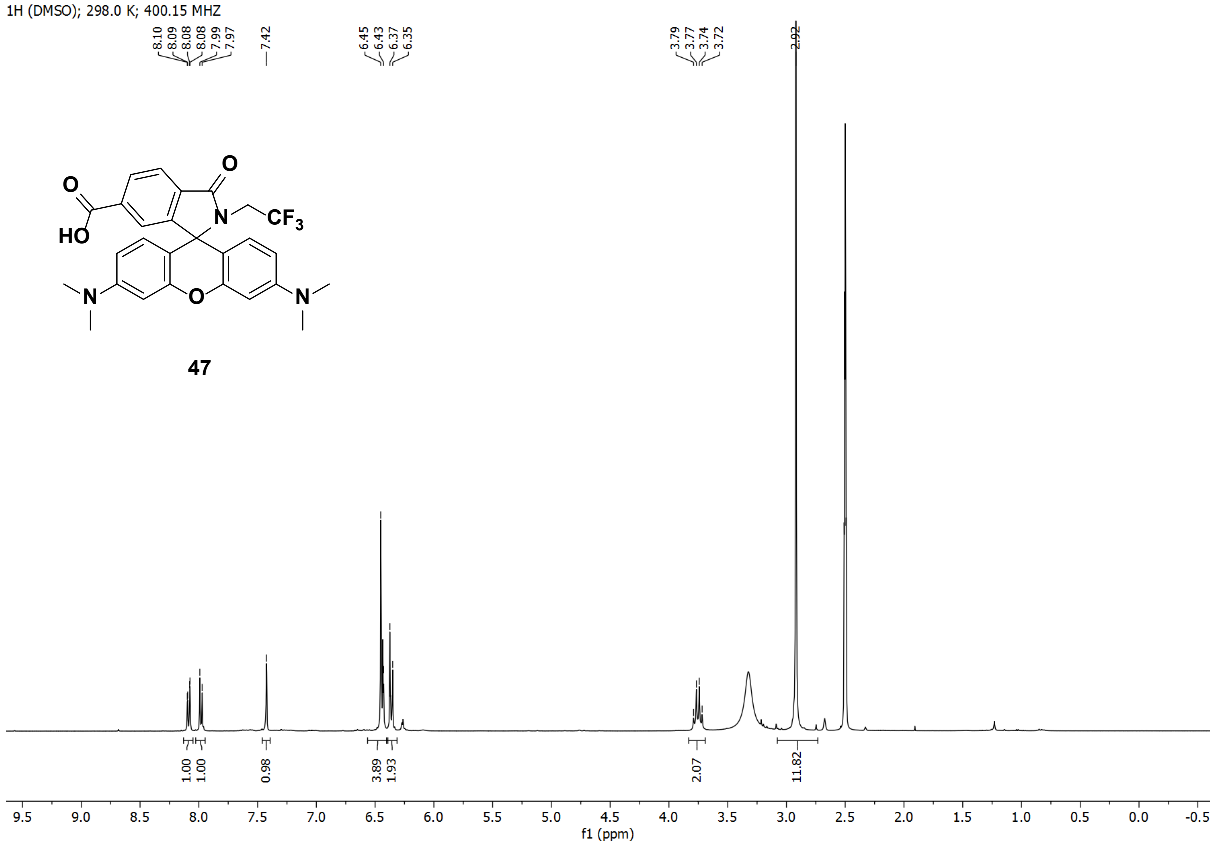

1H (DMSO); 298.0 K; 400.15 MHZ
8.10
8.09
8.08
8.08
7.99
7.97
7.42
6.45
6.43
6.37
6.35
3.79
3.77
3.74
3.72
2.92
1.00
1.00
0.98
3.89
1.93
2.07
11.82
f1 (ppm)


**47**

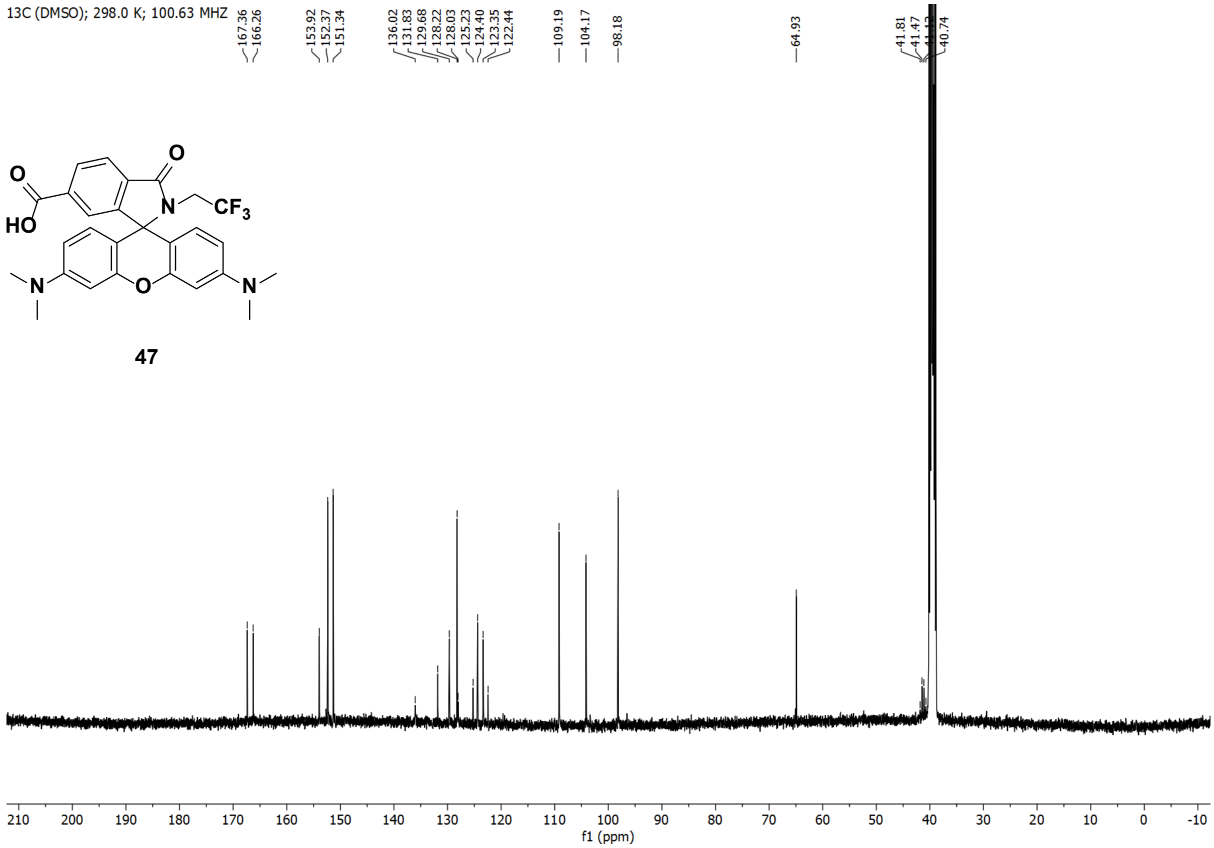
13C (DMSO); 298.0 K; 100.63 MHZ
47
f1 (ppm)

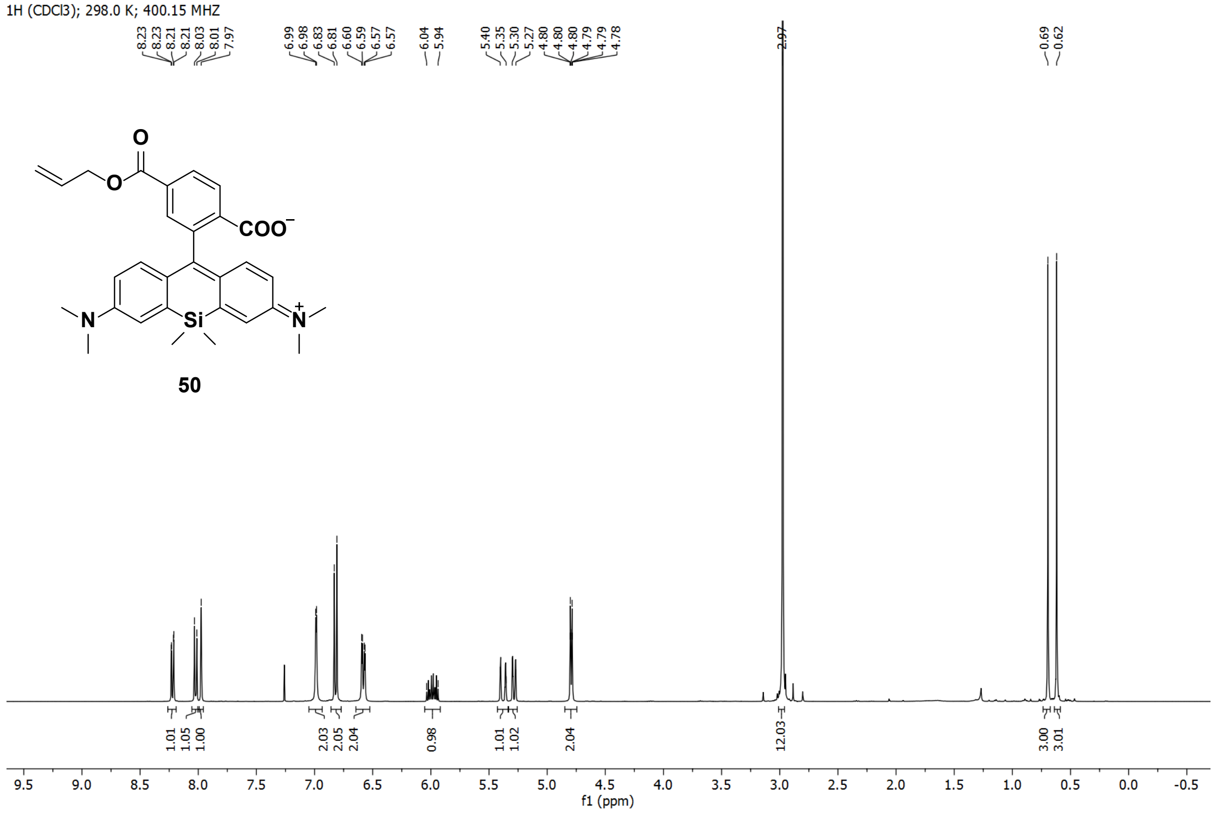
1H (CDCl3); 298.0 K; 400.15 MHZ
50
f1 (ppm)

13C (CDCl3); 298.0 K; 100.63 MHZ

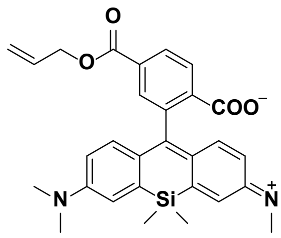


50

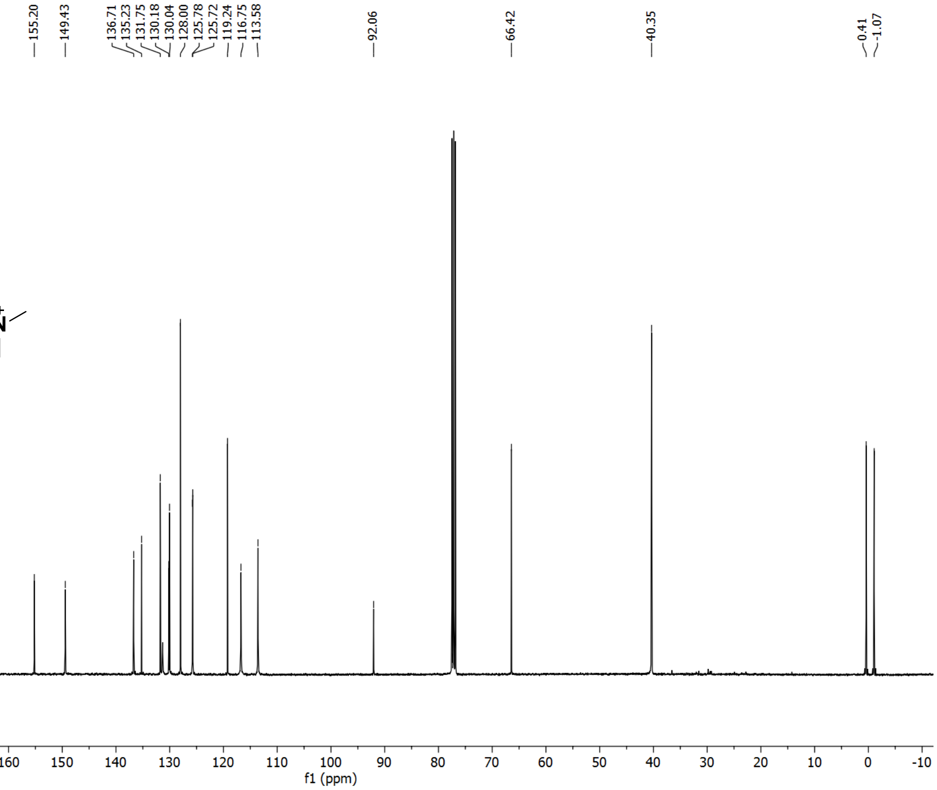
